# Transforming descending input into motor output: An analysis of the *Drosophila* Male Adult Nerve Cord connectome

**DOI:** 10.1101/2023.06.07.543976

**Authors:** H. S. J. Cheong, K. Eichler, T. Stürner, S. K. Asinof, A. S. Champion, E. C. Marin, T. B. Oram, M. Sumathipala, L. Venkatasubramanian, S. Namiki, I. Siwanowicz, M. Costa, S. Berg, Janelia FlyEM Project Team, G. S. X. E. Jefferis, G. M. Card

## Abstract

In most animals, a relatively small number of descending neurons (DNs) connect higher brain centers in the animal’s head to circuits and motor neurons (MNs) in the nerve cord of the animal’s body that effect movement of the limbs. To understand how brain signals generate behavior, it is critical to understand how these descending pathways are organized onto the body MNs. In the fly, *Drosophila melanogaster*, MNs controlling muscles in the leg, wing, and other motor systems reside in a ventral nerve cord (VNC), analogous to the mammalian spinal cord. In companion papers, we introduced a densely-reconstructed connectome of the *Drosophila* Male Adult Nerve Cord (MANC, (Takemura et al., 2024)), including cell type and developmental lineage annotation (Marin et al., 2024), which provides complete VNC connectivity at synaptic resolution. Here, we present a first look at the organization of the VNC networks connecting DNs to MNs based on this new connectome information. We proofread and curated all DNs and MNs to ensure accuracy and reliability, then systematically matched DN axon terminals and MN dendrites with light microscopy data to link their VNC morphology with their brain inputs or muscle targets. We report both broad organizational patterns of the entire network and fine-scale analysis of selected circuits of interest. We discover that direct DN-MN connections are infrequent and identify communities of intrinsic neurons linked to control of different motor systems, including putative ventral circuits for walking, dorsal circuits for flight steering and power generation, and intermediate circuits in the lower tectulum for coordinated action of wings and legs. Our analysis generates hypotheses for future functional experiments and, together with the MANC connectome, empowers others to investigate these and other circuits of the *Drosophila* ventral nerve cord in richer mechanistic detail.

## 1 Introduction

The nerve cord (e.g. spinal cord in mammals or ventral nerve cord in arthropods) is a vital part of the central nervous system in many animals. It receives input both from the brain and from sensors of the body, and integrates these signals to drive local circuits that control motor output to the body and limbs. It also contains ascending neurons that return information about ongoing circuit activity to the brain. Circuits that directly or indirectly control motor output (herein ‘premotor circuits’) in the nerve cord thus must perform many tasks, including decoding descending commands from the brain, locally modulating motor output based on immediate incoming sensory information, controlling the muscles of individual limbs and body parts, and coordinating movements across multiple body parts with millisecond precision. The organizational logic of the individual circuits that accomplish these feats remains largely unknown. In Drosophila, the ventral nerve cord (VNC) carries out these functions to control the fly’s neck, wings, halteres, legs, and abdomen (Figure 1A). Here, we present a system-wide look at the structure of premotor circuits by using the fly Male Adult Nerve Cord (MANC) connectome to examine patterns of synaptic connectivity in the networks connecting brain input, carried by descending neurons (DNs), to motor output, transmitted by motor neurons (MNs).

**Figure 1:**
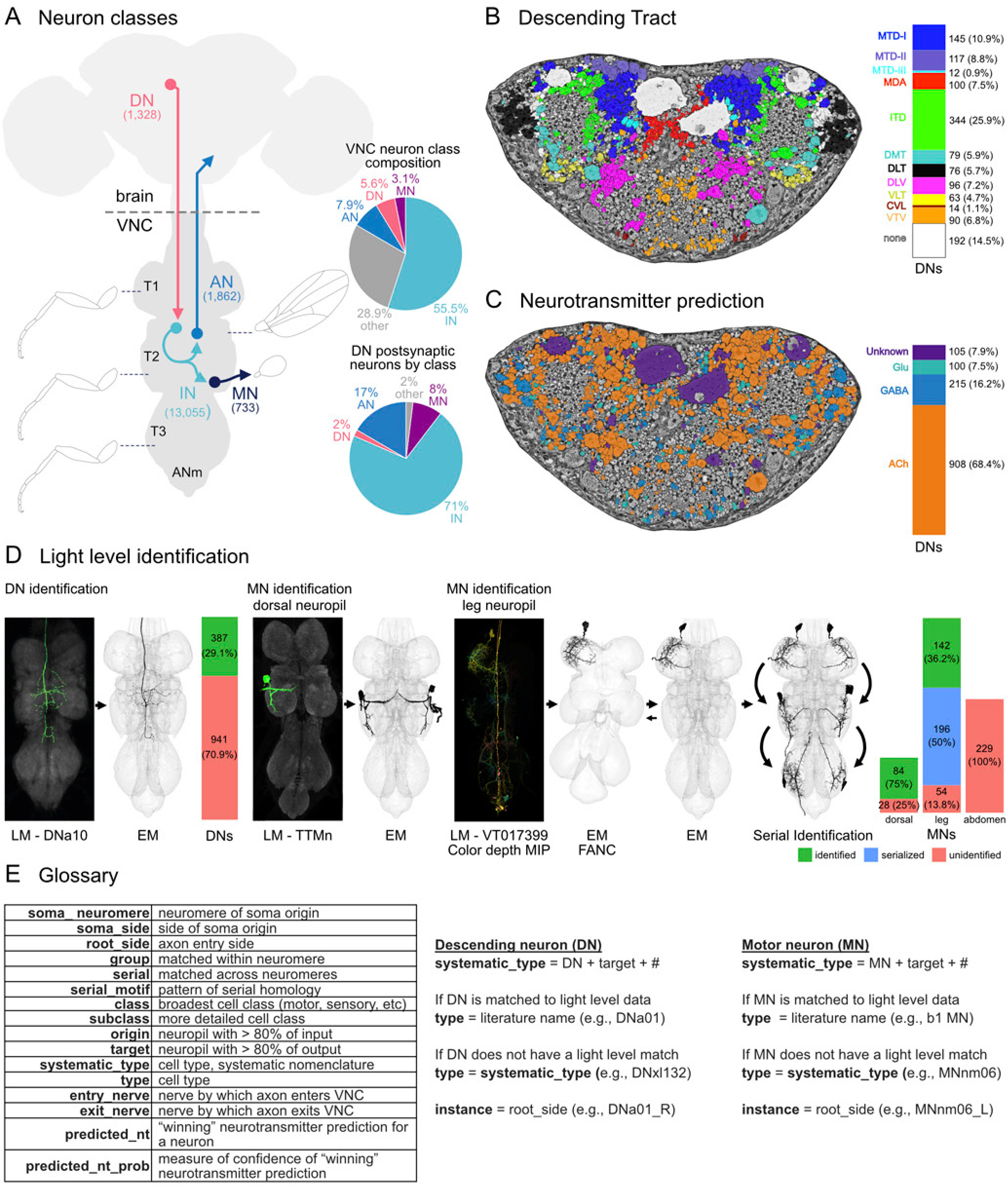
Overview of MANC connectome. **A.** Schematic of the *Drosophila melanogaster* central nervous system, showing brain and ventral nerve cord (VNC) as well as neuron classes which constitute VNC motor circuits: descending neurons (DNs), intrinsic neurons (INs), motor neurons (MNs), and ascending neurons (ANs). Numbers in parentheses indicate the total number of that neuron class found in MANC. Approximate areas indicated by dashed lines correspond to the neuropils controlling several of the fly’s primary motor systems (legs, wings, halteres). The three thoracic segments are indicated with T1, T2 and T3, and the abdominal neuropil is abbreviated ANm. Pie charts show the neuron class compositions of the entire VNC, as well as of postsynaptic partners of all DNs, the majority of which are intrinsic neurons. **B-C.** Cross section of the *D. melanogaster* neck connective at the location of the black dashed line in **A**. DN profiles are color coded by tract membership (**B**, see Figure 2) and predicted neurotransmitter (**C**, see Figure 4). Side bar charts show distribution of DNs per tract or predicted neurotransmitter. Predicted neurotransmitter type is ‘unknown’ if prediction probability was <0.7). **D.** DNs and MNs were identified by matching MANC EM reconstructed neurons to published light microscopic (LM) level descriptions of DNs (mainly (Namiki et al., 2018), for more references see Supplementary file 1) and MNs either manually by eye or aided by neuronbridge queries of color depth maximum intensity projections (MIP) (Clements et al., 2022; Otsuna et al., 2018) of driver lines against the EM dataset. Left, DN identification example: A neuronbridge query for SS02384, which labels DNa10 (Namiki et al., 2018), revealed group 10506 to be a good match. 29.1% of MANC DNs matched a previous DN description in the literature. Center, MN identification example for the dorsal neuropils. Matching to SS02623 (which contains the tergotrochanteral muscle motor neuron, TTMn) revealed group 10068. Right, MN identification example for the leg neuropils. A recent study identified all leg MNs in the T1 compartment (FANC, (Azevedo et al., 2020). T1 leg MNs in this dataset (MANC) were matched to FANC using NBLAST. Cross-matching of leg MNs across FANC and MANC aided in confirming LM identification of leg MNs in both volumes. Furthermore, serially homologous neurons of T1 leg MNs in T2 and T3 were identified in MANC (see Materials & Methods). The bar chart shows the status of LM identification of MNs. 75% of dorsal neuropil MNs are matched to previous descriptions. 35.9% of leg MNs are matched to FANC. By identifying serial homologous neurons of the T1 leg MNs, the muscle targets of an additional 50% of leg MNs were characterized. We did not match abdominal MNs to light level descriptions in this study. Images of SS02384 and SS02623 are from (Namiki et al., 2018), and Color depth MIP of the VT017399 driver are from (Otsuna et al., 2018). **E.** Left, glossary of terms and acronyms used in this study. Right, systematic naming of DNs and MNs. For more information on other neuron classes and terms in MANC, see (Marin et al., 2024).

**Figure 1—Supplement 1:**
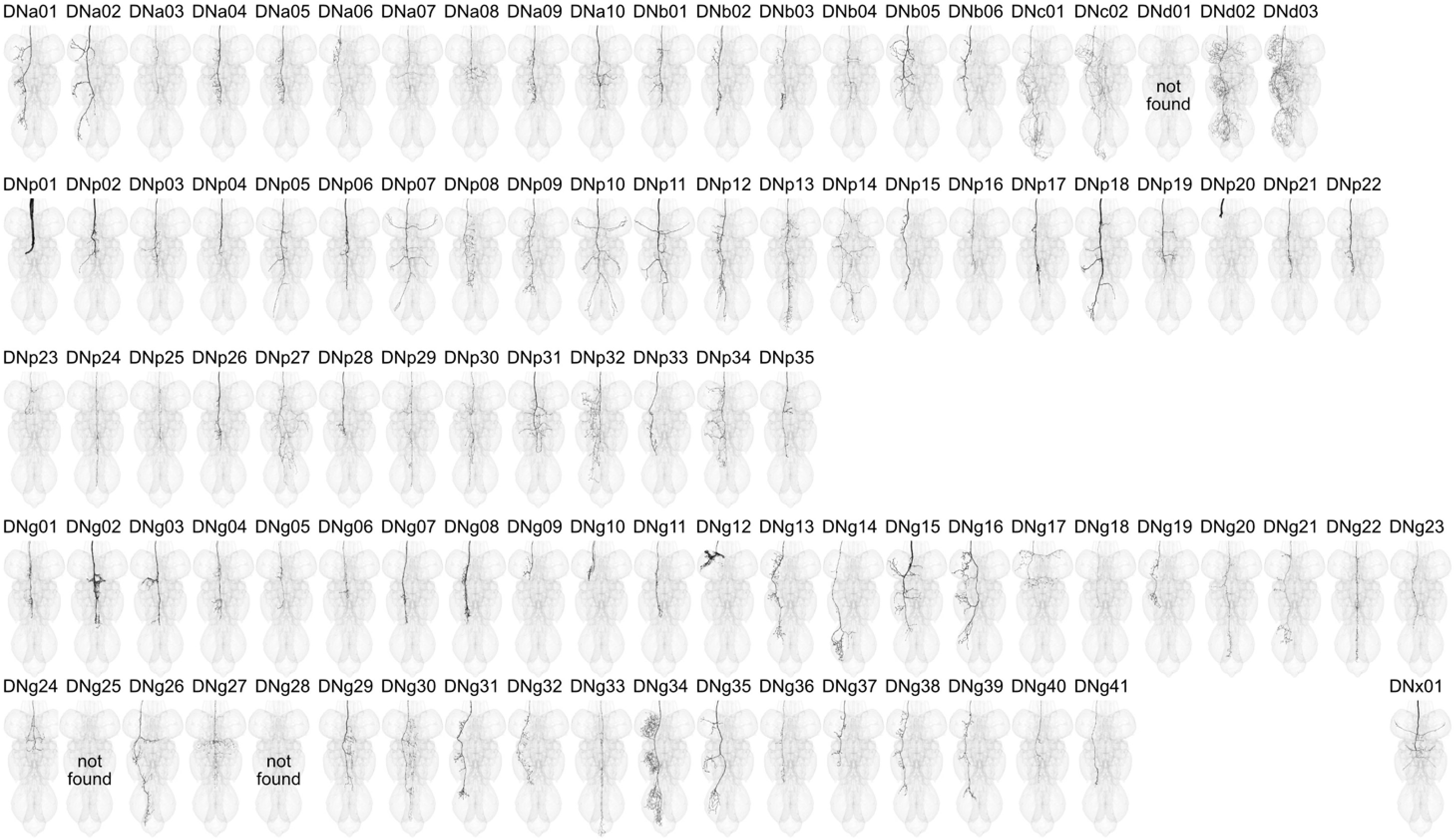
Overview of Identified DNs. Morphology of DN types described by (Namiki et al., 2018) identified in the EM volume. Nomenclature according to (Namiki et al., 2018). Three DN types could not be identified in the EM dataset (indicated by an empty VNC outline and “not found”). See Supplementary file 1 for more information on identification confidence and other synonyms of DNs in the literature.

DNs represent a bottleneck in the transmission of information between sensory processing areas in the brain and premotor circuits in the nerve cord. In the brain, DNs chiefly receive input from several brain regions: the lateral accessory lobe (LAL) and posterior slope (PS), which are regions implicated in navigation and visual motion processing, the posterior lateral and ventrolateral protocerebra (PLP, PVLP), regions implicated in escape and other rapid responses to salient visual stimuli, and the gnathal ganglion (GNG), a region involved in mechanosensory, gustatory and locomotor responses (Hsu and Bhandawat, 2016; Namiki et al., 2018). In the VNC, DNs innervate either VNC neuropil layers that contain wing/neck/haltere MNs or layers containing leg MNs, but rarely both (Namiki et al., 2018). A separate subset of DNs terminate in the posterior neuropil, where motor neurons controlling the abdominal muscles reside, and control abdominal motor behavior such as copulation (Pavlou et al., 2016) and oviposition (Castellanos et al., 2013). Thus, most DNs carry information specific to a particular motor system. One exception is a subset of DNs that targets intermediate VNC layers, which are largely devoid of MNs but likely coordinate behaviors requiring both legs and wings, such as takeoff (Court et al., 2020; Namiki et al., 2018). Previous studies estimated the number of DNs to be roughly 1000, which is 1-2 orders of magnitude fewer than the number of neurons in the brain or VNC networks that they connect to (Hsu and Bhandawat, 2016; Namiki et al., 2018). The brain thus provides input to the VNC in the form of a relatively compressed code across a small number of neurons targeted to specific motor systems.

Many DN types initiate motor activity when activated. In some cases, DN activity triggers clear motor programs such as walking, flight, takeoff, or singing (Bidaye et al., 2014; Cande et al., 2018; Guo et al., 2022; Koto et al., 1981; McKellar et al., 2019; Rayshubskiy et al., 2020; von Philipsborn et al., 2011), while in other cases it modulates ongoing motor activity (Aymanns et al., 2022; Namiki et al., 2022). Some DN types produce uncoordinated actions when optogenetically activated (Cande et al., 2018), suggesting that some behavioral patterning requires the simultaneous activation of multiple DNs or DN activation during the correct internal state. Indeed, the combined action of several pairs of DNs controls distinct leg gestures during walking steering maneuvers (Yang et al., 2023). Further supporting DN population encoding of behavior, silencing DNs often does not eliminate the action that activation of the same DN initiates (Ache et al., 2019). Emerging evidence also suggests that many DNs that act like command neurons can recruit additional sets of downstream DNs when activated, which may aid in carrying out flexible behavioral output (Braun et al., 2024). Thus, a reasonable framework for DN input to the VNC based on current knowledge is that DN activity represents higher-order behavioral modules carried by individual DNs acting more like “command neurons’’ or by groups or small populations of DNs, and both categories of DNs can further interact to produce flexible behavior patterns.

On the motor output end, MNs in the fly VNC support five primary motor systems: the wings, legs, neck, halteres, and abdomen. By patterning activity onto MNs in these systems, VNC premotor circuits are able to control two distinct forms of locomotion (walking and flying) and a wide array of behaviors that combine actions from multiple motor systems (e.g. escape, grooming, courtship, aggression and oviposition). To understand how VNC premotor circuits create these diverse behaviors, it is necessary to consider the biomechanics of each motor system being controlled. In particular, wing and leg motor systems have very different mechanical arrangements, which may suggest that the structure of their underlying control circuits also differ greatly.

In dipterans such as *Drosophila*, the wing motor system includes three classes of muscles: power, steering, and indirect control muscles (Dickinson and Tu, 1997). The power muscles are divided into two types that attach orthogonally relative to each other across the thorax, such that when one set contracts it deforms the thoracic exoskeleton and stretches the other. The muscles themselves are stretch-activated, meaning that once started, the power muscles alternately activate each other, setting up the rhythmic deformations of the thorax that flap the wings, providing power during flight and the primary oscillations used in courtship song (Dickinson and Tu, 1997; Ewing, 1977). Power muscle oscillations will continue as long as activity in MNs continues to provide an influx of calcium to the muscles. However, MN activity is asynchronous and does not need to occur at the frequency of the wingstroke (Dickinson and Tu, 1997). Steering is accomplished through synchronous and direct control of wing hinge elements by the steering muscles, a series of 14 muscles that attach directly to three of the four axillary sclerites that form the wing hinge (the first, third and fourth axillary sclerites: i, iii, and hg) and the basalare sclerite (b) anterior of the wing hinge (Deora et al., 2017; Dickinson and Tu, 1997). Timing is critical for control of these muscles, as the orientation and movement direction of the wing is determined by the phase of the wingstroke at which they are activated (Egelhaaf, 1989; Götz, 1983; Heide, 1983, 1975; Heide and Götz, 1996). Finally, of the indirect control muscles, the pleurosternal (ps) and tergopleural (tp) muscles control thoracic stiffness and resonant properties, thereby indirectly modulating wingbeat amplitude (Dickinson and Tu, 1997), while tergotrochanteral muscle (TTM) activation rapidly depresses the middle legs and ‘jump-starts’ the power muscles during escape takeoffs (Dickinson and Tu, 1997).

In contrast, the leg motor system of the fly includes six separate jointed appendages, each of which is composed of 10 segments (coxa, trochanter, femur, tibia, five tarsal segments and the claw or pretarsus). However, only the proximal joints are “true” joints that can be actively moved, as there are no muscles internal to the tarsal segments. The musculature of the *Drosophila* leg consists of 13 muscle groups confined within the proximal leg segments, and another five in the thorax that insert in the leg (Azevedo et al., 2024; Brierley et al., 2012). These leg muscles are estimated to be innervated by around 70 MNs in each leg, that originate from ∼15 hemilineages (Baek and Mann, 2009; Brierley et al., 2012; Phelps et al., 2021). Muscles in the leg are not innervated uniformly; indeed, the muscles of the T1 legs differ in the the number of MNs innervating each muscle by as much as an order of magnitude (Azevedo et al., 2024; Baek and Mann, 2009). Individual MNs also exhibit considerable heterogeneity in their intrinsic properties, correlating with the speed and force with which they activate downstream muscles as well as the order in which they are recruited by afferents (Azevedo et al., 2024, 2020). Complex movements require the coordinated control of multiple muscles, such as the activation of one muscle at the same time as the deactivation of its opposing counterpart. Premotor circuits that produce fluid patterns of motion in a single leg (e.g. repeated cycles of swing and stance associated with walking forwards) must therefore be capable of activating or suppressing tens of MNs in an appropriate sequence to actuate movement at each of the fly’s leg joints.

Compared to the wing and leg motor systems, less is known about the neck, haltere, and abdominal motor systems. As abdominal MNs in MANC, while reconstructed and grouped, still require future identification efforts to determine their muscle targets, we do not focus on the abdominal motor system in this paper. Neck muscles serve to move the fly’s head, which is critical for gaze-stabilization as *Drosophila* are highly visual animals, yet have compound eyes fused to their head. So, with the exception of small retinal movements (Fenk et al., 2022), neck muscles are responsible for minimizing motion blur during flight maneuvers. The neck muscles and MNs are not yet well-described in *Drosophila*, but a comprehensive description of the neck motor system is available in the blowfly, *Calliphora erythrocephala* (Strausfeld et al., 1987). Of note, fly neck MNs are split between those residing in the VNC and the brain, and so the neck MNs are not complete in the MANC dataset. The halteres are appendages derived from hindwings that in *Drosophila* beat antiphase to the wings during flight. They are equipped with sensors at their base that allow them to detect Coriolis and oscillatory inertial forces, and hence provide fast mechanosensory feedback to the flight system to enable rapid maneuvering. Similar to the wing system, the haltere motor system consists of a reduced set of eight muscles that are homologous to the wing muscles, divided into a single power muscle and seven steering muscles (Dickerson et al., 2019). Movements of the neck and halteres are tightly coordinated with the wings during flight through both mechanical linkages and neural control (Deora et al., 2021, 2015; Dickerson et al., 2019). As such, during the flight state, they may be considered to be under the control of a single “flight” motor system, as suggested by the fusion of neck, wing, and haltere neuropils into a single dorsal “upper tectulum” layer in the VNC.

VNC motor control circuits are further patterned by developmental identity. In the CNS, a fixed set of neuroblasts (stem cells) generate neurons in stereotyped lineages. Single hemilineages often give rise to neurons which are implicated in a single or related set of motor functions (Harris et al., 2015) and in many cases are capable of semi-autonomous patterned motor output. Indeed, experiments with decapitated flies show that the VNC local circuitry can, with mechanical or optogenetic stimulation (or in some cases spontaneously), generate the appropriate signals for patterned motor activities including walking, flight, grooming, and courtship song (Clyne and Miesenböck, 2008; Gao et al., 2013; Harris et al., 2015; Vandervorst and Ghysen, 1980). This arrangement could allow signals from the brain to act in a command or modulatory manner on motor output, independently of the details of the desired patterned motor output.

Despite decades of investigation into DNs, VNC circuits, and their role in behavior, only small portions of VNC circuitry and specific pathways are well understood, and we lack both large- and fine-scale understanding of circuit connectivity and organization across the VNC. In recent years, electron microscopy-based (EM) connectomics has emerged as a powerful tool to dissect and understand neural circuitry, enabling the reconstruction of neurons from EM imagery and quantification of synapse connectivity strength, thus giving us a direct method to analyze neural pathways at synaptic resolution. Several large connectomics efforts have been undertaken in *Drosophila* including the larval CNS (Ohyama et al., 2015; Winding et al., 2023), Female Adult Fly Brain (Dorkenwald et al., 2022; Zheng et al., 2018), Hemibrain (Scheffer et al., 2020), and Female Adult Nerve Cord (Phelps et al., 2021), which are further enhanced by development of bioinformatics tools for analyzing neuroanatomical and connectomics data (e.g. Bates et al., 2020). Of note, the Female Adult Nerve Cord (FANC) (Phelps et al., 2021) is the first complete EM volume of the adult *Drosophila* VNC, providing valuable insight into motor neuron morphology, count and organization in the VNC thoracic segments, as well as allowing labs to further reconstruct and investigate VNC circuits of interest. More recently, work based on proofreading an automated segmentation of the FANC volume has identified all T1 leg MNs and the majority of wing MNs in EM by matching to light-level data (Azevedo et al., 2024). Concurrent to the work presented here,efforts in FANC have further reconstructed and analyzed the connectivity of the direct upstream partners of these MNs, giving us insights into synapse-resolution connectivity and organization of the first layer of premotor neurons upstream of motor output (Lesser et al., 2024).

In the Hemibrain project, advances in sample preparation, EM methods and acquisition, image stack alignment, autosegmentation, synapse prediction, and proofreading software and methods, in combination with efforts of a dedicated proofreading team, made practical the production of large densely reconstructed connectomes where users need only query neurons or circuits of interest through a web or software interface (Plaza et al., 2022; Scheffer et al., 2020). Using these advances, the Janelia FlyEM project team has now produced the *Drosophila* Male Adult Nerve Cord (MANC) connectome (Takemura et al., 2024), a densely reconstructed connectivity map of the entire fly VNC. Proofreading and annotation teams have further added a rich set of metadata to all neurons of the VNC, describing neuron properties including (but not limited to) neuron class, hemilineage, morphological groupings, serial homology, and systematic typing based on morphology and connectivity (Marin et al., 2024; Takemura et al., 2024) The dense reconstruction and neuron annotations of the MANC connectome thus expedites the efforts of the enterprising neurobiologist to mine the connectome for neurons or circuits of interest. This newly available resource complements the existing female VNC connectome (Phelps et al., 2021) with a male counterpart, and its dense reconstruction permits circuit exploration without requiring labs to engage in manual reconstruction or proofreading of neurons.

Here, we carry out focused proofreading and annotation of all DNs and MNs in the MANC connectome, and match their cell types to published light-level imagery when possible. Matching of DNs to additional newly generated light level imagery is separately described in (Stürner et al. 2024). We then broadly survey DN-to-MN connectivity with system-wide analyses examining feedforward information flow from descending input to motor output. We further specifically investigate motor circuit organization in the wing tectulum, leg neuropils, and lower tectulum. Within these analyses, we focus on examining select sets of DNs with known or putative functions in locomotion, navigation and salient visual responses. We present a general overview of VNC premotor network organization as well as detailed connectivity of some of these behavioral circuits of interest.

## 2 Results

### 2.1 Connectome generation and cell typing

Creation and cell type annotation of the MANC connectome are necessary precursors to analysis of DN-to-MN pathways. These aspects of the project are presented in two companion papers: (Takemura et al., 2024), which describes generation of the MANC connectome, and (Marin et al., 2024), which describes cell type annotation. Briefly, MANC used an optimized connectome production pipeline consisting of a system of sample preparation, EM imaging, volume assembly (stitching), automated segmentation, and synapse prediction. Following initial segmentation of the EM volume, trained proofreaders manually examined the morphology of all putative VNC neurons, corrected major false merges, and connected major neurite segments. These efforts captured 42% of the overall synaptic connectivity in the VNC, ranging from 33% to 62% for individual neuropils. MANC synapse completeness is similar to prior reconstruction efforts in the Hemibrain which captured 39% of overall connectivity (ranging from 20% to 85% per neuropil), and is sufficient to capture the vast majority of strong synaptic connections (Scheffer et al., 2020). During proofreading, all MANC neurons were categorized into one of the following classes: intrinsic neuron (IN), descending neuron (DN), ascending neuron (AN), motor neuron (MN), efferent neuron (EN), efferent ascending (EA), sensory neuron (SN) and sensory ascending (SA). In total, ∼23,500 neurons were reconstructed and annotated in this dataset, which comprises 13066 INs, 1328 DNs, 1862 ANs, 733 MNs, 92 ENs, 9 EAs, 5927 SNs, and 535 SAs. All neurons were further categorized by morphology and synaptic connectivity into a ‘group’ comprising right-left homologous neurons, and, where applicable, into a ‘serial’ group that reflected the matching of putative serially homologous neurons across segments. A developmental ‘hemilineage’ was assigned to neurons with soma in the VNC. Neuron groups were then assigned ‘systematic type’ names based on connectivity and hemilineage identity for VNC intrinsic neurons, while DNs and MNs were separately assigned as explained in sections 2.2 and 2.3 (also see Figure 1E). If a neuron was previously identified in the literature, then its published name was defined as its ‘type,’ the primary name by which we refer to it in our analyses. Previously unidentified neurons retained their ‘systematic type’ as their ‘type’. In addition, automated synapse prediction achieved synapse retrieval comparable to human annotation. Finally, neurotransmitter type was predicted for all MANC neurons using a convolutional neural network (Eckstein et al., 2024), trained using 187 neurons with identified neurotransmitter types as ground truth, to assign a probability that a given neuron utilized one of the three major fast-acting neurotransmitters: acetylcholine (ACh), GABA or glutamate (Glu). Thus, MANC neuron reconstruction and synapse prediction efforts have produced a densely reconstructed connectome that allows quantitative analyses of synaptic connectivity across the VNC.

### 2.2 Identification of descending neurons

To systematically annotate all DNs in MANC, we first seeded every profile in a transverse plane through the neck connective (dashed line in Figure 1A) and proofread them to recognition of neuron class (AN, EA, SA, or DN). Classes were distinguished based on the following criteria: ANs and EAs both have a soma in the VNC, but ANs have only one axon branch through the neck connective, whereas EAs may send multiple processes through the neck connective and also enter VNC nerves; SAs and DNs both lack VNC somata, but SA axons enter the VNC through a nerve, whereas DNs enter through the neck connective. In addition to the above classes, we also identified four neck MNs running through the neck connective. Once identified, DNs were further annotated for the descending tract of their axon (Figure 1B, see below) and computer-vision-assigned neurotransmitter type (Figure 1C, see below).

As part of our focus on DN-to-MN pathways, we next proofread the DNs to higher completion in order to reach a greater coverage of presynaptic sites. To aid this process, we mapped all neurons from the left hand side of the dataset to the right using a mirroring registration, and then carried out NBLAST morphological clustering (Bates et al., 2020). This procedure naturally grouped neurons into either left-and-right pairs (59% of all DNs) or larger groups (populations, 39.4% of all DNs) of neurons. Furthermore, it provided a preliminary cell typing which was then used as the basis for comparing neurons with established light level data. We then manually matched DNs to light microscopy stacks (primarily from (Namiki et al., 2018)) showing expression patterns for genetic driver lines targeting single DN types (Figure 1D, Supplementary file 1). Matches were scored for confidence, and only those with high scores were assigned the literature names as their ‘type’. We were able to confidently identify matches for all but three types of published DNs (Figure 1–Supplement 1). This corresponds to 29% of all DNs (Figure 1D) in the EM dataset. Further matching of DN types to newly generated split-Gal4 lines or light level imagery are described in (Stürner et al. 2024), which extends DN identification to 45.9% of DNs.

To provide spatial characterization of each DN, we identified their longitudinal tract and neuropil innervation. DNs run in tightly bundled tracts through the neck connective before diverging to target different VNC neuropils. Previous work described eight main tracts (Court et al., 2020; Merritt and Murphey, 1992; Namiki et al., 2018). To identify the tracts for DNs in the MANC dataset, we took the longest neurite of each DN and performed an NBLAST clustering (Costa et al., 2016). Clusters were then manually compared with published tract morphologies and 3D-rendered tract volumes (meshes from Court et al. 2020, Figure 2A-C). We found that the median tract of the dorsal cervical fasciculus (MTD), which was previously described as containing two subgroups (MTD-I and MTD-II), had an additional subgroup in our dataset, which we named MTD-III. We also found that one of our NBLAST clusters did not match any previously reported tracts. DN axons in this cluster were positioned ventrolaterally but did not align with the ventral lateral tract (VLT). We named this new tract the curved ventral lateral tract (CVL) after its curved morphology. Both of these new tracts contain only a few DNs, which could explain their exclusion from previous light level studies (Figure 2D). In total, we assigned 85% of DNs in MANC to tracts. Some DNs could not be assigned to a tract as they traveled outside of all axon tracts and did not bundle with other DNs. We used these assignments to generate new tract volumes to be compared to future VNC datasets (Figure 2B). The Intermediate tract of dorsal cervical fasciculus (ITD) has by far the largest number of DNs and the highest proportion of population DNs (Figure 2D, E). Similarly, MTD-II also contained a high proportion of population DNs (Figure 2E). Many of the neurons running in these two tracts have somata located in the GNG neuropil (‘G’) of the brain, as observed previously (Namiki et al., 2018)(Figure 2F).

**Figure 2:**
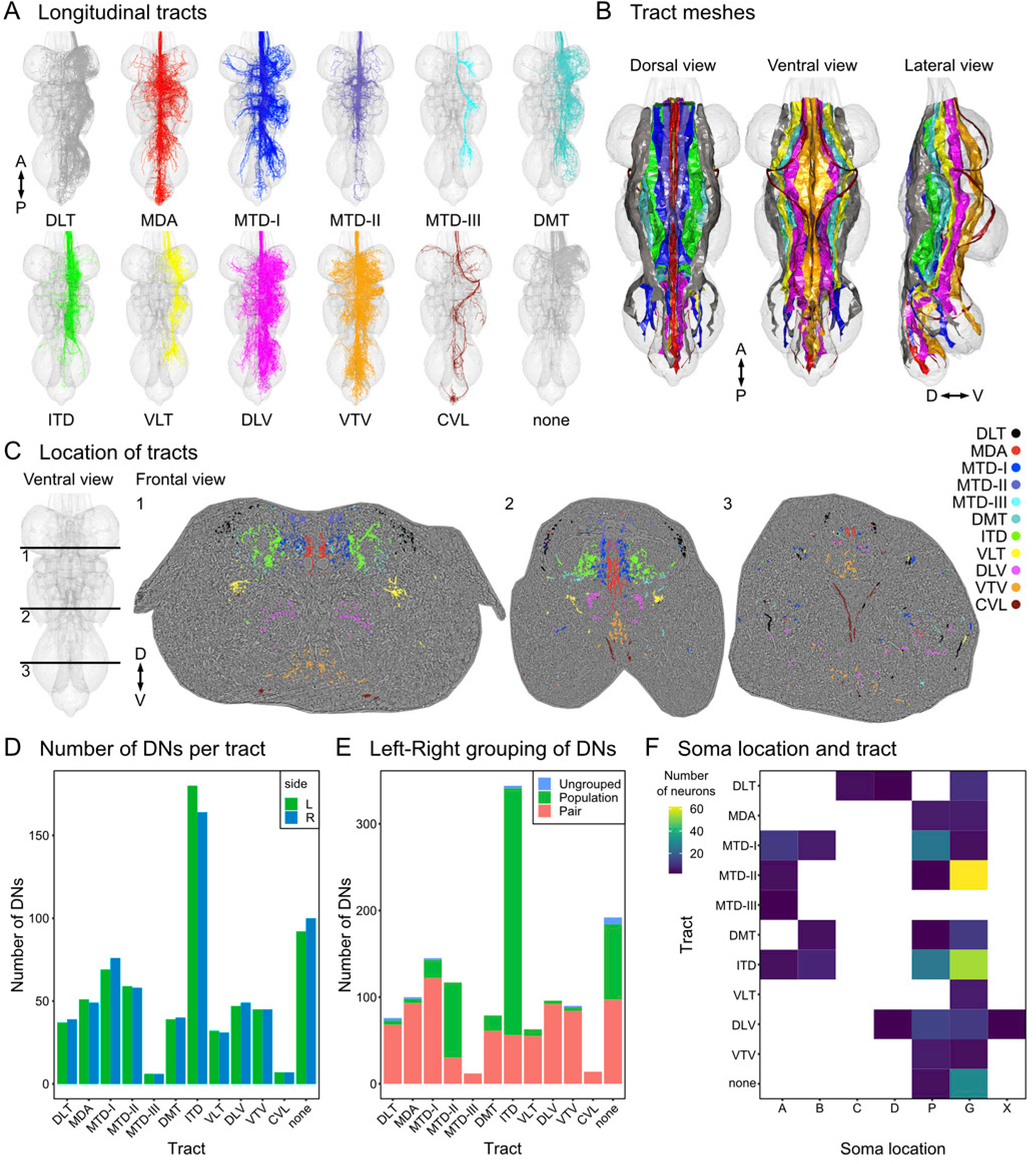
Tract-based analysis of descending neurons. **A.** Tract analysis of all left side DNs (right side DNs are mirror symmetric). DNs run through different tracts in the VNC (Boerner and Duch, 2010; Court et al., 2020; Namiki et al., 2018). Tract names are based on previous publications: DLT, dorsal lateral tract of dorsal cervical fasciculus; MDA, median dorsal abdominal tract; MTD, median tract of dorsal cervical fasciculus; DMT, dorsal medial tract; ITD, intermediate tract of dorsal cervical fasciculus; VLT, ventral lateral tract; DLV, dorsal lateral tract of ventral cervical fasciculus; VTV, ventral median tract of ventral cervical fasciculus. We identified two previously unknown tracts: a third type of MTD (MTD-III) and the CVL, curved ventral lateral tract. DNs with a short longest neurite were not assigned to a tract in agreement with (Namiki et al., 2018). **B.** Tract meshes created from skeletons of DNs assigned to each tract respectively in dorsal, ventral and lateral view. Colors in **B** correspond to **A**. **C.** Cross section (frontal) view of all DN longest neurites color coded by their tract membership (DNs without tract membership were omitted for clarity). Position (1, 2, 3) of section in the VNC indicated by lines on the VNC ventral view. **D.** Number of DNs for each tract separated by cervical connective side. **E.** DNs were annotated with groups based on morphology and connectivity (see Materials & Methods – Identification of descending neurons). Groups can consist of one (pair, 59% of all DNs) or more (population, 39.4% of all DNs) DNs per side. DNs for which we could not identify a group were labeled ‘ungrouped’ (1.6% of all DNs). Shown are the compositions of DN grouping status separated by tract membership. **F.** Correlation of soma location and tract membership for identified DN types (based on light microscopy data, (Namiki et al., 2018)).

We assigned the target neuropils of all DNs by analyzing their synaptic output in the different neuropil areas of the VNC. The VNC is divided along the anterior-posterior axis into three thoracic neuromeres (T1-T3) and one fused abdominal neuromere (ANm), corresponding to the different segments of the fly body (Figure 3A). Along the dorsal-ventral axis, the VNC divides into layers. The dorsal-most layer is the upper tectulum (UTct), which is composed of the neck (NTct), wing (WTct) and haltere (HTct) tectulums that contain MNs for those respective appendages. The ventral-most neuropils are the leg neuropils (LegNp), which are composed of the front (LegNpT1), middle (LegNpT2) and hind leg neuropils (LegNpT3) that contain motor neurons for muscles of the leg in their respective segment. The ventral neuropils also include the leg sensory region, mVAC. In between lie the intermediate tectulum (IntTct), lower tectulum (LTct), and the ovoid (Ov). If a DN had more than 80% of its output in a single neuropil, we designated that neuropil as the DN’s target. If a DN had more than 80% of its output in two neuropils, with each contributing at least 5%, we assigned both neuropils as a target, annotated in the neuPrint database as the two neuropil names separated by a full stop (e.g. IntTct.ANm). Neurons that had over 80% combined output to the three upper tectulum neuropils (neck, wing and haltere tectulum) or to the leg neuropils (LegNpT1, LegNpT2, LegNpT3), we assigned the layer label UTct or LegNp as a target, respectively (Figure 3B). Previous work suggested a correlation between a DN’s tract and its target neuropil (Namiki et al., 2018). We see that this still holds loosely for the entire DN population analyzed at synaptic resolution (Figure 3C, Figure 3–Supplement 1C).

**Figure 3:**
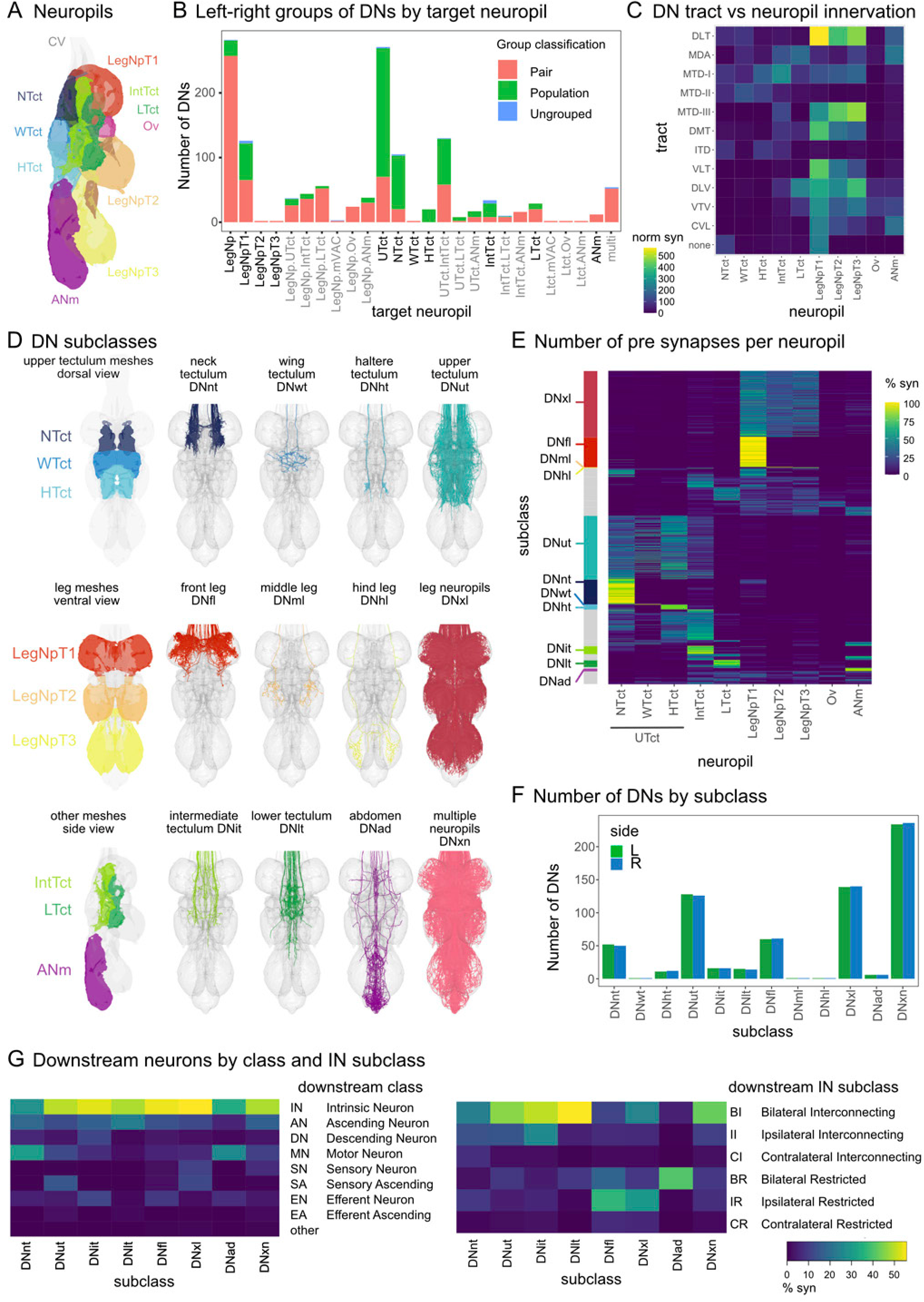
DN subclasses. Descending neurons were analyzed based on neuropil innervation. **A.** VNC neuropils. **B.** X-axis indicates target neuropil designations (the ROI neuropils that together receive more than 80% of the DNs output, see Materials & Methods) and bar color indicates DN grouping status (as in Figure 2). Target neuropil groups that were used to subsequently define DN subclasses are in black, all grey target neuropil groups received the subclass DNxn (see Figure 3–Supplement 1). **C.** The relationship between neuropil location of DN presynaptic sites and tract membership. Total synapse number normalized by the number of DNs in the tract is shown. **D.** DNs are assigned a subclass based on their target neuropil: DNnt, Neck Tectulum; DNwt, Wing Tectulum; DNht, Haltere Tectulum; DNut, upper tectulum, if they target a combination of neck, wing, and haltere neuropils; DNfl, DNml, DNhl, innervating single leg neuropils front leg, middle leg, and hind leg respectively; DNxl, if they target a combination of several leg neuropils; DNit, Intermediate Tectulum; DNlt, Lower Tectulum; DNad, Abdominal Neuromeres. DNs that innervate more widely are referred to as DNxn, for multiple neuropils. **E.** Heatmap representation of DN neuropil innervation, by percent of each DN’s synaptic output. Every row is a single DN output. Bar along the y-axis shows the subclass assigned to different DNs, grey are those in xn category (see Figure 3–Supplement 1). **F.** The number of DNs in a given subclass assigned to either left or right side. There is a single pair of DNs for the subclasses DNwt, DNml and DNhl (see images in **C**). **G.** DN postsynaptic partner composition by neuron class and interneuron subclass.

**Figure 3—Supplement 1:**
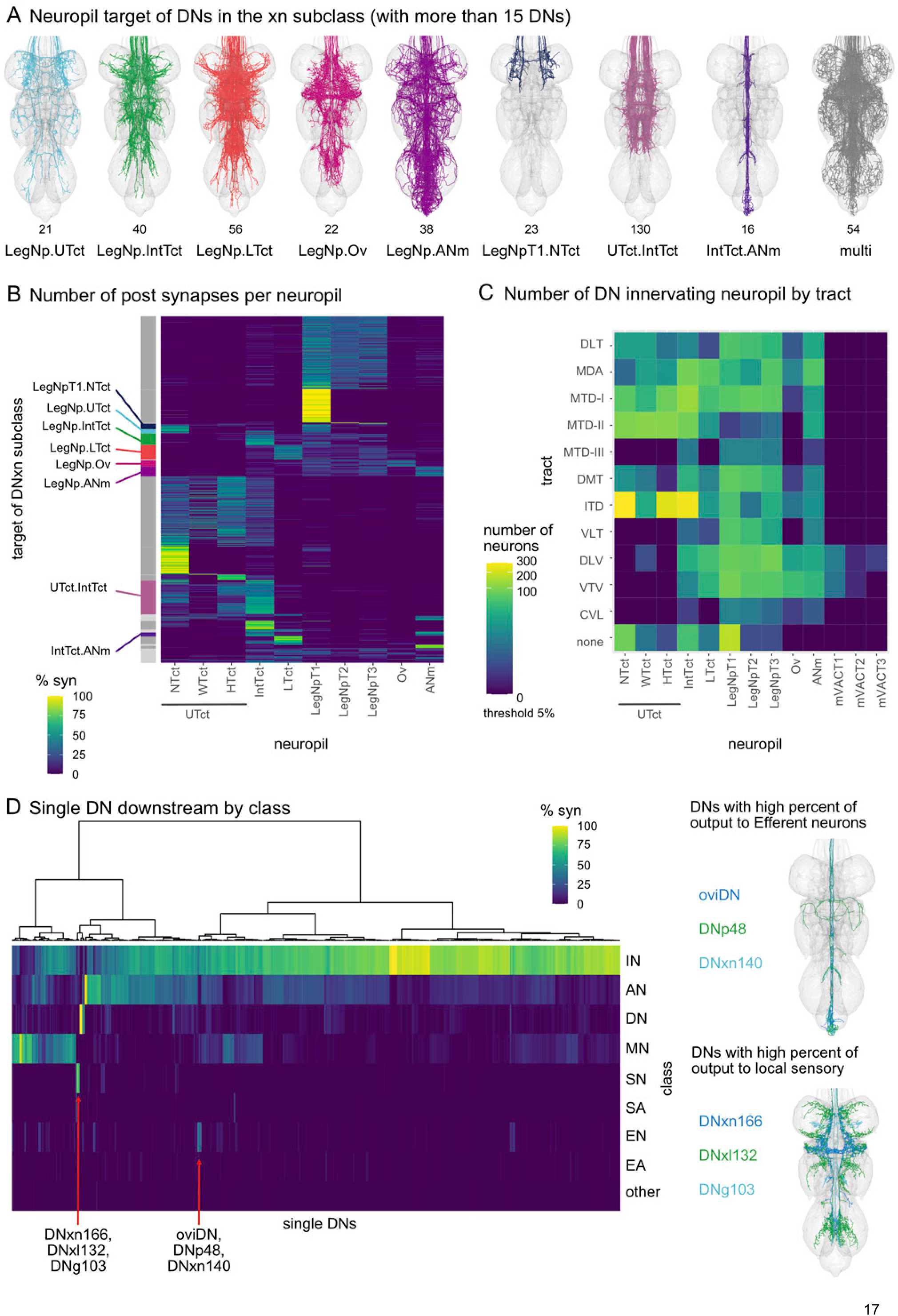
DN subclasses. **A.** DNs of subclass xn, which consisted of large groups (>15 single DNs) of DNs with common target distribution in the VNC. Number of single DNs within each xn subclass and names of combined target neuropils shown below each group. ‘Multi’ DNxns had 80% of the DN output in more than two neuropils. **B.** Heatmap representation of DN neuropil innervation by number of neurons. Every row is a single DN. Bar along the y-axis shows the neuropil targets (with >15 DNs) assigned to subclass DNxn, coloured only the ones shown in **A**. **C.** Number of DNs by tract innervating neuropils. Innervation in a neuropil is counted if the DN has more than 5% of its output to that neuropil. **D.** DN output to a neuron class in percent for every single DN, plotted on the x-axis. Red arrows point to two small groups of DNs, one with a high proportion of output to efferent neurons and the other with a high proportion of output to local sensory neurons.

**Figure 3—Supplement 2:**
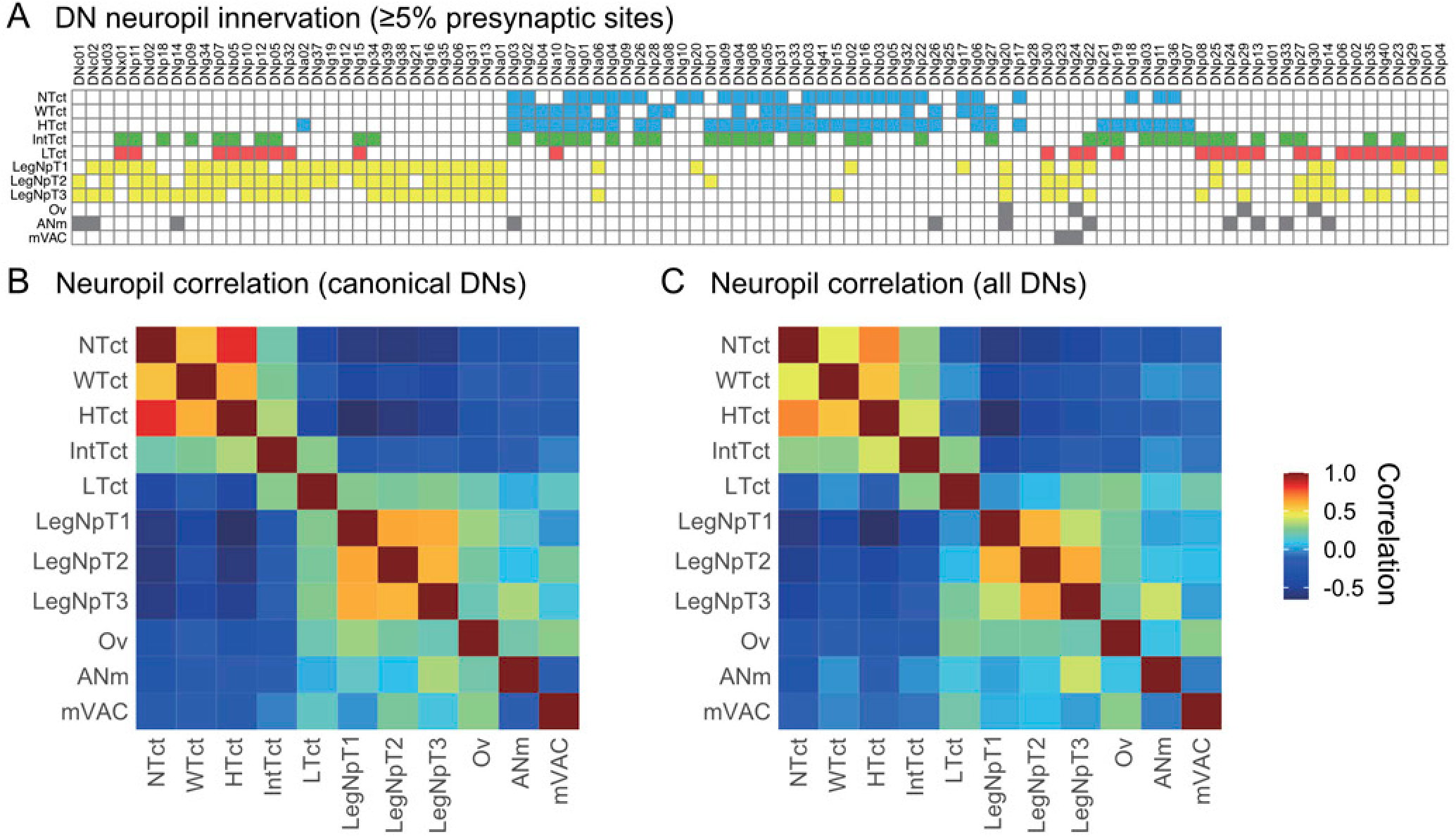
Neuropil innervation for descending neuron types described in Namiki et al., 2018. **A.** Neuropil innervation of DN types. Filled heatmap cells indicate that at least 5% of a DN group’s presynaptic sites are located in a given ROI. X-axis order determined by (Namiki et al., 2018), Figure 7. **B-C.** Autocorrelation matrix of DN innervation pattern in the VNC. The Pearson’s correlation coefficient was calculated for each pair of ROIs between DN innervation of identified canonical Namiki types (**B**) and all types in the MANC EM volume (**C**).

DN neurons were previously named following a convention that divided them into subclasses based on soma location in the brain (Namiki et al., 2018). We sought to systematically name the 376 newly identified DN types in MANC, but did not have soma location available to us to follow the published nomenclature. We therefore adopted a new systematic naming convention based on our annotation of a DN’s VNC neuropil target (Figure 3D, E). This system has the advantage of grouping DNs into subclasses of broadly similar function. Future work should unify these two naming conventions. All DNs annotated with a single target neuropil were given a two-letter subclass abbreviation corresponding to their target neuropil as follows: DNnt (neck tectulum), DNwt (wing tectulum), DNht (haltere tectulum), DNfl (“front leg,” T1 leg neuropil), DNml (“middle leg,” T2 leg neuropil), DNhl (“hind leg,” T3 leg neuropil), DNit (intermediate tectulum), DNlt (lower tectulum), DNad (abdominal neuropil). In the ‘systematic type’ name, individual DN types were then further distinguished within their subclass by a three-digit number (e.g. ‘DNnt001’, see Figure 1E). DNs that had more than one target neuropil were named according to the group of neuropils they targeted as follows: DNut (upper tectulum, including neck, wing, and haltere tectulums), DNxl (all leg neuropils), DNxn (broad targeting of many neuropils) (Figure 3D, E, F). For a further breakdown of DNxn types, see Figure 3–Supplement 1A, B. Previous neuropil correlations, based on light level data of 30% of DNs (Namiki et al., 2018), are representative of the correlations seen for all DNs at synaptic resolution (Figure 3–Supplement 2A, B, C).

These DN subclasses differ not only in the neuropils they target but also in the classes and subclasses of postsynaptic neurons to which they signal (Figure 3G, left panel). While all DNs primarily output to intrinsic neurons (IN), the largest VNC class, a high proportion of DNnt and DNad subclasses also output to MNs in the neck and abdomen, respectively. Within the INs, there is a distinction between the DNs of the dorsal neuropils, which have a higher proportion of their output to bilateral interconnecting INs and DNs innervating the leg neuropils, which mostly target INs local to a single leg neuropil (Figure 3G, right panel). There are small groups of DNs that strongly target some of the other classes of neurons in the VNC, such as efferent neurons or local sensory neurons (see Figure 3–Supplement 1D).

Knowing whether DNs are excitatory or inhibitory is critical for forming functional hypotheses about descending to motor pathways in the connectome. (Takemura et al., 2024) predicted ACh, GABA, or Glu neurotransmitter type for all MANC neurons using a convolutional neural network. We provide examples of the EM data for neurons with both high and low neurotransmitter prediction score (Figure 4A-D). For example, DNp20 (aka DNOVS1) is known to be gap junction coupled to downstream targets (Suver et al., 2016), potentially explaining its low neurotransmitter prediction score. For DNs, predicted use of the three primary neurotransmitters appear associated with their ratio of output to input in the VNC–DNs with high neurotransmitter prediction confidence for these neurotransmitters also have high output to input ratios, while those with low confidence have a lower ratio, perhaps reflecting use of extrasynaptic transmission methods (Figure 4E-G). However, the fidelity of DN neurotransmitter predictions is unclear, as DNs were not included within the initial training dataset. Also, the distribution of predicted DN neurotransmitter usage differed markedly from prior experimental results: EM prediction—68.4% cholinergic, 16.2% GABAergic, 7.5% glutamatergic and 7.9% other or below threshold (with a prediction probability cutoff score of 0.7), co-labeling with neurotransmitter GAL4 lines or antibodies against neurotransmitters or neurotransmitter synthesis proteins—38% cholinergic, 37% GABAergic, and 6% glutamatergic DNs (Hsu and Bhandawat, 2016). These differences could be due to incorrect EM neurotransmitter prediction, but may also be due to incomplete or inaccurate labeling in commonly-used neurotransmitter marker lines, or due to underlying biological details (e.g. marker expression does not always equate to neurotransmitter usage). To evaluate the quality of the neurotransmitter predictions for the DNs, we selected a panel of 53 DN types for which we had a confident match between a MANC DN and a DN targeted in a split-GAL4 line. In this panel of DNs, 11 DN types from the EM volume are matched against newly generated split-Gal4 lines; a more comprehensive description and naming of new DNs identified at the light level, and their EM matches, is described separately (Stürner et al., 2024). We used the set of split-Gal4 lines to perform fluorescent in situ hybridization (FISH, (Meissner et al., 2019)) or Expansion-Assisted Iterative FISH (EASI-FISH) (Eddison, 2022; Wang et al., 2021) against the neurotransmitter markers *ChAT*, *Gad1* and *VGlut* that mark ACh, GABA and Glu, respectively. A subset of 15 DNs that had low neurotransmitter prediction scores or were of interest were also probed for markers for the minor neurotransmitters octopamine, tyramine, serotonin, and dopamine. We found 23 out of 28 cholinergic (additional 4 uncertain), 3 out of 9 GABAergic (additional 1 uncertain), and 9 out of 10 glutamatergic (additional 1 uncertain) predictions to be correct, while 2 out of 6 DNs with low prediction scores were found to use a primary neurotransmitter (additional 2 uncertain, Figure 4H, Figure 4–Supplement 1, Supplementary file 2). Interestingly, we observed possible co-transmission of Glu and ACh for DNg27, DNg40, DNp08 and DNp49, Glu and tyramine for DNd02, and Glu (weakly expressing) and tyramine or octopamine in DNg34, which was a likely match for the known octopaminergic DN aDT8/OA-VPM1 (Busch et al., 2009; Yu et al., 2010). Prior work has shown that ChAT mRNA can be weakly transcribed without producing detectable ChAT protein (Lacin et al., 2019); thus, the possible co-transmission described here will require further verification. Overall, neurotransmitter FISH results suggest that for DNs, ACh and Glu predictions are a good indication of actual neurotransmitter usage, whereas GABA predictions are still unverified. Thus, neurotransmitter predictions allow users a reasonable first guess at DN neurotransmitter identity but must be followed up with independent validation.

**Figure 4:**
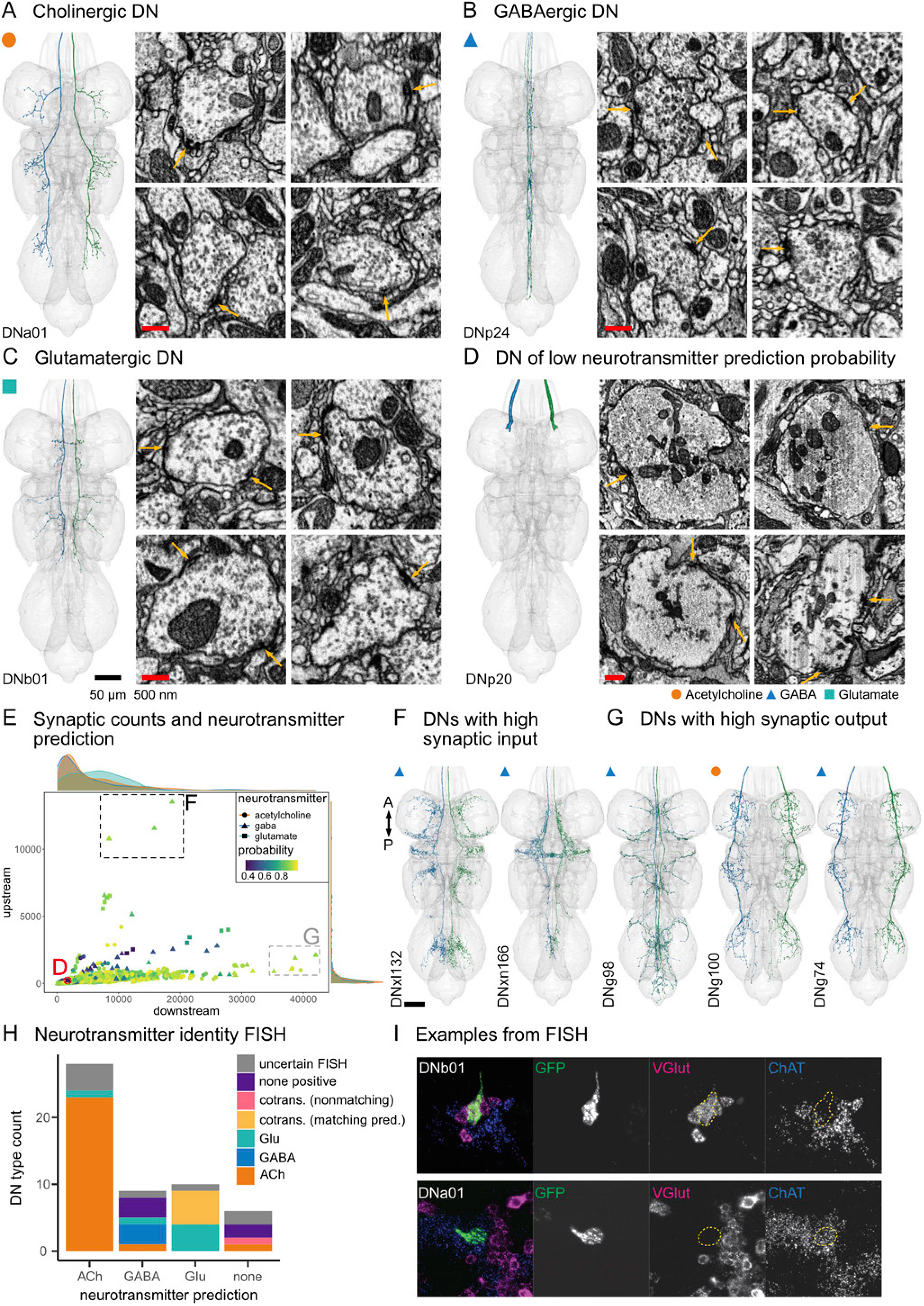
Neurotransmitter prediction for DNs. **A-D.** Morphology of example DNs with high probability neurotransmitter predictions for acetylcholine (**A**), GABA (**B**), glutamate (**C**) and very low probability neurotransmitter prediction (**D**). Four presynaptic sites are shown for each DN with yellow arrows pointing to the presynaptic density (T-bar) and synaptic cleft. **D.** DNp20 has few vesicles and may be electrically coupled (indicated by larger size diameter of the axon). **E.** Counts of pre- and postsynaptic sites. Every shape represents a single DN and its neurotransmitter prediction (acetylcholine, circle; GABA, triangle; glutamate, square). The prediction probability is color coded. Red circle indicates DNp20 shown in **D**, black rectangle indicates the two DN types shown in **F** and the gray rectangle the 3 DN types shown in **G**. **F.** Morphology of two DN pairs with high synaptic input in the VNC. The predicted neurotransmitter for both is GABA. **G.** Morphology of three DN pairs with high synaptic output in the VNC of which the neurotransmitter of two was predicted GABA and one acetylcholine. The type of each DN is indicated in the bottom left. Black scale bars are 50 µm, red scale bars are 500 nm. Scale bars in EM synapse panels are the same if not indicated otherwise. **H.** Neurotransmitter identity for DNs from FISH technique, see Figure 4–Supplement 1 for images and DN identities. **I.** Example images from the FISH analysis, DNb01 (SS02383) and DNa01 (SS00731). Dotted yellow line indicates the DN soma location (see Material and Methods for details).

**Figure 4—Supplement 1:**
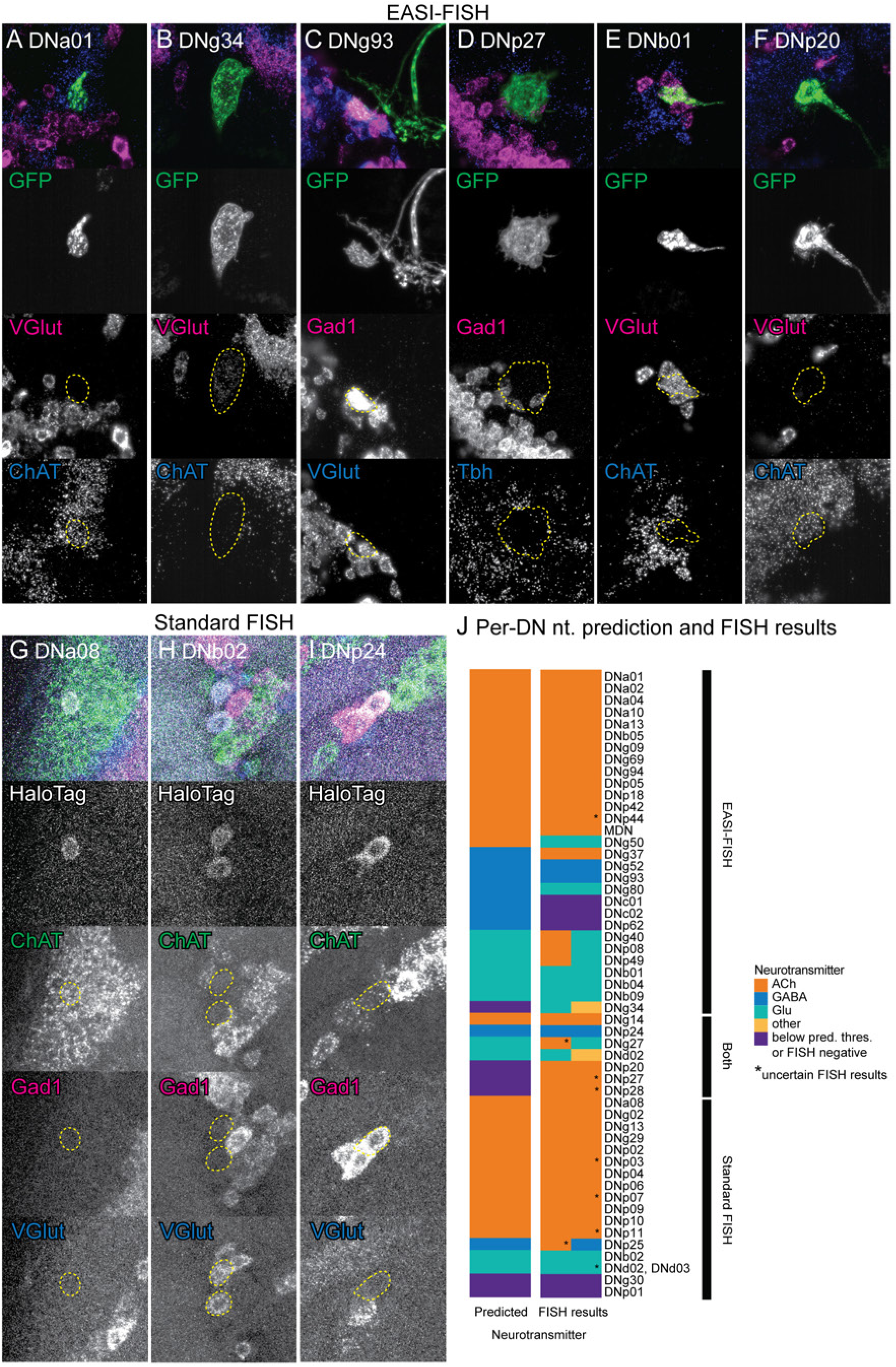
DN FISH for neurotransmitter markers. To survey neurotransmitter usage in DNs, adult Drosophila brains expressing marker genes in individual DN types were probed using either EASI-FISH (Eddison, 2022; Wang et al., 2021) or a standardized FISH technique (Meissner et al., 2019). Example results for nine DNs are shown here. **A-F**. For EASI-FISH, brains from DN split-Gal4 lines expressing UAS-CsChrimson-mVenus were probed for the indicated combinations of two neurotransmitter markers, as well as immunostained for GFP. Labeled DN types and their split-Gal4 lines are as follows: **A.** DNa01 (SS00731), **B.** DNg34 (SS58292), **C.** DNg93 (SS93763, synonym: oval), **D.** DNp27 (SS00923), **E.** DNb01 (SS02383), **F.** DNp20 (SS01078). **G-I.** For standard FISH, brains from DN split-Gal4 lines expressing UAS-7xHaloTag::CAAX were probed for ChAT, Gad1 and VGlut, and stained with fluorescent conjugated HaloTag ligand. Labeled DN types and their split-Gal4 lines are as follows: **G.** DNa08 (SS02393), **H.** DNb02 (SS01060), **I.** DNp24 (SS00732). Soma of DNs of interest are outlined in all FISH channels. Maximum intensity projections in all images are from a Z-range in each stack that closely flanks the soma of interest. **J.** Summary of EM neurotransmitter prediction and FISH-determined neurotransmitter usage for all DN types tested (y-axis). Multiple colors per row for FISH results indicates positive marker expression for multiple neurotransmitters. (*) indicates uncertain FISH results. See Supplementary File 2 for detailed results.

### 2.3 Identification of motor neurons

To identify the motor neurons (MNs), all the nerves exiting the VNC were first identified (Figure 5A), and all neurites passing through these nerves were seeded, proofread and annotated for entry or exit nerve (Marin et al., 2024). MNs were distinguished from the two other neuron classes that have profiles in the nerves by having a soma in the VNC (which sensory afferents lack) and no profile in the neck connective (which efferent ascendings have)(Marin et al., 2024). Additionally, neurons which exit through multiple nerves or were putatively recognized as neuromodulatory were assigned the efferent neuron class; the criteria for this class is broader than the MN class, and may contain currently unrecognized MNs. As with DNs, we carried out additional proofreading and categorization of all MNs of the VNC to obtain a higher degree of upstream completeness (Figure 5B). We note that, although the quality of the EM data underlying the MANC connectome was generally very high, some MNs, mainly of the leg neuropils but also several of the neck and halteres, showed signs of degeneration (dark cytoplasm and irregularly-shaped neurites) due to axonal damage during dissection (Figure 5–Supplement 1). As a result, both morphology and connectivity for affected MNs was sparser than for non-affected MNs, and more processing and proofreading of the EM data for affected MNs was required (see Materials & Methods). These reconstruction issues affect an estimated 49% of leg MNs and 5% of other MNs (Figure 5–Supplement 1A). Overall, we observed that most MNs have their inputs largely confined to a single neuropil, although some MNs arborize more widely (Figure 5C), corroborating light-level neuropil delineations and their likely functional specialization in motor control (Court et al., 2020). Together, MNs densely cover all neuropils except the predominantly sensory neuropils (Ov, VACs, and mVACs) and the intermediate neuropils.

**Figure 5:**
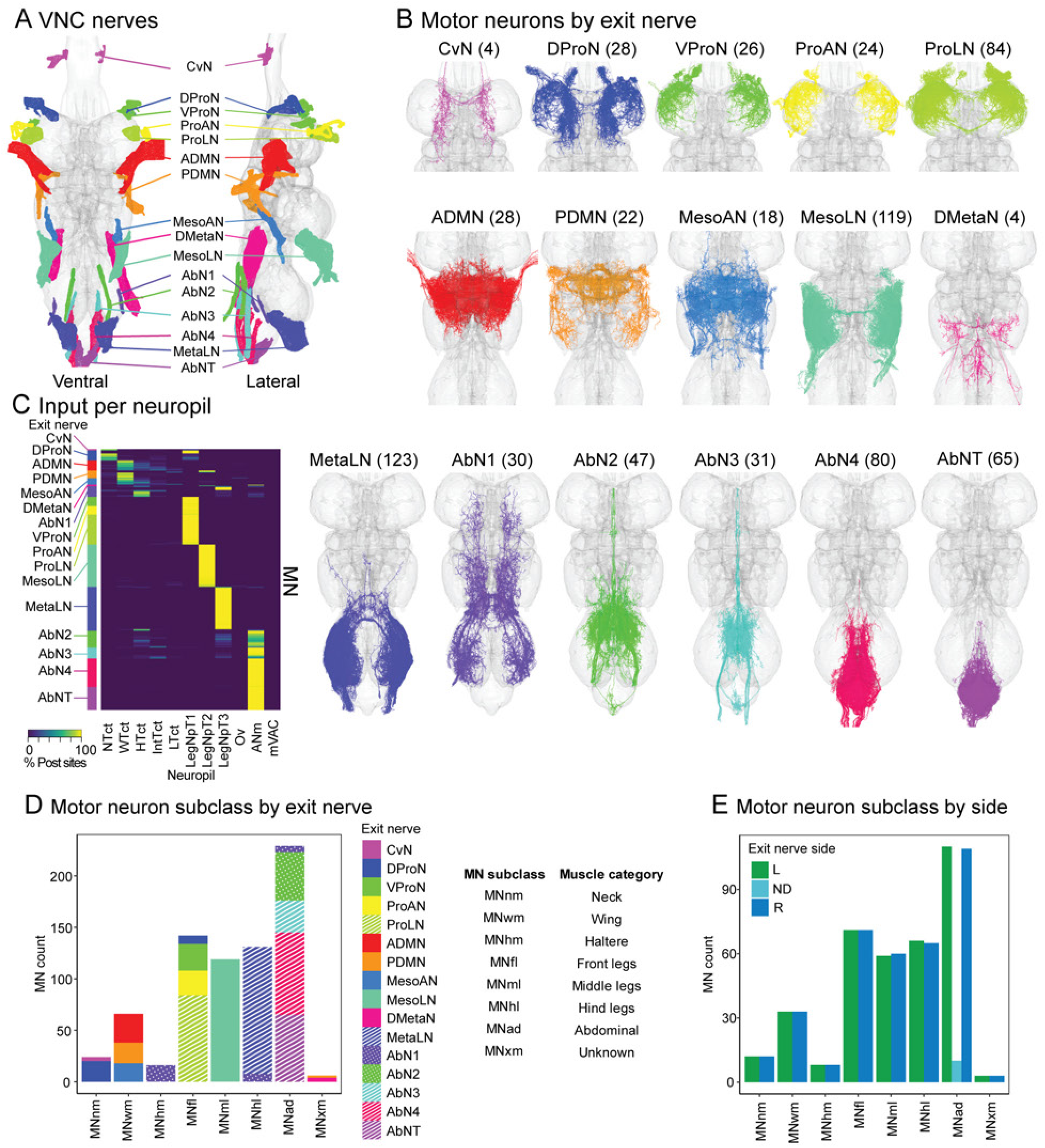
Motor neurons of the Drosophila male VNC. **A.** VNC nerves from which motor neurons (MNs) exit. Nerve abbreviations are as follows: cervical nerve (CvN), dorsal prothoracic nerve (DProN), ventral prothoracic nerve (VProN), prothoracic accessory nerve (ProAN), prothoracic leg nerve (ProLN), anterior dorsal mesothoracic nerve (ADMN), posterior dorsal mesothoracic nerve (PDMN), mesothoracic accessory nerve (MesoAN), mesothoracic leg nerve (MesoLN), dorsal metathoracic nerve (DMetaN), metathoracic leg nerve (MetaLN), first abdominal nerve (AbN1), second abdominal nerve (AbN2), third abdominal nerve (AbN3), fourth abdominal nerve (AbN4), and abdominal nerve trunk (AbNT). **B.** All MNs found in MANC, classified by exit nerve. Number in parentheses indicates cell count. **C.** MN synaptic input per neuropil. **D.** MN subclass classification by broad muscle category (e.g. leg, wing). **E.** MN count per subclass by exit nerve side: left (L), right (R) or not determined (ND). Exit nerve side was not determined only for a subset of MNs exiting near the midline through the AbNT.

**Figure 5—Supplement 1:**
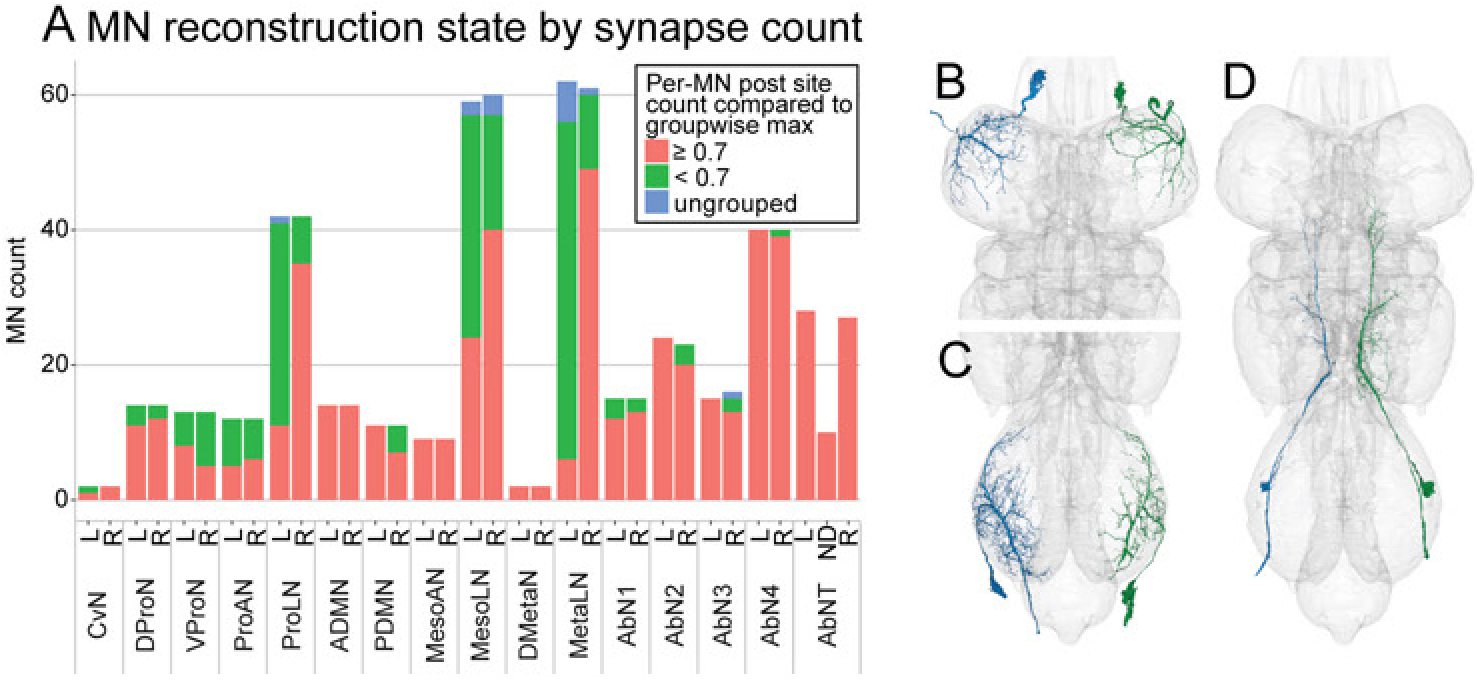
Reconstruction state of motor neurons in MANC. **A.** Motor neurons (MNs) per nerve and side, classified by whether they have less than 70% (0.7) of the predicted postsynaptic site counts compared to their groupwise maximum, i.e. the postsynaptic count of the most well-traced member of each MN’s morphological group in MANC. This classification flags MNs with likely reconstruction issues. Note that MN groups where all members of the group have reconstruction issues may not be flagged. **B-D.** Example MN groups with one well- and one less-well-reconstructed member: **B.** Sternal rotator anterior MN (group 13300), **C.** Ti extensor MN (group 10347), **D.** MNhm43 (group 17216).

MNs were organized by morphology into ‘groups’ that consisted of left-right homologous pairs or populations using the same mirrored NBLAST analysis applied to the DNs (this paper and (Marin et al., 2024), together with exit nerve annotations and connectivity comparisons. In total, we found 733 MNs in MANC, with 362 exiting left side nerves, 361 exiting right side nerves, and 10 that exited through the abdominal trunk nerve (AbNT) near the midline (Figure 5B, E). These numbers were similar to those from the Female Adult Nerve Cord (FANC) dataset, where MNs traced (which excluded AbN2-4 and AbNT MNs) numbered 507 (Phelps et al., 2021)—we found 510 for the MNs in the same nerves. We organized these into 319 groups and 14 singletons (that could not be paired). Similar to the DNs, we assigned both a ‘type’ and ‘systematic type’ to each MN group in the NeuPrint database. For the leg MNs, we additionally organized the groups into ‘serial groups’ of homologous neurons in the T1, T2 and T3 segments. Roughly 150 abdominal MNs were also organized into serial groups (Marin et al., 2024). All MNs were assigned a ‘systematic type’ by serial group, group or singleton cell (in descending order of priority), consisting of the ‘MN’ prefix, followed by a two-letter code denoting the broad muscle category they target (neck, nm; wing, wm; haltere, hm; front leg, fl; middle leg, ml; hind leg, hl; abdominal, ad; unknown, xm) and a two-digit number (Figure 5D, E, also see Figure 1E). For ‘type’ names, MNs were named by prior known names in the literature if identified, and by systematic type otherwise. Lastly, we annotated ‘target’ and ‘subclass’ fields for each MN, where target is the exact muscle target name if known, while subclass is their two-letter abbreviation for broad muscle category as defined in their systematic type.

Each *Drosophila* MN is identified by the single muscle it innervates. However, the muscles were outside our EM dataset, which included only the VNC, so we could not directly observe this correspondence in our data. Instead, to match MANC MNs to their target muscles, we relied on matching the morphology of MN dendrites, which reside in the VNC volume, to images of known MNs in the literature. This was accomplished slightly differently for the different motor systems:

*Wing and haltere MNs*. To determine the target muscle for wing and haltere MNs, we matched the MN dendrite morphology in MANC to light microscopy imagery of MNs with known muscle targets (Ehrhardt et al., 2023), or for three MNs identified their muscle targets in new or existing split-Gal4 lines (Sterne et al., 2021; Wu et al., 2016) (Figure 6–Supplement 1). These efforts allowed us to match all but one of the 26 MNs innervating wing muscles (Figure 6A, B). A single putative wing MN (MNwm36) with similar morphology to the ps1 MN remains unassigned to any muscle; (Azevedo et al., 2024) hypothesizes that it innervates a pleurosternal muscle, likely contributing multiple innervation together with currently known pleurosternal MNs. To determine the MNs of the eight haltere muscles, we used light microscopy data from two papers describing haltere muscles and MN drivers in *Drosophila* (Dickerson et al., 2019; Ehrhardt et al., 2023). We identified seven pairs of haltere MNs, though three of these matches are putative (Figure 6C, D). Morphological identification of these MNs are further supported by their connectivity. Cosine similarity of the synaptic input to these MNs shows that input to wing steering MNs is commonly shared between muscles of the same sclerite (and less commonly between select muscles of different sclerites), while synaptic input is highly shared within the wing power and indirect control MNs, respectively (Figure 6E).

**Figure 6:**
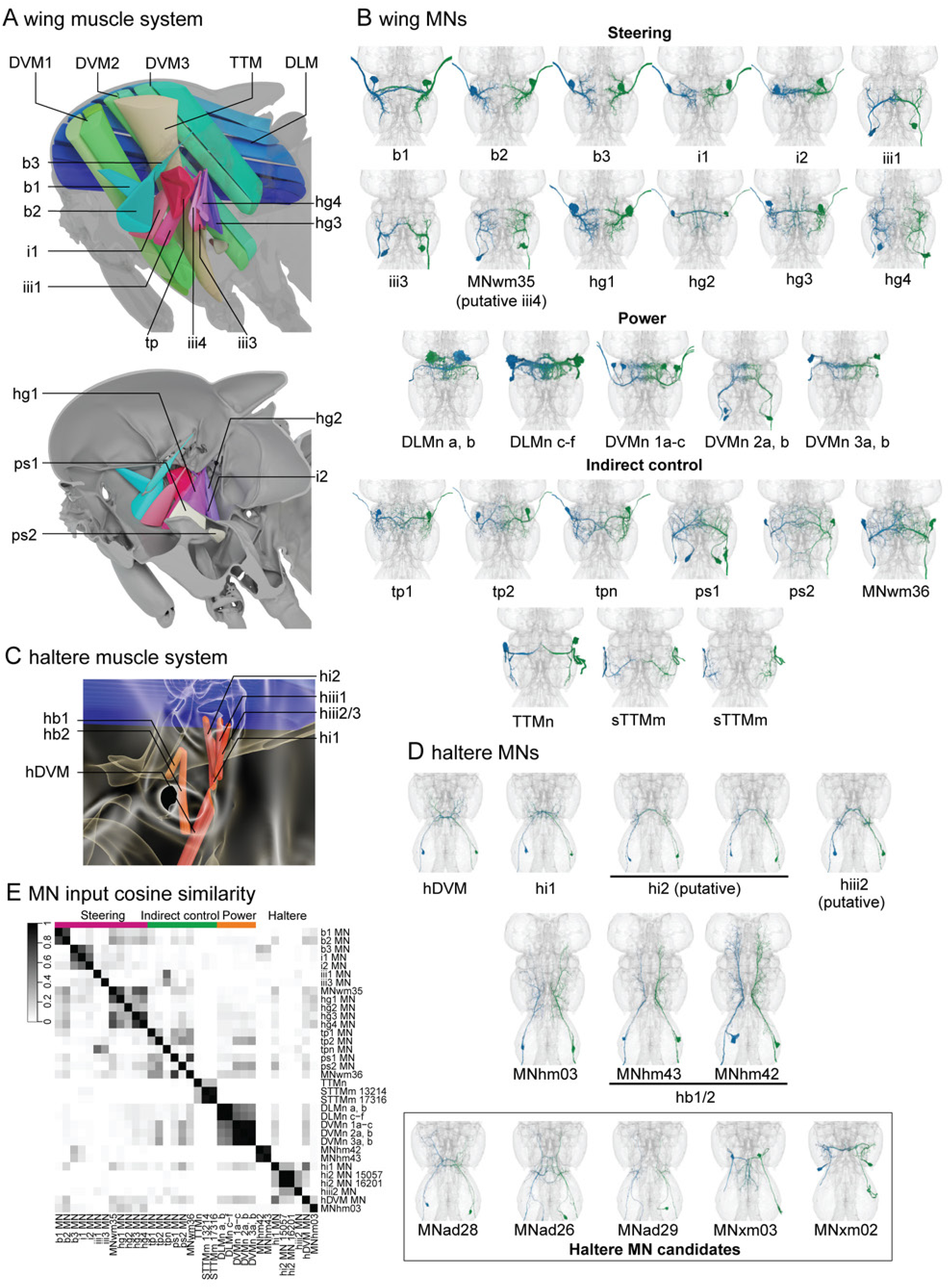
Upper tectular motor neuron identification in MANC. **A.** Thoracic organization of wing muscles shown from a lateral (top panel) or medial (bottom panel) view. Muscle categories and abbreviations are as follows: power muscles (dorsal longitudinal muscles, DLM; dorsal ventral muscles, DVM1-3), steering muscles (basalar, b1-3; first axillary, i1, i2; third axillary, iii1, iii3, iii4; fourth axillary, hg1-4), and indirect control muscles (tergopleural, tp; pleurosternal, ps1, ps2; tergotrochanteral, TTM). The hg2 muscle in the lateral view, and the power muscles and TTM in the medial view are omitted for clarity. **B.** Identification of wing muscle MNs from light-level data (further see Supplementary file 3). The iii4 MN match (MNwm35) is putative, and the muscle target of MNwm36 is unknown. **C.** Organization of haltere muscles. Muscle categories and abbreviations are as follows: power muscle (haltere dorsal ventral muscle, hDVM), and steering muscles (haltere basalar, hb1, hb2; first axillary, hi1, hi2; third axillary, hiii1-3). **D.** Identification of haltere muscle MNs from light-level data. Two candidates for the hi2 MN (groups 15057 and 16201) with similar morphology and connectivity are present in MANC. MNhm43 and MNhm42 innervate the hb1 and hb2 muscles, however the exact target of each MN is not yet known. While additional putative haltere MNs are not easily determined in EM as both haltere MNs and abdominal MNs exit the AbN1 nerve, three potential haltere MNs are subsetted here by their arborization in the haltere neuropil and exit through AbN1, and two potential haltere MNs by exit through the haltere nerve, DMetaN). E. Cosine similarity for wing and haltere MN types by their upstream connectivity to all neurons combined by type or serial set (prioritized over type if assigned).

**Figure 6—Supplement 1:**
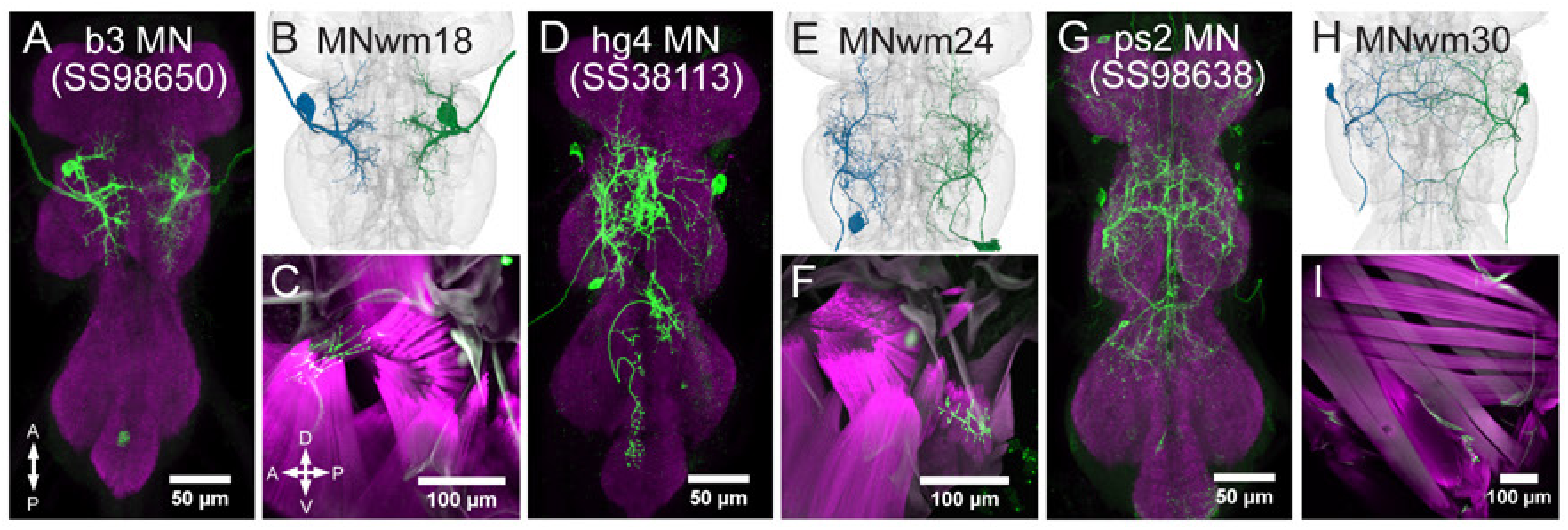
Wing MN drivers and their muscle targets determined in this paper. **A.** VNC expression pattern (green) of the split-Gal4 line SS98650 (VT058382-AD; VT042734-DBD) compared to its MN match in EM (**B**). **C.** b3 muscle innervation (green) in a longitudinal section of the thorax in SS98650. **D.** VNC expression pattern of SS38113 (Sterne et al., 2021) compared to its MN match (systematic type) in EM (**E**). Expression in the wing MN is stochastic in this line, and its MN morphology and muscle innervation was corroborated with another stochastic line, OL0070B (not shown; (Wu et al., 2016)). **F.** hg4 muscle innervation a longitudinal section of the thorax in SS38113. **G.** VNC expression pattern of SS98638 (VT026026-AD; VT064565-DBD) compared to its MN match in EM (**H**). **I.** ps2 muscle innervation in a longitudinal section of the thorax in SS98650. All VNCs are counterstained for mAb nc82 (magenta), while all muscles are counterstained with phalloidin (magenta).

*Leg MNs*. Less light microscopy data was available for leg MNs, so we used a different strategy to identify leg MNs and their muscle targets in three main steps. We first examined light microscopy stacks for previously reported neurons (Enriquez et al., 2018), allowing us to assign several MNs of the T1 leg neuropil with relatively low confidences. We next transformed the T1 leg MNs from the Female Adult Nerve Cord (FANC) EM volume (Azevedo et al., 2024) to MANC space and performed an NBLAST comparison between MNs from the two VNC datasets (Costa et al., 2016). With expert visual evaluation, this allowed us to match up all FANC-identified T1 leg MNs to the MANC dataset (Figure 1D, Figure 7). To extend identification of leg MNs even further, we noted that the fly’s three pairs of legs, and their muscles, exhibit a high degree of serial homology across segments (Miller, 1950; Soler et al., 2004) that is likely reflected in the morphology and connectivity of the leg MNs. As serially-repeating homologs of other neuron classes were identified across segments in MANC (Marin et al., 2024), we leveraged this data to identify serial homologs of the T1 leg MNs in the T2 and T3 segments by comparison of MN connectivity with these serial homologs, as well as connectivity with neurons that span across several thoracic segments (mainly DNs, ANs and INs). Overall, in the MANC dataset, we find 392 leg MNs (142 in T1, 119 in T2, 131 in T3). We identified 142 of the 142 T1 MNs and, through serial matching, putatively identified 198 of the 252 leg MNs in the T2 and T3 segments (Figure 7A, B). Thus, we were able to map leg muscle targets to most of the MNs across the three leg neuropils (Figure 7A, C).

**Figure 7:**
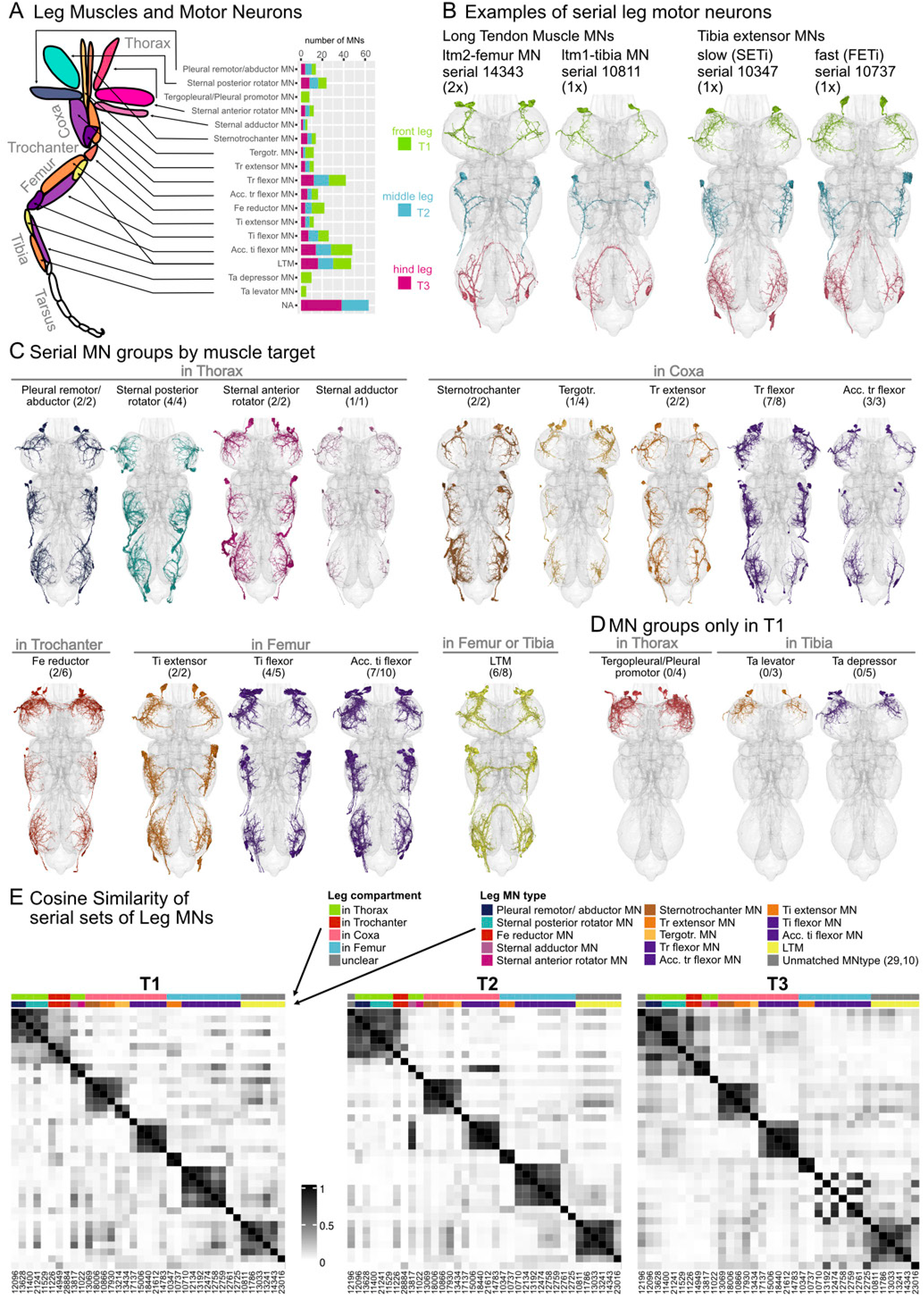
Serial leg motor neurons. **A.** Diagram of a prothoracic leg showing the position of muscle groups innervated by motor neurons (MNs), some of which attach across the thorax, and others of which are intrinsic to the coxa, trochanter, femur, or tibia leg segments. Adapted from (Brierley et al., 2012). On the right the number of MNs that could be assigned a muscle target split into soma neuromere (T1, T2, T3). **B.** Example of two serial MN sets that innervate the long tendon muscles (LTM) and two serial MN sets that innervate the tibia extensor muscles, one for slow and one for fast tibia extension. **C.** All MNs that are in a serial set listed by their muscle target. In brackets the number of serial sets found / number of neurons in T1 pairs. Colors by the muscle they innervate as shown in the cartoon in **A**. **D.** For three of the muscle targets we were not able to find corresponding neurons in T2 and T3 leg neuropils. Thus only the T1 leg neuropils have muscle target assignments. **E.** Cosine similarity of all leg MNs by their connections to serial local neurons in T1, T2 and T3 leg neuropils. MNs are organized by muscle targets and leg compartment, shown in corresponding coloured bars. All MN targets in T1 were assigned by careful matching to the FANC dataset (Azevedo et al., 2024), while all MN targets in T2 and T3 were assigned by serial sets (see Materials & Methods).

**Figure 7—Supplement 1:**
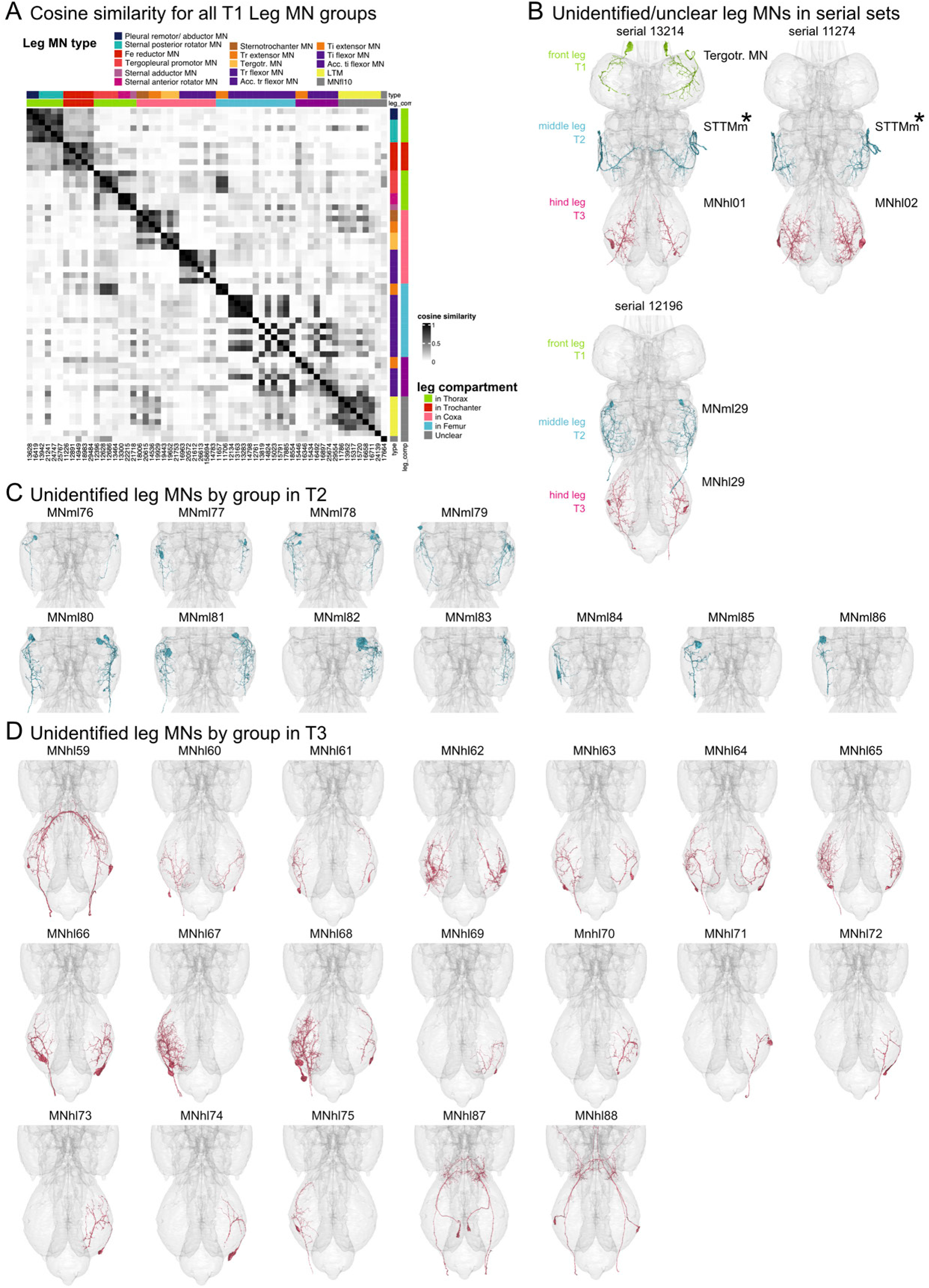
Unidentified leg MNs in MANC. **A.** Cosine similarity of all leg MN groups by their connections to serial local neurons in T1, T2 and T3 leg neuropils. MNs are organized by muscle targets and leg compartment, shown in corresponding coloured bars. All MN targets in T1 were assigned by careful matching to the FANC dataset ((Azevedo et al., 2024), see Materials & Methods). **B.** Four serial MN sets that could not be assigned to the same muscle target across segments. The first two examples each include a T2 pair that has been matched to the literature type satellite tergotrochanteral motor neuron, STTMm (Bacon and Strausfeld 1986)(marked with *). By connectivity and morphology they have a serial pair in T3 (and in T1 a MN assigned to the Tergotr. MN). **C.** All MNs by group in T2 that we were unable to assign a serial set. **D.** All MNs by group in T3 that we were unable to assign a serial set.

We could not identify serially-repeating homologs (in T2 and T3) for the Tarsus levators and depressor MNs as well as the Tergopleural/Pleural promotor MNs (Figure 7D). For the tarsus targeting MNs, this was mainly due to their reconstruction state in T1 which made them hard to distinguish. For the Tergopleural/Pleural promotor MNs that innervate a muscle in the thorax, we could not find MNs in T2 and T3 that were similar in morphology and connectivity, suggesting that these MNs are either not present or have a different function and morphology (for unmatched leg MNs, see Figure 7–Supplement 1C, D). To compare synaptic input of the leg MNs across the serially repeated leg neuropils, we calculated the cosine similarity of the upstream connections of serial MN sets from other serially repeated neurons that were restricted to the leg neuropils (see (Marin et al., 2024), for details on restricted INs). The pattern of cosine similarity for each muscle target and leg segment is generally consistent across all segments (Figure 7E). The inconsistencies seen in the T3 Tibia flexor and Acc. ti flexors are predominantly due to badly reconstructed and segmented MNs which resulted in low synapse counts.

*Other MNs*. For the remaining muscle categories (neck and abdominal), the *Drosophila* literature lacks detailed morphological descriptions and high-resolution LM data for their MNs, precluding identification of most MNs. For the neck MNs, light microscopy level identification of neck MNs are not yet available in *Drosophila*, although a comprehensive description of the neck muscle system and MNs is available in the blowfly, *Calliphora erythrocephala* (Strausfeld et al., 1987). In MANC, we identified 12 pairs of likely neck MNs (compared to 10 pairs in *Calliphora*) by their predominant presynaptic input in the NTct and exit through the cervical nerve (CvN) or dorsal prothoracic nerve (DProN). However, due to cross-species differences in MN morphology compared to *Calliphora*, as well as gross similarity between many *Drosophila* neck MNs, only 3 of these MNs can be tentatively identified (Supplementary file 3). Abdominal MNs exiting through AbN2-4 and AbNT, while reconstructed, grouped, and organized into serial sets, remain unidentified and will require future identification efforts.

Overall, the 733 MANC MNs were categorized into 168 types, composed of 319 groups with more than 1 member, and 14 singletons. We were able to identify 170 out of the 319 MN groups by light-level, EM-to-EM or serial homology matching methods. All MN and efferents with their light microscopy matches and accompanying references are listed in Supplementary file 3. Together, these identifications cover the majority of wing, haltere, and leg MNs, as well as several neck MNs, thus laying the groundwork for connectomics analyses of these motor systems.

### 2.4 DN network analysis

#### 2.4.1 DN to MN connectivity across the VNC

With the DN and MN populations fully identified and annotated, we were next able to examine large-scale organizational features of the networks that connect these inputs and outputs. We first asked whether DNs and MNs were found more commonly as individually-identifiable neurons (as left-right pairs) or as small populations (>2 DNs per group) that were difficult to distinguish by morphology or connectivity. The former is implicated in a more “command-like” form of motor control (von Philipsborn et al., 2011; von Reyn et al., 2014; Wang et al., 2020) and the latter in a more distributed representation (Dombrovski et al., 2023; Namiki et al., 2022). Using the innervation-based DN subclasses established in section 2.2, as well as the MN subclasses from section 2.3, we examined the pair versus population demographics across DN and MN categories (Figure 8A). Interestingly, a high proportion of population type DNs (∼75-100% of DN count) were found among those DN subclasses that broadly innervate the upper tectulum (DNut), or specifically target the neck (DNnt) or haltere (DNht) tectulums within the UTct, as well as among DNs innervating the intermediate tectulum (IntTct). The remaining DN subclasses had higher proportions of pairs (∼50-100%). Notably, a greater proportion of population DNs (∼50%) are found in the DNs innervating only the LegNpT1 (DNfl) compared to those innervating all leg neuropils (∼10%, DNxl).

**Figure 8:**
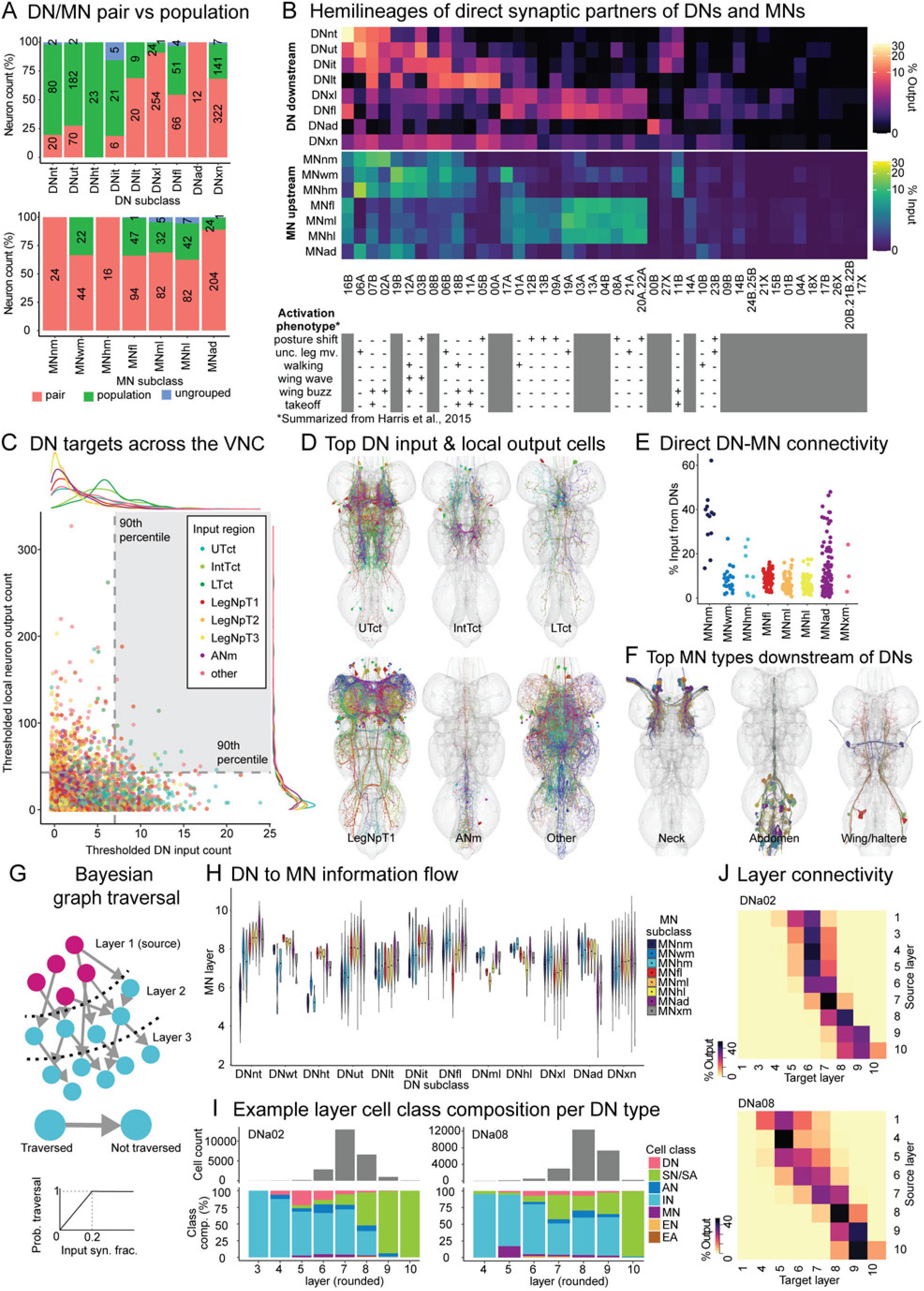
DN to MN connectivity across the VNC. **A.** Pair vs. population classification for all DN and MN types, as determined by the number of cells per neuron morphological group. DN and MN subclasses are defined in Figures 3 and 5. The DN subclasses DNwt, DNht, DNml and DNhl, and the MN subclass MNxm are not shown as they consist of few cells (3 groups or less). **B.** Hemilineage composition of DN downstream and MN upstream partners, compared to hemilineage motor activation phenotype as summarized from (Harris et al., 2015). Motor phenotypes are marked “+” if ≥50% of flies display it upon hemilineage activation in (Harris et al., 2015). **C.** All VNC intrinsic premotor groups (intrinsic, ascending, efferent and ascending efferent neurons) by their DN input count and local output count (Count of groups at a ≥1% groupwise input threshold). Intrinsic neuron groups are categorized by their primary input neuropil (≥50% postsynapses). **D.** Neuron groups with high DN input and local output (≥90th percentile for both) as defined in **C**. **E.** Direct DN to MN connectivity, shown by input fraction per MN group contributed by DNs. **F.** MNs with high DN input (≥20% total MN group input) from **E**. **G.** Bayesian graph traversal model for assessing distance from individual DN groups (source layer) to all VNC neurons (target layer). All neurons are assigned a layer based on their synaptic connectivity with neurons in prior layers, where the probability of traversal is scaled by the input fraction contributed by neurons of prior layers up to a probability of 1 for a synapse input fraction of 0.2 (further see Materials & Methods). Thus, layer assignments are a nonlinear measure of distance combining path length and synapse connectivity strength rather than a strict path length measure. **H.** MN groupwise layer means (rounded) for Bayesian graph traversal runs starting with individual DN groups as the source layer, classified by DN and MN subclasses. **I.** Example per-layer (rounded) cell class composition of Bayesian graph traversal for DNa02 and DNa08. **J.** Example layer-to-layer connectivity of Bayesian graph traversal for DNa02 and DNa08, showing the synapse output fraction of each source layer to all other layers up to 10 layers. For panels I and J, per-neuron layer is rounded, and layers without neurons are omitted from the axes.

**Figure 8–Supplement 1:**
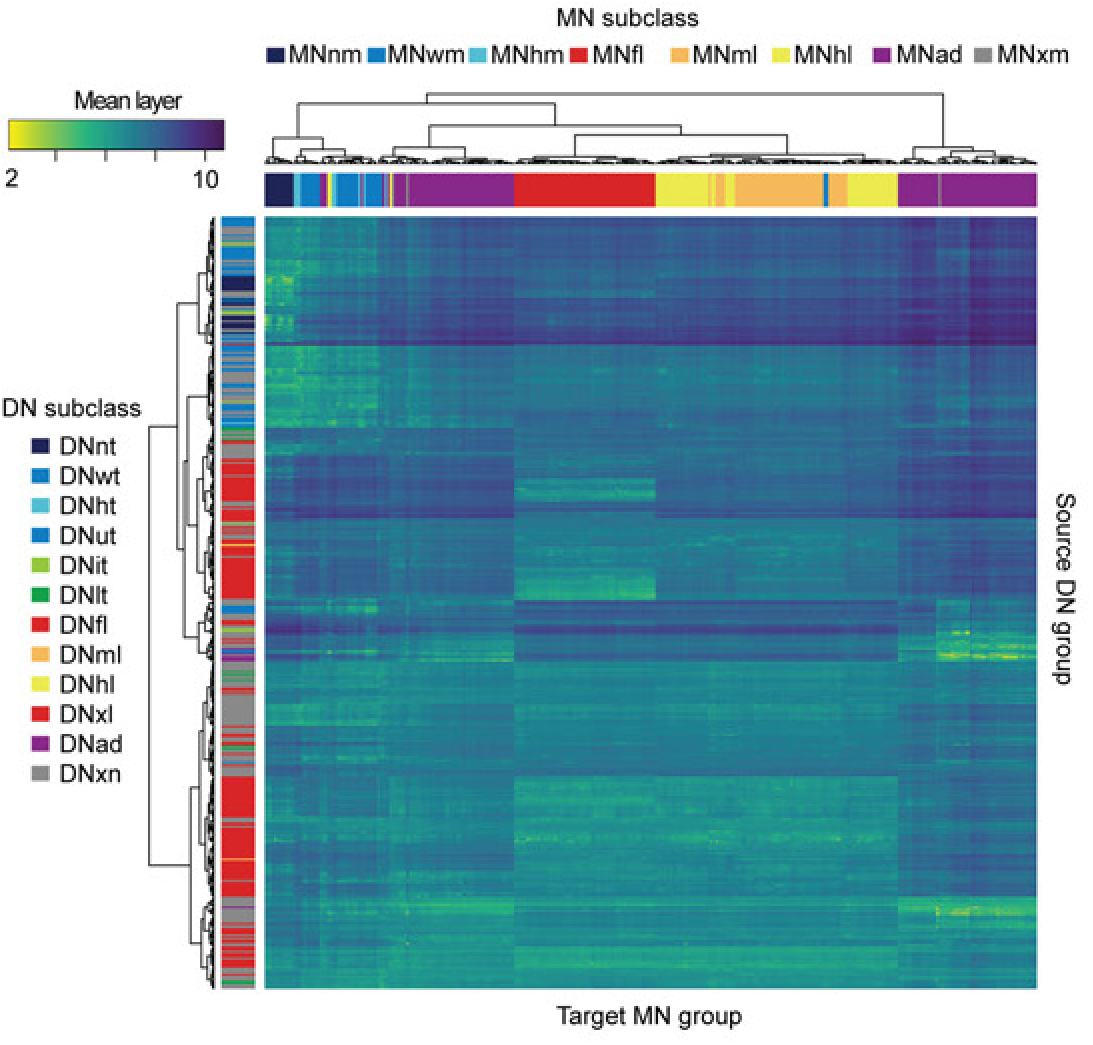
Bayesian graph traversal from DNs to MNs. Heatmap of all-to-all mean DN group to MN group layer distance for Bayesian graph traversal runs starting with individual DN groups as the source layer.

Population-type DNs likely enable population coding of information across individual cell activity (van Hemmen and Schwartz, 2008), allowing finer-grained representation of information sent from the brain compared to pair-type DNs. The preponderance of these DN types in the UTct, IntTct and LegNpT1 thus likely points towards the need for higher precision muscle control in the associated neck, wing, haltere and T1 leg motor systems. Differences in potential control precision between VNC regions can be further illustrated by the ratio of DNs to MNs targeting each neuropil. For example: In leg neuropils there are 404 DNs:392 leg MNs by neuron count, a ratio of about 1:1. In contrast, in the upper tectulum there are 381 DNs:106 MNs, a ratio of close to 4:1. More DNs per MN may indicate finer control of MN action. The differences in pair versus population demographics between wing and leg DN inputs was not reflected at the level of MN outputs, where population types were fewer (<30%) and similar across wing and leg MNs (Figure 8A), indicating that a transformation of population-coded descending information into discrete muscle activation signals occurs in the premotor networks between DNs and MNs.

Next, we asked whether particular DNs or MNs were more closely connected with neurons derived from specific developmental hemilineages, which are thought to define functional units that make up VNC premotor networks associated with specific actions (Harris et al., 2015; Shepherd et al., 2019). The hemilineage identity of MANC intrinsic neurons was determined in (Marin et al., 2024). We found high correspondence between the hemilineages connected to individual DN and MN subclasses that are expected to be functionally related (Figure 8B). For example, both upper tectulum DNs (DNut) and wing MNs (MNwm) share stronger connectivity with hemilineages 6A, 7B, 2A, 19B, 12A and 3B. Four out of the 6 of these hemilineages were documented in a previous optogenetic activation study (Harris et al., 2015) to have wing movement phenotypes when activated (Figure 8B, bottom panel). Similarly, DN- or MN-hemilineage connectivity was largely consistent with expected broad motor roles of individual DN and MN categories.

A few notable exceptions are observed, such as the association of NTct, UTct and IntTct DNs, and neck, wing and haltere MNs, with hemilineages 6A and 6B, whose activation largely causes uncoordinated leg movements, and less commonly wing-related motor activity (Harris et al., 2015). Interestingly, while the 6B hemilineage mainly ramifies the dorsal VNC, it also has relatively sparse projections into the leg neuropils (Marin et al., 2023). These and similar hemilineages may be associated with coordination of multiple motor circuits, e.g. both leg and wing circuits, or may have subpopulations with distinct motor roles. For hemilineages with no activation information, their connectivity with DN and MN subclasses allows a reasonable first guess at their motor function.

We next sought to identify which neurons were the strongest DN postsynaptic partners, with likely strong influence on VNC motor circuits. We plotted all neuron groups of the VNC (excluding DNs, MNs and sensory neurons) by the number of DN groups from which they receive input and the number of downstream intrinsic neuron groups they output to at a ≥1% synapse input threshold (Figure 8C). We color coded neuron groups by the neuropils they received ≥50% input from. Across the VNC, individual neuron groups received input from up to 23 DN groups and provide output to up to ∼350 intrinsic neuron groups. Neuron groups categorized as IntTct or LTct received input from more DN groups on average (∼5 groups) compared to intrinsic neurons from other regions (∼1-2 groups). Neuron groups with a large amount of DN input and/or interneuron output (above the 90th percentile) are further plotted in Figure 8D. Notably, a subset of LTct neurons with high DN input and high output project widely in the VNC, and their role in LTct circuits are further followed up in section 2.7. The top targets of DNs in the UTct and ANm included several MNs, and we thus plotted direct DN input to MNs separately (Figure 8E, F) to examine what fraction of MN input comes directly from DNs. Most neck MNs, a subset of abdominal MNs, and several wing and haltere MNs receive >20% (up to ∼60%) of their synaptic input directly from DNs, suggesting that they are under relatively direct control of DNs. Most other MNs receive a relatively low proportion of their input from DNs (MN groupwise mean of 7%), although ∼80% of MN groups receive direct input from at least one DN group at a ≥1% groupwise input threshold, suggesting that control of most MNs comes from VNC intrinsic premotor circuits, but that direct input from brain signals via DNs still exerts some influence.

We have thus far been considering direct connections between DNs and their postsynaptic partners. To more broadly examine indirect connectivity from DNs to MNs, we next applied a Bayesian graph traversal model (based on (Marin et al., 2024; Schlegel et al., 2021) to the MANC network. Starting from a set of seed neurons, the model traverses the neurons of the entire VNC and assigns a ‘layer’ position to each cell by Bayesian probability, where the traversal probability for each node is determined by linearly scaling the input synapse connectivity between the node and all nodes of prior layers, up to a maximum of 20% synaptic input which corresponds to a traversal probability of 1 (Figure 8G). The layer position is thus a relative nonlinear measure of distance from the source neurons that quantitatively combines path length and synapse connectivity strength, although layer position assignments do not correspond directly to path length. We observed that DN subclasses generally have closer layer positions to MN subclasses that are putatively expected to receive control by those same neuropils. For example, the layer distance from haltere tectulum DNs (DNht) to MNs in the neck, wing, and haltere neuropils (MNnm, MNwm, MNhm) are about half the distance as that from DNht to leg MNs (MNfl, MNml, MNhl) (Figure 8H). In addition, MNs of the neck and abdominal subclasses have the lowest mean layer score from upstream DNs, suggesting that neck and abdominal MNs are more directly controlled by DNs than other MN subclasses, in agreement with our direct connectivity analysis (Figure 8E). The mean layer assignment of all DN-MN group combinations is shown in Figure 8–Supplement 1. Overall, these results indicate that there is likely segregation of premotor circuits by function, e.g. flight, which requires control of neck, wings, and halteres together, that also corresponds to anatomical separation of the DNs by neuropil. Examples of layer cell class composition and layer-to-layer connectivity for two DN types (DNa02 and DNa08) are shown in Figure 8I, J. For both DNs, neuron layer assignment is relatively sparse and specific in initial layers–these cells are likely strong downstream partners of these DNs–while later layers (around layers 7-8) ‘fan out’ to reach most of the connectome. Layer-to-layer connectivity shows that each layer commonly has output to itself and the next layer, consistent with flow of information from one layer to the next. This trend is only noticeably different for the initial few layers and last 1-2 layers, possibly due to the paucity of cells assigned to those layers.

#### 2.4.2 Community structure of VNC networks

Our analyses above indicate that, although some motor control is accomplished by direct connectivity of specifically-targeted DN terminals synapsing onto MNs for a particular limb, most paths from DNs to MNs are indirect, involving multiple layers of intrinsic neurons between DN and MN. As specialization of motor functions is evident in the VNC at the hemilineage and neuropil level (Harris et al. 2015; Court et al. 2020; Marin et al. 2024), VNC neurons may plausibly be organized into premotor ‘modules’ that each control separate motor functions. To gain further insight into the structure of intrinsic premotor circuits (here comprised of all cells except DNs, sensory neurons and MNs), we investigated whether connectivity in this network was uniform, or whether subsets of intrinsic neurons tended to synapse more strongly with each other, forming separable connectivity clusters, or communities. To characterize community structure, we applied a commonly used graph community detection method called the Infomap algorithm. This community detection method partitions the nodes of a weighted, directed graph (such as a connectome) by information flow, namely by minimizing the description length of the movements of a random walker within the network as described by the map equation (Rosvall et al., 2009; Rosvall and Bergstrom, 2011, 2008; Smiljanić et al., 2021). The Infomap method thus provides a coarse-grained description of information flow across neurons of the VNC. We applied Infomap community detection to a directed graph of all VNC intrinsic premotor neurons using synapse weights as edge weights. We used regularization using a Bayesian prior network to account for missing connections and multi-level clustering in the Infomap algorithm to allow splitting into nested communities; however, multi-level clustering showed that the two-level solution was optimal (i.e. splitting the VNC cells once into a non-nested set of communities).

The Infomap algorithm partitioned the VNC into 37 communities (Figure 9A, 9–Supplement 1A, B) (excluding ∼10 with no VNC synaptic input or output), of which 18 consist of 50 cells or more (Figure 9B, neuron community assignments in Supplementary file 4). The connectivity of these communities with each other are shown in Figure 9A, C. Furthermore, the vast majority of these communities have stronger intra-community connectivity than inter-community connectivity (Figure 9–Supplement 1B). Labels for putative motor functions were assigned by community neuropil arborization (Figure 9D) and connectivity with DN and MN subclasses (Figure 9E, F). Notably, we observe one community per leg neuropil per side that largely corresponds to local leg networks, five communities covering the upper tectulum, and three in the ANm. We also observe three communities that each include neurons from multiple intermediate regions of the VNC and one in the mVACs, which all have low direct MN output. To probe the correspondence of Infomap communities with possible motor control functions, we further examined the connectivity of these communities with DNs and MNs by subclass (Figure 9E, F). Overall, most Infomap communities have relatively specific connectivity with DN and MN subclasses, which suggests that each community may be a premotor network for a specific motor role. For example, communities 8, 9, and 10 receive input from upper tectulum DNs (DNut) and provide output to wing motor neurons (MNwm). Similarly, communities 5 and 6 receive input from front leg DNs (DNfl) and output to front leg motor neurons (MNfl). Interestingly, in support of its hypothesized role in coordination of wing-leg behaviors, lower tectulum DNs (DNlt) target communities 7, 8, and 14—community 7 is notable as the largest VNC community (∼1800 cells) and is the top non-self input to the majority of other communities (Figure 9C, 9–Supplement 1B), while community 14 has little to no connectivity to MNs but itself outputs to communities labeled for both wing and leg functions. This raises the possibility that these communities play a key role in controlling or coordinating motor output across the VNC via control of other communities. Indeed, lower tectulum DNs that have been implicated in escape takeoff behavior (which requires leg-wing coordination) preferentially target this community, and are followed up on in section 2.7. Finally, we note that the Infomap method is one of many graph community detection algorithms currently available, and it is not yet known which algorithms are most optimal for application to connectome data. Thus, the community detection results above offer a first look at the organization of VNC circuits but will require further verification of their functional significance.

**Figure 9:**
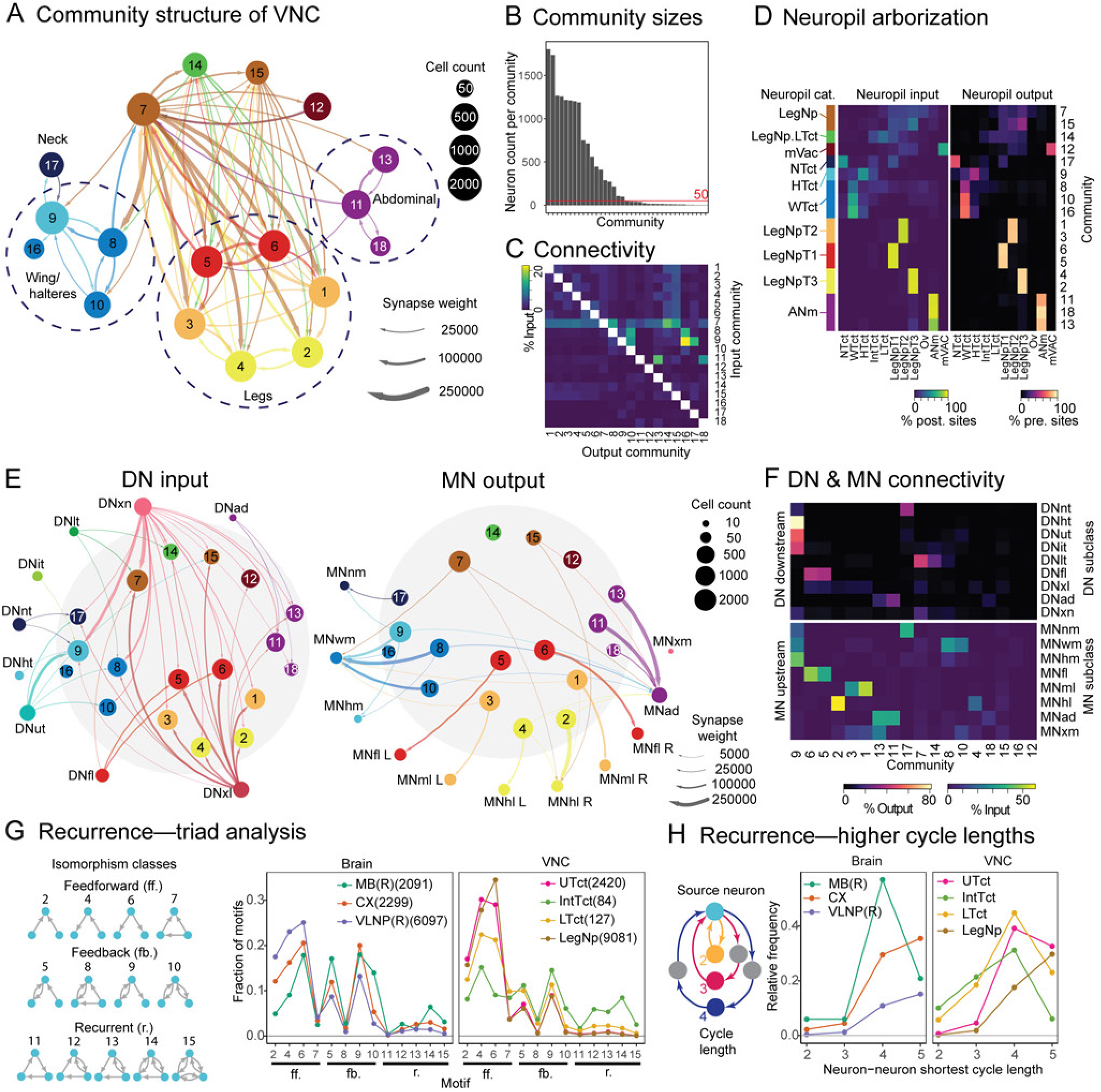
Structure of VNC networks. **A.** Community structure of VNC intrinsic premotor networks (all neurons excluding descending, sensory and motor) as partitioned by the Infomap algorithm (Rosvall et al., 2009; Rosvall and Bergstrom, 2011, 2008; Smiljanić et al., 2021). Communities are numbered arbitrarily, and communities of at least 50 cells are shown on graph and colored by their constituent neurons’ primary input and output neuropils (see **C**). Edges between communities are thresholded at ≥25000 synapses. Dotted circles and their labels indicate putative broad motor function based on neuropil arborization and connectivity with MN subclasses (see **C & D**). **B.** Neuron count of Infomap communities. Communities on the x-axis are rank-ordered by decreasing neuron count. **C.** Infomap community-to-community connectivity for communities of ≥50 cells. Full community-to-community connectivity including intra-community connectivity is shown in Figure 9–Supplement 1B. **D.** Community neuropil input and output, calculated by the % neuropil input and output of all neurons in the community. **E.** Community DN input and MN output by DN/MN subclass. Connectivity in graphs are thresholded at ≥5000 synapses. Light shaded area in graphs correspond to Infomap communities as shown in A. The DN subclasses DNwt, DNml and DNhl consist of only a single DN group each and are not shown. MN subclasses for leg MNs (MNfl, MNml, MNhl) are further split into left and right categories (L and R). **F.** Heatmaps of community DN input and MN output. **G.** Recurrence in VNC networks as evaluated by triad census, compared to brain neuropils with high recurrence (mushroom body, MB, and central complex, CX) and low recurrence (right ventrolateral neuropils, VLNP(R)). Triad motifs are classified by their isomorphism class into feedforward, feedback and recurrent categories. Synaptic connectivity between neurons are thresholded at ≥0.1% input. Neurons per neuropil are subsetted by having ≥50% input and output in the target neuropil, and further excluding DNs, sensory and MNs. **H.** Recurrence in neuropil networks as evaluated by all-to-all neuron shortest cycle length at a synapse threshold of ≥0.1% input. Neurons per neuropil are subsetted by having ≥50% input and output in the target neuropil, and further excluding DNs, sensory and MNs.

**Figure 9–Supplement 1:**
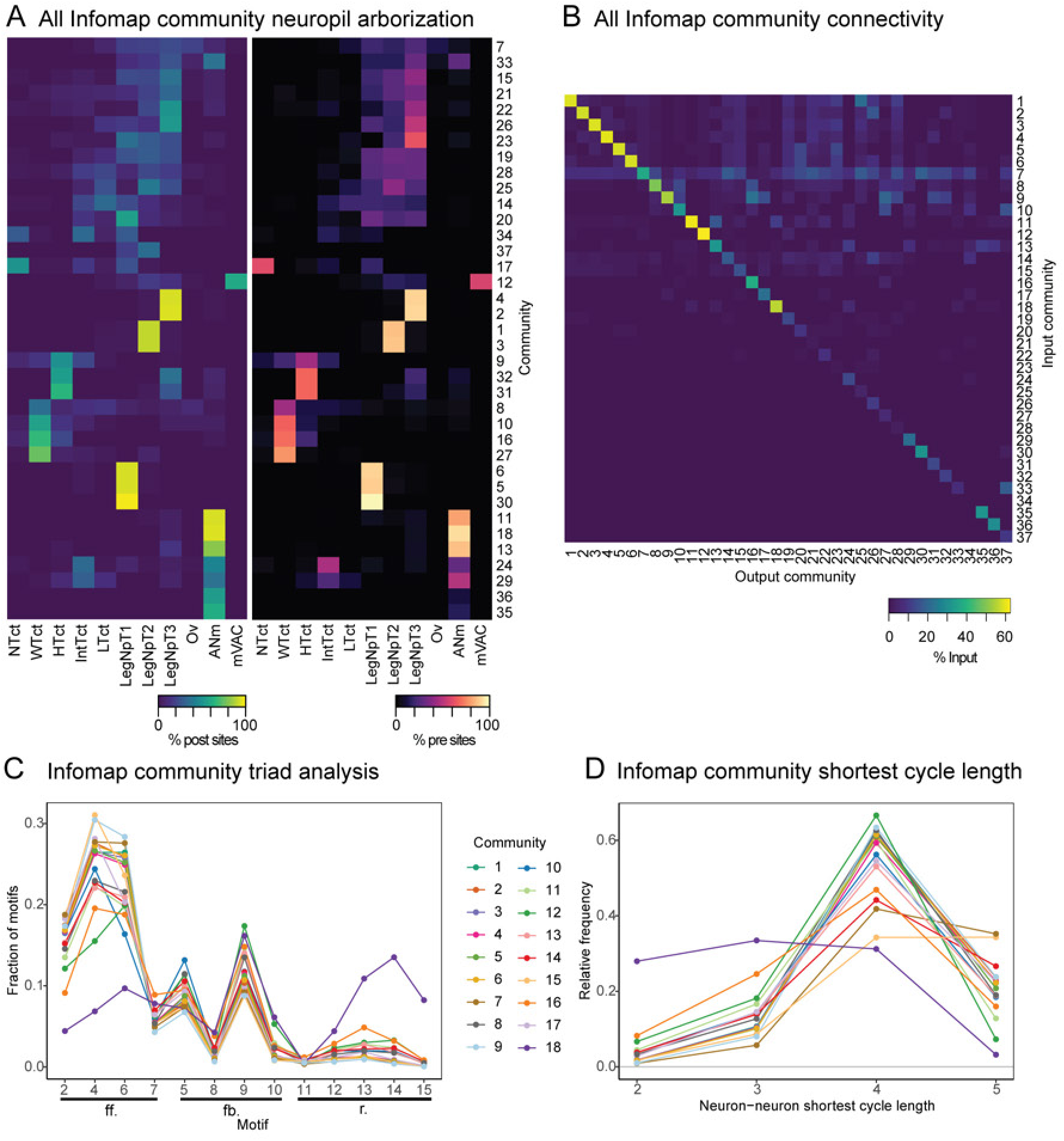
VNC Infomap communities and within-community recurrence. **A.** Community neuropil input and output for all Infomap communities (excluding communities with 1 neuron, or that have zero pre- or postsynaptic sites), calculated by the % neuropil input and output of all neurons in the community. **B.** All Infomap community-to-community connectivity. **C.** Recurrence in communities with ≥50 cells as evaluated by triad census. Synaptic connectivity between neurons are thresholded at ≥0.1% input. **D.** Recurrence in communities with ≥50 cells as evaluated by all-to-all neuron shortest cycle length at a synapse threshold of ≥0.1% input.

Our community analysis looked at the macrostructure of the VNC network, but what about the microstructure? Recurrent connectivity may plausibly play roles in local modulation of VNC circuits and generation of the patterned motor signals required for rhythmic motor actions. To probe the degree of recurrence present in VNC circuits, we quantified the proportion of all triad motifs in different sets of neuropils of the intrinsic VNC network (Figure 9F). A neuron was defined to be within a neuropil if ≥50% of its input and output resided in the neuropil. For comparison, we performed the same analysis on brain regions from the Hemibrain connectome (Scheffer et al., 2020) with known high recurrence such as the mushroom body and central complex, or expected low recurrence such as the ventrolateral neuropils. We found that the entire VNC in general, as well as the UTct and all six leg neuropils have a lower proportion of cyclic triad motifs than the mushroom body and central complex, and comparable levels to the ventrolateral neuropils (Figure 9G), suggesting that VNC circuits generally operate in a more feedforward manner. Yet, the relatively small set of LTct-and IntTct-restricted cells have higher cyclic triad motifs at a similar frequency to the mushroom bodies and central complex, perhaps suggesting that these neurons are higher order circuits that utilize recurrence to carry out more complex motor functions. LTct circuits are further discussed in section 2.7. These results are corroborated by a different analysis in which we looked at all-to-all neuron shortest cycle length (Figure 9H). This analysis showed that short-length recurrence occurs with lower frequency in UTct and LegNp circuits and higher frequency in IntTct and LTct local circuits, at rates comparable to the mushroom body and central complex. Interestingly, LTct restricted neurons are included in the Infomap communities (e.g. 7 and 14) without direct links to MNs that we hypothesized to be higher level premotor circuits, whereas IntTct restricted neurons are largely part of community 9, which contributes to wing and haltere motor control. This observation suggests that similar microstructure of connectivity (i.e. recurrent motifs between individual neurons) may play different roles at different macrostructural levels of the network (lower vs. higher-level control of motor output)–recurrence may plausibly play roles in coordination or rhythmic control of wing and haltere muscles in the IntTct, and coordination of different motor systems or even computations determining behavioral output in the LTct. However, Infomap partitioning suggested that some premotor network communities cross neuropil boundaries–for example, LTct-restricted cells are a small part of larger communities that span the LTct and all leg neuropils. Thus, we also examined recurrence in Infomap communities with ≥50 cells (Figure 9–Supplement 1C, D). While overall recurrence in the communities that include IntTct and LTct-restricted cells are less obvious due to inclusion of cells not restricted to these neuropils, we further observe high recurrence in an abdominal community (community 18). The function of recurrence in abdominal circuits remains to be investigated.

In summary, the direct synaptic partners of DNs and MNs are hemilineage-restricted in a manner consistent with putative DN and MN subclass function, and some MN subclasses (in particular neck and abdominal) further receive strong direct input from DNs. By a Bayesian graph traversal model, we help further define sets of DNs that target individual MNs and MN subclasses, as well as compare relative distances of MN subclasses from DNs. Community structure of VNC intrinsic premotor neurons suggests that VNC premotor circuits are organized into communities that are each dedicated to control of specific MN subclasses, as well as several communities that have little direct MN control but instead have strong output to other communities, potentially for higher-order motor control. Finally, by analyses of recurrence, larger VNC regions like the UTct and all leg neuropils likely operate in a relatively feedforward manner, but the small circuits consisting of IntTct and LTct restricted cells instead have more recurrent motifs. This microstructure in these regions may aid in their respective motor roles, i.e. as effectors of motor output vs control and coordination of motor systems. In the following three sections we take a closer look at the microstructure of connectivity within different circuits of the VNC: the wing circuits contained in dorsal communities, the leg circuits contained in ventral communities, and the potential higher order circuits of the intermediate neuropil.

### 2.5 Organizational logic of DNs onto wing circuits

#### 2.5.1 Descending neurons and networks underlying wing motor control

In the fly, the wing motor system is crucial to behaviors such as flight (Dickinson and Tu, 1997), aggression (Kravitz and Fernandez, 2015; Zwarts et al., 2012) and courtship (Greenspan and Ferveur, 2000; Yamamoto and Koganezawa, 2013). The VNC houses premotor circuits that govern the control of the wing motor system and patterning of its motor output during these diverse behaviors (Harris et al., 2015; Yu et al., 2010). The MANC connectome allows us comprehensive access to their synaptic connectivity to begin dissecting the architecture of wing motor circuits and their control by descending signals. Here, we first broadly survey the direct and indirect connectivity of DNs to the wing MNs, and specifically focus on describing circuit connectivity and structure in relation to potential roles in flight. Wing control circuits in relation to courtship in MANC are discussed in detail in (Lillvis et al., 2024).

As previously discussed, *Drosophila’s* wing muscle system consists of three classes of muscle: the indirect power (PMs), steering (SMs) and indirect control muscles (ICMs) (Dickinson and Tu, 1997). The asynchronous PMs act to generate power for wing vibrations–the dorsal longitudinal muscles (DLMs) and dorsal ventral muscles (DVMs) function antagonistically, each taking turn to contract while stretch-activating the other to vibrate the thorax (Dickinson and Tu, 1997). These thoracic vibrations are thought to be transmitted to the wings through a clutch and gearbox mechanism formed by wing hinge elements, which functions to couple or isolate the wing from thoracic vibrations and further modulate wingbeat amplitude (Deora et al., 2015; Miyan et al., 1985; Miyan and Ewing, 1988; Nalbach, 1989). The wing hinge is in turn controlled by the relatively small SMs, which fire either tonically or phasically during flight in a manner phase-locked to the wingbeat cycle (Fayyazuddin and Dickinson, 1999, 1996; Trimarchi and Murphey, 1997) to accomplish steering (Deora et al., 2017; Dickinson and Tu, 1997) (also see Figure 6A). The mechanics of the wing hinge is complicated, and a comprehensive model of how SMs control flight maneuvers remains elusive. However, lateralized activation patterns of SMs during turning in flight have been well-documented (Table 1). The action of two of the three ICM categories–the pleurosternal (ps) and tergopleural (tp) muscles—are less well understood but are thought to function during flight to control thoracic stiffness and resonant properties, thereby indirectly modulating wingbeat amplitude (Dickinson and Tu, 1997). The tergotrochanteral muscle (TTM) functions in flight initiation during escape and is quiescent during flight (Dickinson and Tu, 1997), and thus its control is separately addressed in section 2.7. Lastly, wing control during flight is intimately tied to mechanosensory input from the halteres, which possess a reduced set of muscles that are homologous to the wing muscles, which indirectly impact wing steering during flight (Dickerson et al., 2019). Thus, we also examine the descending control of haltere MNs below.

**Table 1:** Dipteran steering muscle function. Activity of steering muscles in *Drosophila* and other dipterans during turning in flight on the ipsilateral (inner side of turn) vs contralateral (outer side of turn) sides. ‘Tonic’ and ‘phasic’ designations as defined in Lindsay et al., 2017: tonic muscles show elevated activity throughout flight, while phasic muscles are intermittently active only during specific flight maneuvers. Evidence compiled from (Egelhaaf, 1989; Heide, 1975; Heide and Götz, 1996; Lehmann and Götz, 1996; Lindsay et al., 2017; Nachtigall and Wilson, 1967). The iii2 muscle, which is not present in *Drosophila,* is omitted.

| Muscle | Evidence type | Activity during flight | Activity during turning |  |
| --- | --- | --- | --- | --- |
|  |  |  | Ipsilateral | Contralateral |
| b1 | Electrophysiology, calcium imaging | Tonic | Phase delay | Phase advance |
| b2 | Electrophysiology, calcium imaging | Phasic | Inactive | Active |
| b3 | Calcium imaging | Tonic | Activity increased | Activity decreased |
| i1 | Electrophysiology, calcium imaging | Phasic | Active | Inactive |
| i2 | Calcium imaging | Tonic | Active/minor change | Active/minor change |
| iii1 | Electrophysiology, calcium imaging | Phasic | Inactive | Active |
| iii3 | Calcium imaging | Tonic | Activity decreased | Activity increased |
| iii4 | Calcium imaging | - | Minor/inactive | Minor/inactive |
| hg1 | Electrophysiology, calcium imaging | Phasic | Active | Inactive |
| hg2 | Calcium imaging | - | Minor/inactive | Minor/inactive |
| hg3 | Electrophysiology, calcium imaging | Phasic | Active | Inactive |
| hg4 | Calcium imaging | Tonic | Activity increased | Activity decreased |

To examine DNs that likely contribute to wing motor control, we first narrowed down DN types that provided a groupwise contribution of at least 10% and 100 presynaptic sites (raw count) to the WTct and HTct. The full set of WTct/HTct DNs is shown in Figure 10A, with example morphologies of several DNs implicated in flight-related behaviors shown in Figure 10B and followed up on in the next sections. Of the WTct/HTct DNs, only a single pair of DNs (DNa08) has presynaptic sites restricted to only the WTct, while most other DNs that innervate the WTct also innervate the NTct and HTct (Figure 10C), suggesting that descending motor signals often simultaneously control neck, wing and haltere motor systems. Similarly, neurons that are directly downstream (≥1% groupwise input) of these DNs largely target the WTct and HTct, with smaller sets targeting other neuropils (Figure 10C). Both WTct/HTct DNs and their direct downstream cells target mainly intrinsic neurons (INs), with MNs as the next strongest targets (Figure 10C). To further probe the top targets of WTct/HTct DNs, we examined downstream interneuron targets of WTct/HTct DNs by the group-to-group number of ‘strong’ DN inputs (≥1% synaptic input and ≥50 raw synapse weight, groupwise) they receive, as well as the Infomap community assignments established in section 2.4 (Figure 10D). Indeed, the communities that receive the largest number of WTct/HTct DN input groups are expected to be associated with wing and haltere motor control–i.e. communities 8, 9, 10 and 16–with individual neuron groups receiving input from up to 19 DN groups (example morphologies in Figure 10E). Next, to further examine finer details of community structure and their relation to DN input, we carried out a series of Infomap community detection using individual communities as input graphs and selecting an option to prefer modular solutions (as the two-level clustering from section 2.3 is the ‘optimal’ solution). This method divided up communities 8, 9 and 10 into further subcommunities (Figure 10F), and WTct/HTct DNs could be hierarchically clustered by their input to subcommunities to yield five clusters. Wing-related communities appear specialized in their wing motor functions, with community 8 and 9 most strongly targeting steering MNs, and community 10 targeting power MNs. Both communities 8 and 10 further target indirect control MNs. The breakdown of subcommunity connectivity with wing and haltere MNs is shown in Figure 10–Supplement 1A. Furthermore, community 8 and its subcommunities may be implicated in courtship-related behaviors, as many neurons that are likely positive for *fruitless* (identified by (Lillvis et al., 2024)), a transcription factor necessary for sex-specific neuronal development, are found within these communities (Figure 10–Supplement 1B).

**Figure 10:**
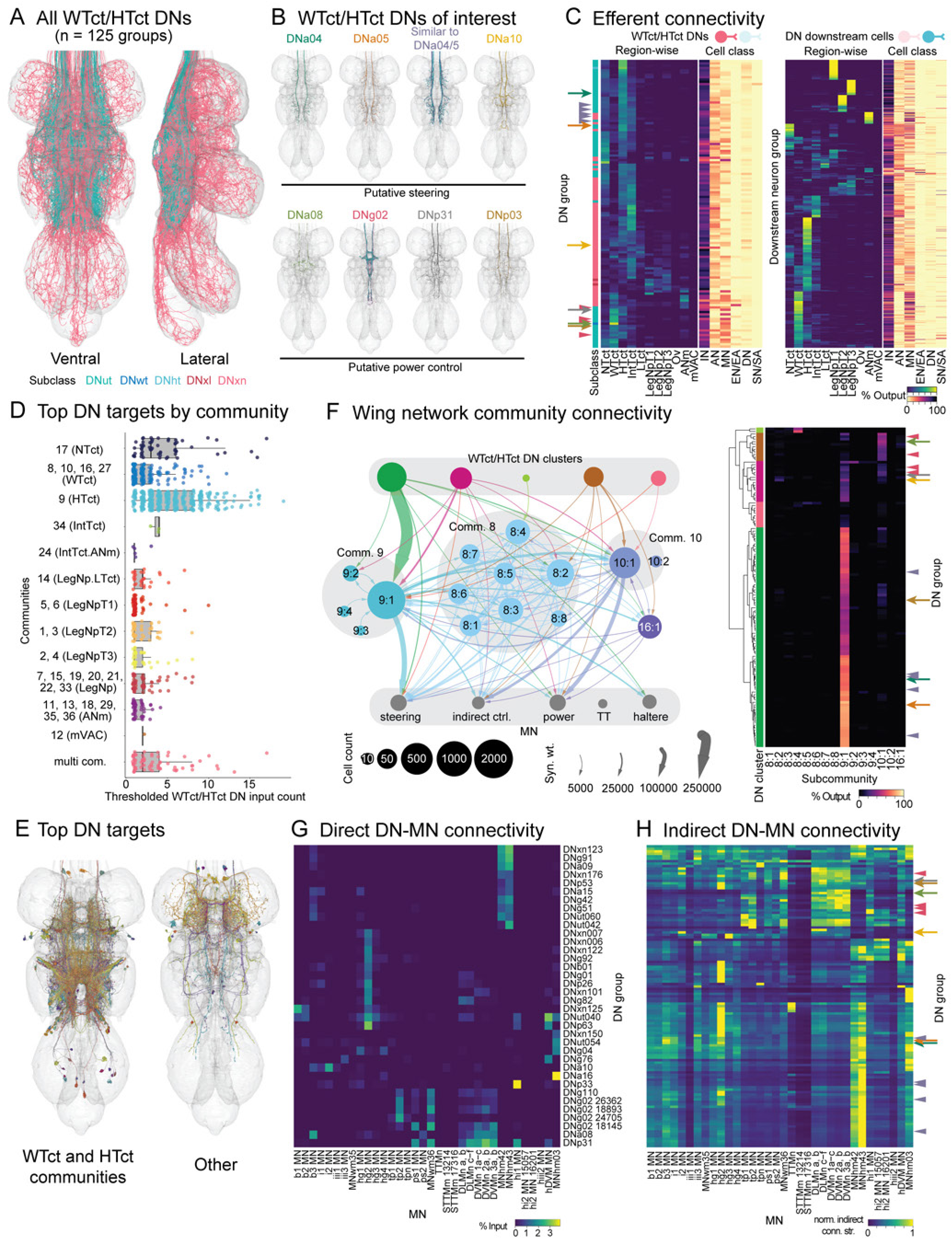
Overview of wing and haltere-related DNs and networks. **A.** DN groups with ≥10% output (groupwise mean) and ≥100 presynaptic sites (groupwise sum) in the wing tectulum (WTct) and haltere tectulum (HTct). DNs are categorized and colored by their subclass as defined in Figure 3. **B.** WTct/HTct DNs of interest that may be implicated in flight motor control. The DN category ‘similar to DNa04/05’ comprises the DN types DNa09, DNa15, DNut060, DNut037 and DNxn123. **C.** Neuropil-wise and cell class-wise efferent connectivity of WTct/HTct DNs (left) and their direct downstream targets that receive ≥1% groupwise input from WTct/HTct DNs (right). In this and subsequent heatmaps, DNs of interest from **A** are shown with arrows (DN pairs) or arrowheads (DN populations) with the same color as DN labels in **A**. **D.** DN input count (number of WTct/HTct DN groups providing ≥1% groupwise input) of WTct/HTct DN downstream intrinsic premotor neuron groups (IN/AN/EN/EA), classified by top input/output neuropil of their VNC Infomap communities (see Figure 9). Groups in the “multi com.” category have individual neurons assigned to different communities. **E.** Morphologies of top WTct/HTct DN targets from **C** that have input from 10 or more DN groups with ≥1% input from each DN group. **F.** Community structure of wing-related Infomap communities and their connectivity with WTct DNs and wing/haltere MNs. Subcommunities were defined by a second round of Infomap community detection on cells of each community. DN grouping is defined by hierarchical clustering of individual WTct/HTct DN groups’ output to subcommunities (right heatmap). Edges on graph are thresholded at ≥2000 synapses. The T2 tergotrochanteral MNs (TT) are shown separately from the other indirect control MNs as their input is largely distinct (see Figure 6E). **G.** Direct connectivity from WTct/HTct DNs to wing and haltere MNs. DN groups shown have ≥1% input to at least 1 MN group. DNs and MNs are named by type, with group appended if multiple groups share a type. **H.** Indirect WTct/HTct DN-MN connectivity by normalized indirect connectivity strength (see Materials & Methods). DN groups shown have ≥5 normalized indirect connectivity strength values to a MN group. Normalized indirect connectivity strength is then rescaled to a maximum of 1 row-wise (DN groupwise) for display here.

**Figure 10—Supplement 1:**
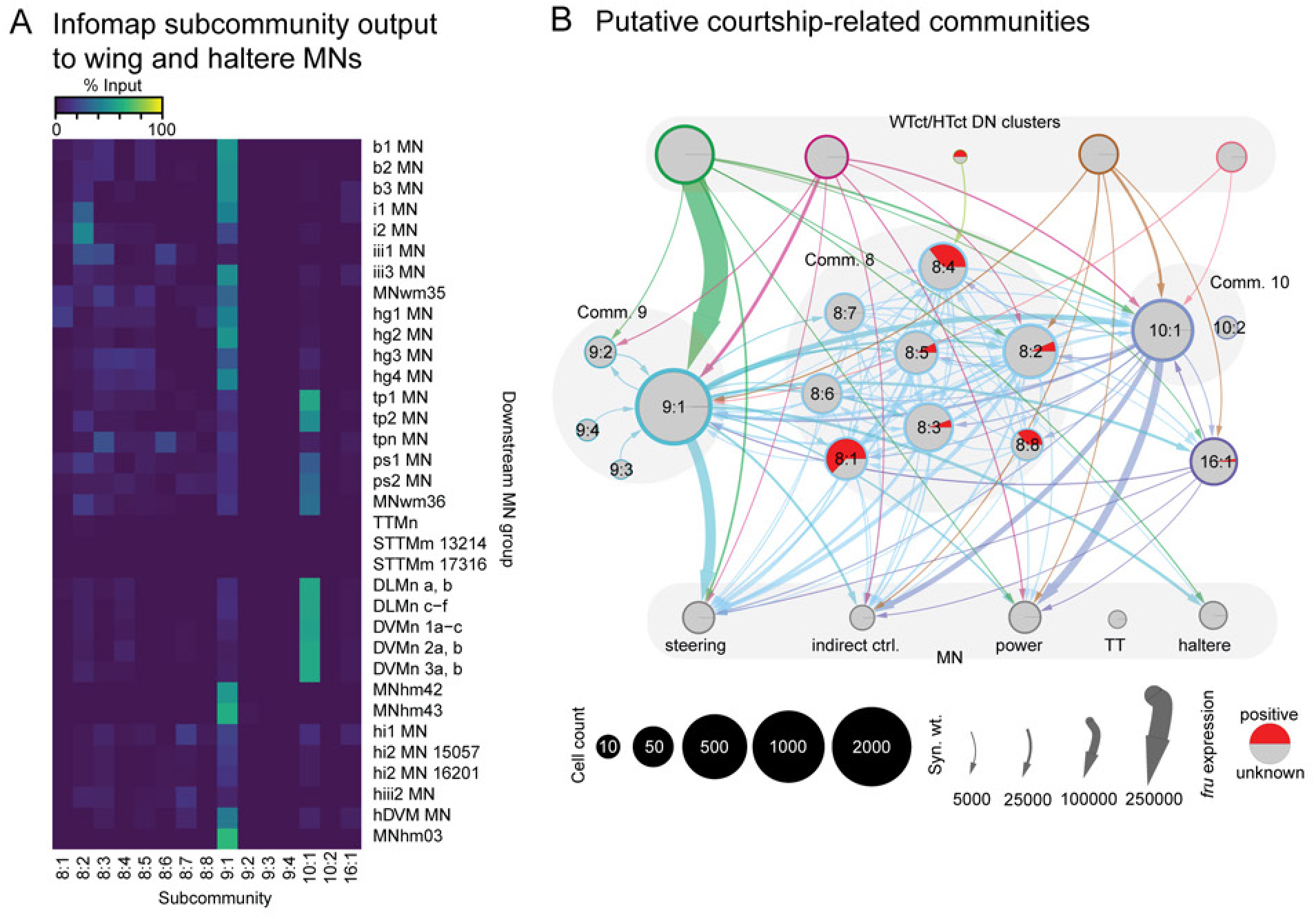
Connectivity of wing network Infomap subcommunities to MNs. **A.** Synaptic input fractions contributed by individual Infomap subcommunities to wing and haltere MNs of a group. **B.** Putative courtship-related Infomap subcommunities as suggested by LM-EM matching of neurons to *fruitless*-positive neuron types (Lillvis et al., 2024). Edges on graph are thresholded at ≥2000 synapses.

Lastly, we sought to define at the DN type level the combinations of wing MNs and haltere MNs (also implicated in flight control) that each DN controls. To do this, we plotted direct connectivity from WTct/HTct DNs to MNs via groupwise synaptic input fractions, and indirect connectivity via a measure of indirect connectivity strength. Briefly, the metric of indirect connectivity strength (Li et al., 2020) is calculated by multiplying the synaptic input fractions for all cells forming a path between a source and target cell for a given path length, and summing them across all paths of same length between source and target cells (accomplished by matrix multiplication of adjacency matrices). To compare indirect connectivity strength across multiple path lengths, we further normalized the raw values per path length per source neuron by their non-zero mean, and the largest normalized value among all path lengths (in practice up to 5) is taken as the indirect connectivity strength. For direct connectivity, roughly 30% of WTct/HTct DNs provide substantial direct input (≥1% groupwise input) to one or more wing or haltere MNs (Figure 10G), most commonly with single DN groups dedicated to steering MNs or a mixture of power MNs and indirect control MNs. These trends are similar for indirect connectivity strength between WTct/HTct DNs and MNs (Figure 10H), but with several DNs providing more widespread input across MN muscle categories. Thus, WTct/HTct DNs are specialized in the MN categories and intrinsic neuron communities they target, which likely reflects their individual roles in wing-related behaviors.

#### 2.5.2 Laterized descending control of steering muscles mediated by the wing contralateral haltere interneurons

Flies are exceptionally nimble during flight, capable of executing aerial maneuvers involving minute wing kinematic changes (Dickinson and Muijres, 2016) to change their trajectory in the air. The steering muscles (SMs) are key to such maneuvers, and left-right asymmetric activation of up to nine of these muscles has been documented during flight turns (Figure 11A, Table 1). Prior work identified a DN (named AX) whose activity correlated with rapid, saccadic turning maneuvers in flight in response to visual motion and is thus a candidate command neuron for flight saccades (Schnell et al., 2017). While this DN’s identity in MANC has yet to be determined, its rough morphology is similar to both DNa04 and DNa05, which are part of a set of DNs containing at least 5 other members with similar VNC morphology (Figure 10B). Both DNa04 and DNa05 are cholinergic DNs that receive input from the posterior slope in the brain, an area implicated in wide-field motion responses, and DNa04 further receives input from the central complex which encodes fly heading direction (Hulse et al., 2021; Namiki et al., 2018). We thus set out to investigate the connectivity downstream of DNa04, DNa05 and similar DNs to determine their potential role in flight motor control and steering.

**Figure 11:**
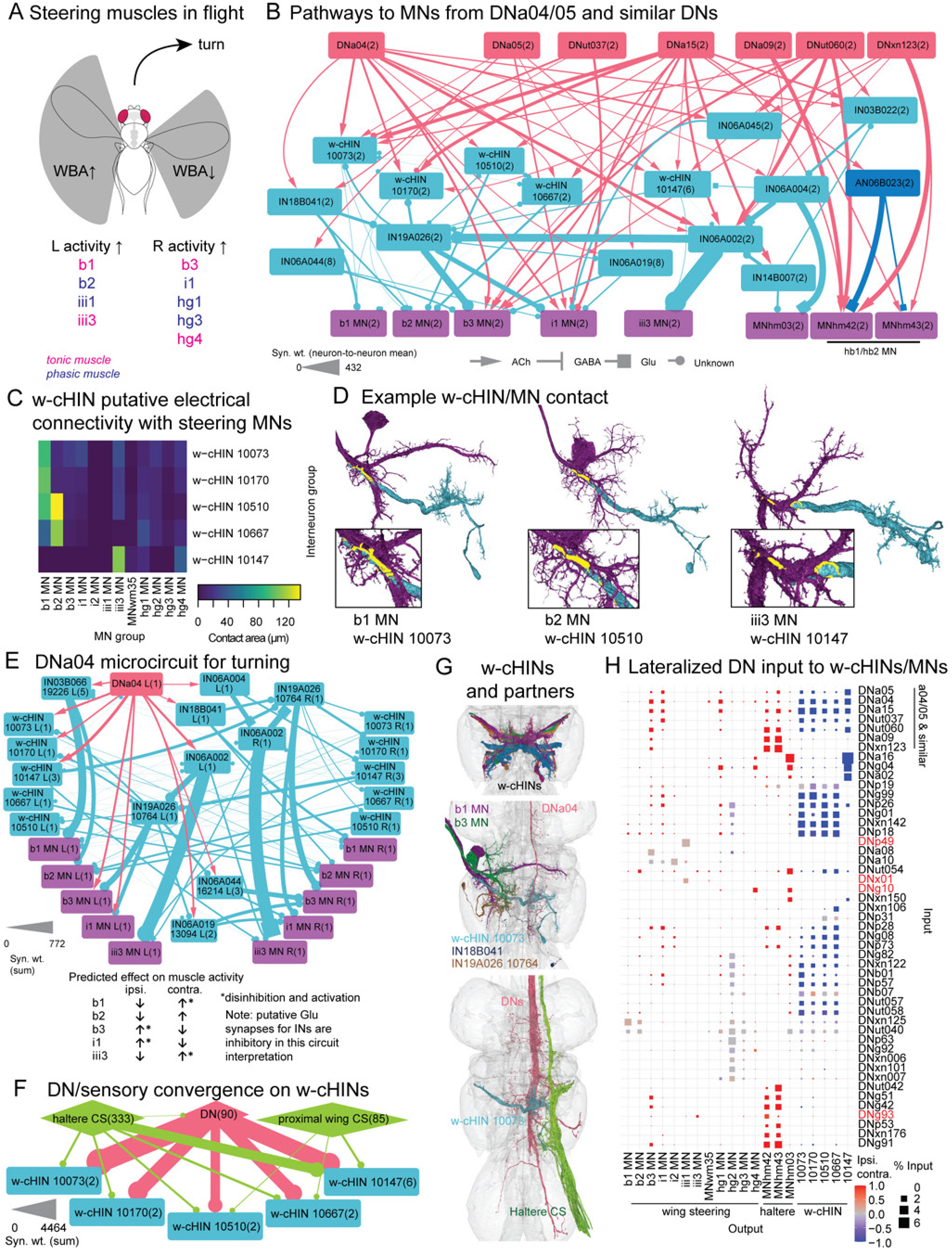
Putative flight steering circuits through DNa04/05 and w-cHINs. **A.** Modulation of fly wingbeat amplitude (WBA) and steering muscle (SM) activity during turning in flight. SM activity is simplified from (Lindsay et al., 2017) (also see Table 1). **B.** Pathways of DNa04, DNa05 and other DNs with similar VNC morphology to steering MNs and haltere MNs. Node labels are neuron type, appended with a five- or 6-digit number for neuron group if a subset of neurons of a type are shown, while number in parentheses indicates cell count per node. Synapse weight (neuron-to-neuron mean) is thresholded at ≥20, except for connectivity of wing contralateral haltere interneurons (w-cHINs) to MNs which may have electrical synapses (see **C**). **C.** Mean surface contact area of single w-cHINs of a group to all MNs of a group, suggestive of electrical synaptic connectivity. Contact area is calculated using 3D neuron meshes derived from EM segmentation; for each MN/w-cHIN pair, meshes are expanded by 0.1 μm, and the intersection surface area is calculated. **D.** Examples of w-cHIN contact with MNs, with intersection area (yellow) shown through the neuron meshes. **E.** Lateralized pathway from DNa04 to steering MNs through w-cHINs and associated cells as a candidate steering microcircuit, and predicted effect on lateralized control of steering MNs. For clarity, the right side DNa04, IN18B041, IN06A004, IN03B066 group 19226, IN06A044 group 16214, and IN06A019 group 13094 are omitted. While neurotransmitter prediction for several INs are below probability threshold (<0.7; round edge arrowheads), hemilineage 18B is thought to be cholinergic, while 3B, 6A and 19A are thought to be GABAergic (Lacin et al., 2019). However, IN03B066 group 19226, IN06A002, IN06A004, and IN06A019 group 13094 have moderate prediction probability (≥0.5) for glutamate, which may be excitatory or inhibitory. An inhibitory role of these cells is more consistent with a role of DNa04 in mediating asymmetric muscle control (E, bottom panel). Side annotation is neck side for DNs, output nerve side for MNs, and soma side for others. Synapse weight (neuron-to-neuron mean) is thresholded at ≥50, except for connectivity from w-cHINs to MNs. **F.** Convergence of DN and sensory input (including ascending sensory) onto w-cHINs. All DNs contributing ≥1% groupwise input to a w-cHIN group are shown as a single node, while sensory neurons grouped by anatomical origin contributing ≥1% input to w-cHIN groups are shown. **G.** Morphology of w-cHINs (top, colored by group), pathway of DNa04 to contralateral b1 and b3 MNs through w-cHIN group 10073, IN18B041 and IN19A026 group 10764 (middle), and convergence of SNs (including ascending sensory) and DNs onto w-cHINs (bottom). **H.** Direct lateralized DN input to steering MNs, haltere MNs (from **A**) and w-cHINs, for DNs providing groupwise input of ≥1% to any. Ipsilateral-contralateral index is calculated by ipsilateral minus contralateral connectivity, normalized by their sum. w-cHIN side is considered to be their output side, contralateral to dendrites. DN labels colored red are not within the WTct/HTct DN set defined in Figure 10.

**Figure 11—Supplement 1:**
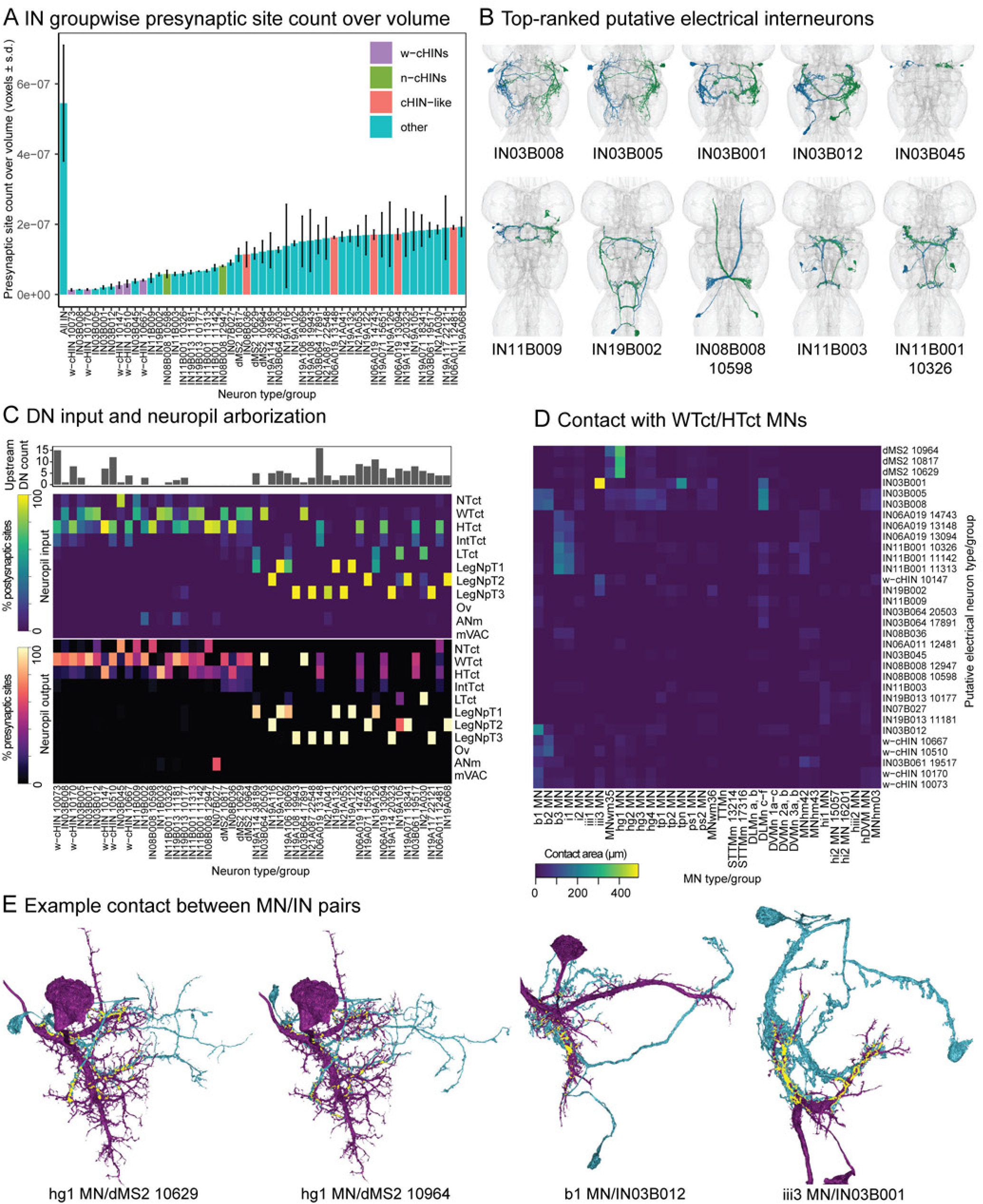
Putative electrical interneurons in the Upper Tectulum. **A.** Top 50 intrinsic neuron groups in MANC rank-ordered low to high by presynaptic site count per neuron volume (groupwise mean), which is suggestive of non-chemical synaptic transmission such as electrical synaptic connectivity. The mean of all VNC intrinsic neurons is also shown. Within the top 50, groups with dense core vesicles (which suggests extrasynaptic neurotransmitter release) or significant reconstruction issues were manually evaluated and excluded (see Supplementary file 5). Neurons are named by type, with their group appended if multiple groups share a type. **B.** Morphology of top 10 rank-ordered putative electrical interneuron groups besides the w-cHINs shown in Figure 11. **C.** Direct upstream DN count (thresholded at ≥1% groupwise input), and neuropil input and output of the top 50 putative electrical interneuron groups. **D.** Mean surface contact area between single putative electrical interneurons of a group and all wing and haltere MNs of a group. Contact area was determined by expanding each interneuron and MN by 0.1 μm and calculating the surface area of intersection meshes between every interneuron-MN pair. **E.** Example interneuron-MN pairs with high contact area as determined in **D**. Intersection area is shown in yellow on top of the neurons. All views shown are ventral.

**Figure 11—Supplement 2:**
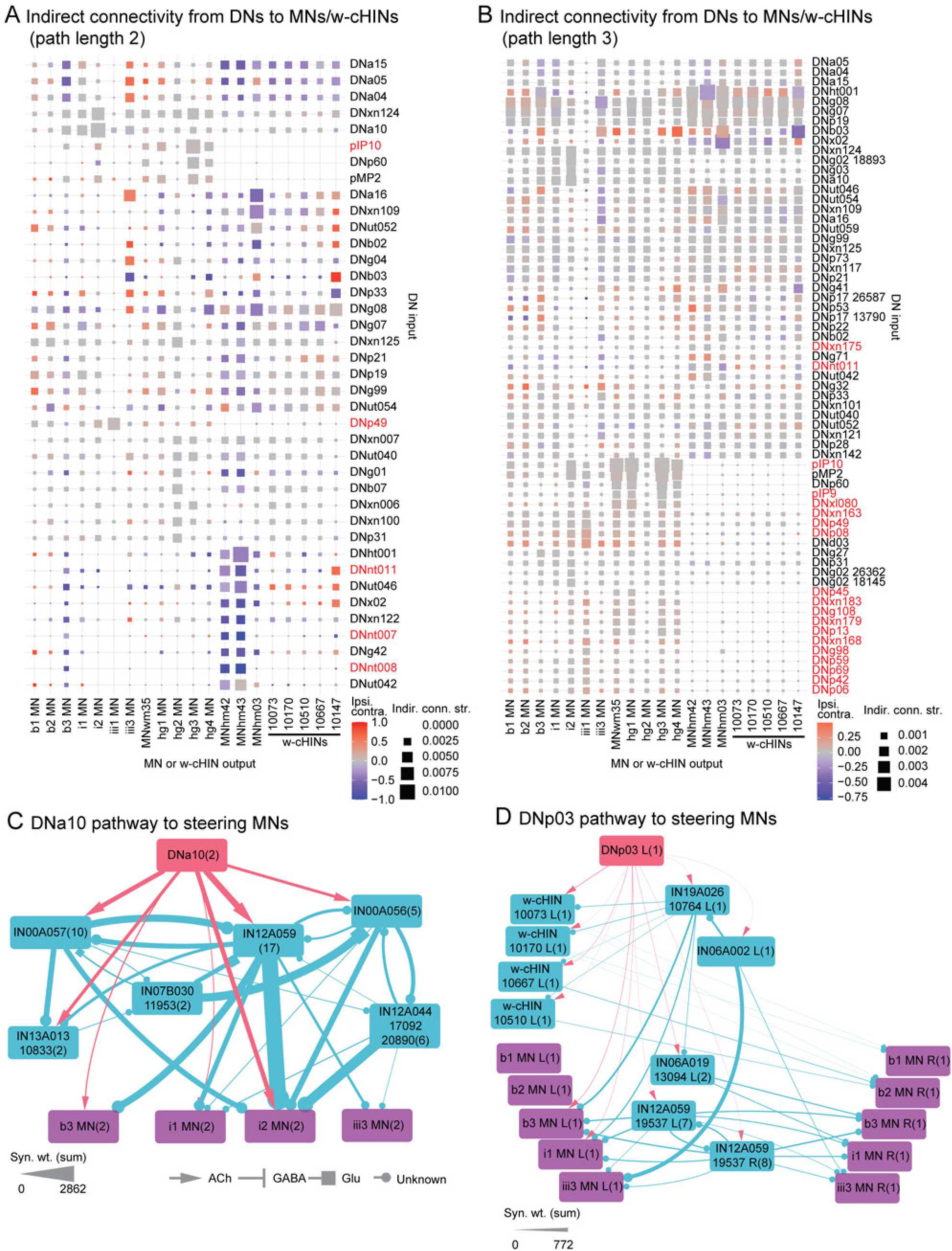
Additional connectivity of DNs controlling steering MNs. **A-B.** Lateralized indirect connectivity of DNs to steering MNs, haltere MNs and w-cHINs, for path length of 2 (A) and 3 (B). Un-normalized indirect connectivity strength is used, as a comparison between path lengths is not carried out. Ipsilateral-contralateral index is calculated by ipsilateral minus contralateral connectivity strengths, divided by their sum. For this calculation, w-cHIN side is considered to be their output side, contralateral to dendrites. Only DNs above a cutoff value for indirect connectivity strength (≥0.003 for path length 2, ≥0.0009 for path length 3) with MNs/w-cHINs are shown. DN labels colored red are not within the WTct/HTct DN set defined in Figure 10A. **C.** Highly bilateral pathway of DNa10 to steering MNs. Synapse weight (group sum) is thresholded at ≥100. **D.** Lateralized path of DNp03 to steering MNs through w-cHINs and other cells. For clarity, only the left DNp03 and INs (by dendrites side) are shown, except for IN12A059 group 19537, a bilateral population cell type. Synapse weight (group sum) is thresholded at ≥20. Side annotations in node names are neck input side for DNs, output nerve side for MNs, and soma side for others.

**Figure 11—Supplement 3:**
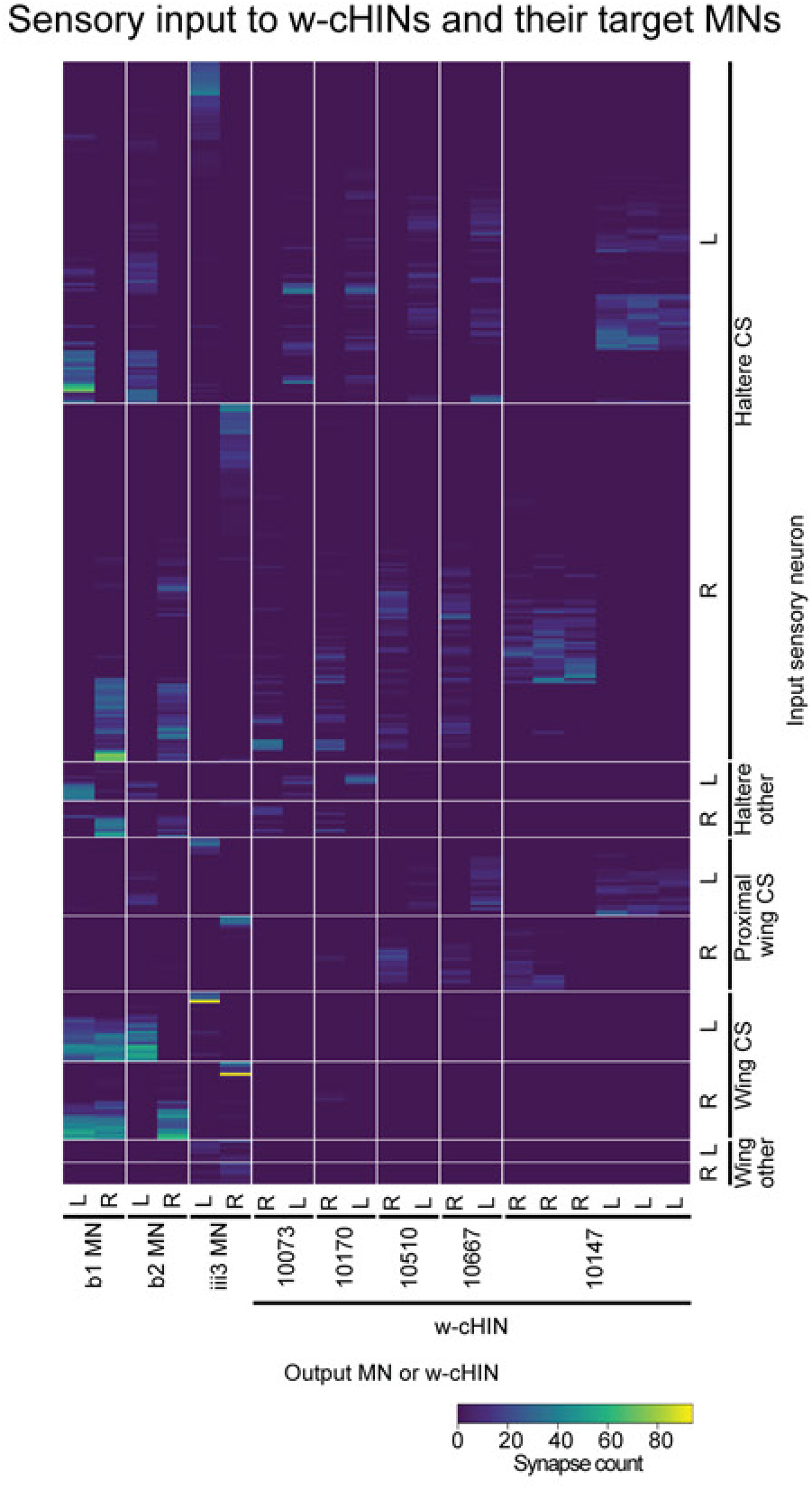
Sensory input to w-cHINs and their target MNs. Adjacency matrix of synapse weights of individual sensory neurons (including sensory ascending) to individual w-cHINs or w-cHIN target MNs. Only sensory neurons which contribute 1 or more synapses to individual w-cHINs or MNs are shown. X-axis labels for sensory neurons denote their literature type (if known) as annotated in the synonyms NeuPrint field. For sensory neurons without a synonym annotation, their entry nerve field was used to denote presumptive identity: ‘wing other’ neurons enter through the anterior or posterior dorsal mesothoracic nerve, and ‘haltere other’ enter through the dorsal metathoracic nerve. Side annotations are entry nerve side for sensory neurons, exit nerve side for MNs, and soma side for w-cHINs (same side as their dendrites, contralateral to axon terminals).

To examine MN control by these DNs, we first plotted their most direct pathways to wing and haltere MNs (Figure 11B). Indeed, multiple steering MNs are found downstream of these DNs, notably b1, b2, b3, i1 and iii3, as well as the hb1 and hb2 haltere MNs (MNhm42 and 43; exact muscle assignment unknown), which are the haltere homologs of the basalar wing MNs. Despite similar morphology in the VNC, notable differences are observed in connectivity between these DNs–they may roughly be split into two sets, namely one set (DNa04, DNa05, DNut037, DNut038, DNut060) with direct connectivity to wing contralateral haltere interneurons (w-cHINs, morphology recognized from (Trimarchi and Murphey, 1997)), and one set (DNa09, DNxn123) with no connectivity to w-cHINs but that have strong connectivity to haltere MNs, which may indirectly affect steering (Dickerson et al., 2019).

The w-cHINs are excellent candidates as intermediaries for DNs to carry out steering during flight, as they have been documented to contribute lateralized electrical input to at least the b1 MN (Trimarchi and Murphey, 1997). Thus, we further looked into their connectivity and possible functions in wing motor control. In MANC, we observe five groups of large w-cHINs consisting of 14 cell pairs, all of which receive substantial DN input (also see Figure 11H). Visual examination in the EM volume indicates physical contact between the w-cHINs and several MNs, suggesting that the five w-cHIN groups may each provide electrical synaptic input to distinct sets of MNs. Electrical synapses are too small to be resolved using our current EM method. However, where they occur the cell membranes of the two cells they connect run in close parallel contact to each other. Thus to unbiasedly evaluate physical contact of w-cHINs with wing MNs to form hypotheses for putative electrical connections, we retrieved EM-derived 3D models of the w-cHINs and all wing MNs and quantified the area of contact between w-cHINs and steering MNs (Figure 11C, D). This method suggests that two w-cHIN groups have electrical connectivity with the b1 MN, two groups have electrical connectivity with both the b1 and b2 MNs, and one group have electrical connectivity with the iii3 MN. In all cases, the w-cHINs connect to contralateral steering MNs (relative to w-cHIN dendrites), insert at the base of the MN dendritic tree, and are thus highly suited to drive steering MN activity while bypassing the MN dendritic integration process (Figure 11D).

How might DNs use w-cHINs and their synaptic partners for steering? To examine this, we looked at the example of DNa04. We plotted the ipsilateral and contralateral pathways of DNa04 to steering MNs through w-cHINs and other cells (Figure 11E, also see Figure 1G). Upon activation of a DNa04 cell on one side, we would expect that w-cHINs are directly activated and indirectly disinhibited to mediate a contralateral increase in activity in the b1, and iii3 MNs. The contralateral b1 and b2 MNs may also be activated through IN18B041, while the contralateral iii3 MN may be further disinhibited through a two-hop synaptic connection. At the same time, we expect an ipsilateral increase in b3 and i1 MN activity through direct DNa04 input, as well as disinhibition via a two-synaptic hop connection. Separately, the contralateral b3 and i1 MNs are expected to be inhibited through one- and two-hop connections. Note that this circuit interpretation is dependent on several neurotransmitter predictions below threshold (<0.7) and the interpretation that Glu usage is inhibitory, which must be validated in future work. All in all, the pattern of ipsilateral-contralateral MN activation suggests an effect of turning towards the ipsilateral side of DNa04 activation, although not all steering MNs associated with turning are found in this pathway (notably hg MNs do not receive substantial input from DNa04, at least at shorter path lengths). Thus, DNa04 is indeed likely to play a steering role during flight.

In investigating the connectivity of w-cHINs, we also observed substantial chemical synaptic input to all w-cHINs from haltere campaniform sensilla as well as lesser input from proximal wing campaniform sensilla (Marin et al., 2024), corroborating connectivity previously identified in the literature (Dickerson, 2020; Trimarchi and Murphey, 1997) (Figure 11F). Within the wing and haltere sensory neurons, distinct subsets provide input to w-cHIN groups and their target MNs (Figure 11–Supplement 3). During flight, haltere campaniform sensilla send oscillatory signals that encode the inertial forces acting on the haltere as it moves in antiphase to the wing, and in separate sensory neuron subsets, signals encoding angular changes to the fly from the action of Coriolis forces on the haltere (Heide, 1983; Pringle and Gray, 1948). Oscillatory input from haltere campaniform sensilla is critical for determining the phase-locked timing of SM firing relative to the wingbeat cycle (Fayyazuddin and Dickinson, 1999, 1996; Trimarchi and Murphey, 1997), as the asynchronously-contracting nature of PMs does not allow the fly to determine its phase in the wingbeat cycle from power MN circuit activity alone. The observed convergence of DN and sensory input onto w-cHINs and steering MNs thus suggests a relatively direct method of controlling steering: in tonically active muscles such as b1 and iii3, oscillatory haltere sensory input is the primary driver of wing MN firing, and subthreshold DN input to w-cHINs serves to alter the timing of MN firing in the wingbeat cycle, either causing phase advance from excitatory DN input or phase delay from inhibitory DN input. For phasic MNs such as b2, haltere input to w-cHINs or w-chIN input to the b2 MN is likely subthreshold, and the combined activity of descending and haltere input allows the driving of the MN to threshold while maintaining accurate timing relative to the wingbeat cycle. DN and sensory connectivity is thus consistent with prior observations of phase tuning to wingbeat as well as phase changes during flight maneuvers in the basalar muscles and other SMs (Egelhaaf, 1989; Götz, 1983; Heide, 1983, 1975; Heide and Götz, 1996). Given that the b1 and b2 muscles both exert similar effects on the wing in isolation, i.e. pulling the basalar apophysis in the anterior direction to move the wing forward (Nachtigall and Wilson, 1967), the shared circuitry through w-cHINs may allow recruitment of variable number of muscles by the same descending steering signals, with smaller magnitude turns only requiring phase shifting of the tonic b1 muscle, while larger magnitude turns further requiring recruitment of the phasic b2 muscle. This wiring also explains the observation that firing of the b2 muscle is almost always concomitant with phase advance of b1 muscle firing, while the reverse is not always true (Tu and Dickinson, 1996).

To further define DNs that may play a role in steering during flight, we examined direct and indirect DN input to steering MNs and w-cHINs by direct input (Figure 11H) and indirect connectivity strength (Figure 11–Supplement 2A, B). As lateralization of steering MN control is key for steering, we also calculated an ipsilateral-contralateral index for both direct and indirect DN-MN connectivity (ipsilateral minus contralateral connectivity divided by their sum). Overall, both direct and indirect DN input to the SMs provide a mix of bilateral and unilateral connectivity that may allow muscle combinatorial control underlying the range of aerial maneuvers available to the fly. DNs that have lateralized SM control such as DNa04 and DNa05 likely control turning maneuvers involving yaw and roll, whereas other DNs, such as DNa10 (which also receives input from navigation-related brain areas), provide bilateral input to SMs (Figure 11H, Figure 11–Supplement 2A-C), and thus may be involved in wing maneuvers involving bilateral muscle control, such as in changes of thrust, lift, or pitch. The w-cHINs are sites of convergence of direct, highly lateralized input from many DNs, further indicating their likely importance in steering during flight, although they are not the only lateralized pathways from DN to steering MNs.

Interestingly, while most WTct/HTct DNs mainly arborize in the upper and intermediate tectulum (Figure 10A, C), several also innervate the leg neuropils. Of note, the cholinergic DNa02 is thought to control turning during walking (Rayshubskiy et al., 2020) and has strong connectivity to leg circuits (see section 2.6), yet also has indirect connectivity with the iii3 MN through a w-cHIN (Figure 11H). Why a walking DN has connectivity with a wing MN is unknown, and indeed DNa02 activation in landed flies does not result in wing motion (Cande et al., 2018). It is possible that DNa02’s role in flight is gated by haltere sensory input required to drive the w-cHINs and the iii3 MN, allowing this DN to perform a dual role in steering during walking and flight depending on the behavioral state of the fly–the highly specific connectivity with only the iii3 MN suggests a much more restricted flight role that is perhaps limited to only slow turning, as activation of the iii3 muscle is predictive of slow but not fast increases in wing amplitude (Lindsay et al., 2017).

Visual observation of the EM volume suggests that, in addition to the w-cHINs (and the related neck-cHINs), there are additional groups of putatively electrical neurons that also have large axon diameter and few chemical synapses, including many in the upper tectulum. Trained annotators labeled these neurons ‘putative electrical’ in the neuPrint ‘transmission’ field, while known electrical neurons from the literature are labeled ‘electrical’ when identifiable ((Marin et al., 2024), this work). To further systematically discover putative electrical INs, we rank-ordered all IN groups of the VNC by mean presynaptic site count over neuron volume (low to high) to find neurons which may carry out neurotransmission through non-chemical synaptic means (Figure 11–Supplement 1A, B). These top neuron groups were manually examined to rule out alternative explanations for low chemical synapse count including significant reconstruction issues or the presence of dense core vesicles (Figure 11–Supplement 1A, B, Supplementary file 5). The top-ranking groups largely ramify the upper tectulum, with some ramifying leg neuropils (Figure 11–Supplement 1C). In addition, many are direct targets of DNs, suggesting that some may play roles in wing control similar to w-cHINs (Figure 11–Supplement 1C). To evaluate if any of these cells play a role in electrical wing or haltere motor control, we calculated the physical contact area with wing and haltere MNs for neuron groups within the top 50 that also ramify in tectular regions (Figure 11–Supplement 1D, E). Indeed, many putative electrical cells also come into contact with wing MNs, chiefly steering MNs such as the b1, b2, b3, i1, iii3, and hg1 MNs. Together, these observations suggest that electrical connectivity is a widespread mechanism of wing and upper tectular motor control, and many are part of pathways linking DNs to MNs. Indeed, as the high frequency of wing beats during flight (∼200 Hz in *Drosophila*) demands rapid and repeated activation of many wing MNs, electrical synapses are well-suited for control of wing MNs without the synaptic fatigue issues associated with chemical neurotransmission. However, as electrical synapses cannot be directly observed at the MANC EM resolution, these putative electrical neurons await experimental confirmation.

Overall, descending and sensory input converges onto w-cHINs, making them highly suited to carry out a role in steering through subsequent electrical input to the b1, b2 and iii3 MNs. Descending input such as those from DNa04 and DNa05 to w-cHINs, combined with oscillatory sensory input from halteres, likely allows a flight maneuver signal to be converted into accurately timed muscle activations relative to the wingbeat cycle. Besides the w-cHINs, other putative electrical INs are prevalent in the upper tectulum and perhaps serve similar roles in wing motor control.

#### 2.5.3 Wing power output is likely driven by diffuse excitatory connectivity

In flies, power generation for the high frequency wingbeats necessary for flight are driven by action of the PMs–namely the dorsal lateral muscles (DLMs) and dorsal ventral muscles (DVMs), two sets of asynchronous, stretch-activated muscles which function antagonistically to each other (also see Figure 6A). In this system, the power MNs do not have direct control over PM spike timing, and thus do not directly time wingbeats. Instead, power MN spiking (∼5Hz) is over an order of magnitude lower than the *Drosophila* wingbeat frequency (∼200Hz, Figure 12A; (Gordon and Dickinson, 2006)), and is thought to play a permissive role to enable PM contraction, as well as control calcium availability in PMs to modulate their power output (Gordon and Dickinson, 2006; Lehmann et al., 2013; Wang et al., 2011).

**Figure 12:**
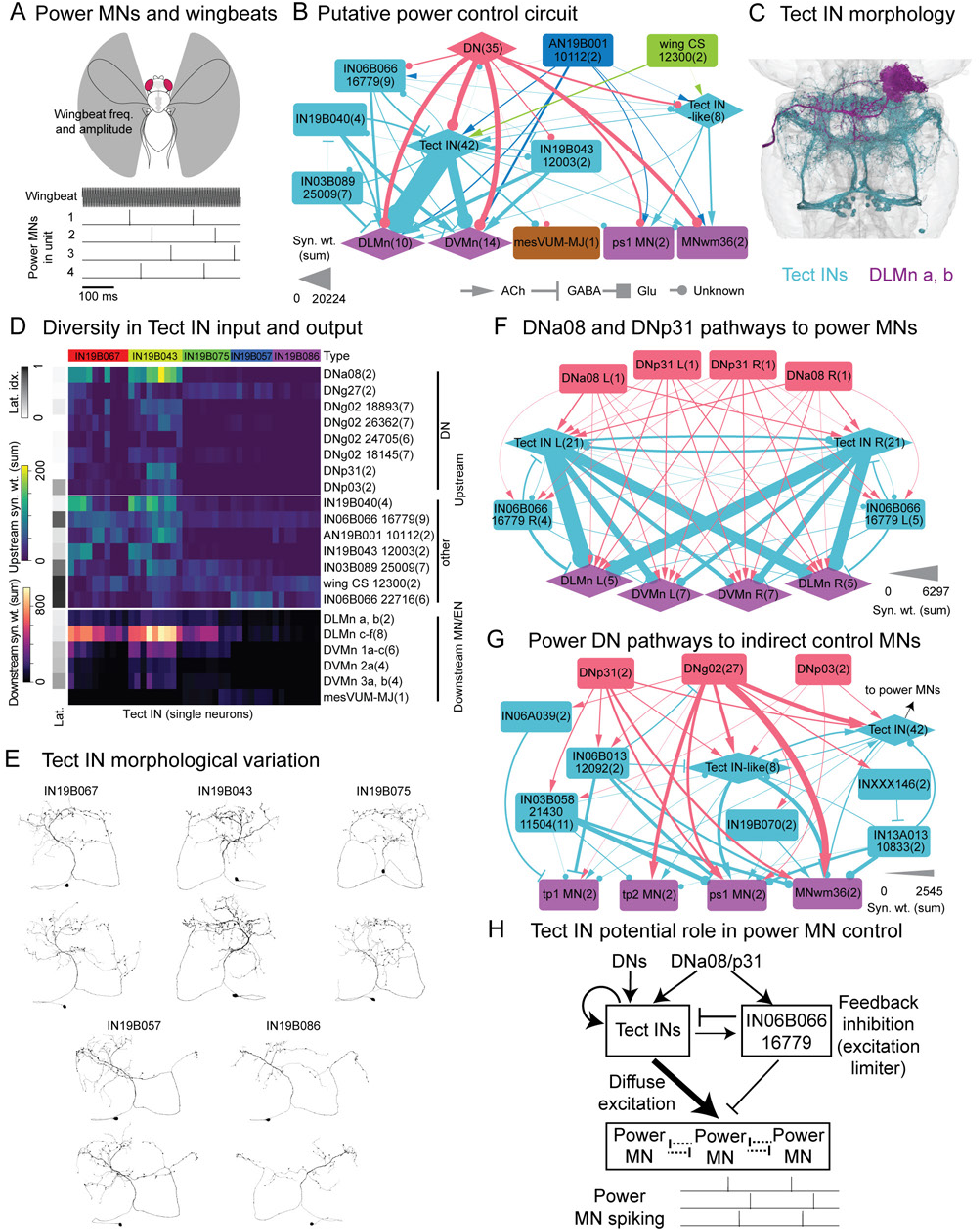
Putative power muscle control circuit for flight. **A.** The power muscles (PM) drive wingbeats during flight in Drosophila. These wingbeats occur over an order of magnitude faster than firing of power MNs, which are thought to enable muscle firing and control muscle power output. The power MNs within the same motor unit fire in a loose, staggered pattern, likely to smooth out PM calcium changes and wing power output. Wingbeat and power MN traces artistically rendered from prior descriptions (Harcombe and Wyman, 1978, 1977; Tanouye and Wyman, 1981; Wyman, 1966). **B.** Upstream connectivity of the power MNs (dorsal lateral, DLMns and dorsal ventral, DVMns). The bulk of upstream connectivity of the power MNs is contributed by 42 tectular intrinsic neurons (Tect INs), with many DNs (here subsetted by ≥1% groupwise connectivity with Tect IN groups) and other neuron types with connectivity to Tect INs. Upstream connectivity to the power MNs are also associated with inputs to the ps1 and MNwm36 wing MNs, and the mesVUM-MJ which targets the dorsal lateral muscles. Edges are thresholded at ≥100 synapses. The Tect INs comprise neuron groups 15113, 13060, 14502, 18519, 21307, 15337, 22521, 21381, 22289, 15788, 14625. The Tect IN-like neurons comprise groups 13874, 18143, 22530, 18638. **C.** Morphology of the Tect IN population, shown with a single DLMn. **D.** Connectivity of the Tect INs (individual neurons on x-axis) with upstream DNs and other major inputs, and downstream MNs/efferents (combined by group). Laterality index is the mean of the difference between side-separated connectivity, normalized by per-side cell count and sum of side-separated connectivity. The Tect INs show variation in the combinations and strength of their upstream and downstream connectivity, which is reflected in their morphological variation (see **E**). Neuron groups shown on y-axis are top input or output neuron types/groups connected to Tect INs, and named by type, with group (5- or 6-digit number) appended if more than 1 group share a type. **E.** Examples of morphological variation in Tect INs. **F.** Lateralized connectivity of DNa08 and DNp31 with Tect INs, IN06B066 group 16779 and power MNs, forming a motif where the same DNs appear to excite and inhibit power MNs. Side annotation is neck side for DNs, nerve side for MNs, and soma side for others. **G.** Connectivity of three example power MN-associated DNs (DNp31, DNg02 and DNp03) with indirect control MNs. **H.** Schematic of putative power MN control circuit. In flight, DN input controls Tect IN activity, which excites power MNs through ‘diffuse’ connectivity (individually small synaptic connections). DNa08 and DNp31 further excites IN06B066 group 16779, which may form an inhibition-stabilized with Tect INs to limit runaway excitation. Power MNs sum ‘diffuse’ excitation from Tect INs to individually reach their spiking threshold, and weak electrical connectivity between power MNs further enforce an inhibition-like connectivity between them, such that power MNs spike in a slow, staggered manner.

**Figure 12–Supplement 1:**
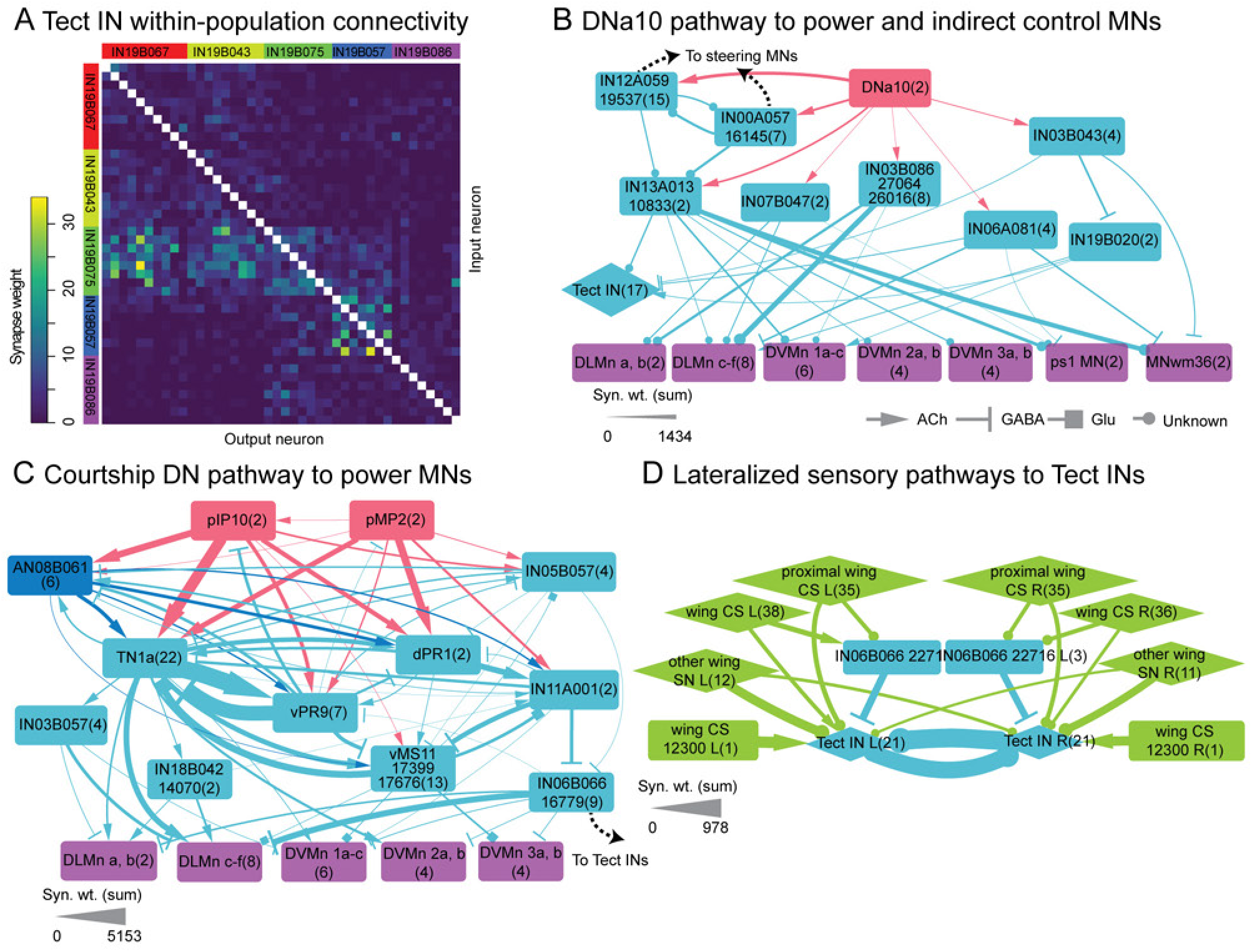
Additional connectivity from DNs to power MNs. **A.** Tect IN connectivity within the population. Tect INs provide weak recurrent input to each other. Order of neurons on both axes is the same as for x-axis in Figure 12D. **B.** Pathways of DNa10 connectivity to power and indirect control MNs. IN00A057 group 16145 and IN12A059 group 19537 also have connectivity to steering MNs, see Figure 11–Supplement 1. Tect IN connectivity to power MNs not shown for clarity. **C.** Pathways of the courtship DNs pIP10 and pMP2 to power MNs. IN06B066 group 16779 has connectivity with Tect INs (see Figure 12F). Identification of the TN1a, dPR1, vPR9 and vMS11 cell types implicated in courtship in MANC by (Lillvis et al., 2024). **D.** Lateralized connectivity of wing campaniform sensilla (CS) and other wing sensory neurons (including sensory ascending) to Tect INs and IN06B066 group 22716. Node labels for the sensory neurons denote their literature type as annotated in the synonyms NeuPrint field, and presumed wing sensory neurons without synonym annotation that enter through the anterior dorsal mesothoracic nerve are placed in the ‘other wing SN’ nodes. The wing CS group 12300 is shown separately due to its exceptionally strong connectivity with Tect INs. Node labels for other neuron classes for all graphs are neuron type, appended with five- or 6-digit numbers for neuron groups (if a subset of groups within a type are shown), while number in parentheses indicates cell count per node. Synapse weights of all graphs are thresholded at ≥100.

To begin examining power MN function in MANC in relation to descending control, we examined the upstream connectivity of power MNs, particularly in relation to connectivity from DNs of interest that are directly or indirectly upstream of power MNs such as DNa08, DNa10, DNp31, DNg02 and DNp03. The upstream connectivity of power MNs show strong similarity, suggesting that power MNs are largely controlled together (see Figure 6E). Strikingly, when considering population-wise input, all power MNs receive heavy shared input from a population of ∼21 pairs of wing upper tectular intrinsic neurons (‘Tect IN’, Figure 12B-D) that were categorized into 5 types and 11 groups by variations in their morphology and connectivity. While each individual Tect IN contributes relatively low input to power MNs (mean of 22 synapses per Tect IN with each power MN), together they comprise 7-23% of total input to each power MN. These cells also have input or output with many other top contributors of input to the power MNs, and receive input from most DNs that are upstream of power MNs, at least at shorter path lengths (Figure 12B). In addition, the Tect INs target the mesVUM-MJ/T2VUM1, an efferent neuron that likely provides modulatory octopaminergic input to the DLMs (Ehrhardt et al., 2023; Schlurmann and Hausen, 2003). DNs and other cells associated with Tect INs commonly have input to the ps1 MN and the likely ps-related MNwm36, suggesting that control of thorax rigidity/tension by the ps muscles is intimately tied to control of wing power output. The Tect IN pathway is one of three routes from DNs to power MNs. Two other pathways are taken to power MNs by the courtship song DNs (pIP10 and pMP2) and the putative navigation-related DNa10, respectively–these routes also have some indirect connectivity with the Tect IN pathway (Figure 12–Supplement 1B, C).

Prior work has observed that the power MNs exhibit slow out-of-phase spiking during flight, with a loose order enforced on the MNs of the fibers of each PM motor unit (Harcombe and Wyman, 1978, 1977; Tanouye and Wyman, 1981; Wyman, 1966), which is a hypothesized mechanism to convert a discrete flight signal, such as from DNs, into more smoothly graded power changes. The out-of-phase firing of power MNs led experimenters (Harcombe and Wyman, 1977) to suggest that each power MN may summate many small excitatory postsynaptic potentials from a common source over time to independently reach their spiking threshold, as opposed to single strong excitation sources which would lead to synchronous spiking. The large population of Tect INs is an excellent candidate to carry out such a function–they are likely cholinergic (excitatory) by their 19B hemilineage (Lacin et al., 2019), and their input from DNs may serve to activate ‘diffuse’ activity within the population, which is then contributed to the power MNs via relatively small individual synapse counts. Furthermore, 7% of Tect IN input is contributed by weak intra-population recurrent connectivity (Figure 12–Supplement 1A), likely leading to self-reinforcing activation and signal amplification of DN input. Out-of-phase firing of the power MNs are likely then further promoted by weak electrical connectivity between the power MNs themselves, which through experimental and modeling studies are thought to lead to a limited possible set of DLMn spiking orders (Hürkey et al., 2023). The minimal circuit elements described here may be sufficient to drive the CPG-like control of out-of-phase power MN spiking, as well as provide a means of modulating power MN spike rate through which other neurons or circuits act.

Interestingly, the Tect INs are not a homogenous population. Rather, they include a range of connectivity strengths with their top upstream and downstream partners (Figure 12D) as well as a range of morphologies (Figure 12E). Clustering of neurons by connectivity and morphology separates the Tect INs into 5 types (Marin et al. 2024) with differences in connectivity strength with top upstream and downstream partners of these cells (Figure 12D). DN input to Tect INs is largely concentrated within two Tect IN types (except DNg27 input), and the same types have higher output to power MNs (although all types have power MN output). The other types have higher input from a single wing campaniform sensilla pair (group 12300), as well as IN06B066 group 22716, which receives input from wing SNs (Figure 12–Supplement 1D)—these cells represent a major route of sensory input to Tect INs. In addition, a degree of left-right asymmetry of calcium activity in PMs has been observed during flight and is thought to modulate PM power output to support turning (Lehmann et al., 2013). To examine if lateralization of Tect IN inputs and outputs may play a role in asymmetric power modulation in PMs, we calculated a lateralization score for Tect IN connectivity ranging from fully bilateral (score of zero) versus fully lateralized (score of one, Figure 12D, left bar). Overall, Tect IN connectivity to power MNs are weakly lateralized, suggesting that lateralized Tect IN activation likely generally activates excitatory drive to all power MNs but potentially still causes slightly asymmetrical power changes. However, direct DN input to Tect INs are almost fully bilateral, except for connectivity from DNp03; although its connectivity is weaker than other DNs, this putative looming responsive DN (Namiki et al., 2018) also has connections to w-cHINs and steering MNs (Figure 11–Supplement 1D), which together may support concurrent turning with increase in wing power during flight that perhaps aids in evasive maneuvers. Of other lateralized upstream partners, the sensory paths from the wing SN group 12300 and IN06B066 group 22716 are highly lateralized (Figure 12–Supplement 1D), suggesting that wing sensory input may trigger partially lateralized power MN activity through Tect INs. Lastly, connectivity from IN06B066 group 16779 to Tect INs is also highly lateralized.

Of Tect IN inputs, IN06B066 group 16779 is unusual in that it is likely inhibitory (GABAergic) by its 6B hemilineage (Harris et al., 2015) and neurotransmitter prediction. It has reciprocal connectivity with Tect INs, and is a target of two cholinergic DNs, DNa08 and DNp31, which provide input to Tect INs and are also the strongest direct DN inputs to power MNs (Figure 12F). DNa08 is likely implicated in navigation as it receives input from the lateral accessory lobe and posterior slope in the brain (Namiki et al., 2018), while DNp31 is likely implicated in responses to salient visual stimuli as it receives input from lobula columnar neuron types LC22/LPLC4 and LPLC3 (Namiki et al., 2018). Together, these neurons form an unusual motif where the same descending input appears to both excite and inhibit power MNs (Figure 12F, H). While the actual function of this motif is unknown, one intriguing possibility is that the recurrent excitatory and inhibitory connections in the Tect INs and IN06B066 group 16779 form an inhibition-stabilized network, where a stable level of excitatory activity can be maintained in the excitatory subnetwork with feedback inhibition from the inhibitory subnetwork serving to prevent runaway excitation (Sadeh and Clopath, 2021). Such networks are notable for the paradoxical observation that excitation to the inhibitory neurons decreases the steady-state activity for both the inhibitory and excitatory neuron populations (Sadeh and Clopath, 2021)–the input from DNa08 and DNp31 to the Tect INs and IN06B066 group 16779 may thus serve to shift the steady-state activity of the network by decreasing IN06B066 group 16779 activity and counteracting the corresponding decrease in Tect IN activity. Alternatively, short-duration DN input to both Tect INs and IN06B066 group 16779 may serve to transiently increase Tect IN activity with delayed feedforward inhibition from IN06B066 group 16779, causing momentary changes in wing power output. In addition, several DNs upstream of power MNs such as DNp31, DNg02 and DNp03, (Figure 12G) as well as Tect INs themselves, take pathways to indirect control MNs–namely to the tergopleural and pleurosternal MNs which may control thoracic stiffness or tension. While the roles of the ICMs in flight have not received as much investigation as other muscle categories, it appears that concurrent control of PMs and ICMs are important for control of wing power output by DNs such as DNg02 which modulate wingbeat amplitude (Namiki et al., 2022; Palmer et al., 2022).

In summary, we observe that the power MN upstream network is largely shared among all power MNs and is highly bilateral. The Tect INs provide diffuse excitatory connectivity to the power MNs, potentially facilitating staggered firing of power MNs and control of power MN power output, and is also the main, but not only, target of descending control to the power MNs. Descending input often targets Tect INs and power MNs directly, and two excitatory DNs (DNa08 and DNp31) target both the WTct INs and the potentially inhibitory IN06B066 group 16779 which we hypothesize to form an inhibition-stabilized network with Tect INs (Figure 12G). Thus, wing premotor networks are partitioned into distinct circuits that control power and steering, an organization reflective of the functional specialization of each wing muscle type and also clearly present at the DN level. In addition, the pathways from courtship DNs to wing MNs appear largely distinct from pathways from flight-related DNs. Excitatory and inhibitory connections feature in the circuits upstream of both power and steering MNs, but appear to play different roles, at least in the pathways we explored in detail–these connections are commonly lateralized in steering circuits to allow left-right asymmetrical control of steering MNs, while for the power MN circuit, inhibition appears to function mainly as a means of excitation control. Lastly, circuits controlling indirect control MNs appear mixed with circuits controlling steering or power, suggesting that indirect control muscles may play a supporting role to the function of steering or power muscles.

### 2.6 Organizational logic of DNs onto leg circuits

We next turn our attention to premotor circuits of the leg motor system. Our focus is on circuits that control walking, but of course there are numerous other leg behaviors including grooming or take-off. Rhythmic motor output produces alternating leg movements termed stance (the power phase when legs are in contact with the ground) and swing, which together allow the animal to move efficiently. The movements of all six legs must be coordinated, and flies display a range of walking gaits in which different patterns of inter-limb coordination are observed. Although these motor patterns respond to changing environmental conditions (Büschges et al., 2011), flies have been shown to walk without proprioceptive feedback, suggesting that inter- and intra-leg coordination is not strictly dependent on sensory feedback loops from the legs (Mendes et al., 2013). While headless flies have been shown to generate locomotor patterns, the movements are often uncoordinated, possibly due to the lack of input from DNs (Gao et al., 2013; Mendes et al., 2013).

In the following sections, we carry out an initial analysis of leg locomotor circuits. We first identify a repeated premotor-MN circuit that is consistent across all three pairs of legs; we refer to this as the ‘standard leg connectome’. We then describe the neurons upstream of this circuit that interconnect different leg neuropils; these are likely responsible for inter-leg coupling. Finally, we identify DNs providing input to leg neuropils and examine how they modulate this leg circuit.

#### 2.6.1 The leg premotor circuit is serially repeated across segments

To identify neurons that might contribute to premotor walking circuits, we first examined neurons upstream of the leg MNs (defined in Figure 7). Unlike the upper tectulum motor systems, leg anatomy and its muscular organization is segmentally repeated across the fly’s three thoracic segments; this is also reflected at the level of the leg MNs and their upstream circuits. We found that over 75% of the presynaptic partners of leg MNs are intrinsic neurons restricted to a single leg neuropil (Figure 13A). 52% of these restricted neurons (totalling about 1200 per leg neuropil) have been serially matched across the six neuropils (Figure 13B) (Marin et al., 2024). Our Infomap community detection (Figure 9) also found six very similar communities, each consisting of ∼1200 neurons mostly restricted to single leg neuropils. Together, this suggests that the muscles of each individual leg are controlled and coordinated by local circuits within each hemisegment; inter-leg coordination presumably depends on the much smaller set of neurons with inter-neuropil connectivity.

**Figure 13:**
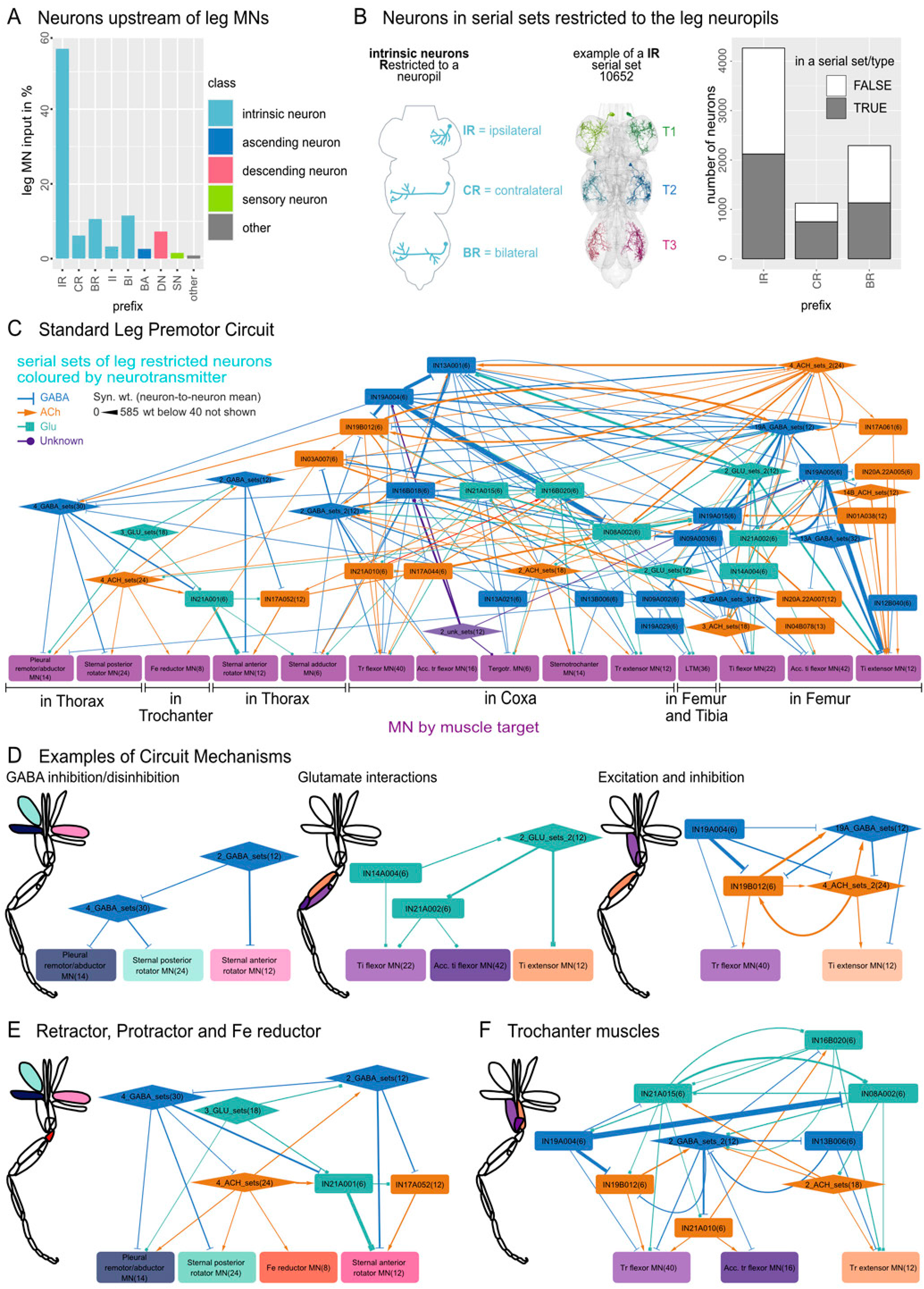
Standard Leg Connectome. Leg premotor circuit local to all three leg neuropils. **A.** Input to leg MNs in percent. Colour indicates the class of the input neurons, and the intrinsic neurons are additionally split up along the x-axis by their subclass (prefix). **B.** Over 75% of input onto leg MNs comes from intrinsic neurons restricted to the leg neuropils. Schematic illustrates the distinction between the three types of restricted neurons: IR, CR, BR. Next to it an example of a IR serial set (10652, type IN08A006) colored by leg neuropil. On the right the number of neurons in the VNC with those prefixes. Color indicates if the neuron is sorted into a serial set or a serial type (serial sets that contain more than 2 neurons per hemisegment). **C.** Premotor circuit of all Leg MNs that are in serial sets. All connections between serial leg restricted neurons and MNs are shown (weight > 40), collapsed by serial set (see Figure 13–Supplement 1 for differences between leg circuits). Color of intrinsic neurons indicates the neurotransmitter prediction. Squares are single serial sets, hexagons are grouped serial sets (See Supplementary file 6 for more information). **D.** Examples taken from **C** to show a GABA, Glutamate and GABA/Cholinergic mechanism of controlling the leg MNs. **E.** Interconnection between the MNs in the Thorax with the Fe reductor. **F.** Major serial sets controlling the trochanter MNs. MN nodes in **D, E** and **F** are colored by the color of the leg muscle they control in the inset leg schematics.

**Figure 13—Supplement 1:**
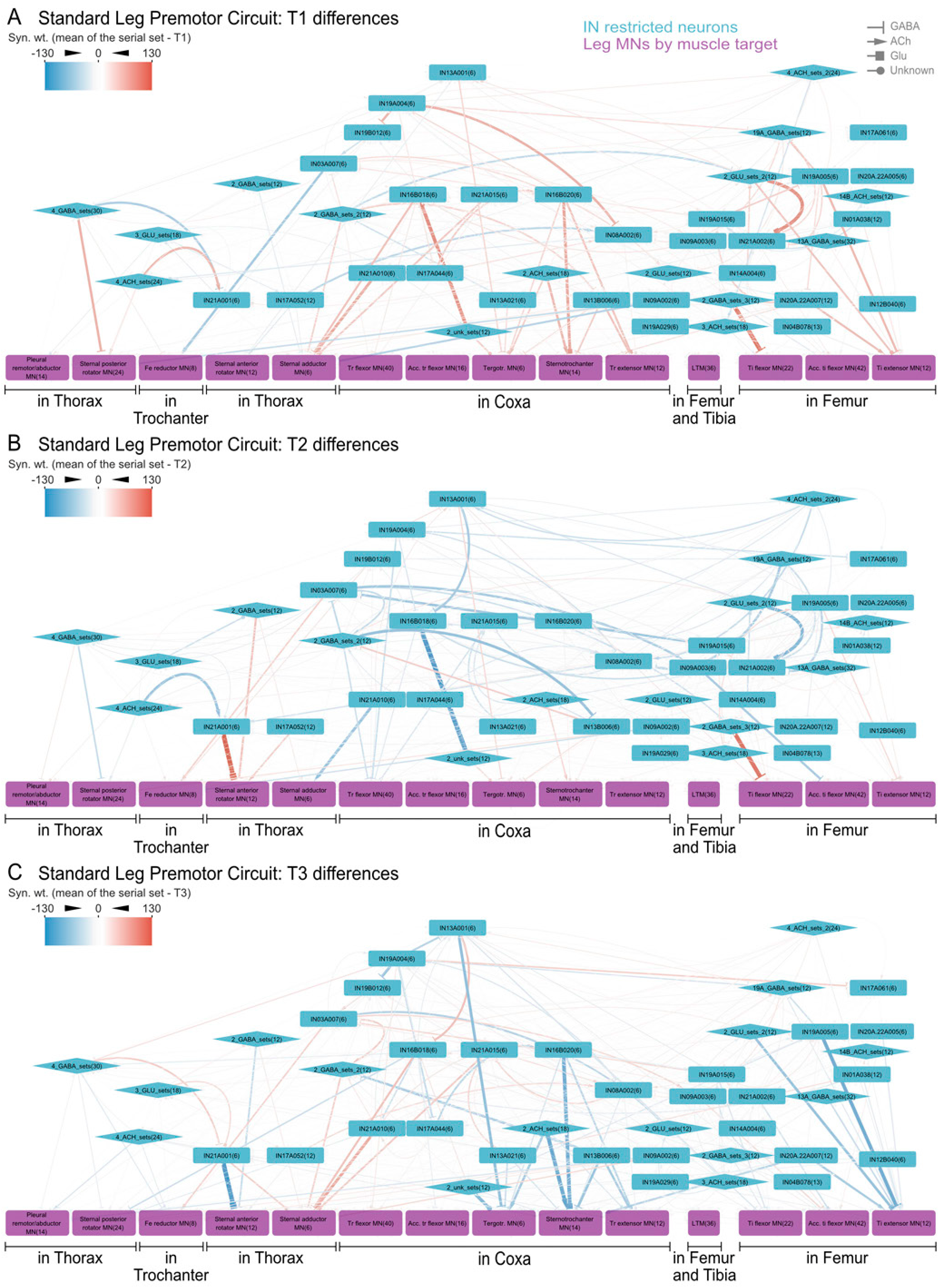
Differences in the Standard Leg Connectome across segments. Difference between the mean connection of the serial set and T1, T2 or T3 specific connections in **A, B, C** accordingly. Weight difference is color-coded, with red indicating stronger connections and blue meaning weaker connections compared to the mean connection of the serial set. Intrinsic serial sets are in light blue, motor neuron serial sets by muscle target in purple. Squares are single serial sets, hexagons are grouped sets.

This modular and repeated organization means that we can use serial homology between leg neuropils to simplify the representation and analysis of leg circuits; this approach also helped us to overcome some inconsistencies in the reconstruction state of leg MNs. The standard leg connectome is therefore defined as the serially repeated units common to all legs including the presynaptic neurons directly upstream of serially identified leg MNs (Figure 13C). For this analysis, we grouped the serial MNs by their leg muscle target as shown in Figure 7 (see Supplementary file 6). The premotor circuit is large and complex even when considering only the strongest connections between presynaptic neurons and serial leg MNs (using a threshold of 40 synapses onto each MN group averaged across each leg) (Figure 13C, Supplementary file 7). The Tarsus MNs could not be matched across segments and are therefore not included in this analysis. This standard leg connectome was very similar across legs, but there were small deviations between T1, T2, and T3 legs, as shown in Figure 13–Supplement 1, some of which may be due to differences in MN reconstruction issues across the three thoracic segments.

Neurotransmitter predictions in the dataset greatly assist in interpreting the circuit logic within this standard leg connectome (Figure 13D). For example, there are five GABAergic restricted neuron sets that strongly inhibit MNs innervating the thorax muscles responsible for the rotation of the leg. Two of these GABAergic IN sets (2_GABA_sets: IN03B042 and IN08A006) inhibits the Sternal anterior rotator MNs that rotate the leg forwards during the swing phase of forwards walking (Azevedo et al., 2024, 2020; Baek and Mann, 2009). The other three GABAergic sets (4_GABA_sets: IN09A002, IN09A010, IN19A022, IN19A016) target the opposing MNs (Pleural remotor/abductor and Sternal posterior rotator MNs), active in the stance phase that propel the body forwards (Figure 15D, left panel). These strong inhibitory input neurons are characteristic of most leg MNs; furthermore, the interneurons also receive strong inhibitory input themselves, suggesting that inhibition and disinhibition are key motifs in leg premotor circuits.

We also find many glutamatergic interactions. In the vertebrate CNS, glutamate is the major excitatory neurotransmitter; in *Drosophila* both excitatory and inhibitory receptors are present but existing evidence suggests that glutamate is principally inhibitory in the CNS (Liu and Wilson, 2013; Marin et al., 2024). There are two glutamatergic INs (grouped 2_GLU_sets: IN08A005, IN08A007) upstream of the Tibia extensor MNs and two upstream of the Tibia flexor MNs (IN14A004, IN21A002), and these two sets also target each other, likely forming a reciprocally inhibitory microcircuit (Figure 13D, central panel).

The GABAergic and glutamatergic circuit motifs that we have just described involve neurons targeting the same jointed segment of each leg; this is typical for the direct inputs to MNs. For example, each MN group has upstream partners that can specifically activate or inactivate that group (Figure 13E, F). However, each step requires coordination between the different segments of the leg. We observe some circuit motifs where INs target MNs that control different leg segments. These include a set of three GABAergic (IN19A004 and grouped 19A_GABA_sets: IN19A007, IN19A011) and five cholinergic (IN19B012 and grouped 4_ACH_sets: IN01A015, IN03A010, IN21A012, IN21A017) IN serial sets that interact upstream of the Trochanter flexor and the Tibia extensor (Figure 13D, right hand panel). The Femur reductor MNs are another exception: they share glutamatergic INs and one GABAergic IN with the Pleural remotor/abductor and Sternal posterior rotator MNs (Figure 13E) suggesting a simultaneous inactivation. However, these neurons do not share the same cholinergic activation. Since the flexor and accessory flexor muscles have a similar basic action on the legs joints, it is not surprising to see that there are many similarities in their connectivity (Figure 13F).

There are additional INs that specifically interact with a single leg premotor circuit (Figure 14–Supplement 1B, see Supplementary file 8). Most of these are again restricted to the given leg neuropil but some project from other areas of the VNC to the T1 or T3 leg circuits. One of these is the previously described wing grooming neurons, wPN1, that project from the Ov to the T3 leg premotor circuit (Zhang and Simpson, 2022) (see example INXXX011, Figure 14–Supplement 1B, D).

**Figure 14:**
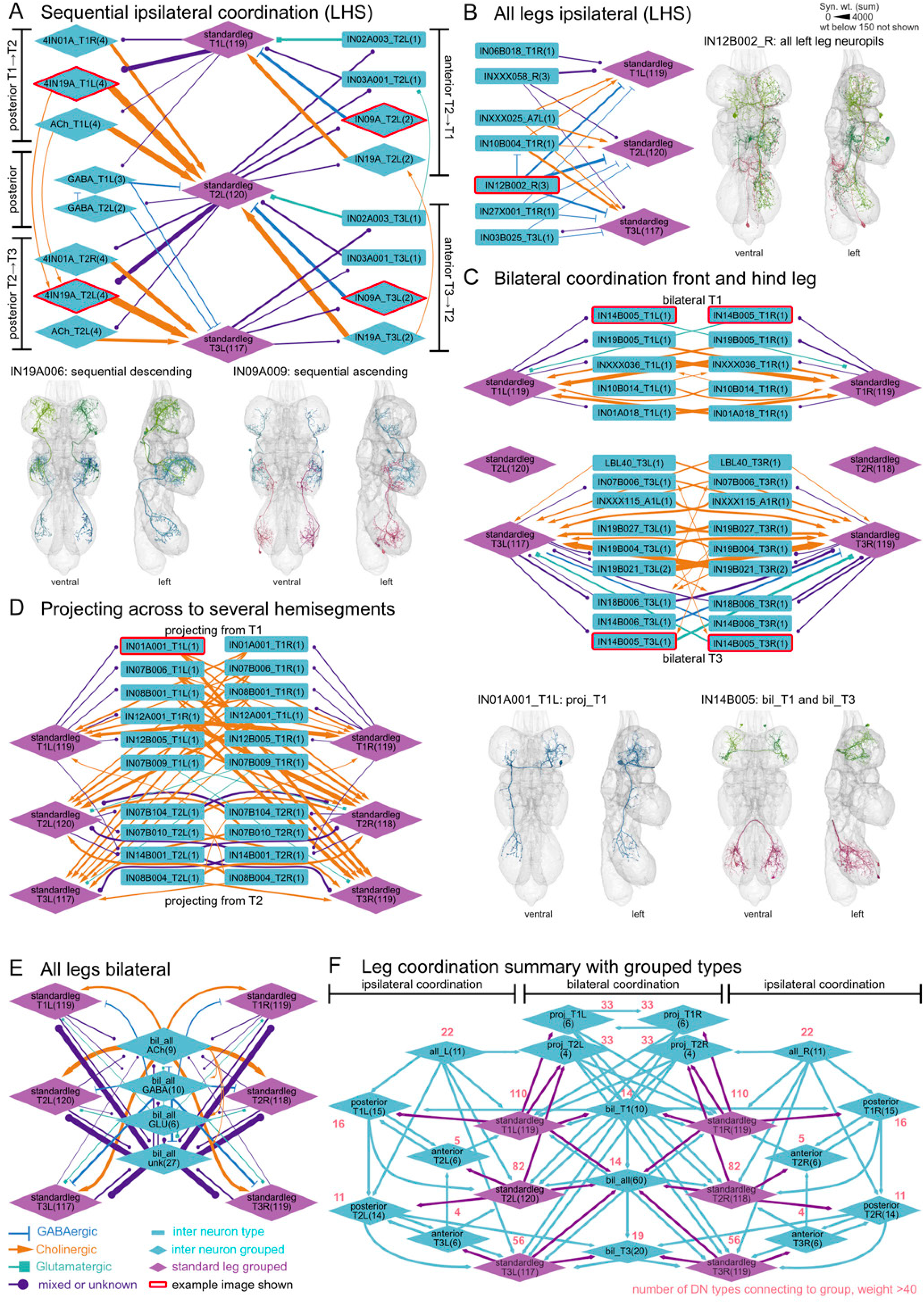
Coordination of leg premotor circuits. Grouped intrinsic neurons upstream of the standard leg premotor circuit with effective connectivity (see Materials & Methods). **A.** Serially repeating intrinsic neurons that ascend or descend a segment with their axonic projections (sequential). Grouped by hemilineage or neurotransmitter, for ungrouped version see Figure 14–Supplement 1. **B.** Ipsilaterally connecting intrinsic neurons, targeting several leg premotor circuits (all_L or all_R). Often projecting from other neuropils, see example IN21B002. **C.** Intrinsic neurons connecting T1 leg or T3 legs bilaterally (bil_T1 and bil_T3), example shown below. **D.** Intrinsic neurons projecting from T1 or T2 on one side to several leg premotor circuits on the other side (proj_T1 or proj_T2). **E.** Intrinsic neurons grouped by neurotransmitter that innervate all leg neuropils on both sides (bil_all). For details and examples, see Supplementary file 1. **F.** Summary of the groups and their connection to the standard leg premotor circuits of the six legs. Numbers indicate the number of DN types that connect to these groups with weight >40. See Supplementary file 7 for neuron types and details.

**Figure 14—Supplement 1:**
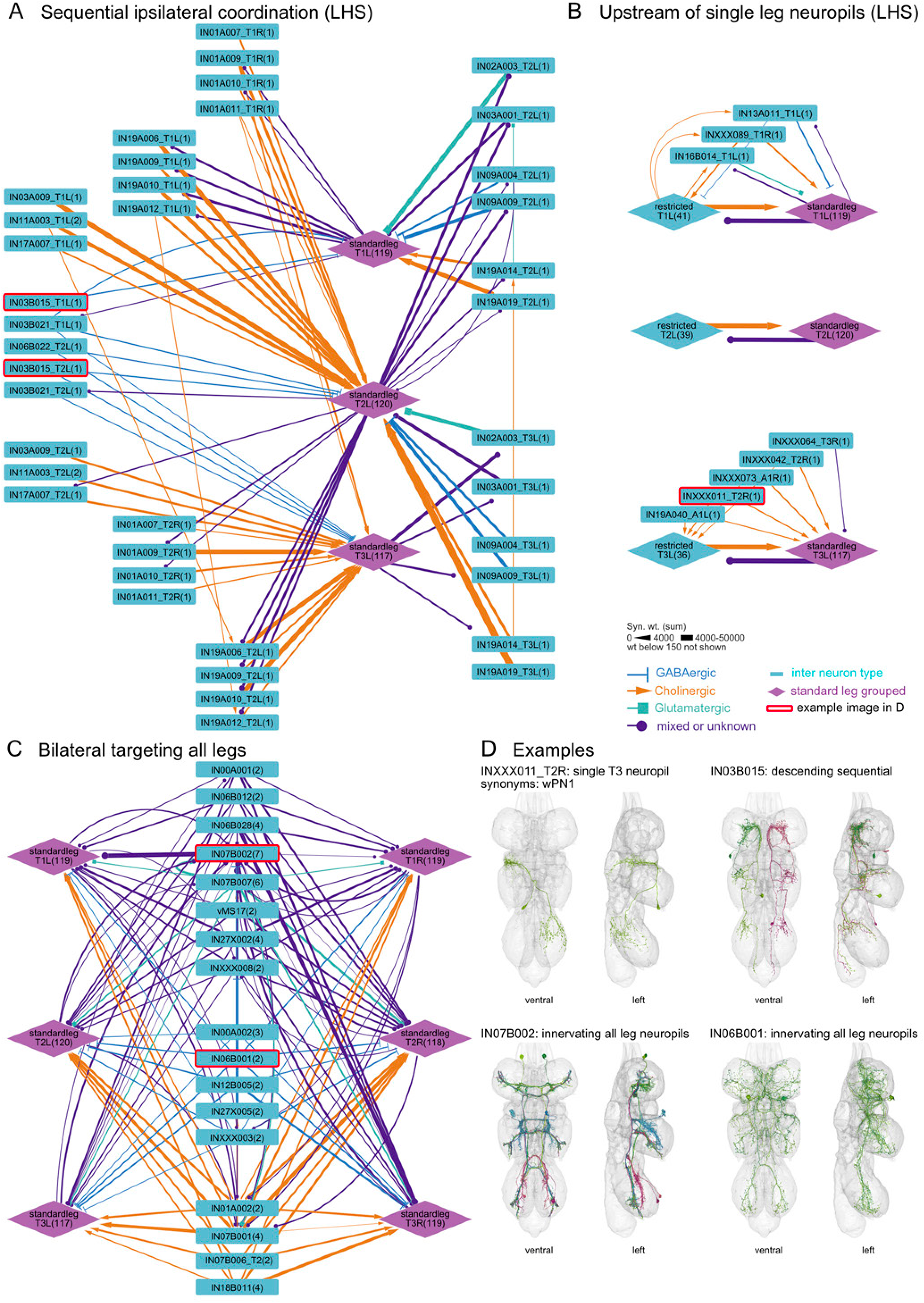
Additional upstream connectivity to leg circuit. **A.** Sequential neurons (see Figure 14A) split by type and hemisegment. **B.** Neurons that project to a single leg neuropil, only the left hand side shown. Most are neurons restricted to the same leg neuropil and the standard leg group (grouped into one node, restricted). A few neurons project from another neuropil to a single leg neuropil. **C.** Neurons that innervate all leg neuropils (see Figure 14E) split by type. **D.** Example images from ventral and side view for neuron types with red stroke around the node in **A, B** or **C.** The first example shows a neuron projecting from the ovoid to the T3 leg neuropil, involved in wing grooming (Zhang and Simpson, 2022).

Our first impressions of the circuits contained within the standard leg connectome can be summarized as follows. First, there is a relatively strong direct inhibitory input onto leg MNs, from over 57% GABA and Glutamatergic serial leg neurons. Layered on top of this are strong inhibitory and excitatory connections from additional leg-specific interneurons, which we expect to be functionally disinhibitory or inhibitory with respect to the MNs. Some inhibitory interneurons show reciprocal inhibitory patterns e.g. to control antagonistic MNs or muscles. However, other interneurons instead show a layered inhibitory organization in which some neurons can control others: this could potentially generate sequential activation of different muscles during the walking cycle. The overall impression is of a high degree of circuit complexity associated within each individual leg neuropil. The extensive inhibitory interactions could easily define circuits with oscillatory activity. Together, this suggests that much of the pattern-generating activity of premotor circuits may operate at the level of a single leg.

#### 2.6.2 Inter-leg coordination is not purely inhibitory

Six-legged walking requires coordination between different legs, both left-right and across thoracic segments. We therefore went on to investigate those neurons that innervate multiple leg neuropils and provide input to the ‘standard leg connectome’ that is repeated for each leg. To do this, we conducted an effective connectivity analysis including both direct and indirect (multi-hop) connections for all the serial MNs and serial INs in the standard leg connectome (see Materials & Methods for details, Supplementary file 8). We grouped the INs by their subclass (see (Marin et al., 2024)). This distinguishes neurons arborizing on one side of the VNC (subclasses II and CI, Figure 14A, B) that likely contribute to ipsilateral leg coordination from INs that may mediate coordination of the two sides (bilateral subclasses BI and BR, Figure 14C, D, E). For ipsilateral coordination, we further distinguished between INs forming two different patterns of neuropil arborization, namely those that were sequentially projecting from a leg neuropil in one thoracic segment to a leg neuropil in an anterior or posterior segment (‘sequential’) and those that ipsilaterally targeted leg neuropils in all three segments (‘ipsi’). We observed that the neurons annotated with the *sequential* serial motif consisted of serial sets of four neurons that connect either the T1 and T2 leg neuropils, or T2 and T3 leg neuropils (Marin et al., 2024). Fourteen sequential groups (and one single pair in T1) receive inputs from a variety of neurons from the standard leg connectome in T1 or T2 and then project their axons posteriorly to the next segment, T2 or T3 (Figure 14A, F, Figure 14–Supplement 1A). Six sequential sets do the opposite, projecting anteriorly from T2 or T3 to T1 or T2 (Figure 14A, F). The morphology and connectivity of these sequential neurons strongly suggests that they coordinate legs across segments. In addition to these sequential neurons, we also found three GABAergic, two cholinergic and two neuron types of unknown neurotransmitter that unilaterally target leg neuropils across all 3 segments equally and are potentially involved in amplifying or attenuating signals to the premotor circuits in response to input from other regions of the VNC (e.g. IN12B002 which receives inputs from different areas of the tectulum in Figure 14B example on the right).

Moving next to bilateral coordination, we expected to see reciprocal connections between the left and right standard leg connectomes for each pair of legs. We observe such strong bilateral connectivity for T1 and T3 leg INs (Figure 14C, see example IN14B005) but such connections do not appear in T2. Coordination between the two front legs or between the two hind legs is thus likely more important for the function of leg premotor circuits. Interestingly, we found a handful of neurons that receive input from the T1 or T2 leg circuit and then project to all leg circuits on the opposite side. We speculate that these might be involved in either turning or providing left-right coordination (Figure 14D, example IN01A001). Finally, we observe a group of neurons that output strongly to all leg neuropils; they are predominantly GABAergic or of unknown (potentially neuromodulatory) neurotransmitter (Figure 14E, Figure 14–Supplement 1C) and could control e.g. leg tone or behaviors besides walking that simultaneously involve all legs.

This analysis of interconnections between the circuits dedicated to each individual leg, gives a framework for how the six legs of the fly might be coordinated (Figure 14F). Many different circuit configurations for a central pattern generator to coordinate walking in *Drosophila* have been proposed, but the neurons that generate this rhythmic pattern of activity have not been identified (Agrawal et al., 2020). Our analysis identifies a relatively small number of strong candidate cell types controlling inter-leg coordination that should be the target of future experiments. However, we observed that the majority of these interconnections are cholinergic, in contrast with previous models which hypothesize that leg locomotor CPGs controlling individual legs are inter-coupled by inhibitory interconnections (DeAngelis et al., 2020; Grillner and Kozlov, 2021; Hiraoka, 2021). However, we propose that this contradiction can be resolved if excitatory interconnections target inhibitory pre-motor neurons (see Discussion and Figure 21B).

#### 2.6.3 Descending neurons provide input to both the intra- and inter-leg circuits

To evaluate how descending input controls leg movements, we used several strategies to identify relevant DNs. The analysis in the previous two sections identified local circuits dedicated to a single leg as well as circuits likely to coordinate different legs. Which elements are targeted by DNs? We find that although some DNs connect strongly to inter-leg circuits, the majority target neurons in the serially repeated standard leg connectomes (Figure 14F). This suggests that the brain controls walking principally at the level of local circuits within individual leg neuropils. We also used a neuropil innervation based analysis to ensure that we examined all DNs likely to have a strong impact on leg movement. In Figure 3D we already identified DNs whose axonal outputs were strongly restricted (>80%) to the leg neuropils. We now used a lower threshold (10% of outputs; >100 synapses) to identify all DN types that innervate leg neuropils and categorized them by the thoracic segment(s) that they innervate (Figure 15). This showed that most leg DNs target either the T1 leg neuropils or leg neuropils for all thoracic segments; just one pair of DNs targets T2 or T3 alone. Furthermore, we observed that direct connectivity between leg-specific DNs (DNfl, DNml, DNhl, DNxl) and leg MNs is rare and usually weak. While for some MNs this may be due to poor segmentation and reconstruction, it seems that most connectivity is indirect through at least one IN. Therefore, we calculated an effective connectivity from each leg DN to all leg MNs (by target muscle); this analysis identifies the strongest multi-hop pathways and the number of layers between them (see Materials & Methods, Figure 15–Supplement 1-4).

**Figure 15:**
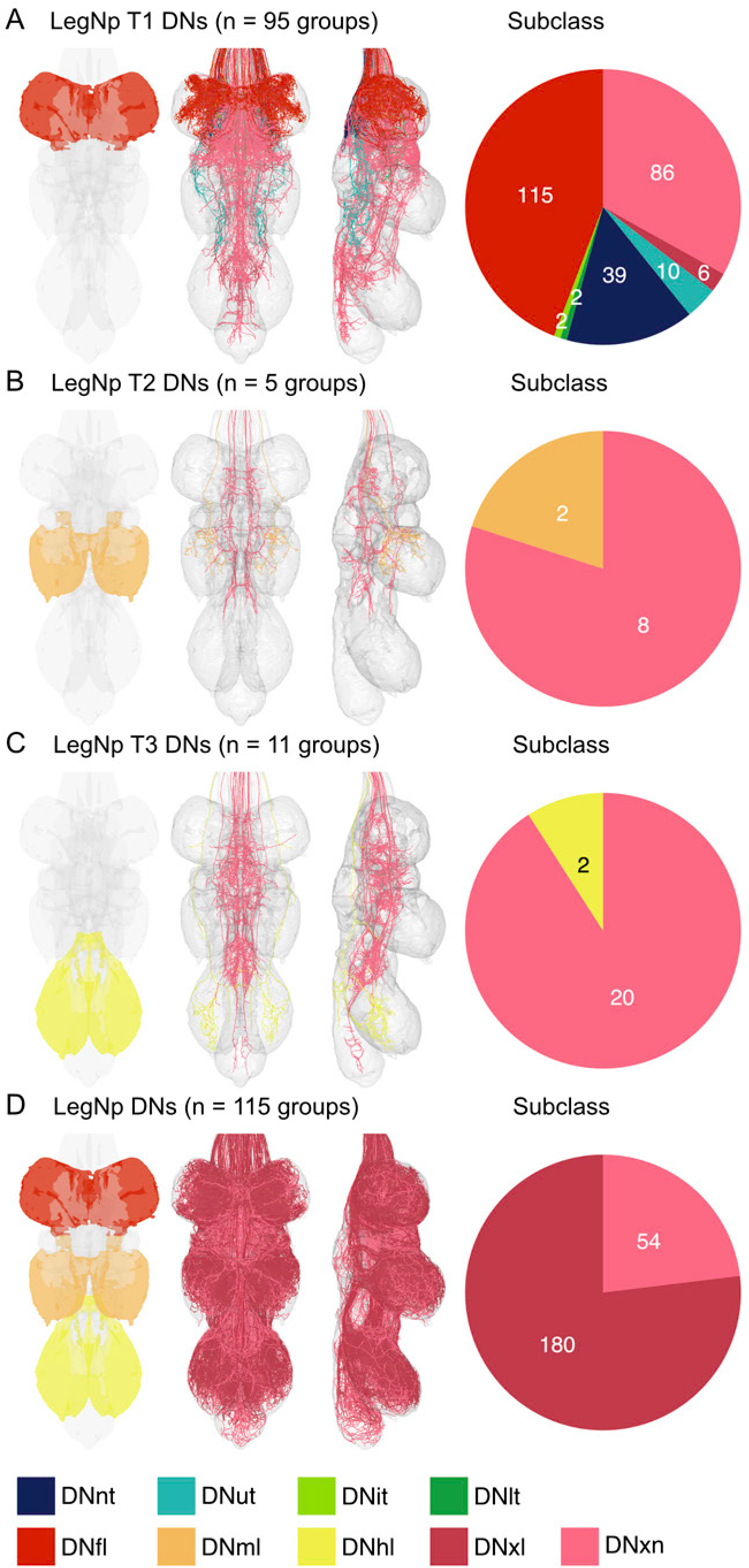
Descending neurons innervating the leg neuropils. DNs that innervate the different leg neuropils with at least 10% of their presynaptic budget and 100 presynaptic sites (group total for both). Their morphology is shown in a ventral and lateral view color coded by their subclass. The number of groups fulfilling the criteria is indicated in brackets and pie charts on the right show subclass (legend at the bottom) composition of the DNs for each innervation type. **A.** DNs innervating LegNpT1 indicated by red area mesh in VNC cartoon on the left. **B.** DNs innervating LegNpT2 indicated by orange area mesh in VNC cartoon on the left. **C.** DNs innervating LegNpT3 indicated by orange area mesh in VNC cartoon on the left. For **A, B** and **C,** DNs that also dedicated more than 10% of their output budget to the respective other LegNps are excluded. **D.** DNs that innervate all three thoracic leg neuropils fulfilling the above criteria for each LegNp.

**Figure 15—Supplement 1:**
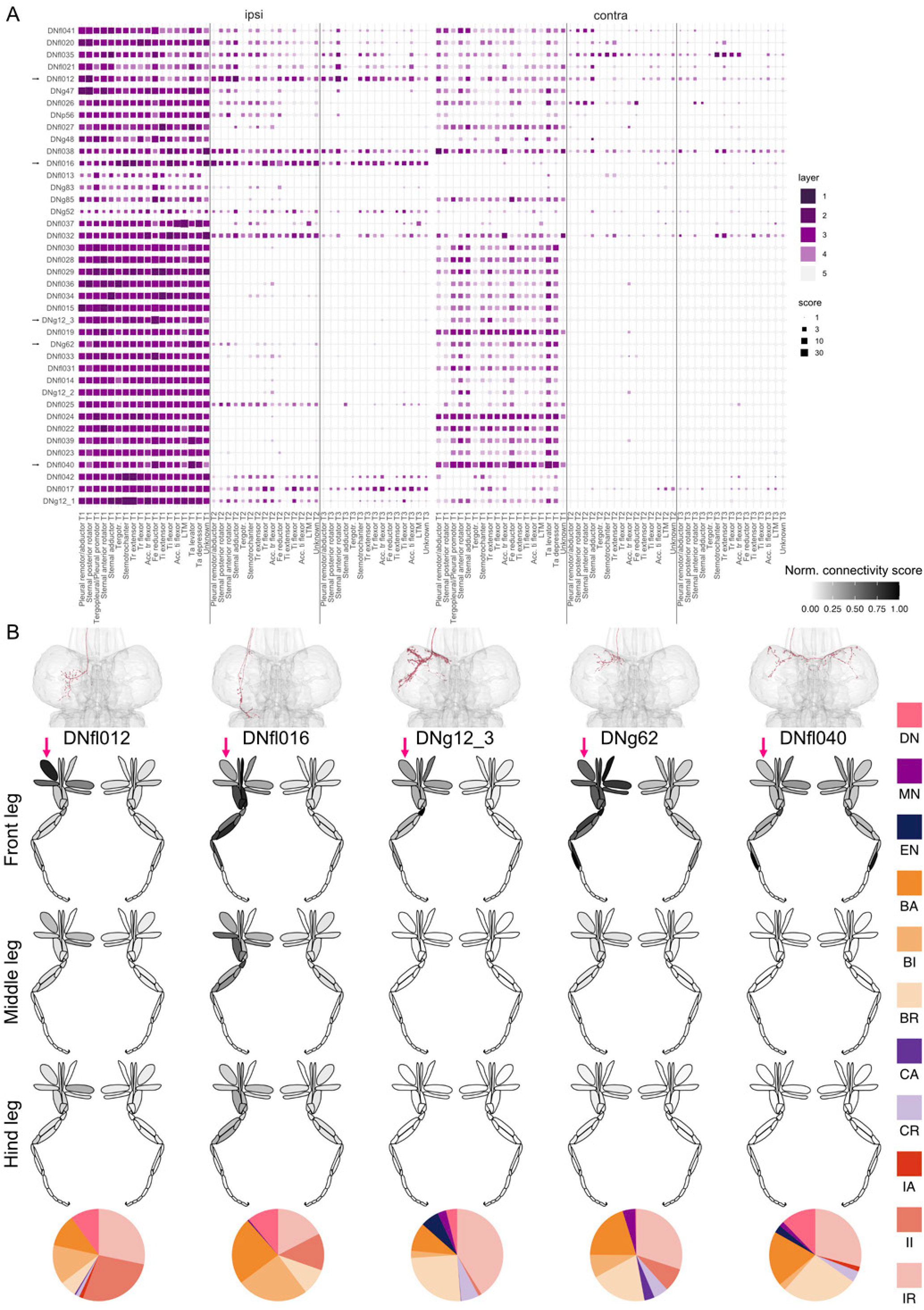
Effective connectivity of DNfl subclass neurons. **A.** The effective connectivity of DN types to the leg MN muscle targets ipsilateral and contralateral to the root side of the DN. The MNs are separated by the three segments T1-T3. The shade of magenta encodes for the layer in which the DN targets the MN (1-5), while the size of the square reflects the connectivity score, which reflects the strength of the connection (see Materials & Methods for details). **B.** Examples indicated with an arrow in **A**. The top row shows the morphology of the DN. The pink arrow indicates the root side of the DN. Underneath are schematics of the 6 leg muscles with the normalized connectivity score to the different muscles highlighted in shades of grey. The bottom row shows the percent direct output to neuron classes and subclasses in the form of prefixes (Marin et al., 2024).

**Figure 15—Supplement 2:**
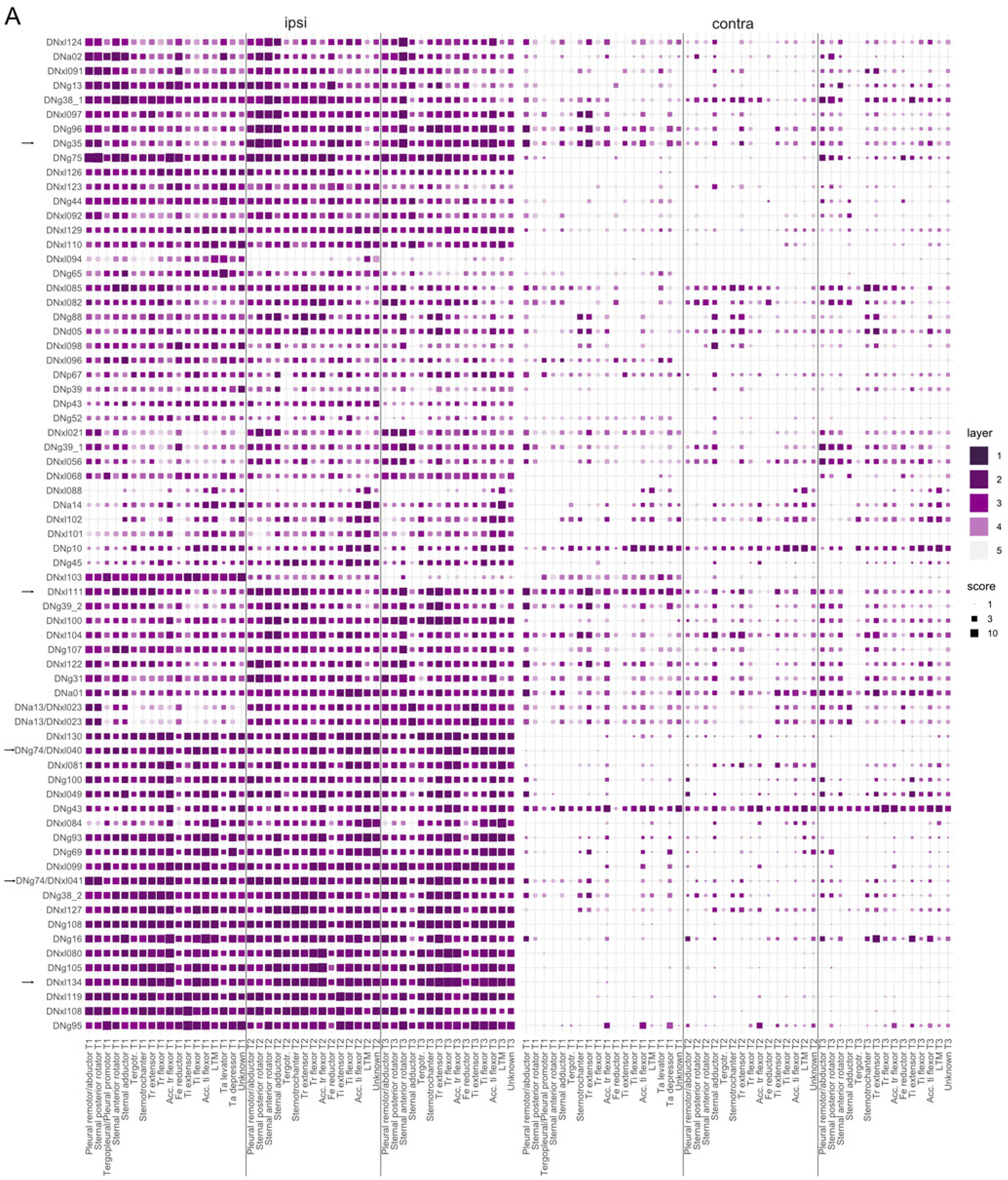
Effective connectivity of DNxl subclass neurons. **A.** The effective connectivity of DN types to the leg MN muscle targets ipsilateral and contralateral to the root side of the DN. The MNs are separated by the three segments T1-T3. The shade of magenta encodes for the layer in which the DN targets the MN (1-5), while the size of the square reflects the connectivity score, which reflects the strength of the connection (see Materials & Methods for details).

**Figure 15—Supplement 3:**
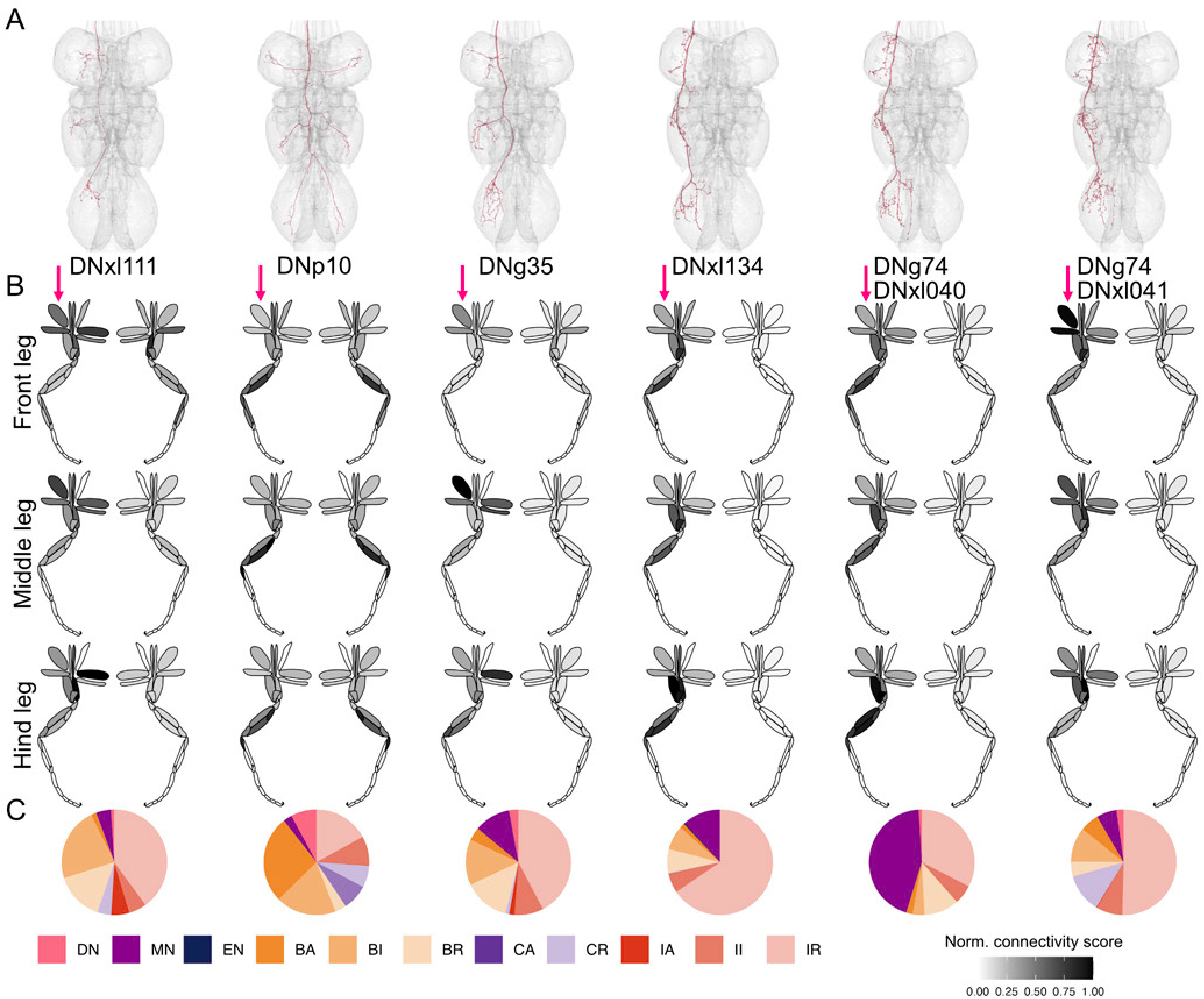
Examples of DNxl subclass neurons. Examples indicated with an arrow in Figure 15–Supplement 2. **A.** Morphology of the DNxl neurons. The pink arrow indicates the root side of the DN. **B.** Schematics of the six leg muscles with the normalized connectivity score to the different muscles highlighted in shades of grey. **C.** The percent direct output to neuron classes and subclasses in the form of prefixes (Marin et al., 2024).

**Figure 15—Supplement 4:**
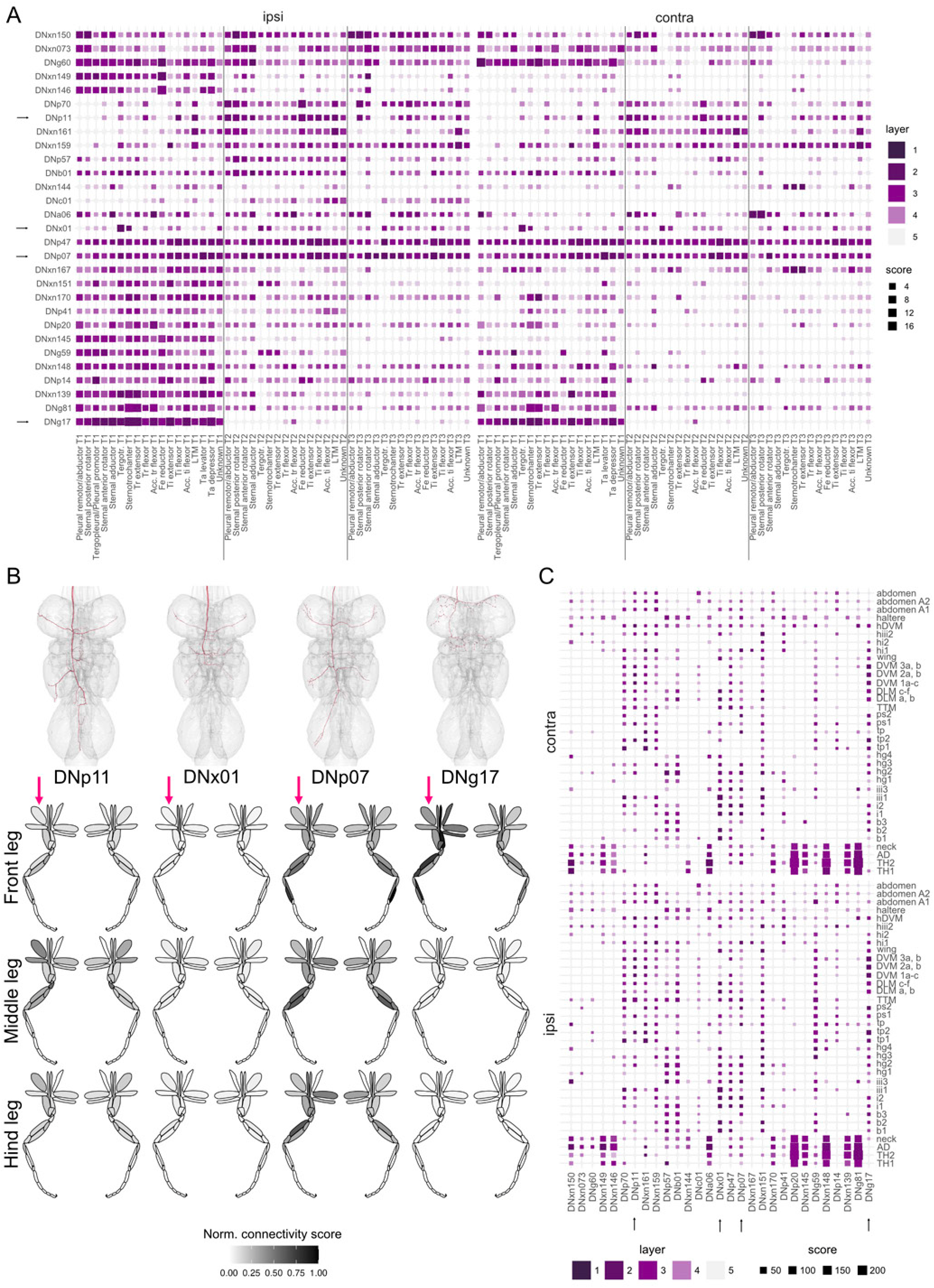
Effective connectivity of DNxn subclass neurons. **A.** The effective connectivity of DN types to the leg MN muscle targets ipsilateral and contralateral to the root side of the DN. Only those DNxn neurons were chosen that had the highest effective connectivity scores to leg MNs. The MNs are separated by the three segments T1-T3. The shade of magenta encodes for the layer in which the DN targets the MN (1-5), while the size of the square reflects the connectivity score, which reflects the strength of the connection (see Materials & Methods for details). **B.** Examples indicated with an arrow in **A**. The top row shows the morphology of the DN. The pink arrow indicates the root side of the DN. Underneath are schematics of the 6 leg muscles with the normalized connectivity score to the different muscles highlighted in shades of grey. **C.** DNxn neurons innervate several neuropils. The effective connectivity to other MNs targets are shown, separated by ipsilateral and contralateral.

Of the DNs innervating the T1 leg neuropils (i.e. the DNfl subclass), DNg12 is the only type previously identified at light level (Court et al., 2020) and is implicated in anterior grooming, including front leg rubbing and ventral head sweeps (Guo et al., 2022). The effective connectivity analysis showed both that DNg12 targets almost all leg muscles and that the majority of other foreleg DNfl neurons have similar connectivity (Figure 15–Supplement 1). DNfl040 is one of the few bilaterally projecting neurons, also targeting MNs on the contralateral side. Most DNfl neurons exclusively target MNs in the T1 leg segment, but a few, including DNfl012 and DNfl016, may have multi-hop pathways to MNs in T2 and T3. While we cannot be certain of the function of the uncharacterised DNfl neurons, we speculate that the majority are involved in grooming or reaching rather than walking, given their similar patterns of effective connectivity to DNg12.

The largest subclass of leg DNs are the DNxl neurons innervate several leg neuropils. DNxl effective connectivity is more diverse than the DNfl subclass (Figure 15–Supplement 2). We picked out representative examples that specifically target the Tibia flexor muscle (DNp10); target different thorax muscles in T2 and T3 (DNg35); directly or indirectly target different muscle groups in different leg segments (DNxl111); target similar MNs and with similar morphology (DNxl134 and DNg74-DNxl040); or can be matched to the same light microscopy type but have differences in their connections onto Trochanter extensor MNs in T2 and T3 (DNg74-DNxl040 and DNg74-DNxl041, Figure 15–Supplement 3). We also looked at DNxn neurons (i.e. those that also target non-leg neuropils) that have strong effective connectivity with leg MNs (Figure 15–Supplement 4A). Many also target other MNs in the VNC (Figure 15–Supplement 4C). For example, DNp07 and DNp11 drive escape-related behaviors requiring simultaneous movement of both the wings and legs (Ache et al., 2019; Dombrovski et al., 2023). In section 2.7, we discuss circuits in the lower tectulum that may contribute to this inter-neuropil coordination.

One theme of these analyses is that we can often make predictions for the function of previously uncharacterized DNs by noting similarities in connectivity to previously studied neurons. In the following two sections, we investigate DNs similar to the Moonwalker DN (MDN), which promotes backwards walking (Bidaye et al., 2014; Guo et al., 2022) and DNa02, known for turning while walking (Rayshubskiy et al., 2020).

#### 2.6.4 Turning DNs

DNa02 has been described to increase locomotor activity when activated bilaterally (Cande et al., 2018). However, there is also strong evidence that lateralized DNa02 activity drives flies to turn towards the activated side during walking (Rayshubskiy et al., 2020). We used effective connectivity to identify leg DNs with similar MN connectivity patterns (Figure 15–Supplement 2). Of previously identified DNs, we found that DNg13 showed a highly similar effective connectivity fingerprint. Notably, light-level imagery shows that both DN types have similar dendritic morphology in the brain (Namiki et al., 2018; Rayshubskiy et al., 2020) (Figure 16A). Both receive input from the lateral accessory lobe (LAL), an integration center of multimodal inputs from higher brain areas associated with steering (Namiki and Kanzaki, 2016; Scheffer et al., 2020). However, while DNa02 descends into the VNC on the ipsilateral side, DNg13 crosses in the brain and innervates the contralateral hemisegments in the VNC (Figure 16A). In the published literature, DNg13 is suggested to be involved in reaching movements (Cande et al., 2018), but the following circuit observations are consistent with both types having a role in turning, which has been confirmed by a recent study (Yang et al., 2023).

**Figure 16:**
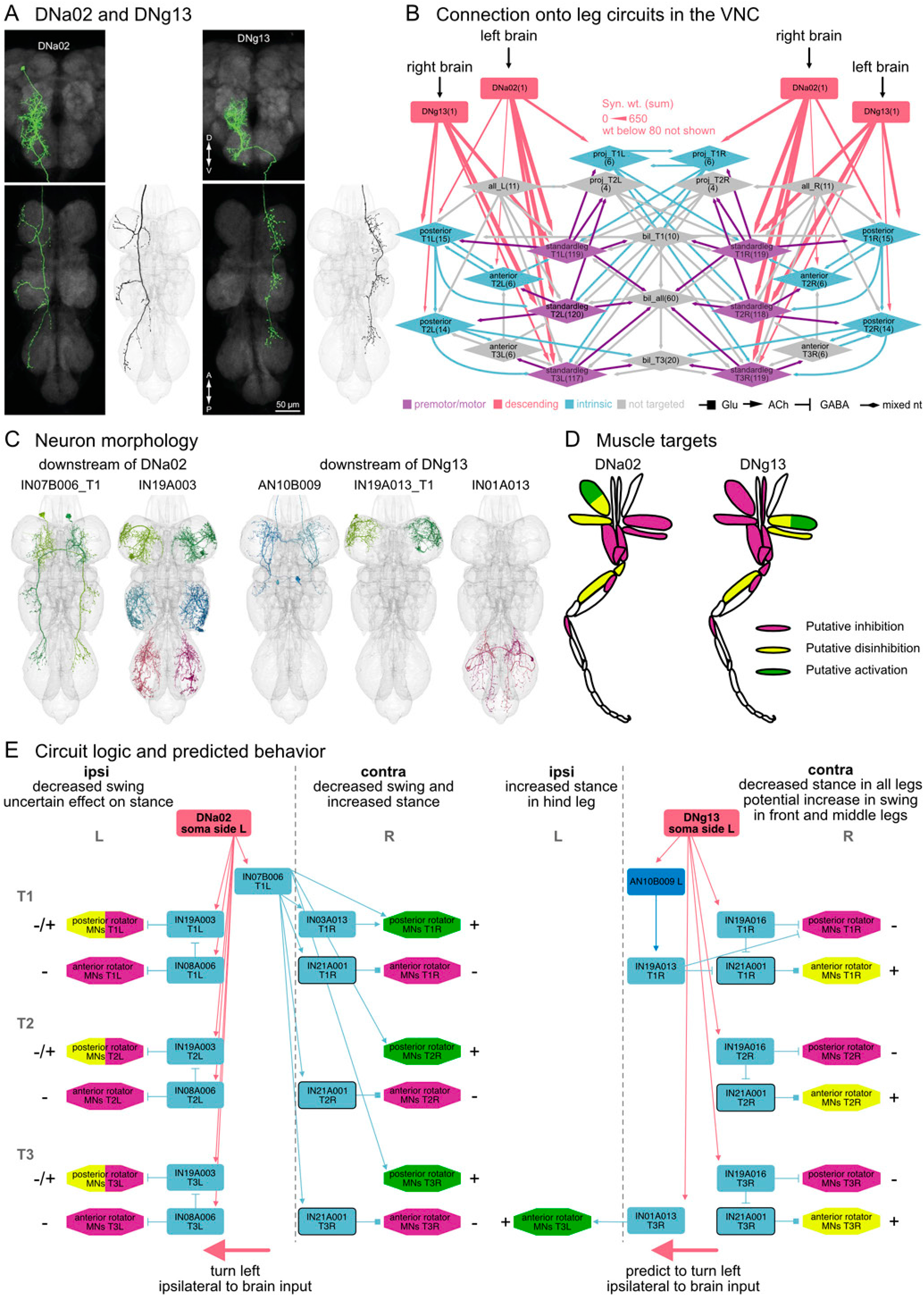
Connectivity of DNa02 and DNg13. **A.** Light microscopy images of DNa02 and DNg13 from (Namiki et al., 2018) with their MANC axonic match. **B.** Connection of DNa02 and DNg13 in MANC onto the leg coordination circuit groups from Figure 14. **C.** Morphology of selected neurons downstream of DNg13 or DNa02. **D.** Leg Muscles targeted by the two DNs, coloured by putative inhibition, activation or disinhibition. **E.** Illustration showing the connectivity across segments and sides for selected downstream circuitry with the predicted putative inhibition, disinhibition and activation of MNs marked on the ipsilateral and contralateral side in respect to the brain morphology and soma location. Note that the connections of only one DN (left side dendrite and soma DN) of the pair is shown, the right side DN has the same mirrored connectivity, but is omitted for clarity. Furthermore, we omitted circuitry of these DNs to MNs other than the swing/stance MNs for clarity. Neurons that are common downstream partners of both DNa02 and DNg13 are marked by a black stroke around the node.

**Figure 16—Supplement 1:**
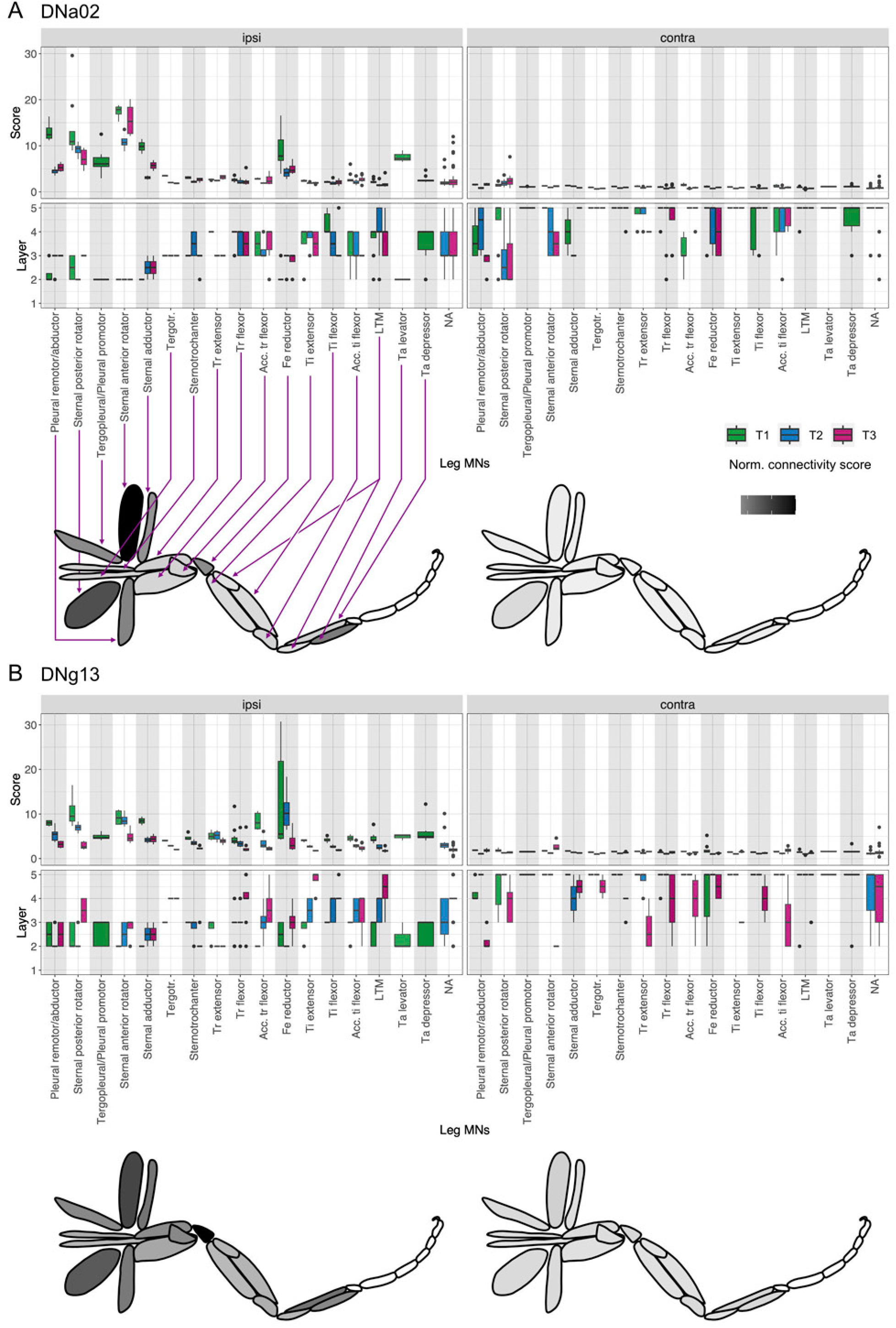
Effective connectivity of DNa02 and DNg13. Connectivity to ipsi- and contralateral leg MNs for the three neuromeres separately. **A.** DNa02 and **B.** DNg13 connectivity. Connectivity score represents the strongest connection in the network (either direct or via interneurons, four layers at maximum) based on the normalized matrix product of percent of input to the receiving neuron (top row, see Materials & Methods). The layer in which the highest connectivity score was present is shown in the bottom row plots. First layer represents direct DN to MN connectivity, second layer is DN→IN→MN, third layer includes two interneurons and so on. Leg muscle cartoons show a normalized score averaged across all three legs for each muscle. Darker color indicates stronger effective connectivity from the DN to MNs controlling the muscle.

The effective connectivity analysis identifies the MNs and muscle groups in the leg that receive the strongest input from DNa02 and DNg13 (Figure 16–Supplement A, B) but does not reveal whether these connections are excitatory or inhibitory. Indeed although the axons of both DNs extend into all three leg neuropils, they have very distinct morphologies, suggesting that their first order partners may be different. Thus, we first established which groups of neurons in our intra- and inter-leg circuits are targets of these two DNs (Figure 16B), and then probed the more complex details of these DN target circuits (Figure 16E).

The targets of DNa02 and DNg13 include many components of the standard leg connectome, encompassing downstream partners from all three leg neuropils (shown in purple in Figure 16B).

However, they differ in the groups that they target upstream of the serially repeated inter-leg circuit: DNa02 targets sequential neurons that project up from T2 to T1, while DNg13 targets sequential neurons that project down from T1 to T2 and from T2 to T3 (illustrated in light blue in Figure 16B). Moreover, DNa02 strongly targets the T1 pair of IN07B006, a group of predicted cholinergic neurons with input from T1 on one side and projections to all 3 contralateral leg neuropils; this suggests a role coordinating premotor circuits on the two sides of the VNC (Figure 16C and E).

A more granular examination of the efferent connectivity of DNa02 and DNg13 reveals clear differences in their connection to MNs in the Thorax and Trochanter (Figure 16E, Figure 16–Supplement 1 A, B). DNa02 (shown on the left) targets two GABAergic serial sets of INs (IN19A003 and IN08A006) that inhibit the MNs involved in the rotation of the leg during stance and swing phase on the ipsilateral side, suggesting a reduction of stride on the ipsilateral side consistent with the previous behavioral study (Rayshubskiy et al., 2020). Through a cholinergic bilaterally interconnecting neuron mentioned above (IN07B006_T1, see Figure 16E), DNa02 directly targets the serial MN set involved in stance (Sternal posterior rotator MNs) and indirectly targets the same MN set in T1 (Sternal posterior rotator MNs) via IN03A013, suggesting an increase in the stance especially in the front legs. The activation of Sternal posterior rotator MNs could result in leg movements that push the body forward on the contralateral side of the DNs dendrites thus moving the body towards the ipsilateral side. Interestingly, IN07B006_T1 connects to a glutamatergic (i.e. predicted inhibitory) set of neurons (IN21A001) on the contralateral side which in turn inhibit the swing driving Sternal anterior rotator MNs; this may coordinate the necessary inhibition of MNs with opposing effect. In contrast, DNg13 (shown on the right) targets the stance/power MNs via the inhibitory serial set IN19A016, contralateral to the brain input. A strong connection to the swing/reach MNs occurs via a cholinergic AN (AN10B009) which potentially disinhibits the MNs through two GABAergic serial sets (IN19A013 and IN21A001), although the connections are only observed in T1. This disinhibition could explain the previously observed reaching when the DN is activated on both sides of the brain (Cande et al., 2018). However, a single DNg13, could contribute to turning during walking by increasing the swing specifically in the front leg and slightly in the middle and hind legs on the contralateral side (IN19A016 also connects to the T2 and T3 pair of IN21A001, potentially disinhibiting the Sternal anterior rotator MNs). We additionally see DNg13 connecting to an excitatory neuron IN01A013 which is only present in the hind legs and projects to the ipsilateral hind leg and abdomen (see Figure 16C for morphology). This contributes to the asymmetric activation of MNs necessary for turning by activating the swing/reaching MNs (Sternal anterior rotator MNs) in the hind legs ipsilaterally (Figure 16E). In summary, we show that while the two DNs target similar MNs, they do so through different circuit patterns, thus likely resulting in opposing muscle activation for the thorax rotator muscles (Figure 16E).

DNa02, as described in (Rayshubskiy et al., 2020), likely elicits ipsilateral turns by reducing stride length on the ipsilateral side and increasing rotational movement on the contralateral side. DNg13 targets the contralateral side, likely decreasing the stance phase in all legs and potentially increasing rotation forwards during the swing phase in the front legs and ipsilaterally increasing stance in only the hind leg. We would expect this to elicit turns ipsilateral to the brain innervation (Figure 16E). Taken together, these results suggest a simple model in which DNa02 and DNg13 may cooperate to elicit turning, with one shortening the stride on the inside and increasing the power phase on the outside of the turn, while the other lengthening the stride on the outside of the turn.

#### 2.6.5 Walking DNs

In *Drosophila*, the Moonwalker Descending Neurons (MDN) act as command neurons to elicit backward walking (Bidaye et al., 2014; Court et al., 2020; Sen et al., 2017), while DNa01 triggers forward walking with a steering component (Cande et al., 2018; Chen et al., 2018; Rayshubskiy et al., 2020). Intriguingly, despite controlling opposite behaviors, both DNa01 and MDN target partially overlapping muscles and INs in the leg neuropils, which are further shared with two DNs, DNxl023 and DNxl024, now identified as DNa13 (Stürner et al. 2024), that have similar VNC morphology as DNa01 (Figure 17A and B, Figure 17–Supplement 1 and 2). Some downstream neurons are targeted by all of these DNs (marked by a black stroke in Figure 17C and E). However, notable differences in muscle targets can be observed between DNs when examining front, middle and hind legs and the two sides independently (Figure 17D and E). DNa01 has been described in steering (Chen et al., 2018; Rayshubskiy et al., 2020) and is similar to DNa02 in its front leg targets, which inhibit the posterior rotator muscles in the thorax. However, the other downstream neurons likely important for DNa02-elicited turning are not shared with DNa01, suggesting that DNa02 utilizes a different circuit mechanism for controlling turning that can act in parallel with DNa02 activity, consistent with previous conjecture and functional experiments (Rayshubskiy et al., 2020). Turning requires an asymmetric circuit control between sides across the 6 legs, as described for DNa02 and DNg13 in the previous section. Indeed, we see this asymmetry in DNa01 onto the tibia extensor MNs. DNa01 inhibits the tibia extensors in all three ipsilateral legs through a serial set of GABAergic neurons, IN12B003, which are disinhibited contralaterally through two bilaterally projecting INs (IN07B009_T1 and IN07B104).

**Figure 17:**
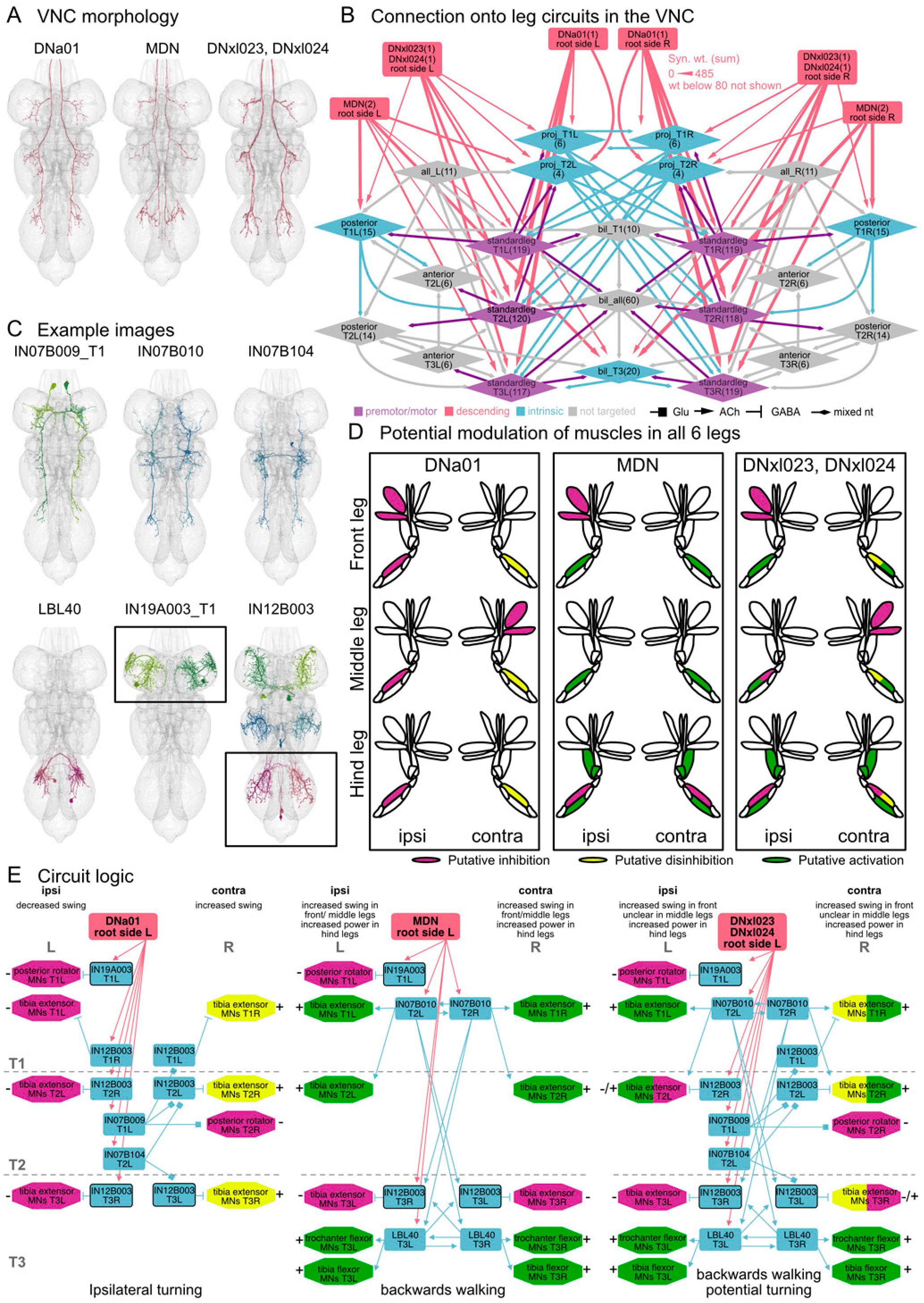
Connectivity of DNa01, MDN and DNa13. **A.** Axon morphology of identified DNa01, MDN and DNa13 that target the three leg neuropils. **B.** Connection of DNa01, MDN and DNa13 in MANC into the leg coordination circuit groups from Figure 14. **C.** Morphology of selected MANC neurons downstream, including LBL40 (Feng et al., 2020; Rayshubskiy et al., 2020). **D.** Predicted modulation of the leg muscles across the 6 legs, coloured by putative inhibition, activation or disinhibition. **E.** Illustration showing the connectivity across segments and sides for selected downstream circuitry of the DNs. Predicted putative inhibition, disinhibition and activation of MNs marked on the ipsi and contralateral side in respect to the brain morphology and soma location of only one DN (left side dendrite and soma DN for DNa01 and MDN, left root side for DNa13). For clarity and comparison the circuitry concentrates on connections from the neurons shown in **C**. Neurons that are common downstream partners of all 4 DN types are marked by a black stroke around the node.

**Figure 17—Supplement 1:**
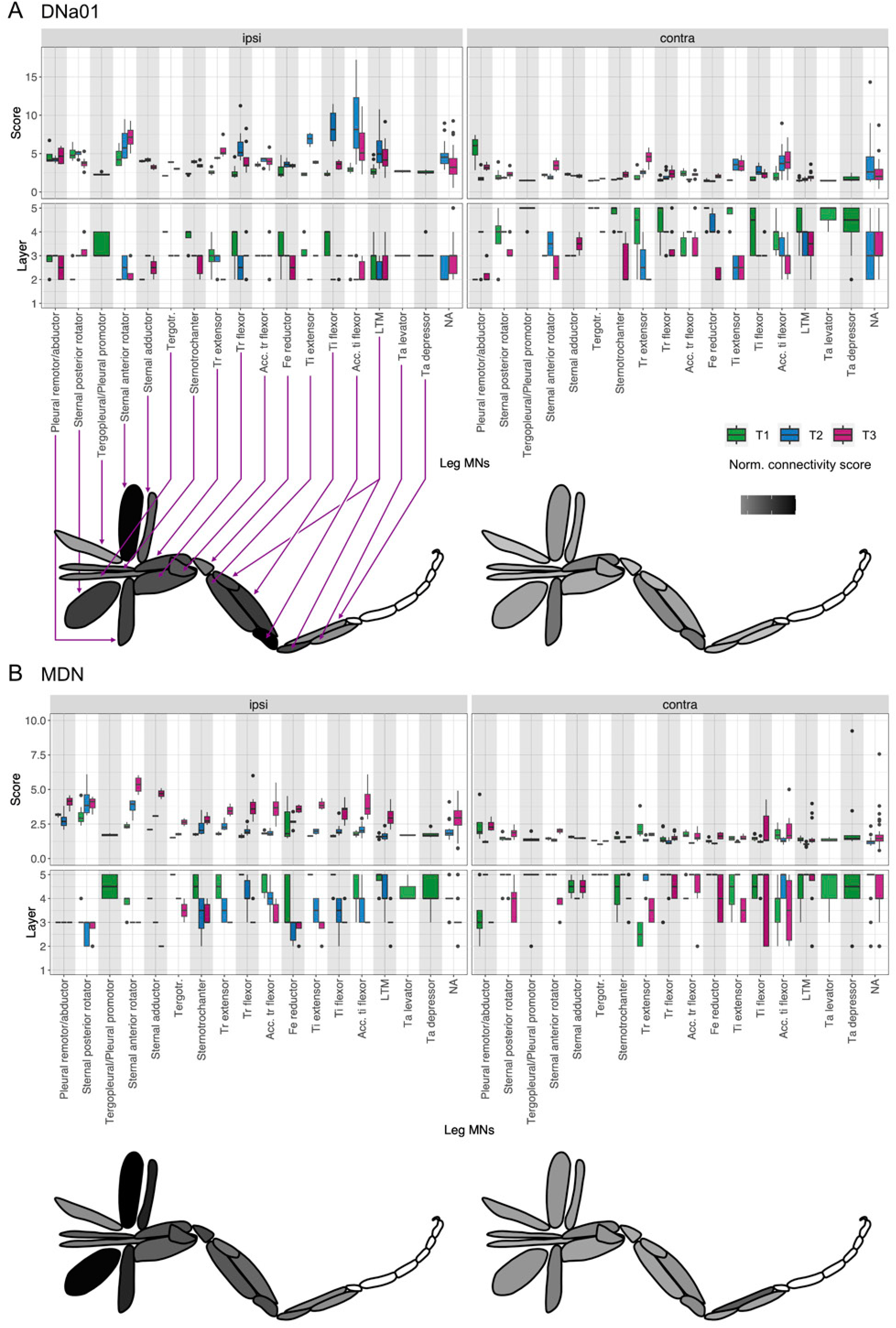
Effective connectivity of DNa01 and MDN. Connectivity to ipsi- and contralateral leg MNs for the three neuromeres separately. **A.** DNa01 and **B.** MDN connectivity. Connectivity score represents the strongest connection in the network (either direct or via interneurons, four layers at maximum) based on the normalized matrix product of percent of input to the receiving neuron (top row, see Materials & Methods). The layer in which the highest connectivity score was present is shown in the bottom row plots. First layer represents direct DN to MN connectivity, second layer is DN→IN→MN, third layer includes two interneurons and so on. Leg muscle cartoons show a normalized score averaged across all three legs for each muscle. Darker color indicates stronger effective connectivity from the DN to MNs controlling the muscle.

**Figure 17—Supplement 2:**
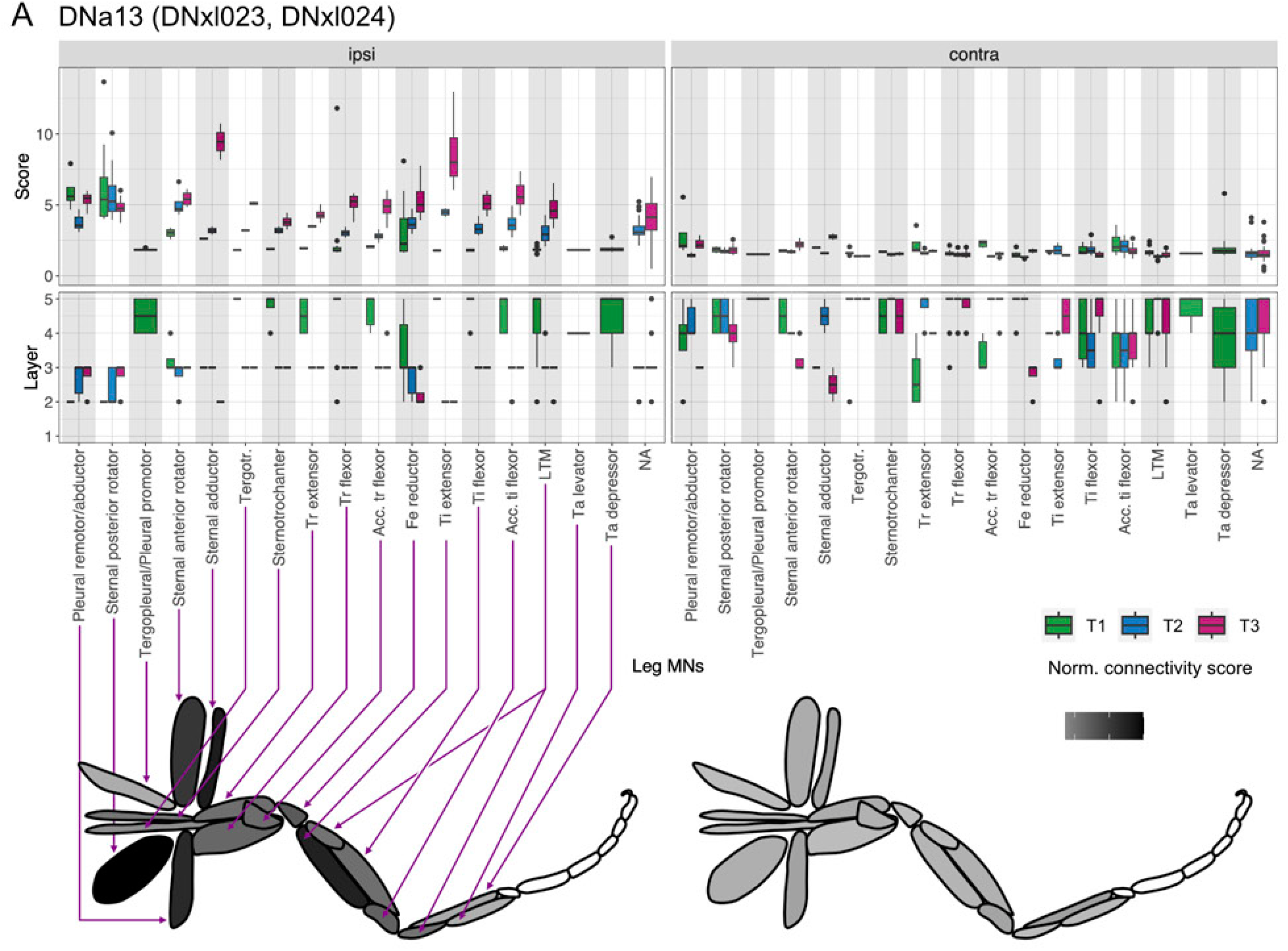
Effective connectivity of DNa13. Connectivity to ipsi- and contralateral leg MNs for the three neuromeres separately. **A. DNa13**connectivity. Connectivity score represents the strongest connection in the network (either direct or via interneurons, four at maximum) based on the normalized matrix product of percent of input to the receiving neuron (top row, see Materials & Methods). The layer in which the highest connectivity score was present is shown in the bottom row plots. First layer represents direct DN to MN connectivity, second layer is DN→IN→MN, third layer includes two interneurons and so on. Leg muscle cartoons show a normalized score averaged across all three legs for each muscle. Darker color indicates stronger effective connectivity from the DN to MNs controlling the muscle.

The two previously unidentified DNs have a strong similarity to MDN in the INs that they target, especially in the Trochanter flexor and Tibia extensor circuits (Figure 17D and E). We find that the strongest connection in the hind legs is through the previously described LBL40 bilateral cholinergic neuron that is one of the top targets of MDN (bodyid: 10994, 11493). This neuron is sufficient to elicit backwards walking (Feng et al., 2020; Rayshubskiy et al., 2020) and activates the tibia and trochanter flexor muscles in the hind legs. We were surprised by how strongly MDN also modulates the leg muscles in the middle and front legs. Backwards walking requires a switch of the protractor retractor muscles in the thorax (Rosenbaum et al., 2015). The specific inhibition of the posterior rotator MNs in T1 through IN19A003 could be the basis of that switch when MDN is bilaterally activated, inhibiting forward movement. Additionally, we speculate that a bilateral interconnecting neuron, IN07B010, strongly targeted by LBL40 and MDN, coordinates the two sides and contributes to the switch by increasing tibia extension in the front legs and decreasing it in the hind legs. The strong targeting of MDN to the hind legs is consistent with the observed strongest impact on the hindlegs when MDNs are artificially activated (Bidaye et al., 2014; Feng et al., 2020). Ipsilateral activation of MDN (in respect to the soma hemisphere) has been implied to elicit contralateral turning during backwards walking (Sen et al., 2017). MDN has bilateral dendrites in the brain with an axon extending contralateral of the soma. We therefore propose that MDN might receive similar input in both brain hemispheres and that the described turning could be rather due to the ipsilateral inhibition of IN19A003_T1 in the front leg, which in turn inhibits the posterior rotators. Please note that Figure 17E shows this connectivity to be ipsilateral because we are looking from the root entry point of the MDN axon into the VNC. This is, however, contralateral to the soma, due to MDN’s brain morphology (Bidaye et al., 2014). IN19A003_T1 is notably the same interneuron that is inhibited ipsilaterally by DNa02, which we propose to be involved in the ipsilateral turning phenotype during forwards walking (Figure 16E).

DNa13 targets most of the major partners of MDN and DNa01 (Figure 17E). Based on the similarity of the connectivity to MDN, we expect them to also elicit a switch to backwards walking. Due to the asymmetry in connectivity across the two sides, similar to DNa01, we suggest these DNs could also play a role in turning. This proposed turning while walking backwards remains to be experimentally confirmed.

The connectivity downstream of DNs can be difficult to interpret without first understanding the general leg coordination circuit. Strong connections from a DN to a given MN, even when the target muscle is known, does not automatically predict which behavior activation of that DN will elicit. By first considering the standard circuit logic of the leg premotor circuits and then analyzing how these circuits are interconnected, we have established a basis for interpreting the effect of single DN inputs on motor behavior. The DNa02 and DNg13 examples illustrate how we can use this dataset and its annotations to understand how a known behavior, ipsilateral turning, can be coordinated, and how we can find other DNs that could be involved in the same behavior. The DNa01, MDN and DNa13 examples show us how targeting the same muscles in different segments can elicit very different behaviors, predicted two new DNs that could be involved in backwards walking, and identified an IN that could be important for the switch from forwards to backwards walking.

### 2.7 Organizational logic of DNs onto intermediate neuropil circuits

#### 2.7.1 Organizational logic of lower tectulum circuits

In the sections above, we analyzed premotor circuits in the dorsal upper tectulum (UTct) controlling the wing motor system and in the ventral leg neuropil (LegNp) controlling the leg motor system. Intriguingly, our community network analysis implicated the remaining neuropil areas intermediate between these dorsal and ventral layers (intermediate tectulum, IntTct, and lower tectulum, LTct) as potential areas of higher order processing because they 1) have little ramification by MN dendrites, and 2) have more microcircuit motifs with local recurrence compared to other VNC areas. IntTct, which is largely associated with community 9, is tightly interconnected with the UTct communities controlling the wings, halteres, and neck (communities 8, 10, 16, and 17, see Figure 9A). It thus likely serves a higher order control purpose specialized for flight or other actions of the wing motor system. The LTct, however, includes several communities (e.g. 7 and 14), which are distinct from those already discussed in that they provide output to neurons in both the wing and leg motor systems. In particular, the lower tectulum (LTct) is innervated by DNs, INs, and MNs critical for initiating takeoff or landing, suggesting that this region is an important hub for initiating behaviors that require coordinated wing and leg motion (Namiki et al., 2018) (Figure 18A). In this section, we use the MANC data to investigate the connectivity of neurons that innervate the LTct and identify pathways linking DNs related to escape behaviors with their motor effectors.

**Figure 18:**
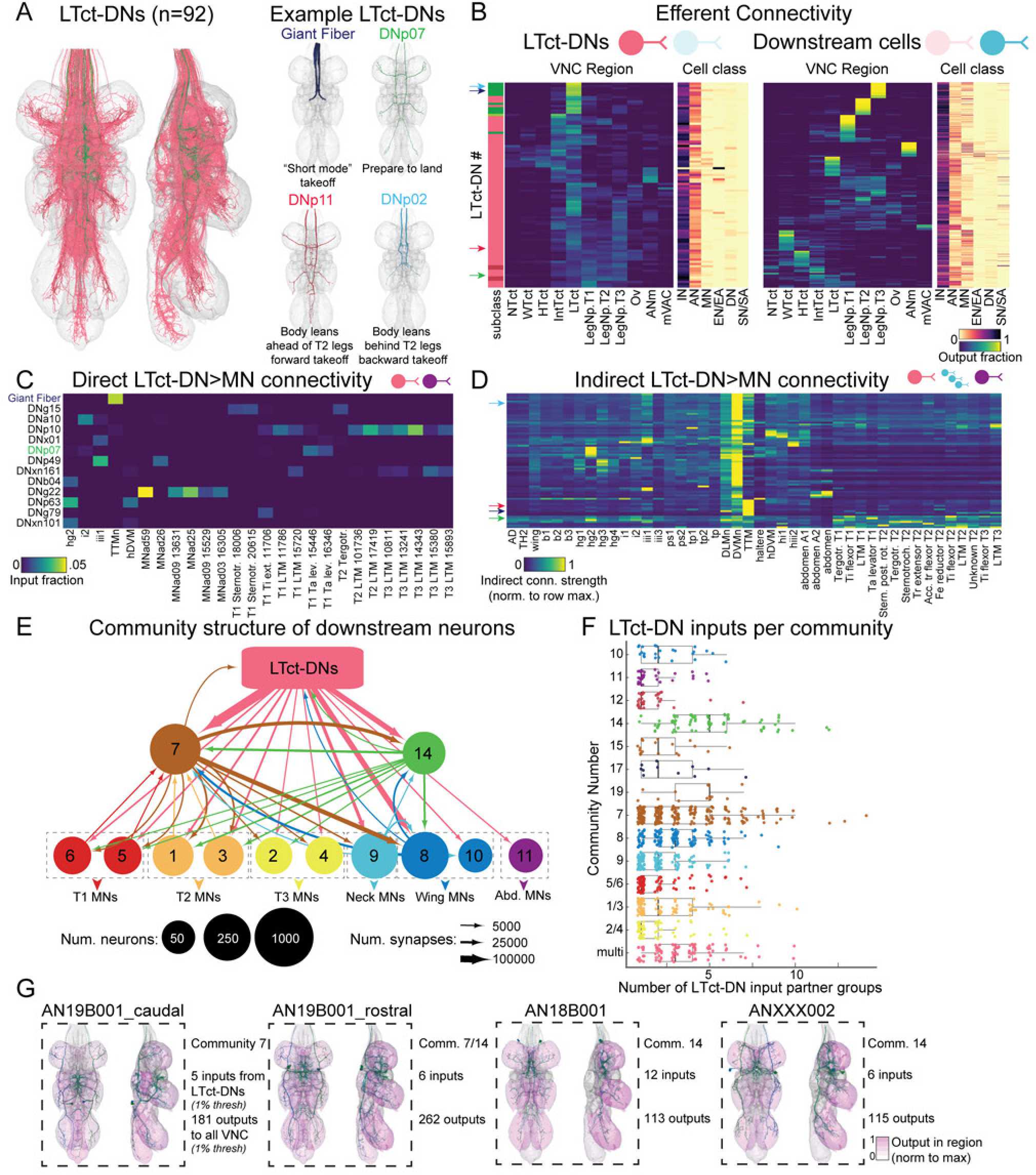
Overview of lower tectulum circuits. **A.** Left: morphology of the LTct-DNs (DN groups with ≥100 synapses and ≥10% of their synaptic output in the lower tectulum). Color corresponds to DN subclasses (e.g. xl, xn, lt) as in Figure 3. Right: morphologies of LTct-DNs with known behavioral functions. **B.** Left: Efferent connectivity of LTct-DNs. Left sidebar, subclass of LTct-DNs, with colors as in **A** (left panel). Colored arrows on the left of the sidebar indicate rows that correspond to examples in **A**. Left heatmap, proportion of each LTct-DN’s synapse-wise output in each region of the VNC. Right heatmap, proportion of LTct-DN’s synapse-wise output, onto each downstream cell class (counting only neurons receiving ≥1% of their synaptic input, minimum 50 synapses, from an LTct-DN). Right: as on the left side, but each row corresponds to one AN, IN, or EN downstream of an LTct-DN (“LTct-DN downstream neurons”, same connectivity criteria as above). **C.** Heatmap depicting all direct connections between LTct-DNs and MNs (groupwise). Any DNs that do not provide ≥1% groupwise synaptic input to a MN and any MNs that do not receive ≥1% groupwise synaptic input from an LTct-DN were excluded. Groups on the x-axis are named according to MN type; if types do not refer to a unique group then group number is appended. **D.** Heatmap depicting normalized indirect synaptic connectivity (as in Figure 10H) from each LTct-DN onto all MNs within five downstream layers. Any DNs that do not have an indirect connectivity score ≥5 onto any MN and any MNs that do not receive indirect connectivity ≥5 from any LTct-DN were excluded. For visualization, all heatmap values are re-normalized based on row-wise maxima. Colored arrows indicate rows corresponding to example DNs in **A**. **E.** Diagram depicting connectivity between neurons directly downstream of LTct-DNs (same subset as in right panels of **B**), binned according to the Infomap community they were assigned to (see Figure 9). Minimum connectivity to display an edge is 5000 synapses, and communities with fewer than 60 LTct-DN downstream members are not displayed. **F.** Communities 7 and 14 receive high levels of LTct-DN input. Number of LTct-DN input partners (neurons providing ≥1% of synaptic input) per downstream neuron, segregated by community assignment on the y-axis. **G.** Examples of neurons from communities 7 and 14 that receive high levels of LTct-DN input (≥1% of input from ≥5 LTct-DNs) and project widely throughout the VNC. Regions in diagrams are colored according to the amount of output they receive from the depicted neuron (normalized to the region with the highest output). Text indicates the community membership of each neuron, the number of inputs from LTct-DNs to each example neuron, and the number of outputs from that neuron to all neurons in the VNC (≥1% threshold).

To begin investigating LTct function and its descending control, we first identified the subset of DNs which output in the LTct (“LTct-DNs”, 92 DN groups with ≥10% of their output and ≥100 synapses in the region, Figure 18A). We note that only a minority of LTct-DNs exclusively targeted the LTct; most sent additional projections to either the UTct or the leg neuropils (but not both) (Figure 18B). Looking into the cell class composition of DN downstream neurons, LTct-DNs make few direct synapses onto MNs. Instead, most LTct-DNs output onto INs and ANs (Figure 17B, C). This synaptic architecture is perhaps surprising, as one might expect that innate, rapid actions such as takeoff or landing sequences (many of which are actuated by LTct-DNs, see Figure 18A) would rely on circuits that minimize synaptic distances between DNs and MNs to increase reliability and decrease latency. Indeed, the most well-studied DN-MN pathway in the VNC, the Giant Fiber (GF) circuit that triggers a rapid escape takeoff, features direct connections between the GF and the tergotrochanteral muscle (‘jump’ muscle) MN (TTMn). We elaborate on this pathway in the following section. However, since our analyses suggested that few of the LTct-DNs follow the GF example and initiate motor activity directly, we next examined the connectivity of the network of intrinsic neurons (INs, ANs, and ENs) that are strongly postsynaptic to an LTct-DN (≥1% groupwise input and ≥50 synapses). We refer to this set as the LTct-DN Downstream neurons. These neurons generally provide most of their output to only 1-2 neuropil regions, although a sizable minority project broadly throughout the VNC (Figure 18B). Furthermore, many of them have significant output to MNs, suggesting that they play a role as downstream motor effectors of LTct-DNs (Figure 18B).

We verified these observations by quantifying both the direct and indirect synaptic input from LTct-DNs onto downstream MN targets, by groupwise input fractions and normalized indirect connectivity strength (Figure 18C, D). While direct DN-MN connectivity was sparse and relatively weak (only 13/92 LTct-DNs provided at least 1% of synaptic input onto any MN; see Figure 18C), the majority of the LTct-DNs have indirect impact on a wider range of MNs (Figure 18D). Notably, many LTct-DNs provide indirect input to both leg MNs and wing MNs, supporting earlier hypotheses that the LTct is a hub for leg-wing coordination (Figure 18D) (Namiki et al., 2018). Although some DNs had both direct and indirect influences onto the same MN groups (e.g. the GF contacts the TTMn both directly and indirectly, and DNp07 contacts tarsal levators both directly and indirectly), overall there was little correspondence between the strongest direct and indirect influences. For example, many LTct-DNs provided strong indirect input to the DLMns and DVMns, but none provided direct input to these MNs.

To survey the circuits to which the LTct-DNs contribute, we used the results of our earlier Infomap community detection analysis (see section 2.4) to examine the community demographics of LTct-DN Downstream neurons. Notably, communities 7 and 14 are enriched among this population and contain neurons that received high levels of LTct-DN input (Figure 18E, F). These communities (in particular 7) have weaker connectivity with MNs but have strong connectivity with communities that output to leg and wing MNs, consistent with a role in coordinating behavior across multiple motor systems and neuropils (Figure 18E). Among the LTct-DN Downstream cells assigned to communities 7 or 14, we identified neurons of interest (Figure 18F) that both received high levels of LTct-DN input and output broadly throughout the VNC. In particular, we identified a set of ANs with high LTct-DN input and broad output; several of these neurons are shown in Figure 18F and are described further in section 2.7.3. Taken together, these findings suggest that LTct-DNs do not rely solely upon direct outputs to neuropils containing wing or leg MNs to coordinate simultaneous leg and wing movements, but instead control them indirectly through multisynaptic motifs such as the broadly-projecting intermediates observed here and the recurrent triad motifs described in Figure 9G, H.

To examine how the general LTct organization we describe above may guide coordinated behavior, we next focused on a previously hypothesized behavioral role for LTct circuits: coordination of visually-induced escape-related behaviors (Namiki et al., 2018). In the next two sections, we look at LTct circuits specifically connected to DNs implicated in the two known types of escape takeoff: short mode (section 2.7.2) and long mode (section 2.7.3).

#### 2.7.2 LTct circuits modulating Giant Fiber-mediated escape

Behavioral responses to the threat of an approaching predator are among the most critical actions a fly performs to ensure survival. When a predator such as a damselfly attacks, its image grows rapidly in size on the fly’s retina, creating a ‘looming stimulus’ to which flies react rapidly. These rapid responses can take the form of several behaviors: the fly may start walking away from the stimulus, or it may perform an escape takeoff. Takeoff requires coordination between leg and wing motor systems: in a ‘long-mode’ takeoff, the wings are first raised, followed by a downstroke performed almost concurrently with extension of the T2 legs in a takeoff jump, while in a ‘short-mode’ takeoff, the T2 legs rapidly extend while the wings are pulled down in a ‘tuck’ against the fly’s body.

Functional studies have determined that a single action potential in a pair of large-axon descending neurons called the Giant Fiber (GFs; also known as DNp01) triggers the short-mode “tuck and jump” takeoff. The GFs have non-branching blunt terminals that end in the LTct and provide both chemical and electrical synaptic input onto downstream LTct neurons, including the TTMn and the peripherally synapsing interneuron (PSI). As mentioned earlier, the tergotrochanteral muscle is an extrinsic muscle of the leg, which extends dorso-ventrally across the thorax parallel to the DVM flight muscles. When stimulated by its single excitatory MN, the TTMn, the tergotrochanteral muscle rapidly extends the fly’s mesothoracic (T2) legs, causing the fly to push off from the ground. In turn, stimulation of the PSI activates downstream MNs controlling the dorsolateral indirect wing power muscles (DLMns), which function as wing depressors. Simultaneous activation of both via the Giant Fiber thus causes the fly to “tuck” its wings and leap into the air. This short-mode takeoff allows flies to get away from a rapidly approaching threat, but the resulting flight is unstable because they do not raise their wings before takeoff (von Reyn et al., 2014; Wyman et al., 1984).The Giant Fiber circuit is one of the best-studied motor networks in the Dipteran VNC, and its downstream and secondary output connectivity is well-documented by previous EM and dye-fill studies (Azevedo et al., 2024; Bacon and Strausfeld, 1986; Kennedy and Broadie, 2018; King and Wyman, 1980; Sun and Wyman, 1997). We validated this known GF connectivity in MANC and identified further components of the GF circuit, including a set of neurons that make strong, direct inputs to the GF axon within the VNC (Figure 19A, B).

**Figure 19:**
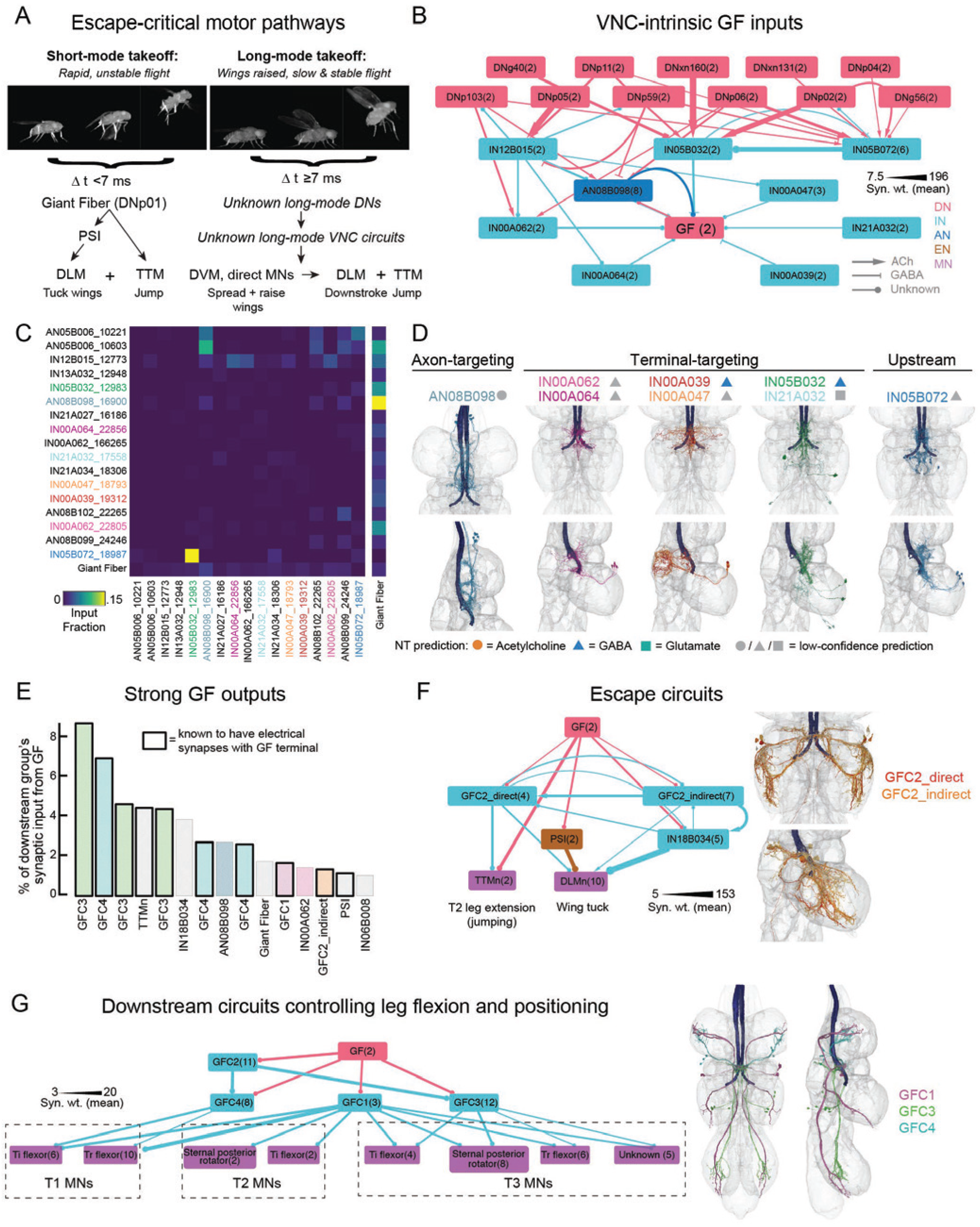
Upstream and downstream circuits of the Giant Fiber in the VNC. **A.** Schematic of differences between short-mode and long-mode takeoffs, including the characteristic patterns of motor activity, the durations of the behaviors, and the neurons known or hypothesized to be involved in each. Montages of escape sequences were collected using the flyPEZ3000 (Williamson et al. 2018). **B.** Diagram depicting connectivity of neuron types that provide at least 1% of the Giant Fiber’s VNC-intrinsic input, as well as a selection of neurons that have strong input to those cells. **C.** Heatmap showing synaptic connectivity between the 16 AN and IN groups that provide at least 1% of the Giant Fiber’s VNC-intrinsic synaptic input as well as IN05B072, a group of important second-order inputs. Connectivity is expressed as the fraction of the postsynaptic cell’s total synaptic input (groupwise). **D.** Morphology of GF-targeting ANs and INs. Top row shows morphology from a ventral perspective and bottom row shows morphology as viewed from the left side of VNC. The axons of the GF are shown in indigo in each image. Symbols next to neuron names indicate putative neurotransmitter identity: circle corresponds to a cholinergic prediction, triangle to a GABAergic prediction, and square to a glutamatergic prediction. Grey symbols indicate that neurotransmitter prediction is low confidence (prediction probability <0.7 for at least one member of group). Left, morphology of the axon-targeting group AN08B098. Middle, six types of terminal-targeting INs divided into three sets based on morphological criteria. Right, morphology of IN05B072. **E.** Major groupwise synaptic outputs of the GF. Synaptic output is plotted as the % of the downstream group’s synaptic input that is provided by the GF. Only neurons that provide at least 1% of input are shown (min. 30 synapses). Neurons with known electrical connectivity to the GF are shown in dark boxes. **F.** Left, network diagram depicting direct and indirect connectivity between GF and two downstream partners that trigger takeoff behaviors: the TTMns, which trigger jumping, and DLMns, which initiate wing tuck or downstroke, via different subsets of GFC2 neurons. Right, morphologies of the GFC2_direct and GFC2_indirect groups. The GFC2_direct set contains the following groups: 13127 and 13645. The GFC2_indirect contains the following groups: 14527, 13479, and 15505. **G.** Left, network diagram depicting connectivity between GF and downstream partners that putatively control post-takeoff positioning and leg flexion. MNs are included in this plot so long as they receive at least 50 synapses (groupwise) from GFC1, GFC3, and GFC34. All three INs contact tibia flexor MNs and trochanter flexor MNs, although they vary in their other targets and in which neuromeres they project to. Right, morphologies of the GFC1, GFC3, and GFC4 groups.

Since a single GF action potential is sufficient to initiate a short-mode takeoff, yet high-velocity escape disrupts ongoing behavior and requires a large amount of energy, we hypothesized that GF spiking is regulated by strong presynaptic inputs. Indeed, we identified many neurons intrinsic to the VNC that directly target the axon of the GF, potentially influencing the GF’s output by enhancing or curtailing the propagation of its spikes. Of the 470 DN groups in the MANC volume, the GF is among the 20 which receive 4000 or more synaptic inputs from VNC-intrinsic neurons. We identified a set of neurons that provide especially strong input to the GF (≥1% the GF’s synaptic input) and organized them based on their connectivity and morphology. Some of these cells, including types IN05B032, IN21A032, and IN00A062, synapse onto the terminal of the GF (Figure 19C, D). Depending on whether these neurons are excitatory or inhibitory (most have neurotransmitter prediction probability below a threshold of 0.7, although all high-confidence predictions are GABAergic), these INs may be capable of either blocking or directly triggering a short-mode takeoff. Other neurons synapse at locations distal to the GF terminals. Type AN08B098, for instance, comprises a group of eight ANs which synapse almost exclusively onto each other as well as the rostral regions of the GF axon; they also receive synaptic input from the GF (Figure 19A-C). Their recurrent co-activation might modulate the propagation of GF spikes.

In addition, we discovered neurons that putatively regulate these GF inputs (Figure 19B-D). For example, type IN05B072 includes six cells that collectively receive strong input (≥1% groupwise synaptic input) from ten DNs. IN05B072 cells output heavily onto IN05B032 which itself targets the GF terminal directly (Figure 19B, C). IN05B032 also receives descending input, including from many DNs that do not output strongly onto IN05B072 (Figure 19B). Given that neurons in hemilineage 5B tend to express GABA, it is likely that DNs targeting IN05B032 promote inhibition of the GF terminal while the DNs targeting IN05B072 permit GF spiking through disinhibition.

We also identified neurons downstream of the GF that, in addition to the TTMn and PSI, may be important for actuating aspects of short-mode takeoffs (Figure 19E). Several of these neurons– GFC1, GFC2, GFC3, and GFC4–were previously identified at the light level and are electrically coupled to the GF (Kennedy and Broadie, 2018). GFC2 synapses onto the TTMn, the DLMn, and to the ps1 indirect control MN according to previous connectomic analysis (Azevedo et al., 2024). Based on variations in morphology and differential downstream connectivity, we identified several subtypes of GFC2 neurons. The most prominent distinguishing feature separating these groups was how directly they were connected to downstream motor effectors. GFC2 groups 13127 and 13645, which we labeled “GFC2_direct” had relatively strong synaptic outputs onto DLMns, DVMns, and the TTMn (Figure 19F). Other GFC2 groups, which we refer to as “GFC2_indirect,” have comparatively few synapses onto these MNs, instead directing their output towards pre-MN INs. For example, GFC2_indirect group 14527 synapses strongly onto IN06B066 and IN18B034 (which projects to the DLMns). GFC2_indirect neurons also project recurrently to GFC2_direct neurons.

We confirmed prior findings that the GFC4 neuron could play a role in post-takeoff leg flexion (Azevedo et al., 2024) and further observe in MANC that GFC1 and GFC3 might serve similar functions. These neurons each contact tibia flexors in one or more legs but vary in their projection pattern, implying that they each control flexion in different sets of leg pairs (Figure 19F). In addition to being driven directly by chemical and electrical synapses from the GF, we observed strong cholinergic input to GFC4 (≥100 synapses) from the GFC2_direct neurons and to GFC3 from one of the GFC2_indirect groups. These findings collectively suggest that the GF actuates short-mode takeoffs not only via direct activation of a handful of motor effectors, but also by conducting a large ensemble of INs joined together by recurrent and feedforward motifs.

Previous work suggested that selection of long-mode vs. short-mode takeoff could be driven purely by a spike-timing mechanism: if a single GF action potential preceded activity in other DNs that evoke long-mode takeoffs, the fly would “tuck and jump” (von Reyn et al., 2014). The MANC data reveal that the GF axon is itself regulated by presynaptic inputs that may initiate, amplify, or prevent spiking, implying that state-based inputs and other descending pathways could also play a role in shaping takeoff action selection (for example, by “vetoing” an earlier GF action potential). If GF action potentials are able to propagate to the terminal, they initiate short-mode takeoffs by activating canonical downstream elements including the PSI and the TTMn. In addition, they activate an intermediate layer of premotor neurons that either amplify these actions (i.e. GFC2_direct) or coordinate more complex and slower-latency movements (i.e. GFC2_indirect, GFC1, GFC3, GFC4) such as leg flexion after jump.

#### 2.7.3 LTct circuits that may control long-mode takeoffs

Instead of the “short-mode” takeoff, flies can respond to looming stimuli by performing a “long-mode” takeoff, in which downstroke and leg extension are preceded by wing raising (Figure 19A) (Card and Dickinson, 2008a; Trimarchi and Schneiderman, 1995; von Reyn et al., 2014). While this sequence takes more time to complete, long-mode takeoffs initiate more stable flight. The VNC circuits underlying long-mode takeoffs are currently unknown, but many of the important downstream motor outputs have been identified (Figure 19A). TTMn and DLMn are likely to be utilized, respectively, for jumps and downstrokes. Wing raising likely involves activation of MNs innervating several muscle groups, including the DVM wing levator muscles, the basalar muscles (which are active prior to flight, possibly to reconfigure the wing hinge), and the tergopleural muscles (which evoke wing raising when driven and are necessary for voluntary takeoffs) (Dickinson and Tu, 1997; O’Sullivan et al., 2018). In contrast to short-mode takeoff, functional evidence suggests that GF activity is dispensable for long-mode takeoffs (Card and Dickinson, 2008a; Trimarchi and Schneiderman, 1995; von Reyn et al., 2014). Thus, activity in other DNs must be sufficient to produce long-mode takeoffs.

We have previously identified a number of DNs which are likely to contribute to long-mode takeoffs based either on anatomical evidence that they are downstream of neurons important for looming detection or functional evidence that they initiate long-mode takeoffs when optogenetically stimulated (Dombrovski et al., 2023; Peek, 2018; Williamson et al., 2018). This subset of DNs (which includes DNp11, DNp02, DNp04, and DNp06) are all LTct-DNs, suggesting that local circuits in the LTct are important for long mode takeoff coordination. We also thus expect that other LTct-targeting DNs, beyond those already identified, may also be involved in coordinating long-mode takeoffs.

How do LTct-DNs coordinate activity in these critical escape effectors to initiate long-mode takeoffs? Previously, we observed that the outputs of LTct-DNs are integrated by ANs assigned to communities 7 or 14 (see above, Figure 18E, F, G), such as AN19B001 (which we divide into rostral and caudal groupings based on anatomical and connectivity differences, see below) and AN18B001, which themselves provide some of their strongest outputs to DLMns, DVMns, TTMn, and the PSI, as well as several MNs innervating direct control wing muscles (Figure 20A). These ANs are all predicted to be cholinergic and hence excitatory. This disynaptic circuit motif (DN-AN-MN) thus represents the most direct and strongest connectivity between long-mode-associated LTct-DNs and MNs implicated in takeoff. While these AN groups all receive converging inputs from the takeoff-related DN subset, their outputs are distinct from one another. This suggests a model in which a population of DNs are each activated by threatening visual stimuli and their outputs in the VNC are integrated by these cells, each of which contribute to the coordination of a specific part of the motor pattern critical to takeoff (Figure 20A, 21C). We next looked at the connectivity of these ANs in more detail (Figure 18G, 20A).

**Figure 20:**
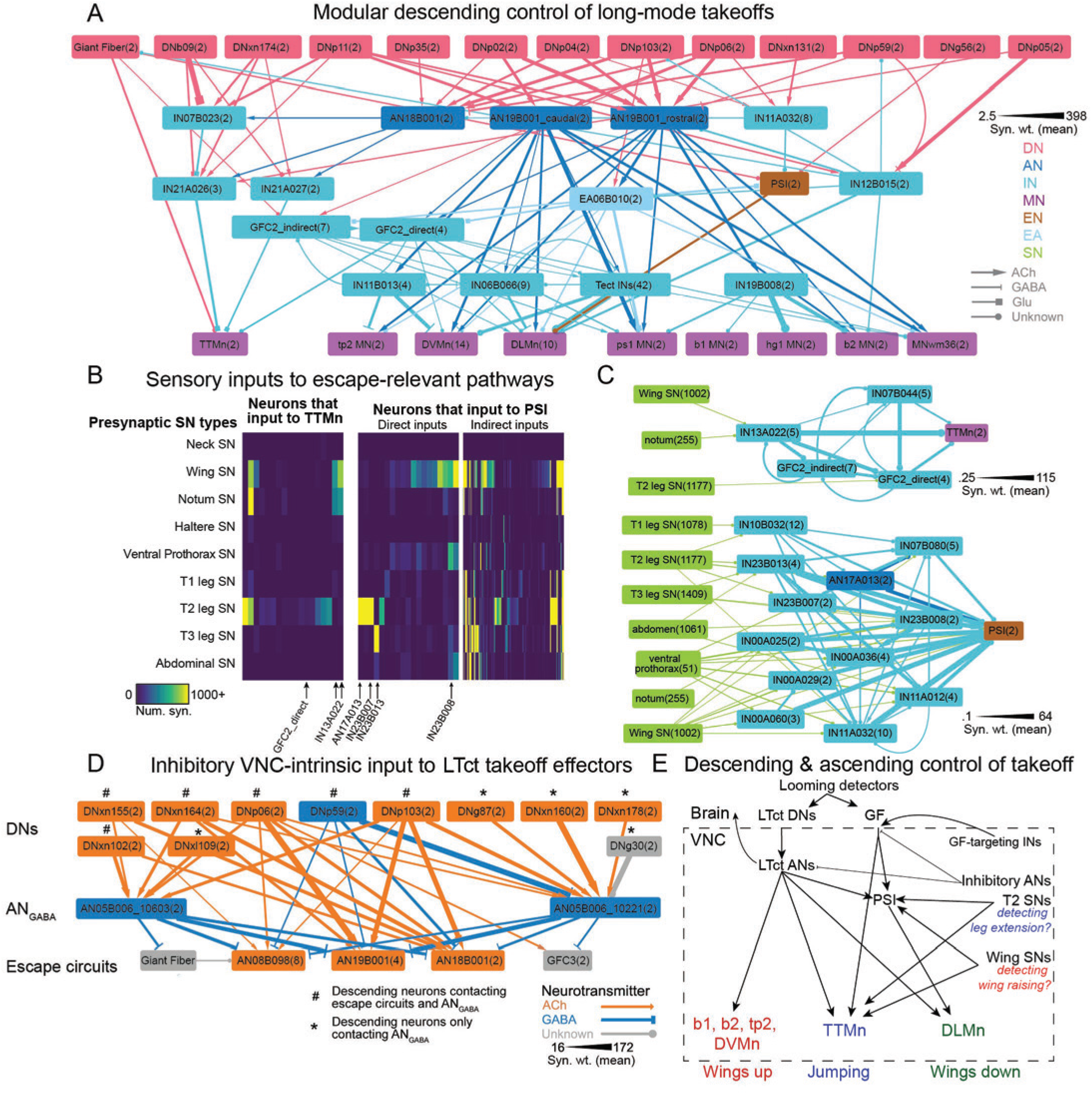
Circuits mediating long-mode takeoff. **A.** Connectivity diagram depicting strong descending connectivity onto MNs critical for long-mode takeoffs. For clarity, several groups are combined into larger categories based on shared connectivity: the “GFC2_indirect” node contains groups 14527, 13479, and 15505, the “GFC2_direct” node contains groups 13127 and 13645, the “Uni. DVMn input INs” node contains groups 17159 and 16088, and the Tect INs node contains the Wing Upper Tectular neurons described in Figure 12. MNs are grouped according to their target muscle. All other nodes contain neurons with a shared type. **B.** Connectivity between SNs and escape-relevant INs and MNs. Left, heatmap depicting connectivity from SNs onto neurons directly presynaptic to TTMn. Connectivity is determined by a 1% input and 25 synapse threshold (groupwise). Rows are grouped by origin, with similar descriptions combined into larger categories (e.g. wing CS SNs and wing margin bristle SNs are both included in the “Wing SN” category). Columns from neurons of interest are labeled with arrows. Right, as above but for neurons one or two synapses upstream of the PSI (i.e., neurons on the right panel provide at least 1% of synaptic input to neurons on the left panel). **C.** Diagram showing strong synaptic connectivity between SNs, neurons of interest, and the TTMn (top) or PSI (bottom). **E.** Connectivity diagram of AN05B006 10603 and AN05B006 10221 (ANGABA),their strongest postsynaptic targets, and upstream DNs. In order to be included on this diagram, presynaptic neurons must be DNs that provide at least 1% of synaptic input to either AN05B006 10603 or AN05B006 10221. Postsynaptic neurons are curated from a set of neurons that receive at least 1% of their input and at least 150 synapses (groupwise) from either AN05B006 10603 or AN05B006 10221. DNs are labeled according to whether they synapse onto both ANGABA neurons and their downstream targets or solely onto one of the ANGABA neurons. **F.** Diagram demonstrating how actions important for both short-mode and long-mode takeoffs could be coordinated by DNs, local neurons in the VNC, and information from SNs.

**Figure 20–Supplement 1:**
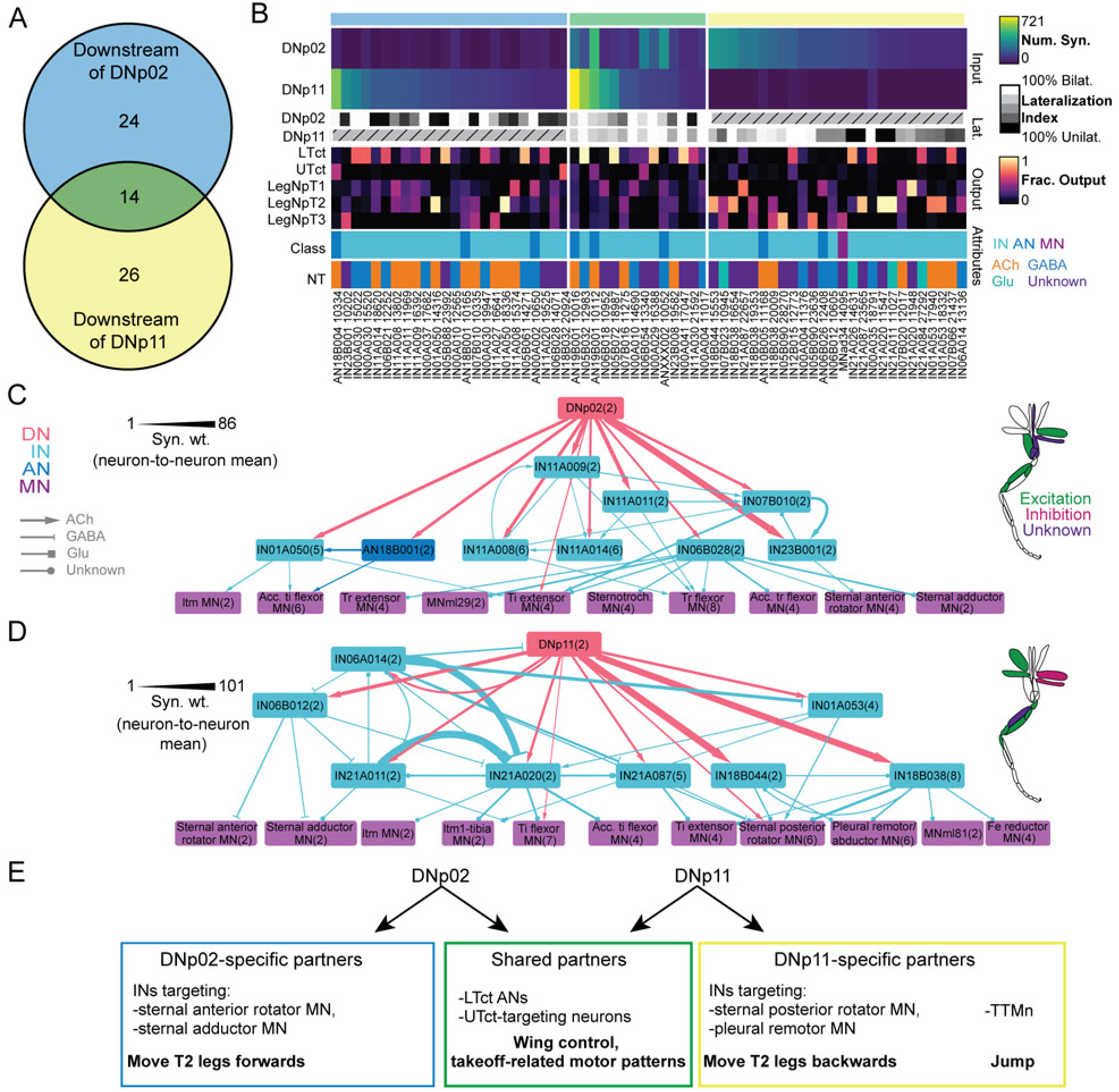
Circuits mediating pre-takeoff postural adjustments. **A.** Venn diagram depicting shared first-order downstream connectivity of DNp02 and DNp11 in the VNC. Numbers refer to the number of groups that receive more than 75 synapses and more than 0.5% of their input from either or both DNs. **B.** Attributes of neurons downstream of DNp02 and DNp11. Top heatmap (“Inp.”): number of synapses onto each group from DNp02 and DNp11. Top bar refers to the group’s assignment in A. All rows are sorted according to the number of synaptic inputs from DNp02 (for the neurons that receive input from DNp02) or from DNp11 (for the neurons that only receive DNp11 input). Second heatmap (“Lat”): Ipsi-contra lateralization index (as in Figure 11H), indicating how bilaterally distributed inputs from DNp02 and DNp11 are. Index was not computed for groups that did not meet the threshold for strong synaptic connectivity. Third heatmap (“Output”): fraction of each group’s output in the indicated VNC regions. Bottom heatmaps (“Attributes”): cell class and neurotransmitter prediction for each group, colored as in Figure 1. **C.** Connectivity between DNp02, all DNp02-exclusive downstream neurons from panels **A** and **B** that synapse onto a MN type in LegNpT2, and those downstream MNs. In order to be included in the diagram MNs must be in LegNpT2 and receive at least 20 synapses from all DNp11-downstream neurons. Downstream neurons must provide at least 5 synapses onto all MNs in this set. Right inset, a diagram of the putative cumulative effects of activating DNp02 on T2 leg muscles based on the connectivity of neurons described in this panel. **D.** Connectivity between DNp11, DNp11-exclusive downstream neurons, and LegNpT2 MNs as in **C**. **E.** Diagram summarizing the putative functional impact of activating synaptic partners downstream of DNp02, DNp11, or both DNs.

AN19B001_rostral (group 10016) provides strong input (≥1% of total synaptic input) to over 250 other neurons in the VNC, including wing MNs (MNwm36, tp2, b2, ps1, DVMn), intrinsic neurons with unilateral or bilateral outputs in single leg neuropils (e.g., types IN06B029, IN21A020, IN01A026), intrinsic neurons that span multiple neuropils (e.g. type IN00A002), and abdominal MNs (Figure 18G, Figure 20A). The intrinsic neuron that receives the most input from AN19B001_rostral, IN19B008, projects to multiple wing MNs including the hg1, hg3, hg4, b2, iii4, and ps1 MNs. Many of the muscles downstream of MNs targeted directly by AN19B001_rostral are involved in escape; for example, the tp2, b2, and DVM muscles are implicated in the processes of wing raising and flight initiation (Dickinson and Tu, 1997; O’Sullivan et al., 2018). AN19B001_rostral synapses extensively on IN06B066 (which, as discussed in section 2.5.3, inhibits the DLMns and, to a much lesser extent, the DVMns). Based on these observations, we speculate that one of the functions of AN19B001_rostral might be to suppress wing depression while directly activating neurons involved in wing raising.

In addition to a more caudal dendritic arborization, we observed that AN19B001_caudal (group 10112) distinguished itself from AN19B001_rostral with a more extensive axonal arborization in the UTct (Figure 17G). This group has strong outputs to wing MNs of multiple classes, including the ps1 MNs, MNwm36, the b2 MNs, and both the DVMns and DLMns. In addition, AN19B001_caudal provides input to intrinsic neurons that themselves directly contact wing MNs including the Tect INs (see section 2.5.3). Its direct and indirect outputs to the ps1 MN and both sets of wing power MNs suggest that AN19B001_caudal might be involved in initiating the flight motor system after takeoff. Unlike AN19B001_rostral, the caudal group does not make any direct excitatory contacts with the tp2 MN and instead synapses strongly onto IN11B013, a set of GABAergic interneurons presynaptic to both the DVMn and the tp2 MNs (Figure 20A). This differential connectivity suggests that, despite their morphological similarities, AN19B001_caudal and AN19B001_rostral are likely to control distinct wing motor outputs.

AN18B001 devotes most of its axonal projections to the LTct and the leg neuropils (Figure 18F). In the LTct, AN18B001 contacts three intrinsic neuron groups, IN07B023, IN21A027, and IN21A026, which themselves output to the TTMn both directly and indirectly. All of these cells are predicted to be glutamatergic, meaning that they could be either excitatory or inhibitory. However, there is good reason to believe that this network has an excitatory influence on TTMn: DNp11 neurons, which are cholinergic and evoke jumping when activated (Dombrovski et al., 2023), make at least 100 synapses onto two of these three neuron groups (representing the most direct connection between DNp11 and TTMn). Several other DNs (DNb09, DNxn174) make strong outputs onto two or more of these neuron groups, and also presumably initiate jumping when activated.

It is still unclear how CNS circuits control the timing of the motor sequences critical to long-mode takeoffs. Two models could explain this action sequencing: either 1) sets of DNs are dedicated to triggering distinct VNC circuits that govern each of these actions and are activated sequentially in the brain, or 2) “long-mode” DNs collectively activate multiple VNC neurons governing takeoff; VNC-intrinsic circuits (including both lateral interactions between descending circuits and ascending sensory feedback) then instruct transitions from one phase of the takeoff sequence to the next.

To substantiate the first model, it would be necessary to identify individual DNs that receive input from looming-sensitive neurons in the brain and initiate only a single action from the suite of motions necessary for takeoff. These DNs would have substantial synaptic influence on the motor pathways associated with one action, but not others. We did identify some DNs which may fit these criteria. For instance, DNb09 and DNxn174 target neurons presynaptic to TTMn without synapsing onto neurons presynaptic to wing MNs (see above) and would presumably initiate jumping with limited wing movement when activated on their own (Figure 20A). Since it is uncommon for *Drosophila* to leap without any wing movement in response to looming stimuli, these DNs may be activated in coordination with other DNs responsible for initiating the other actions associated with long-mode takeoffs.

Notably, however, the majority of LTct-DNs that contribute to long-mode takeoffs exhibit efferent connectivity patterns likely to influence activity in multiple escape-related effectors including the direct wing MNs, the indirect wing MNs, and the TTMn (Figure 20A). For example, DNp02, DNp04, DNp06, and DNp11 have secondary and tertiary synaptic connectivity to all of the MNs listed above. Furthermore, flies that take off after optogenetic activation of these DNs perform full long-mode takeoff sequences (Dombrovski et al., 2023) rather than individual actions.

In support of a model in which local interactions within the VNC pattern instruct the sequencing of takeoff behavior, we discovered evidence that sensory neurons from the wings and legs synapse onto intrinsic neurons presynaptic to escape effectors (Figure 20B, C). This suggests that sensory input might influence the progression of activity in VNC escape circuits. SNs from the notum and the wing, including campaniform sensilla neurons which could detect wing deformation, synapse onto IN13A022 (Figure 20C, D). These INs have direct synaptic input to the TTMn and also output to several other intrinsic neuron groups presynaptic to the “jump” MN, including the GFC2 neurons (Figure 20D). In addition to receiving input from the GF, GFC2 neurons receive descending inputs from other looming-sensitive DNs (such as DNp06) and other neurons associated with takeoffs (such as AN19B001_caudal, Figure 20A) as well as centripetal inputs from T2 leg SNs (Figure 20B, C).

The GFC2 cells are not the only neurons implicated in takeoff initiation that receive strong afferent inputs from both SNs and DNs. AN19B001_caudal, for example, receives extensive monosynaptic inputs from wing SNs. The PSI receives both disynaptic and trisynaptic input from SNs via several groups of INs (Figure 20B, C). While some of these INs are sensitive to only a single sensory modality, others integrate input from multiple types of SNs, or from groups of INs that receive input from SN types spanning two or more modalities. We hypothesize that AN19B001_caudal and neurons presynaptic to the PSI and TTMn serve as sites for the convergence of descending and ascending information streams, and that these two types of inputs may interact with one another (Figure 20F). For example, descending input may modulate ascending sensory pathways, rendering takeoff-related motor effectors more sensitive to cues such as wing raising after activation by LTct-DNs.

What is the role of this sensory information? Some sensory pathways might bypass the central brain to trigger escape circuits directly, minimizing response latency to threats that are close enough to touch the fly. These sensory streams may also be tuned to sensations important to checkpoints in the takeoff process; for example, sensory inputs from the T2 leg could detect the initiation of a jump and trigger activity in the PSI, initiating a downstroke. Wing campaniform sensilla SNs could also detect wing deformation and delay downstroke or jumping until the appropriate time through the polysynaptic pathways described above. Consistent with this hypothesis, we have previously observed that the timing of certain motor subprograms within the long-mode takeoff sequences are always tightly coupled. For example, the initiation of the jump and the downstroke almost always occur with very short latency after the completion of wing raising ((von Reyn et al., 2014); unpublished data). We remain unable to predict the neurotransmitter expressed by many neuron types in these circuits. However, it is likely that sensory information would evoke activity in both excitatory and inhibitory pathways to effectively gate the rigidly-timed activation of escape motor effectors.

A number of LTct neurons which are likely to be involved in coordinating takeoffs, including the set of ANs that integrate synaptic input from looming-sensitive LTct-DNs, are also governed by AN05B006, a set of GABAergic ANs that can also be subdivided into rostral and caudal groups with overlapping efferent and afferent connectivity. Both groups target AN08B098, AN19B001_rostral, and AN18B001; AN05B006_caudal (group 10221) also synapses onto AN19B001_caudal and AN05B006_rostral (group 10603) inhibits the axon of the GF (Figure 20D, Figure 19C). The AN05B006s might serve either as a persistent “brake” to prevent any takeoffs in particular behavioral contexts or as a “switch” by which one behavioral program might laterally inhibit others that would normally evoke takeoffs (Figure 21C). Alternatively, these neurons might have a role in instructing normal takeoff sequences by providing temporally precise inhibition of takeoff-related behavioral modules. Consistent with these many possible roles, AN05B006s receive strong inputs from a diverse set of DNs, many of which will selectively target either the rostral or the caudal group (Figure 20D). This afferent connectivity includes both DNs that do and others that do not also synapse onto some of the targets of AN05B006 (Figure 20D).

**Figure 21:**
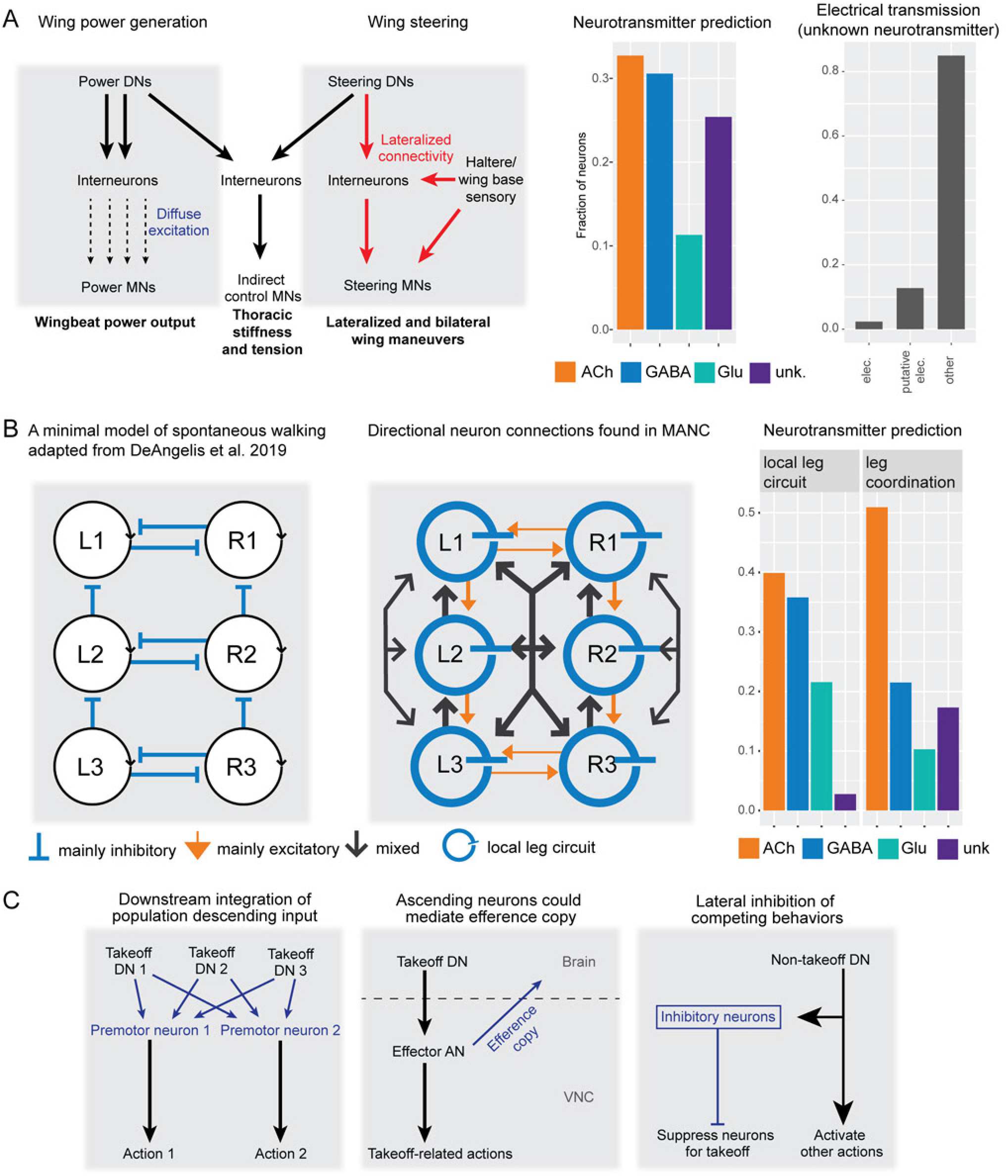
Summary of wing, leg and LTct motor control. **A.** Wing motor control summary. On the left, wing control circuits are segregated into power and steering circuits, which each also contribute to the activity of indirect control MNs. On the right, neurotransmitter predictions and electrical transmission assignments for neurons of VNC origin (except MNs) that are predicted to play a role in wing motor control (Infomap communities 8, 9, 10, 16). Neurotransmitter predictions are thresholded at ≥0.7 probability and assigned to the unknown (unk.) category if below threshold. Known or putative electrical transmission (from the ‘transmission’ neuprint field) are shown for neurons with unknown neurotransmitter prediction. **B.** Leg coordination summary. On the left, a schematic adapted from (DeAngelis et al., 2019) showing a potential minimal model to coordinate the 6 legs to allow tripod and tetrapod movement. In the center, a summary of the actual connections seen upstream of the leg premotor and motor circuits, summarized from Figure 14. Line is thicker if there are more than 5 pairs of neurons in that group of the US connectivity (see Figure 14). Glutamatergic neurotransmitter predictions are considered to be inhibitory for this schematic. On the right, comparison of neurotransmitter predictions for the restricted leg premotor neurons and the leg coordination neurons. **C.** Summary of LTct circuits that likely contribute to escape-related behavior. On the left, distinct DN types converge onto a set of motor effectors that each control overlapping facets of an escape behavior. In the middle, LTct-DN targets ascending neurons that likely play a dual role in actuating motor output and relaying motor information back to the brain (i.e. efference copy). On the right, a subset of DNs unrelated to the control of takeoffs activate inhibitory neurons that suppress takeoff-related actions, thus mediating descending action selection.

Some of the LTct-DNs that contribute to long-mode takeoffs have also been shown to actuate related defensive motor patterns. DNp02 and DNp11, for instance, initiate shifts in the positioning of the T2 legs that influence the direction that the fly jumps in response to a threat (Figure 18) (Card and Dickinson, 2008b; Dombrovski et al., 2023). Notably, they initiate opposing movements: DNp11 promotes a backwards shift of the T2 legs relative to the center of mass (increasing the likelihood of the forward takeoff) while DNp02 promotes a forward shift of these legs (increasing the likelihood of a backwards takeoff) (Dombrovski et al., 2023). Our findings described in section 2.7.1 suggest that this functional divergence might be explained by differential first order connectivity of DNp02 and DNp11 in the VNC.

Indeed, we observe that the majority of neurons downstream of DNp02 do not receive substantial input (here, ≥75 synapses and ≥ .5% input) from DNp11–and vice versa (Figure 20–Supplement 1A,B). Many of the neurons that receive this level of synaptic input from both DNs (n=14 groups) have outputs in the LTct, including several of the LTct AN “integrators” described above (Figure 18G, Figure 20–Supplement 1B). In contrast, neurons that receive strong input from only one of the DNs (n=24 groups from DNp02 and n=26 groups from DNp11) are more likely to output onto leg neuropils including LegNpT2 (Figure 20–Supplement 1B). We focused on a subset of cells that receive input from only one of the DN groups and also have direct output onto a T2 MN (Figure 20–Supplement 1C, D). These neurons could comprise hubs sufficient to coordinate positional shifts of the T2 legs. For instance, DNp11 synapses onto three putatively cholinergic neuron types (IN18B038, IN18B044, IN01A053) that all target T2 sternal posterior rotator MNs that function to rotate the coxa posteriorly. Two of these types also have ≥15 synapses onto the T2 pleural remotor MNs that might serve a similar role. At the same time, DNp11 also synapses onto IN06B012, a GABAergic IN that inhibits both T2 sternal anterior rotator MNs and sternal adductor MNs, neurons that actuate the opposing coxa motion. In contrast, one of the strongest outputs of DNp02 is onto the putatively cholinergic IN23B001 which directly synapses onto the sternal anterior rotator MN in T2. In addition, DNp02 outputs to IN06B028 which provides direct input to the sternal anterior rotator MN and the sternal adductor, among other leg MNs (Figure 20–Supplement 1C).

Differential first-order connectivity between DNp11 and DNp02 also accounts for why the former neuron is more likely to promote takeoffs when optogenetically activated: unlike DNp02, it connects directly to the previously-described trio of INs (IN07B023, IN21A027, and IN21A026) upstream of the TTMn (see above) (Dombrovski et al., 2023).

Flies do not make pre-jump postural shifts in just the rostrocaudal axis; indeed they can take off at any angle relative to their body axis (Card and Dickinson, 2008b). Flies could introduce a lateral bias in their takeoff direction through unilateral forward or backward preparatory movements of the T2 legs. Interestingly, DNp02 and DNp11 differ in how lateralized their outputs to the T2 leg neuropil are (Figure 20–Supplement 1B). For instance, IN23B001 receives nearly all of its input from the ipsilateral member of the DNp02 pair. In contrast, IN18B038 and IN18B044 each receive at least a third of their input from the contralateral member of the DNp11 pair. Intriguingly, DNp11 makes highly lateralized synapses onto IN06A014, a GABAergic IN that itself targets IN01A053, IN21A020, IN21A087, neurons that are each downstream of DNp11 and upstream of T2 MNs (Figure 20–Supplement 1D). This suggests that, unlike DNp02, DNp11 could rely on lateralized feedforward inhibition to produce unilateral movements of the T2 legs.

Experiments using constitutive inactivation and/or optogenetic activation of DNs suggest that, while activity in each of many different DNs is sufficient to initiate patterns of motor activity, very few DNs are singularly necessary for the expression of these behaviors (Ache et al., 2019; Cande et al., 2018; Guo et al., 2022; Rayshubskiy et al., 2020). This is true, as well, of DNp02 and DNp11: flies in which either neuron has been silenced perform normal pre-takeoff postural adjustments (Dombrovski et al., 2023). This implies that these DNs normally act as a population to collectively coordinate action, with each member making distinct contributions to the evoked movement. Consistent with these findings, it is possible to identify distinctions in the activation phenotypes of even closely related DNs (Ache et al., 2019). Our investigation of the downstream connectivity of DNp02 and DNp11 suggest that these distinctions are correlated with unique efferent connectivity in the VNC.

#### 2.7.4 Circuits for the flexible coordination of takeoffs

In summary, we identified synaptic pathways explaining how flies are able to perform coordinated movements of their wings and legs such as long-mode takeoffs (Figure 20E, 21C). We discovered that this coordination rarely emerges from monosynaptic connections between DNs and MNs, which would imply action sequencing occurs solely in the brain, but instead relies upon VNC-intrinsic nodes that have both a high input and a high output degree. These neurons are likely to integrate input from a population of DNs that are activated by similar kinds of threatening features (Figure 20A, 21C). They then synapse across multiple neuropils in the VNC to drive coordinated behaviors. Many of these neurons are ANs and, as such, also send projections back to the brain and/or premotor areas such as the sub-esophageal zone. A broad functional imaging survey of ∼250 AN types suggests that ANs commonly convey high-level information about a variety of behavioral states (Chen et al., 2023). Future connectomic volumes that include more of the central nervous system may elucidate how this efferent connectivity impacts escape-related behaviors. Notably, in *Drosophila* larvae, ascending projections have been shown to synapse in the brain onto DNs that may be activated sequentially in a behavioral sequence (Winding et al., 2023). It remains to be seen if these, or similar motifs, may play a role in the coordination of long-mode takeoffs.

Our analysis suggested that SNs, like ANs, play an outsized role in the control of behavior. Many VNC neurons, including a number that likely influence the activity of takeoff-critical MNs, received strong afferents from both sensory and descending input. We hypothesize that this convergence is critical for establishing precisely timed action sequences.

Finally, our analysis identifies VNC neurons that are poised to gate expression of takeoff behaviors. We identified an AN type, AN05B006, that broadly inhibits many neurons that we believe to be important for long-mode takeoffs. We also observed a number of INs that directly target the axon of the GF, presumably exerting a strong influence on the successful propagation of GF spikes (and, therefore, the initiation of short-mode takeoffs in response to threats) (Figure 19B-D). These inhibitory motifs may be the substrate for enforcing reciprocal inhibition between different action pathways, ensuring that only a single movement is carried out during action selection, or they may be elements required to maintain the correct timing and sequencing of behavioral submodules in a complex action sequence like takeoff. Future functional studies are required to test these hypotheses and further flesh out how connectivity patterns in the VNC contribute to the motor control of the fly’s wide range of behaviors.

## 3 Discussion

Within the last decade, the accelerated development of EM connectomics has greatly improved our ability to probe the organization and function of neural circuits. The densely-reconstructed Hemibrain connectome has allowed researchers to carry out large-scale and open-ended interrogation of *Drosophila* brain circuits. The release of the MANC dataset now permits a similar level of inquiry into nerve cord circuits. To begin describing and understanding the connectivity from brain to VNC and motor output, we have systematically reconstructed, cataloged and annotated all DNs and MNs of the VNC, and linked 29% of DNs and 55% of MNs to neurons identified at the light level. While the remaining DN and MN types in MANC are yet to be identified at the light level, the availability of EM morphologies will expedite neurobiologists’ attempts to target and characterize them through tools such as color depth MIP mask search (Otsuna et al., 2018) and NBLAST (Costa et al., 2016). Next, using the densely-reconstructed MANC connectivity (Takemura et al., 2024), as well as a rich layer of annotations and metadata (Marin et al., 2024), we examined descending input all the way to motor output connectivity broadly across the VNC as well as specifically in leg, wing, and lower tectulum neuropils and circuits.

Our annotation and reconstruction of neurons passing through the neck connective reveal a DN cell count of 1328, greater than prior estimates that range from 700 to 1100 cells (Hsu and Bhandawat, 2016; Namiki et al., 2018). In comparison, the AN cell count of 1864 is only about 1.5 times greater, highlighting the bidirectional nature of information flow between the brain and VNC. We also found and annotated 737 MNs in MANC (roughly half of the DN count), setting an upper limit on the number of motor units found across the fly’s thoracic and abdominal motor systems. Overall, direct DN input to most MNs remains fairly low (<10% of their input synapses), although roughly 80% of MN groups receive direct input from at least one DN group at a 1% input threshold. There are a few major outliers to this pattern. Most prominently, DN input contributes up to 60% of the total upstream synapse count of the neck MNs and a subset of abdominal MNs. As observed previously (Namiki et al., 2018), DNs are divided anatomically into those with axons terminating in the dorsal neuropils (Upper Tectulum and Intermediate Tectulum, subclasses DNnt, DNwt, DNht, DNut, DNit encompassing ∼400 DNs), those with axons terminating in the ventral neuropils (Leg Neuropils, subclasses DNfl, DNml, DNhl, DNxl encompassing ∼400 DNs), or those targeting the most intermediate layer, the Lower Tectulum (subclass DNlt, encompassing 32 DNs). Our proposed systematic nomenclature for the DNs was based on these clear divisions. A further set of DNs (DNxn, ∼500 DNs) target multiple combinations of the above regions and may be involved in control of multiple motor systems simultaneously.

To dissect the structure of VNC intrinsic networks, we used several methods to examine both large- and fine-scale connectivity in these networks. We used the Infomap algorithm to partition VNC circuits into subsets of highly interconnected neurons called “communities”. This partitioning suggests that VNC premotor circuits have a modular structure, with individual communities dedicated to controlling specific sets of MNs. Leg neuropil-restricted neurons largely form a single community in each hemisegment, and the serially-repeating structure of leg control circuits allowed us to generate a single ‘standard leg connectome’ representing connectivity of these restricted neurons in each leg hemisegment. In contrast, upper tectular motor systems are more mixed, with several communities controlling both wing and haltere MNs, and an additional single community controlling neck muscles. Interestingly, Infomap analysis also identified three large communities with low direct MN output—neurons of these communities commonly have input and output in the intermediate neuropils but also often cross neuropil boundaries to arborize widely in the VNC. These communities may thus serve as ‘higher order’ behavioral control centers through their connectivity with other communities, and two of these communities are highly associated with DNs targeting the lower tectulum.

Leg control networks are hierarchically organized, with each leg neuropil containing a local circuit with extensive intra-neuropil connectivity, upon which an inter-neuropil circuit is layered (Figure 21B, middle). For each leg, the large number of intrinsic neurons (∼1200) dedicated to single leg neuropils (Marin et al., 2024) reflects the complexity of a potential leg central pattern generator (CPG) that must control and coordinate muscles across the many segments of each individual leg. This complexity likely reflects the flexibility required for the fly to be able to carry out locomotor behaviors that adapt to varying environmental stimuli ranging from the unevenness of the ground to the different attack strategies of predators. These leg neuropil-intrinsic neurons included a large proportion of serially repeated neurons, consistent with the interpretation that these leg-intrinsic communities comprise a CPG motif that is repeated for each leg.

These intra-leg networks are connected by neurons that project between leg neuropils. Our analysis of these interconnecting neurons allows us to compare our putative premotor circuits to previous predictions for circuit control of six-legged insect walking (Figure 21B, left) (Bidaye et al., 2018; DeAngelis et al., 2019; Rosenbaum et al., 2015; Schilling and Cruse, 2023, 2020; Tóth and Daun, 2019). Based on these existing locomotor models, we expected that a large proportion of inhibitory neurons would control posterior-anterior and inter-segmental inhibition between and across distinct leg neuropils (one example shown in Figure 21B, left). Instead, we found that most inhibitory neurons target all leg neuropils across thoracic segments on one or both sides (Figure 21B, middle). Again contrary to previous predictions, we do not see strong bilateral connections between neurons in the T2 leg neuropils. While we found serial sets of neurons that project from the posterior to the anterior on the same side as predicted, we also found anterior to posterior projections which are exclusively excitatory. Indeed, the inter-leg circuit is more heavily cholinergic compared to the intra-leg circuit (Figure 21B, right). However, it is possible that some of the posterior-to-anterior connections have a net inhibitory effect by exciting inhibitory neurons that are presynaptic to leg MNs. Theories regarding the architecture of leg locomotor circuits must now be updated to reconcile with connectome data to account for the inter-leg neuropil connectivity observed, that for example show both posterior-to-anterior and anterior-to-posterior connections between leg neuropils, as well as relative lack of interaction between the left and right mesothoracic leg neuropils. Taken together, our results provide a platform for detailed investigation of potential inter- and intra-leg CPGs upstream of the leg motor neurons and the modulatory input of different DNs on these circuits. They also suggest that leg CPGs may primarily operate at the level of single legs and are synchronized through numerically much weaker inter-leg connections. One of these inter-leg connections is the wing grooming neuron, wPN1 (Zhang and Simpson, 2022), and we expect that several of the inter-leg coordination neurons are specific to a given leg behaviour. Indeed, while both intra- and inter-leg circuits receive DN input, most leg DNs act directly on the intra-leg circuits, which suggests that most leg descending input primarily controls intra-leg circuit computations which then secondarily affect inter-leg coordination. Future work will be able to combine this leg motor circuit with the thorough sensory neuron annotations provided in our companion paper (Marin et al., 2024)

The organization of the dorsal neuronal communities putatively involved in wing, haltere, and neck control is very different to that of the leg networks. The upper tectulum neuropil is segmentally specialized anterior-to-posterior for control of the neck, wing and haltere motor systems respectively. The wing control system that likely plays a role in flight is further functionally separated into two networks: one largely dedicated to the control of the power muscles, the other to control of the steering muscles (Figure 21A, left). Each of these networks differ in their synaptic organization onto MNs. The power-muscle circuits provide input to MNs that are more bilateral and often ‘diffuse’ (individually weak), which may be suited to generally drive power MNs. This observation is consistent with prior theory that power MNs summate many weak excitatory inputs to drive staggered spiking (Harcombe and Wyman, 1977). In contrast, steering circuits more commonly have stronger, lateralized inputs that are suited to asymmetrically control steering MNs. Descending signals converge with oscillatory sensory input from wing base and haltere sensilla to achieve steering while maintaining phase-locked spike timing of steering muscles relative to the wingbeat cycle, consistent with prior observations (Fayyazuddin and Dickinson, 1999, 1996; Trimarchi and Murphey, 1997). The steering network also appears intimately associated with haltere motor control, and both networks further have shared control of the wing indirect control muscles. A third circuit contains most of the identified *fruitless*-positive neurons in tectular regions(Lillvis et al., 2024), and may be implicated in courtship. DNs that provide input to wing circuits appear largely specialized in their connectivity to one of these three circuits, suggesting that motor control for different stereotyped actions such as flying or singing is controlled by largely parallel motor networks that are already separated at level of DN command. Future studies should investigate whether this functional specialization of wing motor control extends to the brain as well.

Neurotransmitter distribution across neurons in the wing control circuits is similar to that of the leg restricted circuits, albeit with a higher proportion of unknown neurotransmitter predictions. Our analysis suggests, however, that wing control circuits likely further incorporate a number of electrically-coupled interneurons (Figure 21A, right), which may be advantageous to a motor system demanding rapid, repeated muscle activation. Although the connectome does not allow direct assessment of electrical connectivity, efforts were made to identify putative electrical neurons throughout the VNC (Marin et al., 2024). Importantly, putative electrical neurons appear common in the upper tectulum, and some are also found in leg neuropils. Indeed, prior findings show that the innexin ShakB (the main gap junction protein in neurons) is strongly expressed in the medial VNC, haltere afferent tracts and sections of the leg neuropils (Ammer et al., 2022). The prevalence of electrical synapses will pose a challenge for interpreting circuit function, requiring additional work to characterize electrical synapse connectivity in the VNC. For example, serial-section transmission EM in the CNS combined with immunogold labeling (Ando et al., 2016; Shahidi et al., 2015) potentially allows labeling of gap junction proteins in CNS EM volumes for the creation of an ‘electrical synapse connectome’.

In the wing steering system, many of the steering muscles that control a given sclerite of the wing hinge share common tendons and may act as motor units to differentially control force exertion, while some others with separate tendons can exert similar action on a sclerite (Dickinson and Tu, 1997). Indeed, prior work shows differential recruitment of muscles per sclerite depending on the magnitude of turning during flight, reflecting control of force generation for executing wing kinematic changes (Lindsay et al., 2017). This is partially borne out in the connectome: shared upstream connectivity is observed for some (but not all) MNs for muscles of the same sclerite. For example, we find that the tonically active b1 MN and the phasically active b2 MN partially share upstream DN input and electrical input from w-cHINs, upon which DNs and haltere campaniform afferents converge. It is likely that depending on connectivity strength and activation threshold of the MNs, the same descending steering signals may shift the activation phase of the b1 MN in the wingbeat cycle during lower magnitude turns, and further activate the b2 MN during higher magnitude turns.

Both the leg and wing motor systems utilize circuits that are relatively directly associated with MNs controlling a single limb or body part. In contrast, DNs that innervate the Lower Tectulum (LTct) target Infomap communities that may play a role in higher-order motor control by coordinating actions across the different motor systems. For example, a significant fraction of these “LTct-DNs” are implicated in controlling escape takeoff, which involves both wing raising and rapid leg extension. We examined connectivity within these circuits to identify motifs that might play a role in mediating takeoff motor actions, as well as action selection between different modes of escape takeoff. We learned that the outputs of looming-sensitive LTct-DNs are pooled by a common set of broadly-projecting integrator neurons, each of which provides output to distinct leg and wing motor circuits (Figure 21C, left). This schema suggests an explanation for the observed degeneracy of many of the individual DNs associated with the mode of takeoff involving wing-raising: each of these DNs can activate the same takeoff-essential integrators. Surprisingly, many of the LTct integrator neurons that synapse most broadly in the VNC are also ANs. Thus, these integrators may serve as local effectors of motor output while also sending an efference copy of these ongoing actions back to the brain (Figure 21C, middle) or triggering additional motor sequences through feedforward activation of central circuits.

Neurons with a high input and output degree, such as the LTct integrators identified above, are poised to serve as an important site of neural circuit regulation (Figure 21C, right). Inhibition of these cells might play a role in action selection by breaking the link between looming detection in the brain and the initiation of escape behaviors in the VNC. Interestingly, we discovered a type of putative inhibitory neurons (AN05B006) that receives input from DNs–both looming-sensitive DNs and others which are currently uncharacterized–and synapses directly onto multiple LTct integrators. We also identified related motifs controlling the output of the Giant Fiber DN, which is itself necessary for the initiation of an alternative type of escape behavior (the short mode takeoff, see Figure 19) (von Reyn et al., 2014; Wyman et al., 1984). The local VNC input to the GF includes many putative inhibitory neurons that themselves receive strong input from other DNs. This connectivity raises the hypothesis that these other DNs employ this circuit to modulate GF excitability and engage directly in action selection. These examples illustrate how VNC circuits could instruct behavioral choices by integrating local computations with descending information. Indeed, while DNs are generally thought of as sources of information from the brain, most MANC DNs receive significant axonal synapse inputs at a mean of ∼11:1 output-to-input ratio, suggesting that local VNC circuits commonly play an active role in the regulation of DN output.

Our analyses give readers a broad overview of DN-to-MN connectivity and premotor pathways in the VNC as well as specific details on a few key circuits. However, it is not yet a comprehensive interpretation of all VNC connectivity. The scale of the connectome means that large-scale analyses do not always capture nuances of connectivity, while small-scale circuit examination still remains quite manual. In addition, functional data is still lacking for most VNC cells, which is vital to circuit interpretation. A key future goal is to combine connectomics and experimental approaches for meaningful interpretation of circuits across the VNC. Many of our analyses focused on DNs with available functional data, since this enabled us to carry out more meaningful circuit analysis. Encouragingly, this often allowed us to predict the function of unstudied DNs based on their participation in the same circuits.

There are of course numerous ways in which our study (in common with others analyzing EM connectomes) will be extended in future. Our analyses do not take into account non-synaptic neurotransmitter and neuromodulator release, differences in neuron physiology, and also make simplifying assumptions about the equivalence of synapse counts and synapse weights. Thus, we emphasize that our current analyses of VNC-wide premotor circuits are very much a starting framework both to aid future hypothesis-driven experimental exploration of VNC function, and to help define further connectome analysis.

In summary, the *Drosophila* VNC receives a plethora of descending signals from the brain, and VNC circuits serve to generate patterned motor output, coordinate between different motor systems and likely mediate behavioral choices. Our work here in proofreading, curating and identifying DNs and MNs of the VNC lays the groundwork to enable neurobiologists to easily find and examine descending-to-motor pathways of interest. The comprehensive access to VNC connectivity as well as the wealth of neuronal annotations and metadata now empowers neurobiologists to explore VNC circuits and unravel their secrets.

## Acknowledgements

This collaborative project was jointly initiated and supervised by GMC and GSXEJ. The Card lab was primarily responsible for analysis of the wing and lower tectulum circuits. The Jefferis lab was primarily responsible for light level matching, typing and analysis of leg circuits. We thank the following people for their contributions to this work: Feng Li, Emily Tenshaw, Julie Kovalyak, Christopher Dunne, Griffin Badalamente, and Marina Gkantia for contributions to focused MN or pre-MN proofreading. Erica Ehrhardt for pre-publication sharing of wing and haltere MN light-level data and type information, and discussion/verification of their MANC matches, and Joshua Lillvis for help with wing MN identification. Joshua Lillvis and Kaiyu Wang for pre-publication access to fruitless-positive neuron identifications in MANC. Jonathan Enriquez for providing image stacks of leg MNs and expert evaluation of leg MN identification. Tony Azevedo, John Tuthill, Wei-Chung Lee for generous pre-publication sharing and discussion of T1 leg MN identities and nomenclature in FANC. Ellen Lesser for cross-volume corroboration of wing MN identifications with FANC. Mark Eddison, Yisheng He, Jenn Jeter, Gudrun Ihrke, and the Janelia Project Technical Resources team for EASI-FISH. FlyLight team and Alyson Petruncio for standard neurotransmitter FISH work, and Jennifer Jeter and Christina Christoforou for FISH imaging. Janelia FlyLight Project Team and Janelia Project Technical Resources for screening split-Gal4 lines for wing MNs and DNs. Hideo Otsuna for color maximum intensity projections of all MANC neurons and making available EM color MIP match search of MANC neurons on the neuronbridge preprint server. This work was supported by Wellcome Trust collaborative awards 220343/Z/20/Z and 221300/Z/20/Z to GSXEJ and GMC, GM Rubin, S Waddell and M Landgraf; by the Howard Hughes Medical Institute (GMC, FlyEM project team); core support from the MRC (MC-U105188491) to GSXEJ; a Walter-Benjamin-Fellowship from Deutsche Forschungsgemeinschaft to TS (STU 793/2-1); NSF 2127379 to Kevin C. Daly and Andrew M. Dacks supporting the work of HSJC.

## Figure legends

**Supplementary file 1:** DN typing, tracts and light-level matching in MANC

**Supplementary file 2:** DN neurotransmitter FISH

**Supplementary file 3:** MN and efferent light-level matching in MANC

**Supplementary file 4:** MANC Infomap community assignments for intrinsic premotor neurons (all neurons with soma in the VNC except MNs)

**Supplementary file 5:** Putative electrical neuron groups by presynaptic site count over volume

**Supplementary file 6:** Serial Leg MN groups

**Supplementary file 7:** Standard Leg groups

**Supplementary file 8:** Leg Coordination groups

## Materials & Methods

### EASI-FISH protocol

Expansion-Assisted Iterative FISH (EASI-FISH) (Wang et al., 2021) was used for probing neurotransmitter-related genes with improved spatial resolution and more sensitive detection of transcripts compared to standard FISH techniques. Briefly, flies of DN split-Gal4 lines were crossed to UAS-CsChrimson-mVenus. Brains of F1 offspring were prepared for EASI-FISH with GFP antibody detection of Gal4-labeled DNs as described in (Eddison, 2022). For each single brain (1-3 total brains per split-Gal4 line), EASI-FISH was carried out for pairs of neurotransmitter-related genes (*ChAT, Gad1, VGlut, Tbh, Tdc2, SerT*, marking the neurotransmitters ACh, GABA, Glu, OA, OA/TA, and 5-HT respectively). The combinations of gene pairs probed per line were selected based on predicted neurotransmitter identity in MANC: for DNs with high prediction probabilities (≥0.7) for one of the three primary neurotransmitters (ACh, GABA, Glut), the probe combination covered the likely positive neurotransmitter and another primary neurotransmitter as a negative control. DNs of interest or with low primary neurotransmitter predictions were probed for genes for more minor neurotransmitters. A set of brains for DNp20, DNp24, DNd02, DNg34 and DNp27 that were probed for Tbh and Gad1 were further stripped using the protocol described in (Eddison, 2022) and reprobed for ChAT and VGlut. Neurotransmitter usage was determined based on all DN cells of the given type labeled in each brain (a pair or more), unless noted in Supplementary file 2.

### Standard FISH protocol

Standardized neurotransmitter FISH was carried out as previously described (Meissner et al., 2019) with neurotransmitter probes from one of two possible sets: *ChAT, Gad1*, and *VGlut* for ACh, GABA and Glu respectively, and *Tbh, SerT* and *pale* for OA, 5-HT and DA respectively. For each DN split-Gal4 line, 2-3 CNSes were probed for a given combination of neurotransmitter probes. Imaging was carried out on a Zeiss LSM 880 confocal microscope with a 63x/1.40 Oil Plan Apochromat DIC M27 objective with laser excitation at 488, 561, 594 and 633 nm.

### CNS immunohistochemistry

Flies were anesthetized on ice and dissected in ice-cold PBS. Isolated CNSes were fixed in 4% formaldehyde in PBS with 0.5% Triton X-100 (PBST) at room temperature for 30 minutes. After washing with PBST, CNSes were incubated with anti-GFP sheep polyclonal antibody (3:5000, AbD Serotec, #4745-1051) and nc82 mouse monoclonal antibody (1:50, DSHB) in 5% normal donkey serum in PBST for 2 days at 4°C, then following another set of washes in PBST, Dylight 488-conjugated anti-sheep donkey antibody (3:5000, Jackson ImmunoResearch, #713-485-147) and Dylight 649-conjugated anti-mouse donkey antibody (1:250, Jackson ImmunoResearch, #715-495-150) for 2 days. Following washing, CNSes were mounted on poly-L-lysine-coated coverslips in Vectashield antifade reagent (Vector Laboratories, H-1000-10) and imaged on a Zeiss 980 confocal microscope with a 20x/1.0 W Plan-Apochromat DIC M27 75mm objective with laser excitation at 488 and 639 nm.

### Thoracic muscle staining and immunohistochemistry

Flies were anesthetized on ice and briefly washed with 70% ethanol. Thoraces were isolated under 2% paraformaldehyde/PBS/0.1% Triton X-100 and fixed in this solution overnight at 4°C. After washing in PBS containing 1% Triton X-100, the samples were embedded in 7% agarose and sectioned on a Leica Vibratome (VT1000s) sagitally at 0.2 mm. The slices were incubated in PBS with 1% Triton X-100, containing Texas Red-X Phalloidin (1:50, Life Technologies #T7471) and anti-GFP rabbit polyclonal antibodies (1:1000, Thermo Fisher, #A10262) and a chitin-binding dye Calcofluor White (0.1 mg/ml, Sigma-Aldrich #F3543-1G) at room temperature with agitation for 2 days. After a series of four ∼1h-long washes in PBS containing surfactants, the sections were incubated for another 24h in the above buffer containing secondary antibodies (1:1000, goat anti-rabbit, Thermo Fisher #A32731). The samples were then washed in PBS/1% Triton X-100 and fixed for 4 hours in 2% paraformaldehyde to reduce leaching of bound phalloidin from muscles during the subsequent ethanol dehydration step. To avoid artifacts caused by osmotic shrinkage of soft tissue, samples were gradually dehydrated in glycerol (2-80%) and then ethanol (20-100%) (Ott, 2008) and mounted in methyl salicylate for imaging.

Serial optical sections were obtained at 1 µm intervals on a Zeiss 980 confocal microscope with a LD-LCI 25x/0.8 NA objective, or at 0.5 µm with a Plan-Apochromat 40x/0.8 NA objective. Calcofluor White, Anti-GFP/anti-rabbit Alexa 488 antibodies and Texas Red phalloidin-treated samples were imaged using 405, 488 and 594 nm lasers, respectively. Images were processed in Fiji (http://fiji.sc/), Icy (http://icy.bioimageanalysis.org/) and Photoshop (Adobe Systems Inc.).

### Identification of descending neurons

We chose a perpendicular plane through the neck connective posterior to the cervical nerve and anterior to the prothoracic neuropil entrance ((Power, 1948); z = 67070) and annotated every neuronal profile passing through this plane. We then systematically proofread the automatic segmentation as described in (Scheffer et al., 2020) and (Takemura et al., 2024) with focus on the descending neurons (no soma in VNC). DNs were reconstructed by a human proofreader to capture the full extent of their axonic branches in the VNC. We then mirrored the left hand side neurons passing through the neck connective by transforming them through a symmetrized version of the dataset using the <u>malevnc</u> <u>symmetric_manc</u> function. We then ran NBLAST clustering (Costa et al., 2016) analysis on the mirrored left hand side and non-mirrored right hand side neurons. This allowed for systematic high quality proofreading and grouping of left-right homologous pairs. Many left/right neuron pairs had unique morphological characteristics and branching patterns were grouped into a pair of a single neuron on each side and given a unique DN type. However in some instances larger groups containing multiple neurons with similar morphology were combined into groups containing more than one neuron per side (DN populations). In these cases, if the DN populations with similar morphology had different connectivity, then they were split into different DN types.

To match DNs to light-level identifications, we obtained segmentations of DN Gal4 expression patterns (Namiki et al., 2018) and transformed them as described below for tract meshes into MANC space. Overlaying EM reconstructed DNs with the LM segmentations aided the identification of previously described DNs. We further used NeuronBridge (Clements et al., 2022; Meissner et al., 2023) to match EM reconstructed DNs to Janelia’s Gal4 and Split-Gal4 collection. For every match, we indicated the line and slide code used for the matching and employed a confidence system of 1 to 5 from worst to best match (see Supplementary file 1). Curation of prior reported DN split-Gal4 lines revealed that the line SS02542 does not label DNb01 as previously reported (Namiki et al., 2018), but a new DN with high morphological similarity. This new DN is thus named DNb09. To further identify previously published DNs from other studies, we queried virtualflybrain.org for the term “descending” with filters “adult” and “neuron”. We then identified DNs with available VNC LM morphology in MANC. This process is not exhaustive and thus we could have missed previously published canonical types. We kindly ask the *Drosophila* community to contact the authors with missing identifications.

### Tract identification

We obtained the VNC longitudinal tract meshes generated by (Court et al., 2020) from https://www.virtualflybrain.org/ (Court et al., 2023) (ids: VFB_00104642, VFB_00104635, VFB_00104636, VFB_00104637, VFB_00104634, VFB_00104638, VFB_00104633) and transformed them from their original JRC2018VNC_U space into MANC space. We then simplified all DN axon skeletons to their longest neurite starting from their VNC entrance in the neck connective and NBLAST clustered them. Through manual review of the resulting clusters and checking for overlap with the longitudinal tracts from (Court et al., 2020) we assigned tract membership to all DNs. In accordance with the definitions in (Namiki et al., 2018), short DNs that terminated in T1 or did not join a tract for notable distance were not assigned a tract membership. We choose the tract nomenclature from (Namiki et al., 2018) and include a table below of tracts identified in this paper compared to prior identifications and their nomenclature (Table 2). Overlaying the longest neurite skeletons onto the EM images further helped the assignment of the tracts (Figure 2A, C). We found that DMT from (Court et al., 2020) does not overlap with type-I MTD from (Namiki et al., 2018) and thus kept them separate with their original published name. Additionally, we identified two smaller tracts that were previously undescribed (type-III MTD and CVL). While type-I MTD takes a ventral and type-II a dorsal route, type-III originates in type-II, leaving the tract between T1 and T2 to run more ventral and lateral. The CVL tract starts in the neck connective ventro-medially and then curves around the T1 leg neuromeres ventrally towards the lateral posterior end of T1, it then continues medially followed by dorsally in T2, repeating the same pattern in T3 and the abdomen neuromeres.

**Table 2:** VNC tracts according to Court et al., 2020 and Namiki et al., 2018.

| This study | VFB_id | Court et al. 2020 | Namiki et al. 2018 |
| --- | --- | --- | --- |
| MDA | VFB_00104636 | MDT | MDA |
| DLT | VFB_00104633 | DLT | DLT |
| DMT | VFB_00104635 | DMT |  |
| type-I MTD |  |  | type-I MTD |
| type-II MTD |  |  | type-II MTD |
| type-III MTD |  |  |  |
| ITD | VFB_00104638 | ITD | ITD |
| VLT | VFB_00104637 | VLT | VLT |
| DLV | VFB_00104634 | DLV | DLV |
| VTV | VFB_00104642 | VTV | VTV |
| CVL |  |  |  |

We then created new tract meshes for MANC based on the neurons identified for each tract. First, we choose three longest neurites of DNs in a given tract that represented the tract structure the best. We then subsetted all longest neurites to those three for each tract, keeping all points with a maximum distance of 10µm. We calculated the 3D α-shape from the xyz coordinates of the remaining points (R, alphashape3d package) and smoothed the meshes using Blender version 2.82. The tract meshes are available in the malevnc R package (https://natverse.github.io/malevnc).

### Identification of motor neurons

Motor neurons and other neurons that enter or exit through VNC nerves were identified by direct annotation of bodies passing through nerves, or for certain MNs with degradation issues, by morphological identification or matching to MNs on opposite side of VNC, or by assessment of input/output ratio (assuming MNs have high input and low output synapse count). The initial segmentation for some MNs were considered poorer due to MN degradation issues. Thus, a limited resegmentation of MNs was carried out by retraining the flood-filling network model used for EM segmentation (Januszewski et al., 2018; Takemura et al., 2024) with additional ground truth from six MN reconstructions with human-curated merges. When leg MNs could not be fully traced out through the leg nerves due to axonal damage or degradation, exit nerve was putatively annotated based on MNs in the same morphological group. For all MNs, a team of trained proofreaders carried out proofreading and reconstruction, as well as group assignments by manual assessment of morphology and NBLAST matching across the midline. Groups were then refined by serial homology matching by connectivity and morphology for leg MNs.

During cell class assignment, we distinguished between MNs and a more generic ‘efferent’ class based on several criteria: cells are considered MNs if cell morphology and muscle target is known in the literature; otherwise, cells were putatively classified as MNs if they send a single axon out of the VNC via a single nerve, and were considered efferents if they send multiple projections out of the VNC or have known or suspected neuromodulatory roles. The latter includes the ventral unpaired medians, which leaves the VNC via multiple nerves to innervate muscles on bilateral sides of the body and play a neuromodulatory role (Coggshall, 1978; Ehrhardt et al., 2023; Schlurmann and Hausen, 2003), and we note their identity wherever possible (Supplementary file 3).

To identify MNs across the VNC, we primarily matched MNs to light-level data from (Ehrhardt et al., 2023) for wing MNs, (Dickerson et al., 2019; Ehrhardt et al., 2023) for haltere MNs, and (Enriquez et al., 2018) for leg MNs. Secondarily, we cross-checked MN matches by a plethora of light-level imagery and descriptions available in the literature (Bacon and Strausfeld, 1986; Dickerson et al., 2019; Ehrhardt et al., 2023; King and Wyman, 1980; Namiki et al., 2018; O’Sullivan et al., 2018; Schlurmann and Hausen, 2007; Sun and Wyman, 1997; Trimarchi and Schneiderman, 1994). For wing MNs, we further identified the hg4, b3 and ps2 MNs from new or published split-Gal4 lines (Figure 6–Supplement 1) (Sterne et al., 2021; Wu et al., 2016). For the haltere hi2 and hiii2 MNs, three pairs of MNs were present in the split-Gal4 line SS36076 (Dickerson et al., 2019), which innervates the hi2 and hiii2 muscles, of which two near-identical pairs matched the reported hi2 MN morphology (Ehrhardt et al., 2023). It is presumed that the anatomically distinct MN pair in this line is the hiii2 MN, however it is also possible that the hiii2 MN is one of the two hi2 MN pair matches. Furthermore, two MN types targeting the hb1 and hb2 muscles are present in the SS47195 line (Ehrhardt et al., 2023); however, the muscle targets of each MN type have not been distinguished and they thus retain the systematic types MNhm42 and MNhm43. Confidence for all matches were rated on a scale of 1-5, where matches scored 3 and below are putative. In addition, for the T1 leg MNs, we referred to a recent description of all T1 leg MNs at both the light and EM levels (Azevedo et al., 2024). Wing MN identifications were corroborated with a parallel identification effort in FANC (Azevedo et al., 2024). Supplementary file 3 shows the bodyids, type, systematic type, light-level matches and match confidence (if known in literature) for all MNs and efferents found within the MANC volume.

### Motor neuron serial homology matching

To identify T2 and T3 leg MNs that are serially homologous to T1 leg MNs, we initially transformed the T1 leg MNs from the Female Adult Nerve Cord (FANC) EM volume (Azevedo et al., 2024) to MANC space and performed an NBLAST comparison between MNs from the two VNC datasets (Costa et al., 2016). With expert visual evaluation, this allowed us to match up all FANC-identified T1 leg MNs to the MANC dataset. As efforts were made to match serially-repeating homologs of other neuron classes across segments in MANC (Marin et al., 2024), we reasoned that this data could be leveraged to identify serial homologs of the T1 leg MNs in the T2 and T3 segments, by comparison of MN connectivity received from identified serially homologous neurons across the 3 thoracic segments, as well as connectivity from neurons that reach across several thoracic segments (mainly DNs, ascending neurons and intrinsic neurons). This method allowed us to transfer the MN identification from 198 MN of the front legs to the middle and hind leg MNs. The dendritic morphology of the MNs identified by connectivity is highly stereotyped across the three leg neuromeres and their dendritic fields occupy similar yet slightly rotated areas in T1, T2 and T3. MNs that target the same type of muscle but in different leg compartments, such as the long tendon muscle (ltm) in the femur and the tibia, differ slightly also in their dendritic morphology in the VNC. Here, one difference observed is two dendritic branches extending towards the midline for MNs innervating the femur and a single medial branch for MNs innervating the tibia ltm (Figure 7B on the left). However, there are also differences between leg MNs that innervate the same leg muscle target, for example the previously described slow and fast tibia extensor MNs, FETi and SETi (Figure 7B on the right) (Azevedo et al., 2024), which we observed consistently across all leg compartments. Muscles innervated by multiple MNs in T1 were assigned several serial sets, which we pooled together for the standard leg connectome analysis in the later sections of this manuscript.

The non-serially identified MNs in T2 and T3 were either not well constructed enough to make a match to T1 MNs or differed so strongly in their connectivity or morphology that they did not resemble any the T1 MNs. Further, there were some serial sets that could not be assigned a muscle target although there was a serial set present. For two of these serial sets (13214 and 11274) we assume that the function and muscle targets between the segments is distinct, as the T2 pair of these sets are the identified STTMm neurons (Figure 7–Supplement 1B, top). The other examples are a T1 MN pair that we could not identify in FANC, in the serial set 17664 with a T2 MN pair and a serial set that only contains a T2 and T3 pair but we could not find a T1 resemblance (Figure 7–Supplement 1B, bottom).

### DN and MN systematic type naming and subclass

The systematic type is a combination of the prefix ‘DN’ or ‘MN’, the two-letter subclass (defined by the target) and a number that is consistent within the group and serial set. Smaller numbers were assigned to identified DNs or MNs, respectively.

DNs were assigned a two-letter subclass name in accordance with their VNC neuropil target. The subclasses nt, wt, ht, it, lt, fl, ml, hl and ad were given for the DNs innervating a single neuropil corresponding to the targets NTct, WTct, HTct, IntTct, LTct, LegNpT1, LegNpT2, LegNpT3 and ANm. Subsequently DNs were assigned the subclass xl or ut if they targeted a combination of leg (target: LNP) or upper tectulum neuropils (target: UT). In situations where there were inconsistencies for neuropil innervation within pairs or groups of DNs, we calculated the mean percent to determine the assignment. All DNs that innervated more than a single neuropil and not a combination of upper tectulum or leg neuropils were given the subclass xn, for multiple neuropils. In total, 490 systematic types were given to DNs.

MNs were assigned a two-letter subclass name in accordance with their target muscle or target muscle group. For all unidentified MNs, the target muscle group was predicted by the exit nerve. The two-letter subclasses stand for front (fl), middle (ml) and hind leg muscles (hl), neck (nm), wing (wm) and haltere muscles (hm) and abdominal muscles (ad). For 6 MNs, we were unsure about the muscle they target in the periphery; they were assigned the xm subclass, for unknown muscle target. In total, 273 systematic types were given to MNs.

### Connectome Analyses

The neuPrint web application (https://neuprint.janelia.org) provides a graphical front end to explore many aspects of the vnc dataset (Plaza et al., 2022). However a wide range of more sophisticated analyses were enabled by developing small scripts using the natverse suite of tools (Bates et al., 2020) built on top of the R statistical programming language (https://www.r-project.org). Connectome data was queried from MANC v1.2.3 (https://neuprint.janelia.org/?dataset=manc%3Av1.2.3&qt=findneurons) via the neuPrint API using the neuprintr package (https://natverse.org/neuprintr) (Schlegel et al., 2021). For convenience, the most common functionality was wrapped in a specialized malevnc package (https://natverse.github.io/malevnc/) which is the recommended point of entry for detailed analysis.

### Bayesian graph traversal

To complement the identification of pathways between pairs of neurons or neuronal sets using the methods above, we also considered how indirect connectivity stratifies neurons based on distance relative to a given seed set of neurons. For this we used a deterministic Bayesian variant of the graph traversal model from (Marin et al., 2024; Schlegel et al., 2021) implemented as part of the navis Python library (https://zenodo.org/records/8191725). Starting iteratively from a seed set of neurons that are immediately traversed, at each iteration neurons adjacent to the already traversed population have a chance of being traversed. Unlike adjacency matrix multiplication measures of indirect connectivity, the traversal model introduces a nonlinearity in the probability that a neuron is traversed. This probability is linear relative to the proportion of its input synapses whose presynaptic neurons have already been traversed, up to 20% of its total input, but rectified to have a 100% probability of traversal above that. The traversal analysis only considers the distribution of the iteration or “layer” when a neuron is traversed. The deterministic variant treats the traversal graph as a Bayesian network and performs exact belief propagation to determine traversal iteration distributions for each neuron. For traversal analysis from DNs, one traversal was performed for each DN group, with all neurons in that group comprising the seed set. Iteration distributions are summarized by their mean (mean layer), and MN groups are summarized as by the groupwise mean of this mean layer.

### Indirect connectivity strength

The connectome can reveal both the direct connections between any pair of neurons (or between any two groups of neurons) and the indirect pathways via one or more intermediate neurons. In many cases, indirect pathways appear stronger than direct ones; it is therefore helpful to define a measure of connectivity that can quantify indirect connections. We adopted the general approach of (Li et al., 2020). Briefly, we retrieved the all-to-all connectivity matrix for all valid VNC neurons (assigned a neuronal class in the neuPrint ‘class’ field), and normalized all synapse weights to input fractions (sum of inputs for each neuron equals 1). Then, we carried out matrix multiplication of the normalized connectivity matrix with itself for a number of times equal to the path length of interest. For analyses of DN to wing MN connectivity, indirect connectivity strength was calculated at the neuron group level (i.e. synapse counts were summed by group before normalization), with or without separation of neuron side (soma, root or nerve side). Optionally, input fractions to DNs and SNs are set to zero to avoid considering paths through them.

To compare indirect connectivity strength values calculated across different path lengths, we first normalized the values for a given source neuron and path length with their non-zero mean at the given path length. Then, for each target neuron, we compared its normalized values across all path lengths (in practice up to 5); the top value, and the path length it was found at, was taken as the neuron’s normalized indirect connectivity strength value and path length. To determine groupwise normalized indirect connectivity strength value and path length, we then calculate the mean of both these values for all neurons belonging to the same group.

Indirect connectivity strength was also used as an exploratory tool to retrieve neurons or neuron groups contributing to the connectivity between two neurons or neuron groups of interest (e.g. as an intermediary between DN and MN). Here, indirect connectivity strength was calculated from the upstream neuron to each potential intermediary neuron, and separately from the intermediary neuron to the downstream neuron, where the individual path lengths sum to the total path length of interest. The product of the upstream and downstream indirect connectivity strength was then calculated, and top-scoring neurons were considered the best candidates for intermediary neurons. To consider paths through population neuron types where individual connectivity is weak but collectively provide large synapse counts, neurons were optionally collapsed by group (sum groupwise synapse counts before conversion to input fractions) before taking matrix products. Connectivity between DNs, these intermediary neuron or group candidates and MNs were then plotted in Cytoscape via the RCytoscape R package for manual curation and graph plotting.

### Connectome pathway exploration and effective connectivity strength

We first calculated percent of the total input to the postsynaptic neuron and percent of the output budget of the presynaptic neuron for every edge in the VNC connectome. To favor connections that are strong edges for both the pre- and postsynaptic neuron in our connectome analysis we calculated the geometric mean of these two matrices. We then calculated the matrix-vector-product using the geometric mean matrix and a vector that is 1 for a pair or group of neurons of interest and is 0 for all other VNC neurons. Selecting the 20 postsynaptic neurons with the strongest edges reveals the first layer downstream of the neurons of interest. The resulting vector is then multiplied by the geometric mean matrix again to explore the second layer downstream. This step was repeated three more times to describe up to five downstream layers. After selecting the neurons involved in the circuit we used the RCy3 (version 2.17.1) and igraph (version 1.3.0) packages in R to create graphs in Cytoscape (version 3.9.1) for further exploration and visualization.

The VNC connectome matrix of percent of the total input to the postsynaptic neuron was used with the same matrix-vector-product approach to calculate effective connectivity strength between DNs of interest and leg MNs. For comparisons across the layers, we normalized by the non-zero mean of all scores in each layer. For each given DN-MN-pair we picked the highest score of the five layer matrices and recorded the layer in which it occurred. We termed this analysis effective connectivity strength.

### Quantification of contact area between neurons for evaluating putative presence of electrical synapses

To quantify area of contact between putative electrical neurons and wing MNs, we retrieved 3D meshes of all neurons of interest that were generated and simplified from EM segmentation from the MANC clio data store (https://github.com/clio-janelia/clio_website) (Perlman, 2019) using the malevnc R package. Using a custom python script in Blender version 3.4 (https://www.blender.org), all neuron meshes were expanded by 100 nm using the blender shrink/fatten function to create mesh intersection areas between closely-situated areas for pairs of neurons (up to 200 nm apart, but with inaccuracies as neuron meshes are simplified from the underlying segmentation). Then, for each pair of neurons, we calculated pairwise mesh intersections using a hole-tolerant exact method and calculated the area of the intersection mesh, divided by half to obtain the mean of the two surfaces contributed by each neuron mesh.

## Reagents

**Table 3:**
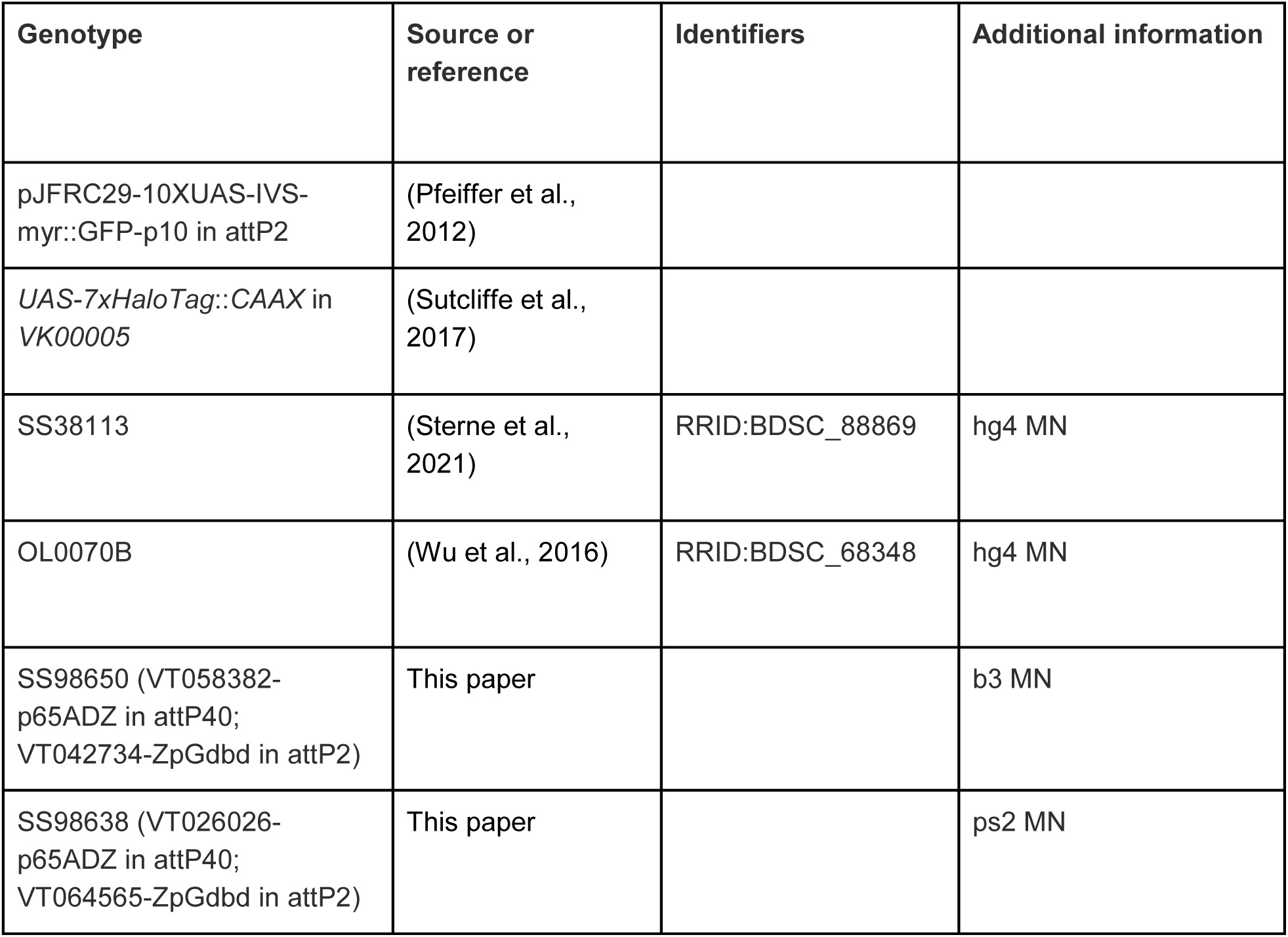
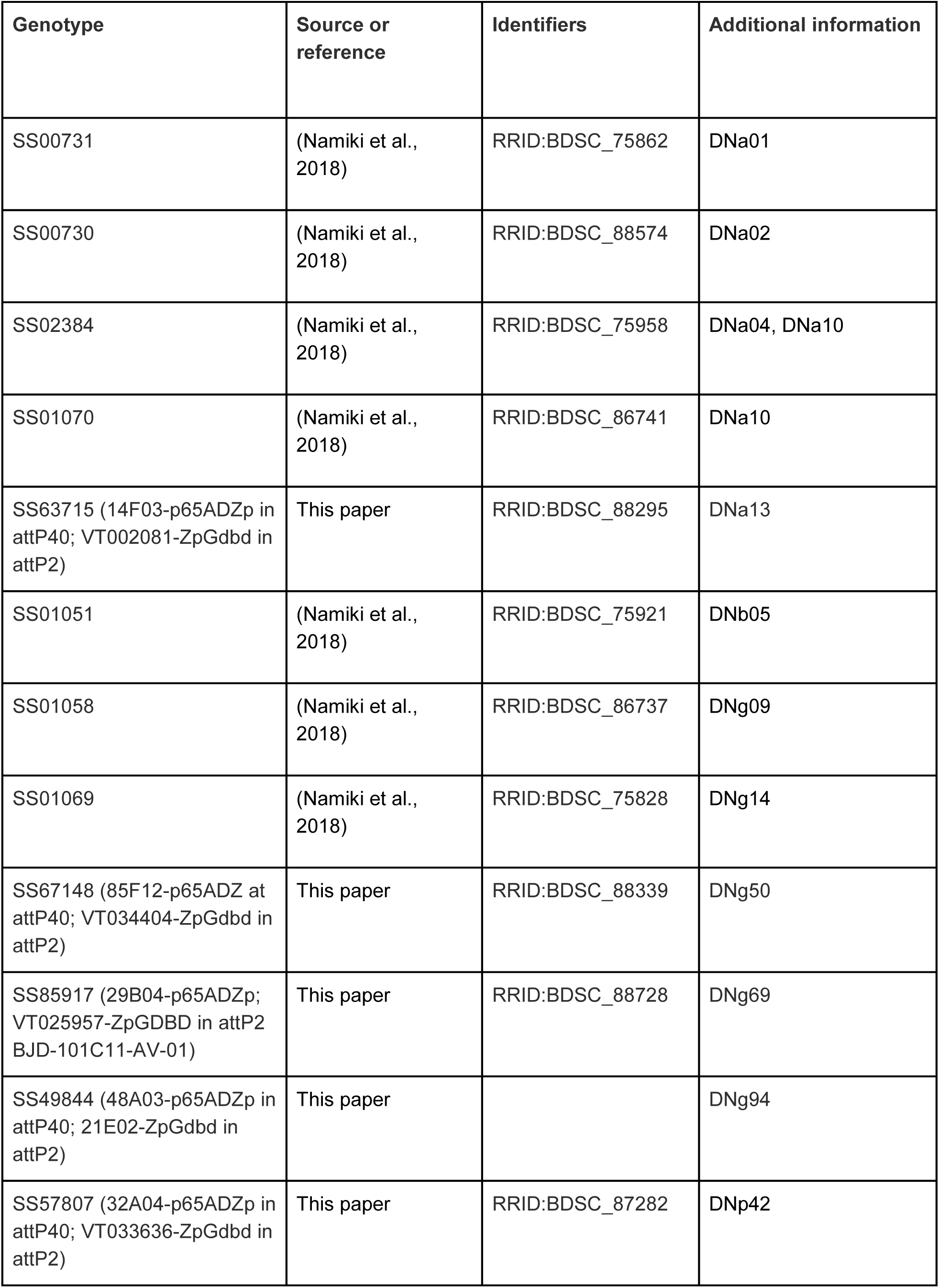
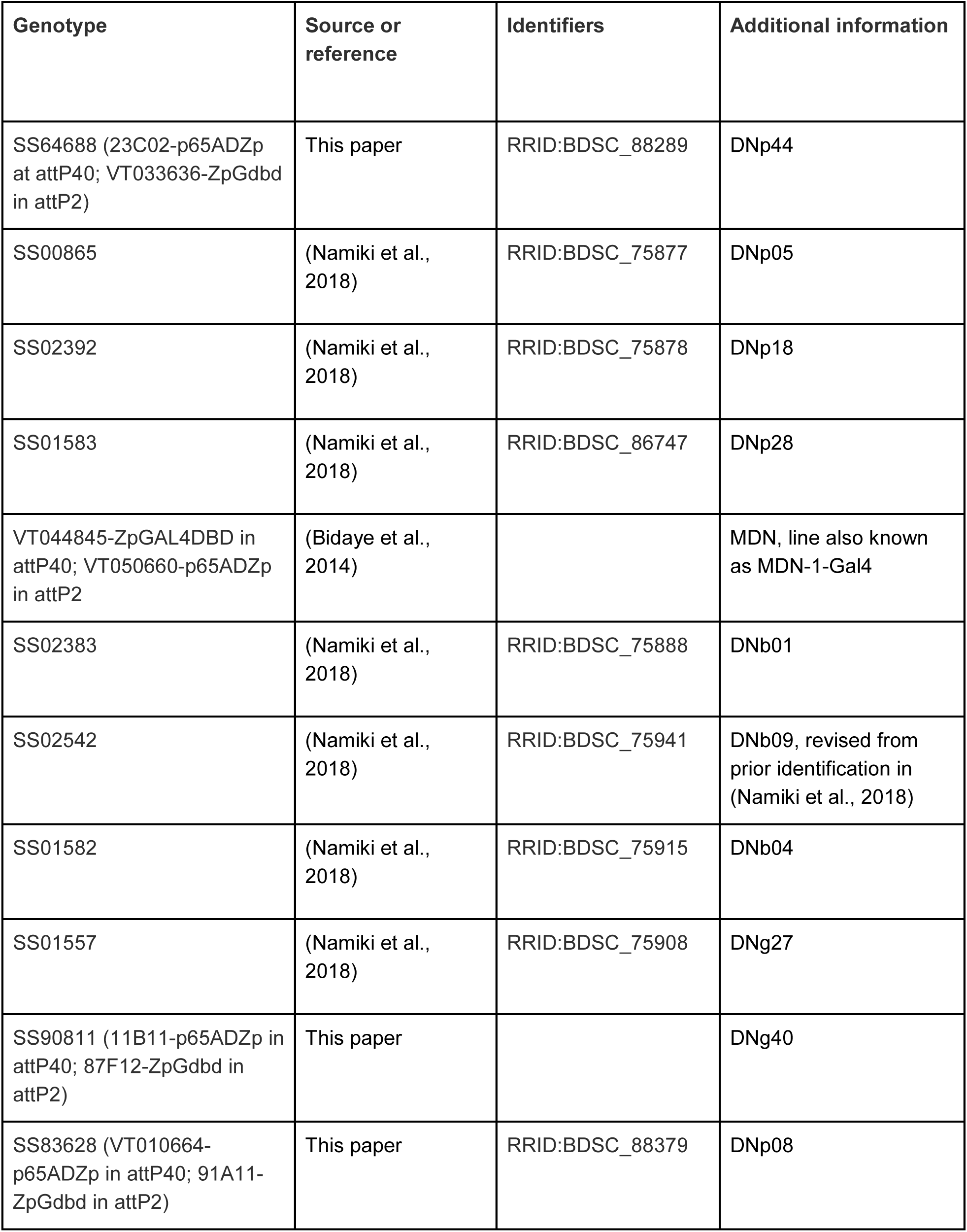
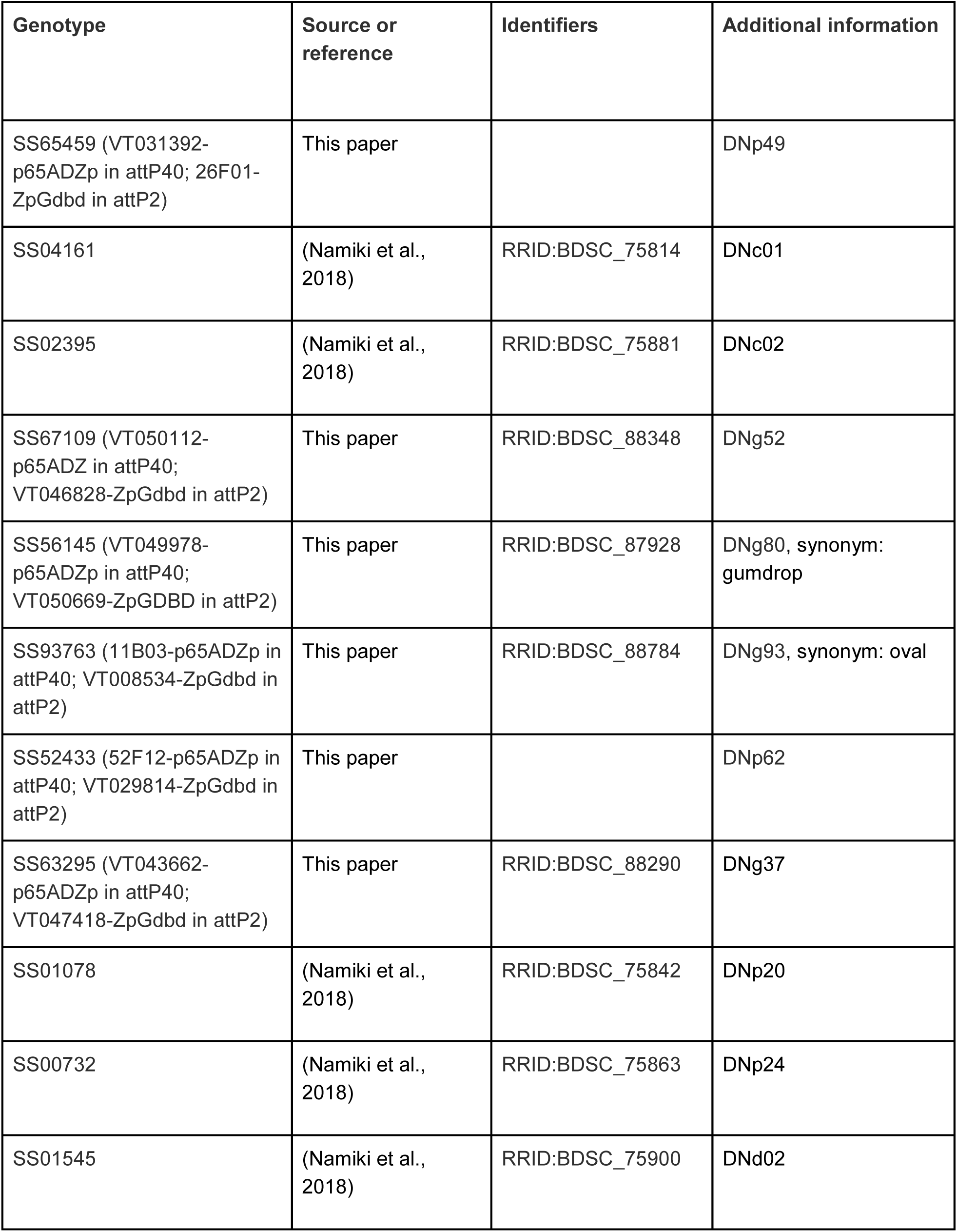
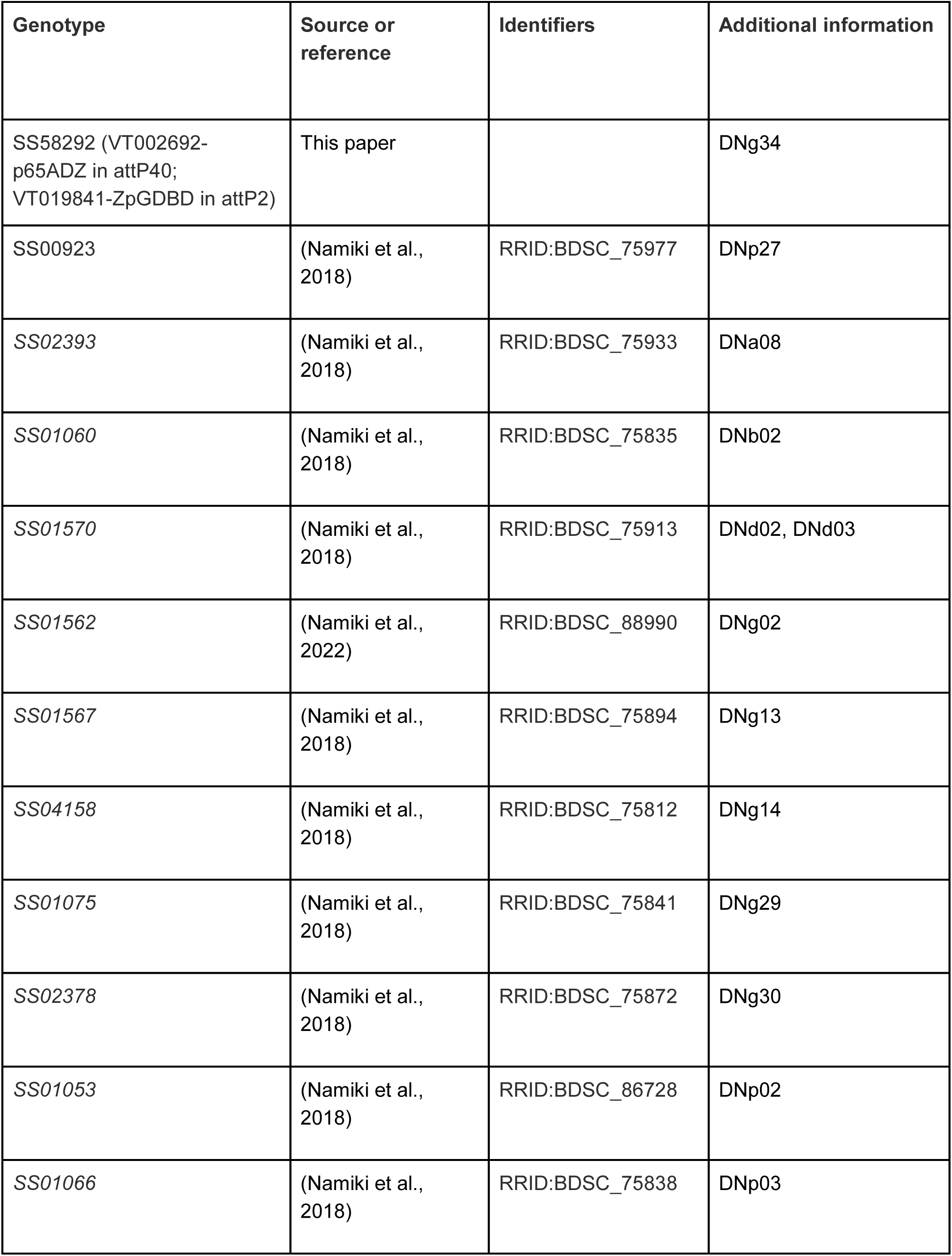
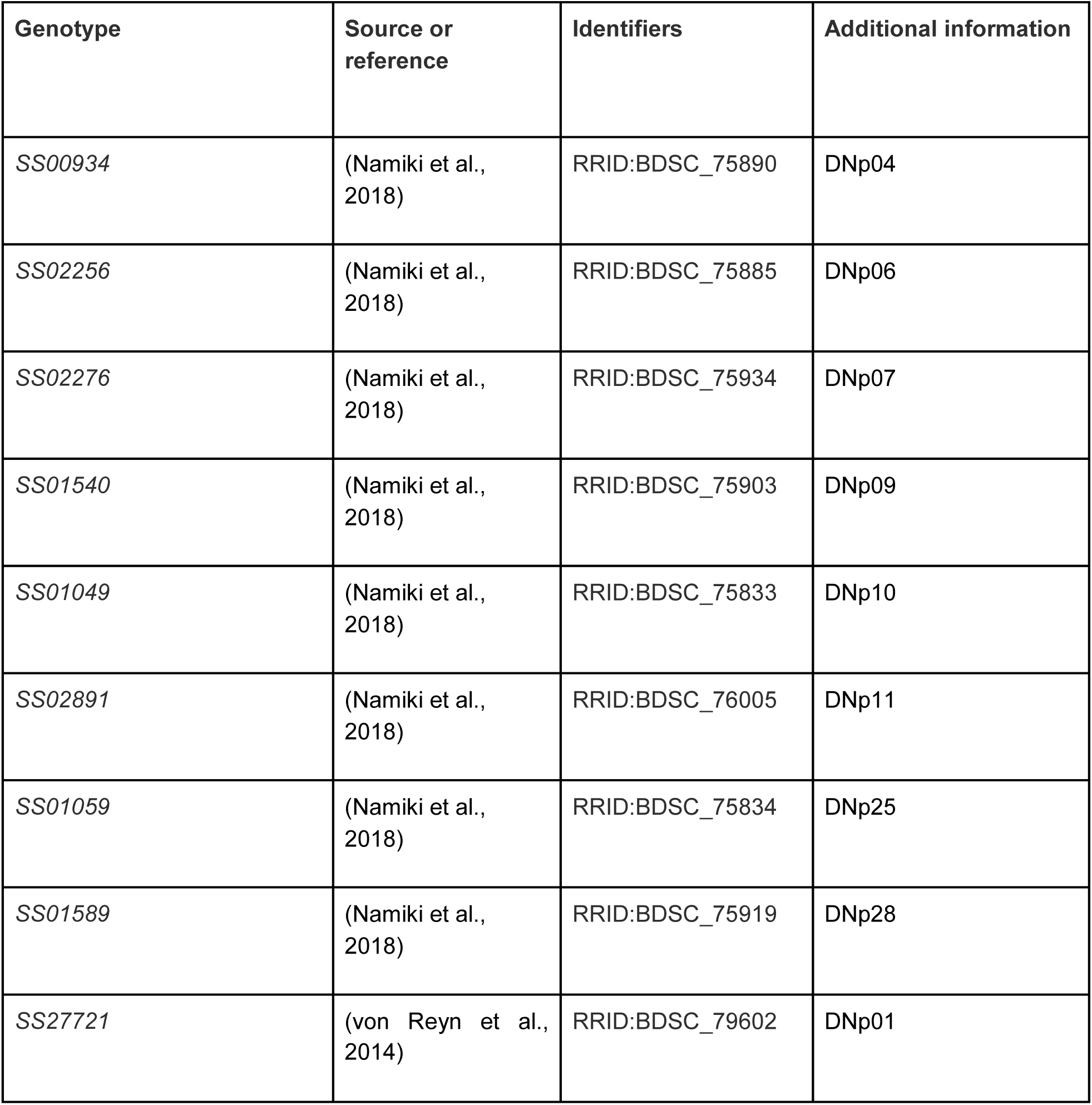
*Drosophila* lines used in this study. Imagery for newly generated split-Gal4 combinations may be found at https://splitgal4.janelia.org/precomputed/Cheong,%20Eichler,%20Stuerner%202023.html

| Genotype | Source or reference | Identifiers | Additional information |
| --- | --- | --- | --- |
| pJFRC29-10XUAS-IVS-myr::GFP-p10 in attP2 | (Pfeiffer et al., 2012) |  |  |
| *UAS-7xHaloTag::CAAX* in *VK00005* | (Sutcliffe et al., 2017) |  |  |
| SS38113 | (Sterne et al., 2021) | RRID:BDSC_88869 | hg4 MN |
| OL0070B | (Wu et al., 2016) | RRID:BDSC_68348 | hg4 MN |
| SS98650 (VT058382-p65ADZ in attP40; VT042734-ZpGdbd in attP2) | This paper |  | b3 MN |
| SS98638 (VT026026-p65ADZ in attP40; VT064565-ZpGdbd in attP2) | This paper |  | ps2 MN |
| SS00731 | (Namiki et al., 2018) | RRID:BDSC_75862 | DNa01 |
| SS00730 | (Namiki et al., 2018) | RRID:BDSC_88574 | DNa02 |
| SS02384 | (Namiki et al., 2018) | RRID:BDSC_75958 | DNa04, DNa10 |
| SS01070 | (Namiki et al., 2018) | RRID:BDSC_86741 | DNa10 |
| SS63715 (14F03-p65ADZp in attP40; VT002081-ZpGdbd in attP2) | This paper | RRID:BDSC_88295 | DNa13 |
| SS01051 | (Namiki et al., 2018) | RRID:BDSC_75921 | DNb05 |
| SS01058 | (Namiki et al., 2018) | RRID:BDSC_86737 | DNg09 |
| SS01069 | (Namiki et al., 2018) | RRID:BDSC_75828 | DNg14 |
| SS67148 (85F12-p65ADZ at attP40; VT034404-ZpGdbd in attP2) | This paper | RRID:BDSC_88339 | DNg50 |
| SS85917 (29B04-p65ADZp; VT025957-ZpGDBD in attP2 BJD-101C11-AV-01) | This paper | RRID:BDSC_88728 | DNg69 |
| SS49844 (48A03-p65ADZp in attP40; 21E02-ZpGdbd in attP2) | This paper |  | DNg94 |
| SS57807 (32A04-p65ADZp in attP40; VT033636-ZpGdbd in attP2) | This paper | RRID:BDSC_87282 | DNp42 |
| SS64688 (23C02-p65ADZp at attP40; VT033636-ZpGdbd in attP2) | This paper | RRID:BDSC_88289 | DNp44 |
| SS00865 | (Namiki et al., 2018) | RRID:BDSC_75877 | DNp05 |
| SS02392 | (Namiki et al., 2018) | RRID:BDSC_75878 | DNp18 |
| SS01583 | (Namiki et al., 2018) | RRID:BDSC_86747 | DNp28 |
| VT044845-ZpGAL4DBD in attP40; VT050660-p65ADZp in attP2 | (Bidaye et al., 2014) |  | MDN, line also known as MDN-1-Gal4 |
| SS02383 | (Namiki et al., 2018) | RRID:BDSC_75888 | DNb01 |
| SS02542 | (Namiki et al., 2018) | RRID:BDSC_75941 | DNb09, revised from prior identification in (Namiki et al., 2018) |
| SS01582 | (Namiki et al., 2018) | RRID:BDSC_75915 | DNb04 |
| SS01557 | (Namiki et al., 2018) | RRID:BDSC_75908 | DNg27 |
| SS90811 (11B11-p65ADZp in attP40; 87F12-ZpGdbd in attP2) | This paper |  | DNg40 |
| SS83628 (VT010664-p65ADZp in attP40; 91A11-ZpGdbd in attP2) | This paper | RRID:BDSC_88379 | DNp08 |
| SS65459 (VT031392-p65ADZp in attP40; 26F01-ZpGdbd in attP2) | This paper |  | DNp49 |
| SS04161 | (Namiki et al., 2018) | RRID:BDSC_75814 | DNc01 |
| SS02395 | (Namiki et al., 2018) | RRID:BDSC_75881 | DNc02 |
| SS67109 (VT050112-p65ADZ in attP40; VT046828-ZpGdbd in attP2) | This paper | RRID:BDSC_88348 | DNg52 |
| SS56145 (VT049978-p65ADZp in attP40; VT050669-ZpGDBD in attP2) | This paper | RRID:BDSC_87928 | DNg80, synonym: gumdrop |
| SS93763 (11B03-p65ADZp in attP40; VT008534-ZpGdbd in attP2) | This paper | RRID:BDSC_88784 | DNg93, synonym: oval |
| SS52433 (52F12-p65ADZp in attP40; VT029814-ZpGdbd in attP2) | This paper |  | DNp62 |
| SS63295 (VT043662-p65ADZp in attP40; VT047418-ZpGdbd in attP2) | This paper | RRID:BDSC_88290 | DNg37 |
| SS01078 | (Namiki et al., 2018) | RRID:BDSC_75842 | DNp20 |
| SS00732 | (Namiki et al., 2018) | RRID:BDSC_75863 | DNp24 |
| SS01545 | (Namiki et al., 2018) | RRID:BDSC_75900 | DNd02 |
| SS58292 (VT002692-p65ADZ in attP40; VT019841-ZpGDBD in attP2) | This paper |  | DNg34 |
| SS00923 | (Namiki et al., 2018) | RRID:BDSC_75977 | DNp27 |
| SS02393 | (Namiki et al., 2018) | RRID:BDSC_75933 | DNa08 |
| SS01060 | (Namiki et al., 2018) | RRID:BDSC_75835 | DNb02 |
| SS01570 | (Namiki et al., 2018) | RRID:BDSC_75913 | DNd02, DNd03 |
| SS01562 | (Namiki et al., 2022) | RRID:BDSC_88990 | DNg02 |
| SS01567 | (Namiki et al., 2018) | RRID:BDSC_75894 | DNg13 |
| SS04158 | (Namiki et al., 2018) | RRID:BDSC_75812 | DNg14 |
| SS01075 | (Namiki et al., 2018) | RRID:BDSC_75841 | DNg29 |
| SS02378 | (Namiki et al., 2018) | RRID:BDSC_75872 | DNg30 |
| SS01053 | (Namiki et al., 2018) | RRID:BDSC_86728 | DNp02 |
| SS01066 | (Namiki et al., 2018) | RRID:BDSC_75838 | DNp03 |
| SS00934 | (Namiki et al., 2018) | RRID:BDSC_75890 | DNp04 |
| SS02256 | (Namiki et al., 2018) | RRID:BDSC_75885 | DNp06 |
| SS02276 | (Namiki et al., 2018) | RRID:BDSC_75934 | DNp07 |
| SS01540 | (Namiki et al., 2018) | RRID:BDSC_75903 | DNp09 |
| SS01049 | (Namiki et al., 2018) | RRID:BDSC_75833 | DNp10 |
| SS02891 | (Namiki et al., 2018) | RRID:BDSC_76005 | DNp11 |
| SS01059 | (Namiki et al., 2018) | RRID:BDSC_75834 | DNp25 |
| SS01589 | (Namiki et al., 2018) | RRID:BDSC_75919 | DNp28 |
| SS27721 | (von Reyn et al., 2014) | RRID:BDSC_79602 | DNp01 |

## Author contributions

Conceptualization: HSJC, KE, TS, SA, AC, GSXEJ, and GMC

Methodology: HSJC, KE, TS, and SA

Software: AC and GSXEJ

Validation: HSJC, KE, TS, ECM, MC, SB, Janelia FlyEM team, GSXEJ, GMC

Formal analysis: HSJC, KE, TS, SA, AC, MS, TO, and GSXEJ

Investigation: HSJC, KE, TS, SA, AC and TO

Resources: ECM, AC, SN, SB, Janelia FlyEM team, GSXEJ

Data curation: HSJC, KE, TS, AC, ECM, LV, MC, and GSXEJ

Writing - original draft: HSJC, KE, TS, SA, GSXEJ, and GMC

Writing - review and editing: HSJC, KE, TS, SA, GSXEJ, and GMC

Visualization: HSJC, KE, TS, SA, AC, and IS

Supervision: MC, GSXEJ and GMC

Project administration: MC, SB, GSXEJ and GMC

Funding acquisition: GSXEJ and GMC

## Data Availability

The MANC connectome, including annotations of DNs and MNs described in this work, are available at https://neuprint.janelia.org. Key annotations for DNs and MNs, as well as match information to neurons identified at the light level, are available as supplementary files. Code used for analyses and to generate figures in this work are available upon request from the corresponding authors.

## Notes

https://neuprint.janelia.org/?dataset=manc:v1.2.1&qt=findneurons

### Competing Interest Statement

The authors have declared no competing interest.

### Summary of Updates

This version of the manuscript has been revised in response to reviewer comments from eLife.

